# A synergistic interaction between HDAC- and PARP inhibitors in childhood tumors with chromothripsis

**DOI:** 10.1101/2021.04.22.440879

**Authors:** Umar Khalid, Milena Simovic, Murat Iskar, John KL Wong, Rithu Kumar, Manfred Jugold, Martin Sill, Michiel Bolkestein, Thorsten Kolb, Michaela Hergt, Frauke Devens, Jonas Ecker, Marcel Kool, Till Milde, Frank Westermann, Joe Lewis, Sascha Dietrich, Stefan M Pfister, Peter Lichter, Marc Zapatka, Aurélie Ernst

## Abstract

Chromothripsis is a form of genomic instability characterized by the occurrence of tens to hundreds of clustered DNA double-strand breaks in a one-off catastrophic event. Rearrangements associated with chromothripsis are detectable in numerous tumor entities and linked with poor prognosis in some of these, such as Sonic Hedgehog medulloblastoma, neuroblastoma and osteosarcoma. Hence, there is a need for therapeutic strategies eliminating tumor cells with chromothripsis. Defects in DNA double-strand break repair, and in particular homologous recombination repair, have been linked with chromothripsis. Targeting DNA repair deficiencies by synthetic lethality approaches, we performed a synergy screen using drug libraries (n = 375 compounds, 15 models) combined with either a PARP inhibitor or cisplatin. This revealed a synergistic interaction between the HDAC inhibitor romidepsin and PARP inhibition. Functional assays, transcriptome analyses, and *in vivo* validation in patient-derived xenograft mouse models confirmed the efficacy of the combinatorial treatment.

## INTRODUCTION

Chromothripsis is a type of genomic instability where tens to hundreds of clustered DNA double-strand breaks occur in a presumably single catastrophic event^1,2^. Rearrangements caused by chromothripsis have been detected in a number of tumor types^3,4^, with varying prevalence rates, reaching up to 100% in specific molecular subgroups^1^. In a subset of these tumor types, chromothripsis was associated with poor prognosis^5–7^. To date no specific therapy targeting chromothriptic cells has been developed and these patients would greatly benefit from more effective therapeutic options. This is especially true in certain tumor entities where the current treatment modalities show limited success, such as in medulloblastoma with mutant *TP53*, where the identification of novel synthetic lethal interactions is highly relevant.

We and others identified a strong link between chromothripsis and DNA double-strand break repair, and in particular homologous recombination deficiency^8,9^. Notably, homologous recombination deficiency may provide vulnerabilities to specifically eliminate tumor cells, as it potentially confers sensitivity to poly (ADP-ribose) polymerase (PARP) inhibition^10,11^. Clinical trials in breast and ovarian cancer first showed the benefits of synthetic lethality between *BRCA*-mutant tumors and PARP inhibition^12,13^. Later, this approach was effectively applied to a range of additional tumor types such as pancreas, bone, skin, and lung cancer, including not only tumors with *BRCA* mutations but also tumors with molecular signatures of BRCAness^14^. In analogy to the successful use of PARP inhibitors in homologous recombination deficient cancers, DNA repair defects of tumor cells with chromothripsis may be exploited to target these cells by synthetic lethality approaches.

Several studies have highlighted the benefit of combinatorial screens to identify synergistic interactions amenable to use for cancer treatment. In Ewing sarcoma, Heske and colleagues used a screening approach to discover a synergistic combination of PARP inhibitors and nicotinamide phosphoribosyltransferase inhibitors^15^. In a chemical screen of DNA damage signaling inhibitors, Henssen and colleagues identified AZD6738 for therapeutic targeting of PGBD5-induced DNA repair dependency in pediatric solid tumors^16^. However, to our knowledge, compound screens in chromothriptic cells have not been performed yet.

In Sonic Hedgehog medulloblastoma with mutant *TP53* as well as in neuroblastoma and osteosarcoma, the prevalence of chromothripsis is high and chromothripsis is linked with poor prognosis^5,6,9,17^. Therefore, focusing on these three childhood solid tumors, we set off to identify vulnerabilities of chromothriptic cells by screening for drugs that kill tumor cells with chromothripsis but spare control cells with functional repair capacity.

## RESULTS

### Screen for drugs that inhibit the metabolic activity of chromothripsis-driven tumors

We searched for compounds that eliminate chromothriptic cells as single agents or as synergistic partners with a PARP inhibitor (BGB-290) or with cisplatin, a chemotherapeutic agent commonly used in medulloblastoma, neuroblastoma, and osteosarcoma. Our library was composed of 375 compounds screened at four concentrations each (5 nM, 50 nM, 500 nM, and 5 μM), with or without addition of PARP inhibitor or cisplatin (Figure 1a, b). This resulted in 750 pairwise drug combinations covering a wide range of targets and processes implicated in cancer biology. The read-out of this primary screen was the metabolic activity of the cells after 96 hours of treatment as compared to control cells with solvent only.

**Figure 1:**
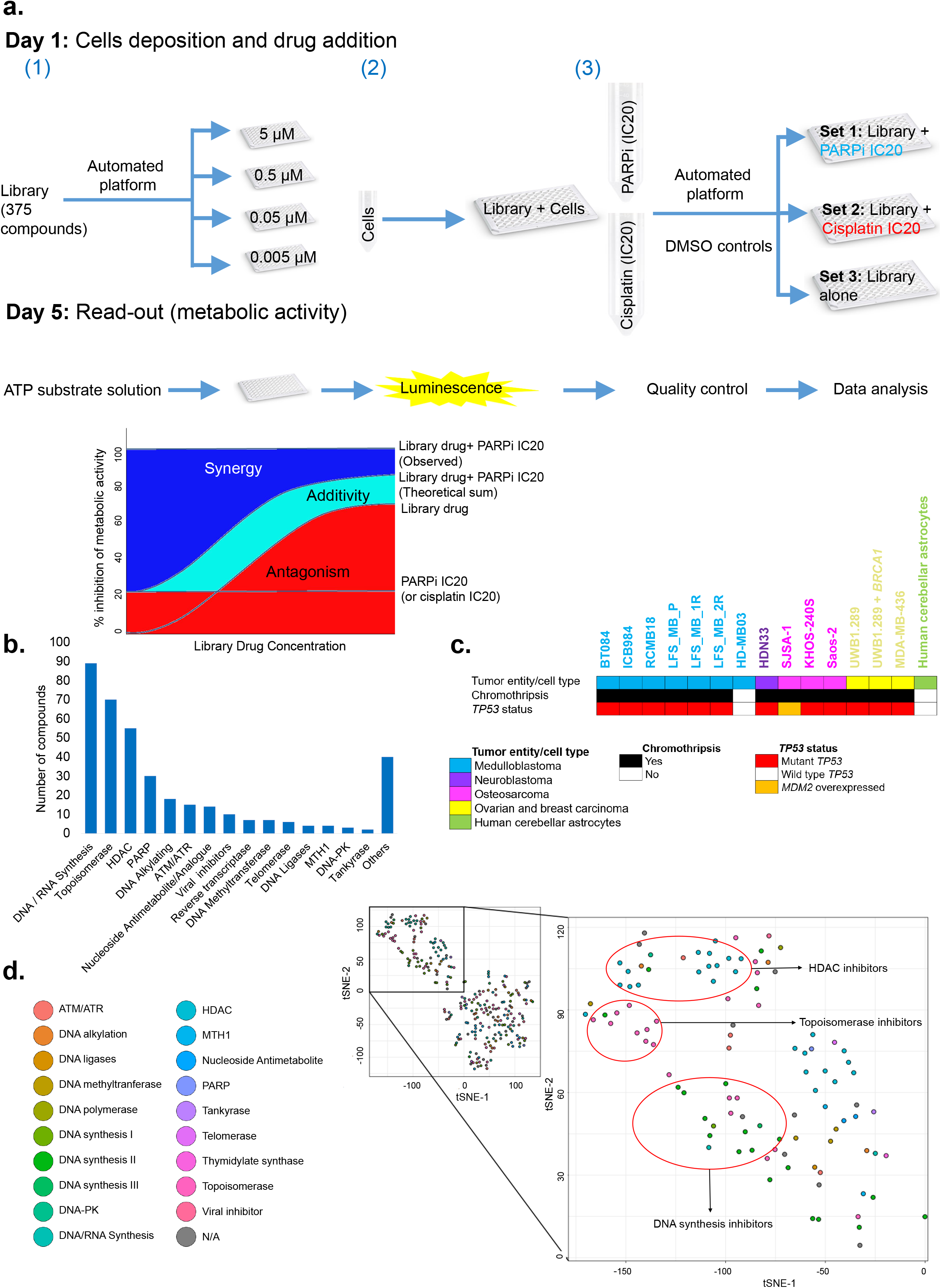
Overview of the primary screen: (a) Schematic representation of the primary screen strategy and graphical representation of synergistic and antagonistic drug interactions. (b) Classification of the library drugs on the basis of their target or mode of action. (c) Characterization of the cell types and tumor entities included in the primary screen. (d) t-distributed stochastic neighbor embedding (t-SNE) plot based on the primary screen data.

We performed the primary screen with a panel of 15 cell lines including medulloblastoma, neuroblastoma and osteosarcoma lines, as well as control cell lines, comprising normal cells (astrocytes), homologous recombination deficient breast and ovarian cancer lines, and non-chromothriptic medulloblastoma cells (Figure 1c). To use representative medulloblastoma models, we established spheroid cultures freshly isolated from patient-derived xenografts. All lines included in the screen were molecularly characterized by methylation arrays and/or whole-genome sequencing (Supplementary Figure 1), except for three models that were previously published^18,19^.

We first examined the quality control parameters, including for instance positive controls leading to complete inhibition of the metabolic activity, and negative controls showing maximal metabolic activity (Supplementary Figure 2a). In addition, we evaluated the response to drugs that were contained in two independent libraries, and therefore were represented twice in the screen, which showed high correlation coefficients (R > 0.88, Supplementary Figure 2b). Finally, comparison of control cell lines analyzed at the start of the screen and repeated in the end showed that the vast majority of the drugs displayed a high stability over time, with only fewer than 10 drugs having a significant loss of activity (Supplementary Figure 2c).

### Synergy screen identifies inhibitors of tumor cells with chromothripsis

We started the data analysis by clustering the drugs based on the similarity of their response profiles across samples (Figure 1d). The t-SNE analysis showed that the profiles of the drugs reflected their mode of action and their target identity. For instance, the response to different types of HDAC inhibitors, topoisomerase inhibitors or DNA synthesis inhibitors was highly correlated across cell lines. In addition, clustering showed disease-specific drug responses, with the major branches in the dendrogram reflecting tumor entities (Supplementary Figure 3a).

We searched for drugs that were highly effective, i) across all tumor entities, ii) entity-specific iii), as single compounds, and iv) in combination with BGB-290 or cisplatin (Figure 2). Compounds were scored as active if they inhibited cell metabolic activity by at least twofold relative to the control (DMSO). Compounds showing additive effects or potential for synergy were rare (Figure 2d, Supplementary Figure 3b). Among the most effective compounds showing high efficiency at low concentrations were the HDAC inhibitor romidepsin, the DNMT3B inhibitor nanaomycin, and the iron chelator Dp44mt. Notably, the class I HDAC inhibitor romidepsin and the DNMT3B inhibitor nanaomycin were more potent than doxorubicin or cyclophosphamide that are used in the clinic for these tumor types. Romidepsin was approved for treatment of lymphoma ^20^ and is in clinical trials for additional indications but is not used in the clinic in the context of medulloblastoma (MB), neuroblastoma or osteosarcoma. Few trials have been conducted using nanaomycin.

**Figure 2:**
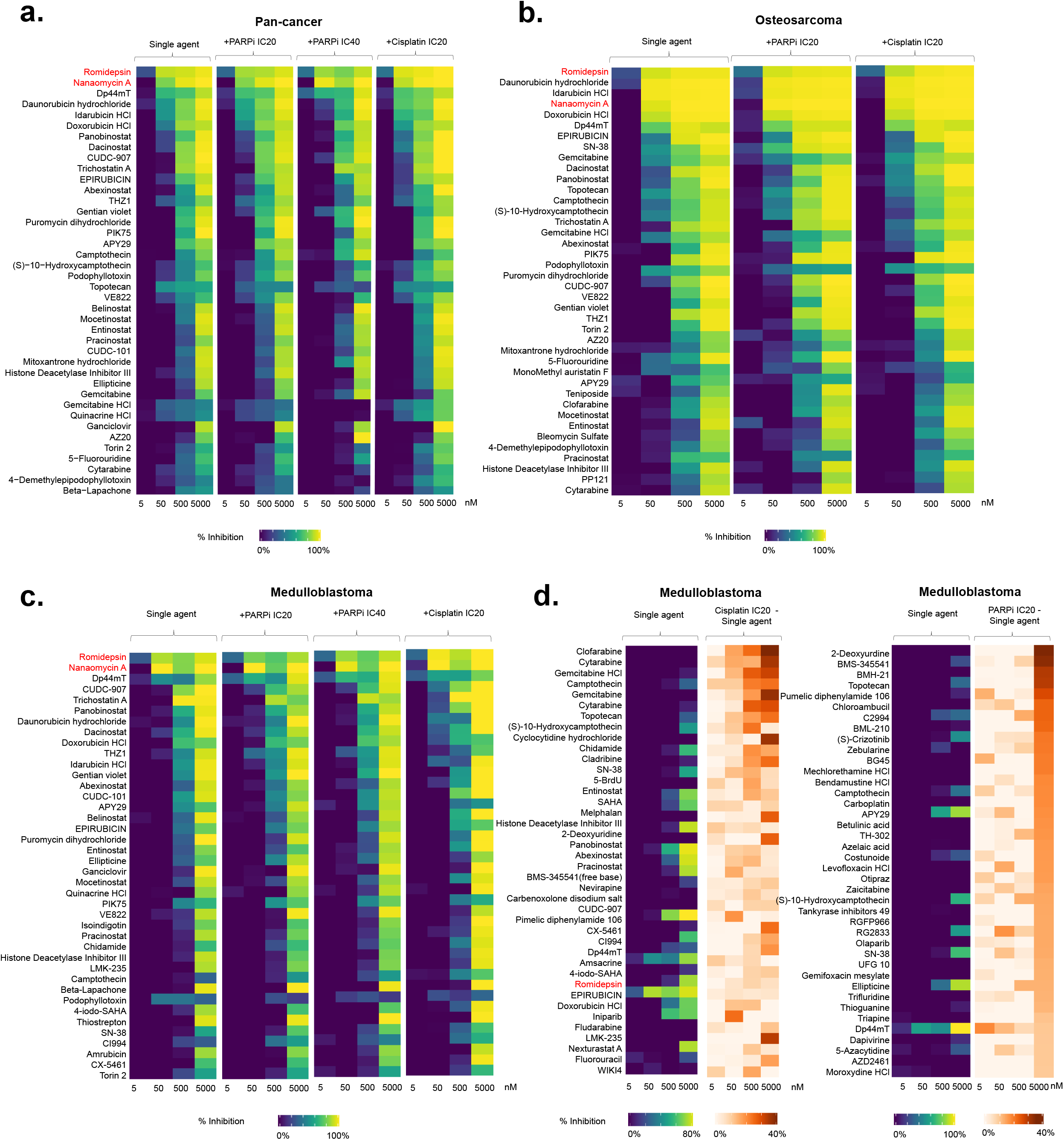
Analysis of the primary screen data: Heatmaps showing the top 40 hits based on the following criteria (a) Maximum overall potency using all tumor lines included in the primary screen (n = 14 lines, non-tumor cells were excluded). (b) Maximum overall potency focusing on osteosarcoma cell lines; KHOS-240S, SJSA-1 and SaOS-2. (c) Maximum overall potency focusing on Sonic Hedgehog medulloblastoma patient-derived xenograft spheroid models (n = 4 lines, for the LFS_MB model only the primary tumor was included here). (a - c) Data are shown as percentages of inhibition of the metabolic activity as compared to the DMSO controls. (d) Synergy with cisplatin IC20 and synergy with BGB-290 IC20 focusing on Sonic Hedgehog medulloblastoma patient-derived xenograft spheroid models.

Having observed potent inhibitory effects (nanomolar potency) of romidepsin and nanaomycin on chromothriptic cells, we then wanted to distinguish bona fide synergy from additive effects. For this, we performed a secondary screen with 6 medulloblastoma models using an 8 by 8 matrix of concentrations where we evaluated the combination effects using the Loewe model (Figure 3a, Supplementary Figure 4). As half of the tumor cell lines included in the primary screen were medulloblastoma cultures, we focused on this tumor type for the secondary screen and the *in vivo* validation. For nanaomycin, no synergy was detected with BGB-290 or with cisplatin. For romidepsin, we detected a strong synergy between romidepsin and BGB-290 in three models, namely LFS_MB_P (primary tumor), LFS_MB_1R (matched relapse from the same patient), and RCMB18 (Figure 3b). Interestingly, there was no synergy or even antagonistic effects in non-chromothriptic medulloblastoma cells, HD-MB03. Phenotypically, the decrease in the metabolic activity in medulloblastoma cells with chromothripsis was accompanied by a massive reduction in the size of the spheroids after combinatorial treatment (Figure 3c, Supplementary Figure 3c). For ICB984 and BT84, additive effects were observed at higher romidepsin concentrations only (> 7.8 × 10^-4^ μM, Supplementary Figure 4).

**Figure 3:**
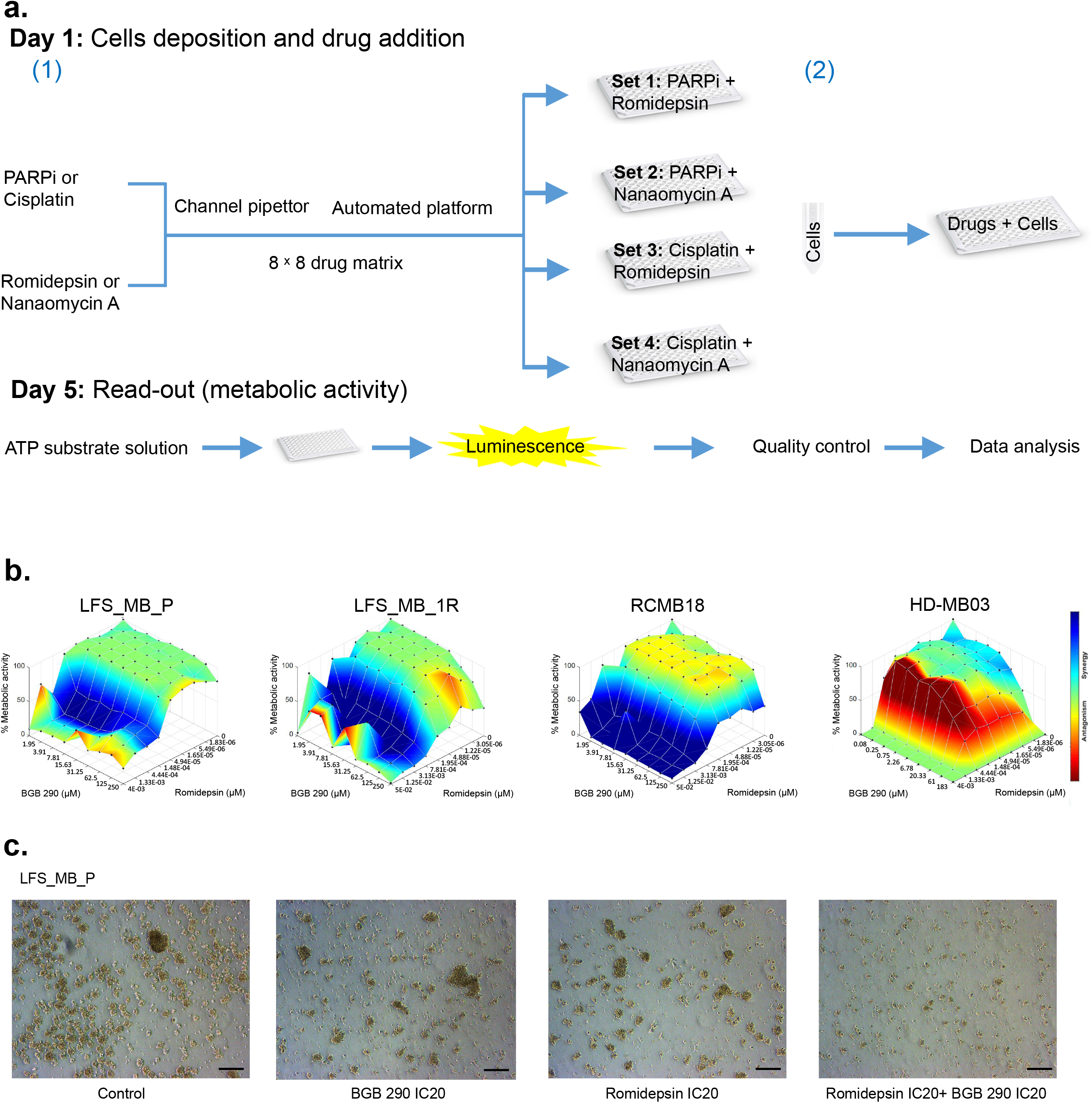
Overview and analysis of the secondary screen: (a) Schematic representation of the secondary screen strategy. (b) Loewe surface plots generated using the combenefit software showing synergistic and antagonistic drug interactions. Synergy and antagonism are plotted on the dose-response curves using the average values for the metabolic activity (percentage from DMSO controls, 3 technical replicates per data point). (c) Bright-field images of spheroids (one representative Sonic Hedgehog medulloblastoma model) after 24 hours of drug treatment. Scale bar in each panel: 100 μm.

### Romidepsin and BGB-290 lead to a p53-independent G2/M arrest

We next investigated the mode of action of romidepsin in combination with BGB-290. First, we tested whether the effects on the metabolic activity were linked with changes in the cell cycle distribution. Due to the strong association of chromothripsis with compromised p53 response and challenges to find efficient treatments in p53-deficient backgrounds, we first tested the p53 dependency of the effects using isogenic wild-type and *TP53* inactivated RPE1 cells (Figure 4). We observed a similar response for RPE1 WT and *TP53* KO cells. Single treatments led to a significant increase in the G2/M fraction and combination treatment induced a strong G2/M arrest, independent of the p53 status. In RPE1 cells, the treatment did not induce apoptosis, as shown by annexin/7AAD staining (Supplementary Figure 5).

**Figure 4:**
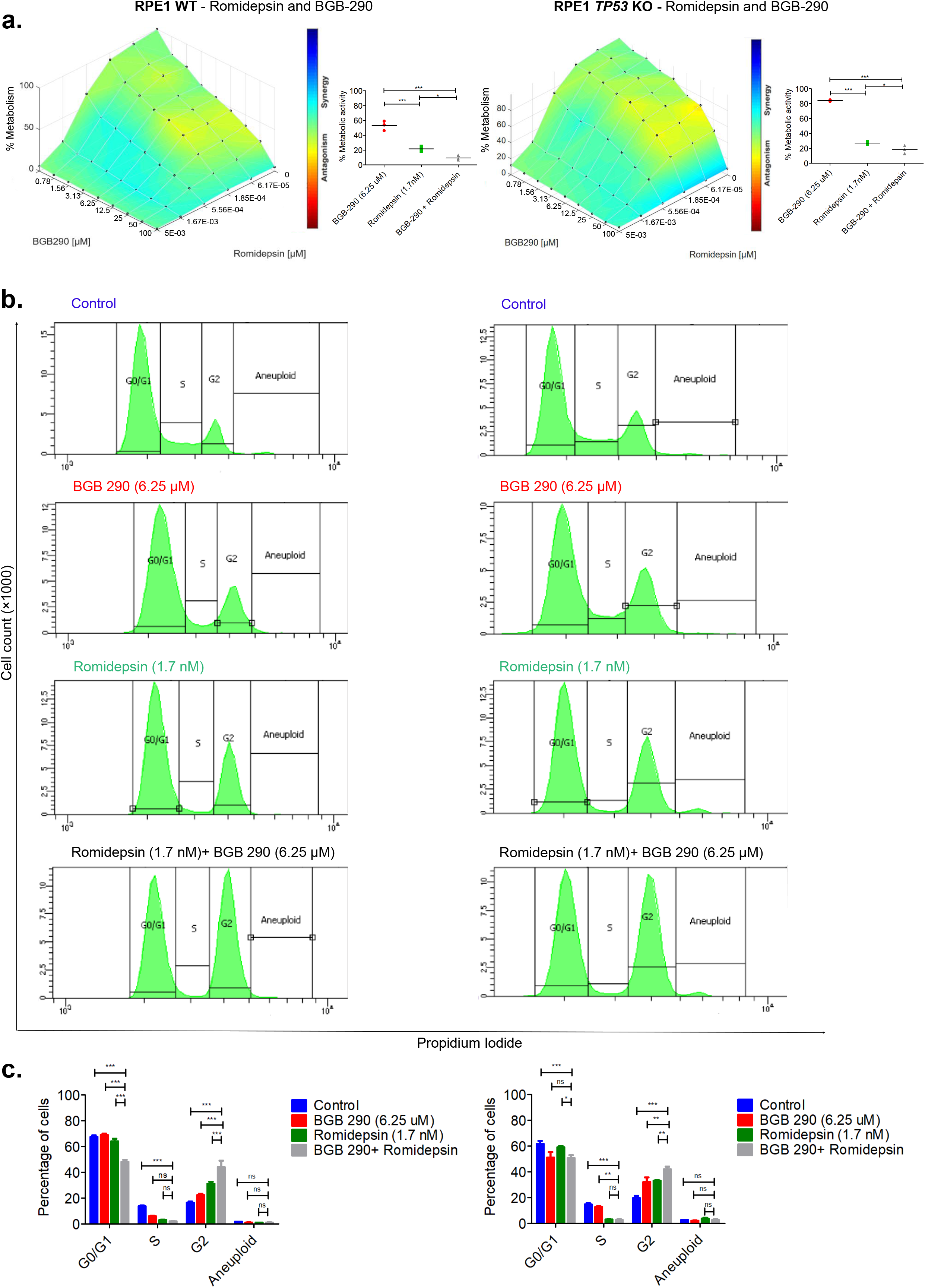
Cell cycle arrest of RPE1 WT and RPE1 *TP53* KO cells upon combination treatment with romidepsin and BGB-290: (a) Loewe surface plot for RPE1 WT cells (on the left) and *TP53* KO RPE1 cells (on the right). Plots were generated using combenefit software. Synergy and antagonism are plotted on the dose-response values using the average percentages of the metabolic activity values for 3 biological replicates per data point. Dot plots on the right of each Loewe surface plot show the metabolic activity values (as percentages of the DMSO controls) for 3 biological replicates. Statistical significance was tested by one-way ANOVA followed by Tukey’s Multiple Comparison Test (*, p < 0.05). Horizontal lines represent the means, n=3 biological replicates. (b) Cell cycle analysis of RPE1 WT cells (on the left) and RPE1 *TP53* KO cells (on the right) using propidium iodide. Cells were treated with drugs at the indicated concentrations for 48 hours before the flow cytometry analysis. Histograms are representative of three biological replicates. (c) Bar graphs showing the quantitative distribution of RPE1 WT cells (on the left) and *TP53* KO RPE1 cells (on the right) in the different cell cycle phases. Statistical significance was tested using two-way ANOVA followed by Bonferroni post-test, *** p < 0.001, ** p < 0.01, * p < 0.05, non-significant (ns) p > 0.05. Error bars represent SEM, n = 3 biological replicates.

To analyze the mode of action of the combinatorial treatment in medulloblastoma cells with chromothripsis, we used DAOY cells (Figure 5). The effects on the cell cycle distribution were much stronger as compared to the ones observed in non-tumor RPE1 cells, with a massive SubG1 apoptotic fraction in addition to the G2/M block. The significant induction of apoptosis in DAOY cells was confirmed by annexin/7AAD stains. Therefore, combining BGB-290 with romidepsin in medulloblastoma cells leads to a G2 arrest and apoptosis induction.

**Figure 5:**
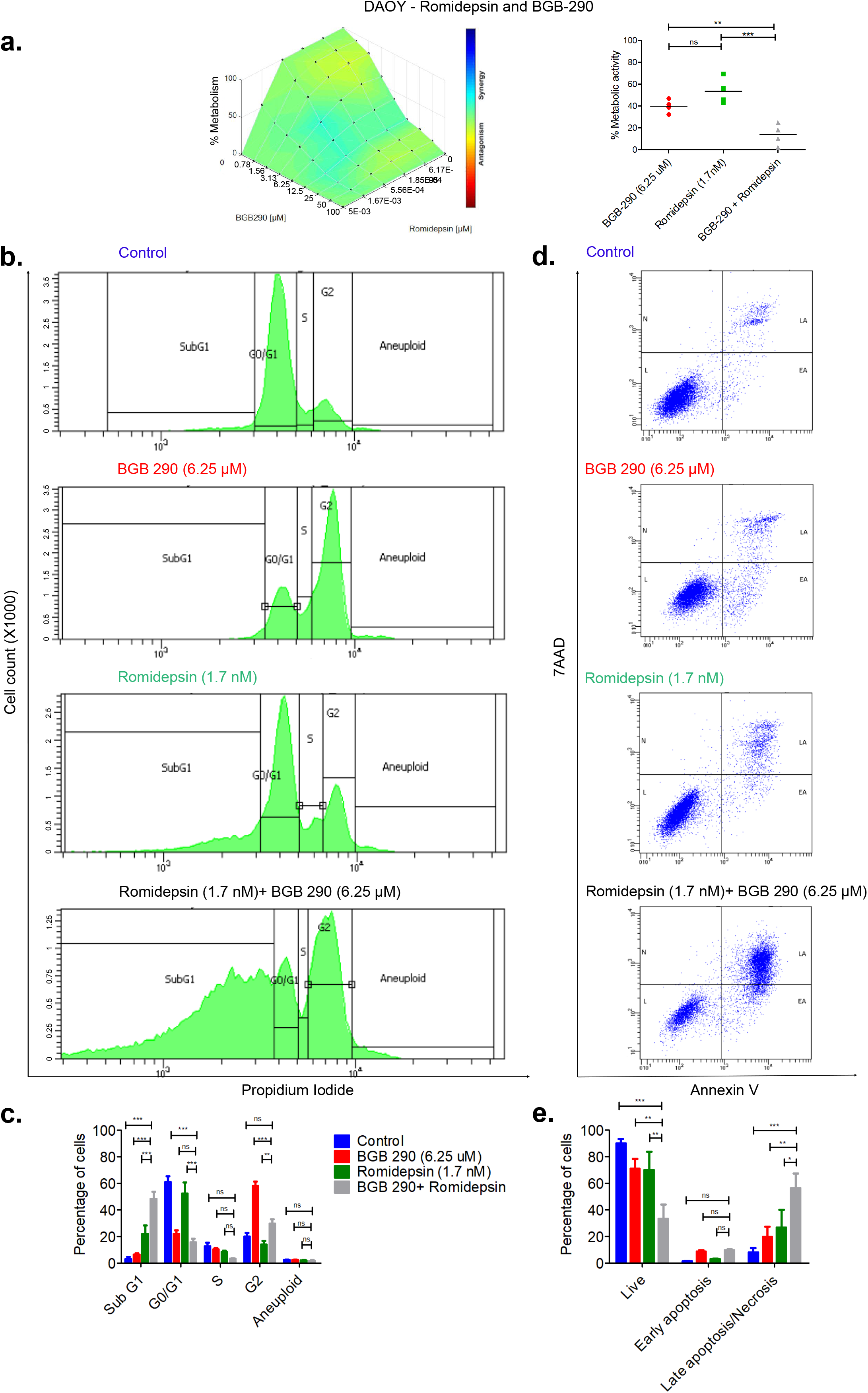
Cell cycle arrest and apoptosis of DAOY cells upon combination treatment with romidepsin and BGB 290: (a) Loewe surface plot for DAOY cells. Plot was generated using the combenefit software. Synergy and antagonism are plotted on the dose-response values using the average of the metabolic activity values (as percentages of the DMSO controls) for 3 biological replicates per data point. Dot plot on the right of the Loewe surface plot shows the metabolic activity for 3 biological replicates at the indicated drug concentrations. Statistical significance was tested by one-way ANOVA followed by Tukey’s Multiple Comparison Test (*, p < 0.05). Horizontal lines represent the means, n = 3 biological replicates. (b) Cell cycle analysis of DAOY cells using propidium iodide. Cells were treated with drugs at the indicated concentrations for 48 hours and subjected to flow cytometry analysis. Histograms are representative of three biological replicates. (c) Bar graph showing the quantitative distribution of DAOY cells in the different cell cycle phases. Statistical significance was tested by two-way ANOVA followed by Bonferroni post-test, *** p<0.001, ** p<0.01, * p<0.05, ns p>0.05. Error bars represent SEM, n=3 biological replicates (ns = non-significant). (d) Apoptotic fractions in DAOY cells were analyzed by FITC-annexin V and 7AAD stains. Cells were treated with drugs at the indicated concentrations for 48 hours and subjected to flow cytometry analysis. Dot plots are representative of three biological replicates. (e) Bar graph showing the quantitative distribution of DAOY cells as living, early apoptotic and late apoptotic/ necrotic. Statistical significance was tested by two-way ANOVA followed by Bonferroni post-test, *** p<0.001, ** p<0.01, * p<0.05, ns p>0.05. Error bars represent SEM, n = 3 biological replicates.

To test the specificity of the inhibitory effect to chromothriptic cells, we compared the response to the treatment to non-chromothriptic medulloblastoma cells (HD-MB03) and to normal cells (BJ cells, normal astrocytes and RPE1 cells). Although all cells were sensitive to the drugs and no cell line was completely unaffected, the sensitivity for non-chromothriptic cells and for normal cells was lower than for chromothriptic medulloblastoma cells. For instance, at a 5 nM concentration of romidepsin, the average inhibition of the metabolic activity of chromothriptic medulloblastoma cells was 1.8-fold higher than for normal cells and non-chromothriptic cells. Altogether, these data suggested that combining romidepsin with BGB-290 inhibits the survival of chromothriptic MB cells and might be a useful therapeutic agent for the development of treatments for these aggressive tumors, with potential activity in additional subgroups as well.

### Molecular effects of romidepsin and BGB-290 on medulloblastoma cells

To gather further insights into the molecular effects of the combination treatment on chromothriptic medulloblastoma cells, we studied the mitotic process by visualizing phospho histone H3 and acetylated tubulin via immunofluorescence (Figure 6a). BGB-290 alone did not lead to any significant change in the mitotic activity of DAOY cells. In contrast, romidepsin treatment and combination treatment induced a significant decrease in the mitotic index, as shown by quantification of mitotic cells.

**Figure 6:**
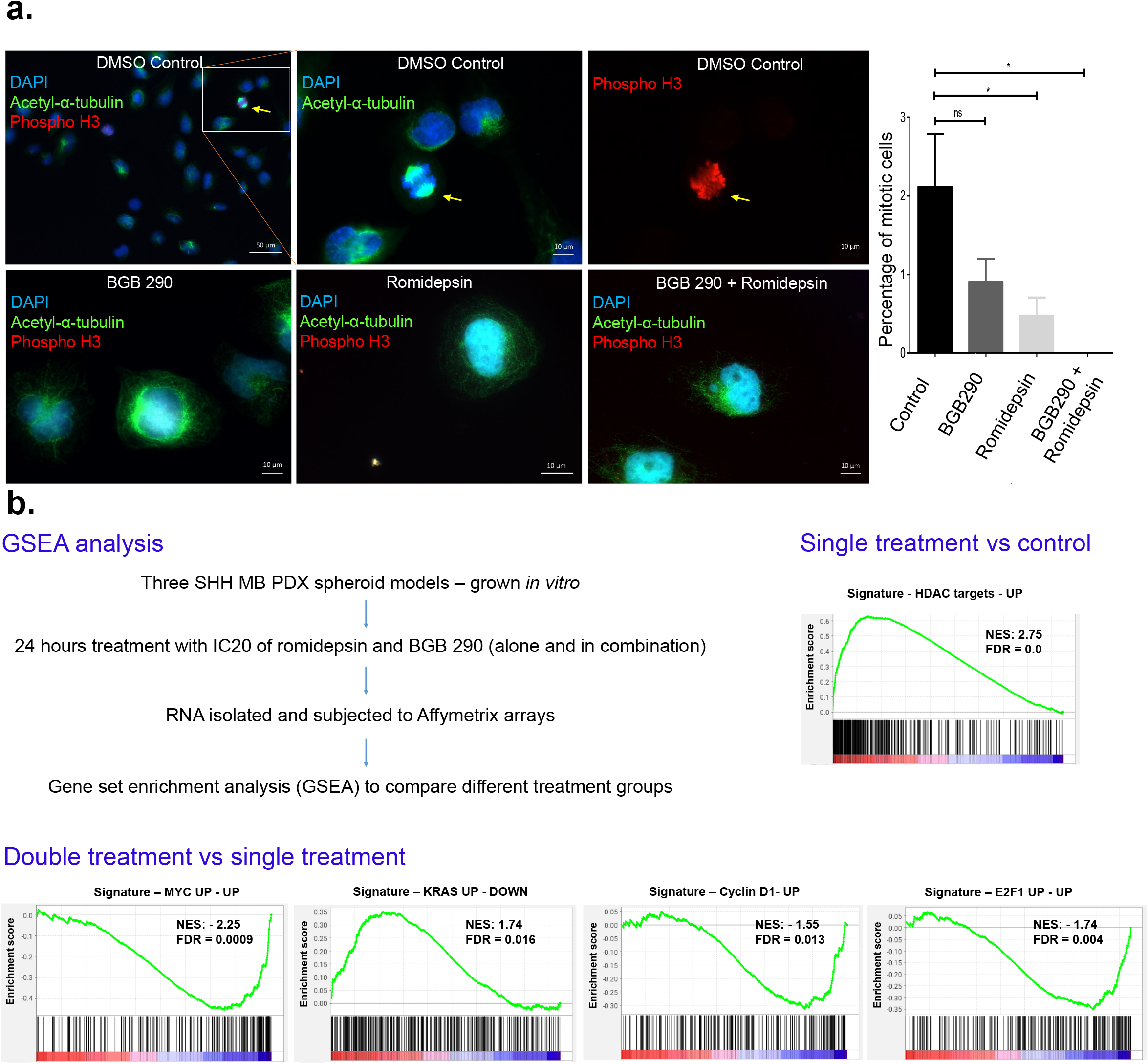
Mode of action of romidepsin and BGB 290: (a) Acetyl-α-tubulin and phospho-histone 3 immunostaining of Daoy cells treated with either vehicle control, 6.25 μM BGB 290, 1.7 nM romidepsin or 6.25 μM BGB-290 with 1.7 nM romidepsin for 48 hours. Scale bar in top left panel: 50 μm. Scale bar in all other panels: 10 μm. Bar graph showing the percentage of mitotic cells in each treatment group. Statistical significance was tested by one-way ANOVA followed by Dunnett’s Multiple Comparison Test, * p < 0.05, ns p > 0.05. Error bars represent the SEM, n= 3 biological replicates. A minimum of 250 cells were counted for each group in each replicate. (b) Schematic representation of the transcriptome data experiment (top left). Top signatures from GSEA analysis from single treatment compared to control (top right, romidepsin versus control) and double treatment compared to single treatment (bottom).

We performed expression profiling using Affymetrix Arrays to get insights into the mechanisms by which romidepsin and BGB-290 suppress growth of chromothripsis-driven MB (Figure 6b, Supplementary Figure 6). MB spheroids from three patient-derived models were incubated with DMSO or drugs for 24 hours, before RNA was isolated for microarray analysis. For each replicate, treatment with romidepsin or combinatorial treatment resulted in a marked change in gene expression, whereas single treatment with BGB-290 did not lead to major transcriptional effects. Using the criteria of a fold change > 1.5 and p < 0.01, we found 115 genes differentially expressed (DE) between the single treatment and the combination treatment. The complete list of DE genes is presented in Table S1. To gain insights into the pathways potentially regulated by romidepsin and BGB-290, we investigated the DE genes using GSEA (Figure 6b). Single treatment with romidepsin was linked with signatures of HDAC targets, as expected. Importantly, signatures specifically linked with combinatorial treatment but not with single treatment included proliferation related genes and pathways, such as *MYC*, *KRAS*, *Cyclin D1* or *E2F1*. To test the role of MYC target genes in the combination treatment, we used a MYC-inducible tumor cell line (Figure 7). Without induction of MYC by doxycycline, romidepsin and BGB-290 showed antagonistic effects, further supporting a possible role for MYC in the synergistic effect.

**Figure 7:**
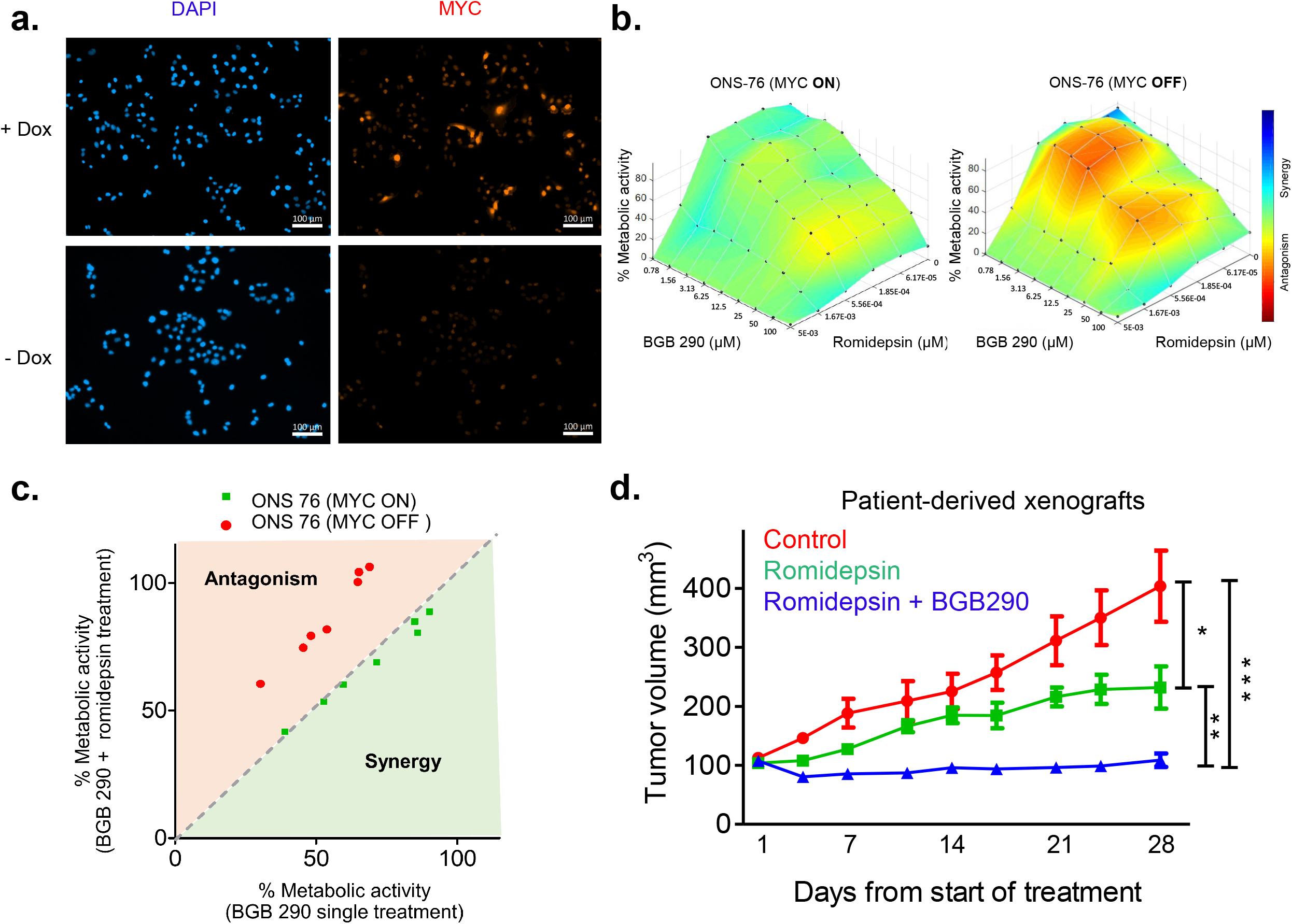
Romidepsin and BGB-290 synergistic effects are linked to *MYC* expression and *in vivo* validation in patient-derived xenografts: (a) MYC immunostaining of ONS76 (*MYC* ON/OFF) cells with and without doxycyclin induction. Scale bar in each panel: 100 μm. (b) Loewe surface plot for ONS76 (*MYC* ON/OFF) cells with and without doxycyclin induction. The plot was generated using the combenefit software. Synergy and antagonism are plotted on the dose-response values using the average of the metabolic activity values (percentage of the DMSO controls) for 3 biological replicates per data point. (c) Scatter plot of the average of the metabolic activity values (percentage of the DMSO controls) for 3 biological replicates per data point for ONS76 (*MYC* ON/OFF) cells with and without doxycyclin induction. The plot was generated with BGB 290 single treatment on the X-axis versus BGB-290 with romidepsin (0.56 nM) double treatment on the Y-axis. (d) Line graph showing the evolution of the tumor volume for patient-derived xenografts in immunocompromised mice. The statistical significance was tested by one-way ANOVA followed by Tukey’s Multiple Comparison Test (* p < 0.05). Error bars represent the SEM, 6 mice per treatment group.

### HDACi and PARPi synergize to inhibit the growth of chromothriptic medulloblastoma *in vivo*

In light of the potent inhibitory effects of romidepsin and BGB-290 *in vitro*, we tested the efficacy of these compounds *in vivo* in subcutaneous patient-derived xenograft models (Figure 7d). Animals were subjected to weekly measurements of tumor volume, and when tumors were clearly detectable (in the range of 100 mm^3^), mice with equivalent tumor volumes were randomized into three treatment groups: vehicle, romidepsin or romidepsin with BGB-290. Animals were treated for a total of four weeks and mice treated with vehicle showed rapid tumor growth (Figure 7d). Treatment with romidepsin slowed down tumor growth, but did not lead to any inhibition of the tumor progression. The combination of romidepsin and BGB-290 significantly inhibited tumor growth, with no detectable increase in tumor volume, suggesting that this might be an effective combination therapy for this highly aggressive form of MB.

## DISCUSSION

Our combinatorial screen across 15 models identified a strong synergy between the HDAC inhibitor romidepsin and the PARP inhibitor BGB-290. Functional assays showed a G2 arrest and apoptosis upon combination treatment. Furthermore, transcriptome analyses revealed MYC target genes as significantly downregulated upon the combination treatment. Finally, we confirmed the efficacy of romidepsin and BGB-290 *in vivo* in PDX mouse models.

Due to the tight link between p53 dysfunction and chromothripsis^1^, the question of the p53 dependency of the effects induced by the double treatment arises. As we observed similar effects in wild-type and *TP53* knock-out RPE1 cells, this suggests a p53 independent effect of the combinatorial treatment. Searching for synthetic lethal interactions in chromothriptic cells, it might seem counterintuitive to identify non-p53 specific hits, which might be of potential relevance in a p53 wild-type background as well. In *TP53* mutant tumors, it is notoriously challenging to find efficient therapeutic strategies. Beyond our initial goal of identifying drug combinations to eliminate chromothriptic cells, a subset of the hits identified in this screen may possibly be applicable outside the p53 deficient context. Therefore, it will be essential to carefully evaluate the specificity of the compounds in a range of genetic backgrounds.

Similarly, the chromothripsis dependency will need to be further investigated. Although we included non-chromothriptic cells as controls in this screen (non-chromothriptic medulloblastoma cells and non-tumor cells), it will be critical to dissect the effects of the combinatorial treatment in a large range of chromothriptic and non-chromothriptic models. Chromothriptic cells show marked DNA repair defects, which motivated our strategy. However, it is conceivable that a subset of the identified compound hits may eliminate non-chromothriptic tumor cells as well as chromothriptic cells. More generally, the specificity to the analyzed tumor types or the potential pan-cancer effects of these compounds may be interesting to consider. Finally, as romidepsin is a class I HDAC inhibitor, but a number of other histone inhibitors (e.g. entinostat, belinostat, panobinostat, dacinostat, abexinostat) also appear in the top hits from the screen (see Figure 2), it will be important to distinguish whether the synergy with PARP inhibition is a compound-specific or a (class I) specific effect.

The transcriptome analyses identified MYC targets as down-regulated upon the combinatorial treatment, with a putative role for MYC in the downstream effects confirmed in a MYC-inducible model. As *MYC* and *MYCN* are frequently amplified and overexpressed in the three tumor types investigated in this screen, it will be important to further investigate the precise role of these factors in the treatment response. The HDACi entinostat was shown to inhibit MYC transcriptional activity in *MYC* amplified MB^21^, in line with the downregulation of MYC target genes observed here upon combination treatment. Interestingly, a previous study identified HDAC and Pi3K antagonists to inhibit *MYC*-driven medulloblastoma^19^. Even though there are ongoing trials combining HDAC with PARPi and HDAC inhibitors are used in pediatric oncology, neither romidepsin nor BGB-290 has been used in medulloblastoma or in other pediatric solid tumors. Therefore, this study bears the potential to develop strategies to exploit the combinatorial effects of these two inhibitors for therapeutic purposes.

## METHODS

### Cell culture

Spheroid cultures from patient-derived xenograft models were prepared by homogenizing the freshly isolated tumors from mice brains by pipetting in NeuroCult^™^ NS-A Proliferation Kit (STEMCELL Technologies, 05751). Homogenized tissue is then filtered to get single cell suspension. Cells were then centrifuged at 1000 rpm for 5 minutes and pellet was resuspended to get 0.5 million cells per 10 ml growth medium. Growth medium was comprised of 46.75 ml Neurobasal™-A Medium (Invitrogen, 10888022) with 46.75 ml DMEM/F-12 (Invitrogen, 11330057), 1 ml MEM Non-Essential Amino Acids Solution (100X) (Invitrogen, 11140050), 1 ml Sodium Pyruvate (100 mM) (Invitrogen, 11360070), 2.5 ml Hepes buffer (1M) (Invitrogen, 15630080), 1 ml GlutaMAX^™^ Supplement (Invitrogen, 35050061), 1 ml Penicillin-Streptomycin (Invitrogen, 15140122), 2 ml B-27^™^ Supplement minus vitamin A (50X) (Invitrogen, 12587001), 40 μl Heparin (50 mg/ml stock) (Sigma, H3149-10KU), 20ng/ml Human FGF rec 154aa (Peprotech, GMP100-18 B), 10 ng/ml EGF (Sigma, E 4127) and 100 μl LIF(Millipore, LIF1010). Spheroids were cultured for 3 weeks in 25 cm^2^ flask (Greiner, 690195) and passaged by trypsinization with 0.05% Trypsin/EDTA once a week. All adherent cells were passaged by trypsinization with 0.25% Trypsin/EDTA. Astrocytes and spheroids were passaged by trypsinization with 0.05% Trypsin/EDTA.

Formulations of culture media for cell lines used in this study were the following: For DAOY, BJ WT and BJ *TP53* KO cells, 1X DMEM high glucose (Gibco, 41965-039) with 10% fetal calf serum (FCS) and 1% L-glutamine. For UWB1.289 and UWB1.289+*BRCA1* cells, 48.5 % RPMI Medium 1640 (Gibco, 21875-034) and 48.5% MEGM media (Lonza, CC-3150) (Supplements were added provided with the kit) with 3% FCS. For UWB1.289+BRCA1, G418 was added to final concentration of 200 μg/ml. For MDA-MB-436 cells, 1X DMEM high glucose (Gibco, 41965-039) with 10% FCS. For KHOS-240S cells, 1X DMEM high glucose (Gibco, 41965-039) with 10% FCS, 1% L-glutamine and 1% MEM Non-essential amino acids (MEM NEAA) (Gibco, 11140-050). For SaOS-2 cells, Roti®-CELL McCoy’s 5A (Carl Roth, 9111.1) with 15 % FCS and 1% L-glutamine. For SJSA-1 cells, RPMI Medium 1640 (Gibco, 21875-034) with 10% FCS and 1% L-glutamine. For HD-MB-03 cells, RPMI Medium 1640 (Gibco, 21875-034) with 10% Heat inactivated FCS, 1% L-glutamine and 1% MEM NEAA (Gibco, 11140-050). For HD-N33 cells, RPMI Medium 1640 (Gibco, 21875-034) with 10 % FCS and 1% L-glutamine. For Normal Astrocytes, astrocyte media (ScienCell, 1801) (Supplements were added provided with the kit). Culture flasks and 96 well plates were coated with poly-L-lysine (Sigma, P1399). For RPE1 WT and RPE1 *TP53* KO, DMEM/F-12 1:1 Medium (Gibco, 11320-074) with 10% FCS and 1% L-glutamine.

### Molecular characterization of the cell lines and PDX models

Copy number states from EPIC and 450K methylation arrays were assessed by the Bioconductor package conumee. For EPIC methylation arrays and 450K methylation arrays, references were taken from the GEO database with accession GSE147740 and GSE68777 respectively. Copy number states were computed per candidate gene by comparing the cell lines with references. Copy number changes were reported on genes with more than 0.15 in log2 fold change. For whole genome sequencing data, copy number analysis was performed using ACESeq. Copy number calls from ACESeq were used to determine the copy number states.

### Primary screen

For each cell line included in the screen (UWB1.289, UWB1.289+*BRCA1*, MDA-MB-436, HD-MB03, HD-N33, normal human astrocytes, SaOS-2, KHOS-240S, SJSA-1, LFS_MB_P (primary tumor), LFS_MB_1R (first relapse), LFS_MB_2R (second relapse), RCMB18, BT084 and ICB984) the IC20 and IC40 values of BGB 290 (Pamiparib, MedChemExpress, HY-104044) and cisplatin (MedChemExpress, HY-17394) were determined. The 375 compounds from 2 drug libraries, namely TargetMol (Catalog No. L3900) and DiscoveryProbe (ApexBio, L1033) were diluted in 96-well plates (Perkin Elmer, 6005689 for adherent cells and Thermo Scientific, 236105 for spheroids) to achieve final concentrations of 5 μM, 0.5 μM, 0.05 μM and 0.005 μM. Cells were then seeded at optimized densities. Each cell line or tumor entity was treated with its respective IC20 concentration of BGB 290 or cisplatin. Spheroids from patient-derived xenograft models were treated with both IC20 and IC40 of BGB 290. Cells were incubated at 37 °C for 96 hours. The metabolic activity was measured after 96 hours with the ATPlite assay (Perkin Elmer, 6016947). Values from the blank measurements were subtracted from the treatment wells and normalized to the vehicle controls. 10% DMSO treatment was used as positive control as measure of 100% metabolic inhibition whereas vehicle DMSO concentration was used as negative control as measure of 0% metabolic inhibition. The effect of single treatments was compared to the combination treatments to identify drugs that have potential additive or synergistic effects with BGB-290 or cisplatin or both. Data from the primary screen was analyzed using a shiny app developed for this purpose.

### Analysis of Primary screen data

A custom R/Shiny^22,23^app was developed for the interactive visualization and analysis of the high-throughput drug screening data. For each cell model, drug-sensitivity profiles were calculated across replicate experiments by averaging the percentage of drug-induced inhibition compared to controls as measured by the metabolic activity of the cells. Hierarchical clustering with ward.D linkage and Pearson’s correlation distance was performed to group 15 cell models (independently for single agent or combination treatments) according to their drug-sensitivity profiles and the resulting cluster was represented as dendrogram. Based on the sensitivity profiles across 15 cell models under single agent or combination treatments, drugs with similar profiles were visualized using t-distributed stochastic neighbour embedding (t-SNE)^24^ with the parameters theta=0, max_iter=20000 and perplexity=10. R packages ‘NMF’^25^ and ‘ComplexHeatmap’ in R/Shiny^26^ were utilized to create heatmaps of drug-sensitivity profiles. Interactive scatter plots were generated with the R ‘plotly’ package^27^.

### Secondary screen

Romidepsin (medchemexpress, HY-15149) and nanaomycin A (Biotrend, A8191) were candidate hits selected from the analysis of the primary screen data to be subjected to a secondary screen. Spheroids from 5 patient-derived xenograft models (LFS_MB_P, LFS_MB_1R, RCMB18, BT084, ICB984) and HD-MB03 cells were seeded in 96-well plates (Perkin Elmer, 6005689 for adherent cells and Thermo Scientific, 236105 for spheroids). Cells were then treated with romidepsin or nanaomycin A in combination with BGB 290 or cisplatin in a 8×8 matrix. Drug concentration ranges for each cell line was selected to cover 80 % to 0% metabolic inhibition. Cells were incubated at 37 °C for 96 hours. The metabolic activity was measured after 96 hours with the ATPlite assay (Perkin Elmer, 6016947). Values from the blank measurements were subtracted from the treatment wells and normalized to the vehicle controls (as described above). The effect of single treatments was compared to the combination treatments to determine additivity or synergy using the Loewe additivity model. Data from the secondary screen were analyzed using the Combenefit freeware software.

### Cell cycle analysis

Cells were seeded in tissue culture dishes (Corning, 353004) and incubated at 37 °C for 24 hours. Cells were then treated with vehicle control, BGB 290, romidepsin or BGB 290 with romidepsin at the indicated concentrations. Cells were incubated for 48 hours at 37 °C. Supernatant was collected in 50 ml falcon tubes. Cells were trypsinized using 0.25 % trypsin and collected. Cells were centrifuged at 1000 rpm for 5 minutes at room temperature. Supernatant was removed carefully. Pellets were washed once with sterile 1X PBS. Pellets were resuspended in chilled 70% ethanol and incubated at 4 °C for 2 hours. Cells were centrifuged at 420 g for 10 minutes at room temperature. Supernatant was removed carefully. Pellets were resuspended in cell cycle staining solution made of 2.5% propidium iodide (Santa Cruz, sc-3541A), 2% DNAse-free RNAse (Sigma, 10109169001) and 0.1% Triton X-100 in PBS. Cell cycle staining solution permeabilizes the cells and stains the DNA. Cells were incubated at 4 °C for 30 minutes. The DNA content was measured using a FACS Fortessa and the data were quantified using the FACSDiva software.

### Apoptosis

Cells were seeded in tissue culture dishes (Corning, 353004) and were incubated at 37 °C for 24 hours. Cells were then treated with vehicle control, BGB 290, romidepsin or BGB 290 with romidepsin. Cells were incubated for 48 hours at 37 °C. Supernatant was collected in 50 ml falcon tubes. Cells were trypsinized using 0.25% trypsin and collected. Cells were centrifuged at 1000 rpm for 5 minutes at room temperature. Supernatant was removed carefully. Pellets were washed once with sterile 1X PBS. Pellets were dissolved in 1X annexin buffer (BD Biosciences, 556454). Suspended cells were distributed in FACS tubes. Cells were then incubated with FITC annexin V (BD Biosciences, 556419), 7AAD (BD Biosciences, 559925), no stain control or FITC annexin V with 7AAD. Cells were incubated at 4 °C for 15 minutes in the dark. Live, early apoptotic and late apoptotic or necrotic cells were quantified using a FACS Fortessa and data were analyzed using the FACSDiva software.

### Doubling time

Cells were seeded in tissue culture dishes (Corning, 353004) at low confluency. Cells were incubated at 37 °C. After 24 hours, cells were trypsinized using 0.25% trypsin. Cells were then treated with 50% trypan blue (Sigma, T8154). Cells were pipetted into counting slides (Bio Rad, 145-0011) and living cells were quantified using an automated cell counter (Bio Rad-TC20). To estimate the average doubling time, exponential linear regression was applied on the doubling time data.

### *In vivo* studies and chemotherapy

Freshly isolated cells from orthotopic patient-derived xenograft mouse model (LFS_MB_1R, first relapse) were injected into the right flanks of six to 10-week-old female immune-compromised mice (NSG, NOD.*Cg-Prkdc^scid^Il2rg^tm1Wjl^*) with 50% matrigel (Corning, 356234). Twice a week, flank tumor volumes were measured using digital caliper and calculated. Once the tumor volumes reached 100 mm^3^, the mice were randomized and subjected to either treatment group, vehicle control, romidepsin single treatment or BGB290 with romidepsin combination treatment. In the romidepsin single treatment group, mice were treated with 2 mg/kg of romidepsin administered i.p. (in saline) twice a week. In the BGB290 with romidepsin combination treatment group, mice were treated with 2 mg/kg of romidepsin administered i.p. (in saline) twice a week and 6 mg/kg of BGB290 administered p.o. (in 0.5% methylcellulose) twice a day. All animal experiments were performed in accordance with ethical and legal regulations for animal welfare and approved by the governmental council (Regierungspräsidium Karlsruhe, Germany).

### Acetyl-α-tubulin and phospho-histone 3 immunostaining

Daoy cells were seeded onto coverslips. After 24 hours the media in each well was replaced with fresh media containing the drug corresponding to the drug treatment group, DMSO control, 6.25 μM BGB 290, 1.7 nM romidepsin and 6.25 μM BGB 290 + 1.7 nM romidepsin. Cells were then incubated for 48 hours. The coverslips were then fixed for 20 minutes with 4% PFA. Next, a blocking buffer (1x PBS, 5% normal goat serum, 0.3% Triton X-100) was prepared and added to the coverslips for 1 hour at room temperature. The blocking buffer was then aspirated and anti-Acetyl-α-Tubulin and anti-Phospho-Histone 3 primary antibodies were diluted to 1:400 and 1:200 respectively in antibody dilution buffer (1x PBS, 1% BSA, 0.3% Triton X-100) and both were added simultaneously to each coverslip. Coverslips were incubated with the primary antibody overnight at 4 °C. Then coverslips were washed thrice for 5 minutes in 1X PBS. Subsequently goat anti-mouse and anti-rabbit secondary antibodies were diluted to 1:500 in antibody dilution buffer and both were added simultaneously to each coverslip. Coverslips were incubated for 2 hours in the dark at room temperature with the secondary antibody and were then washed thrice for 5 minutes in 1X PBS. Then they were rinsed in double distilled H2O followed by 100% ethanol and left to air dry. Coverslips were then mounted onto microscope slides using DAPI fluoromount and left to cure for 1 hour in the dark before imaging. Imaging was performed on the Axio Zeiss Imager.M2 microscope.

Determination of the percentage of mitotic cells per treatment group was done using using a macro designed at the DKFZ Light Microscopy Facility.

### Gene expression analysis

Cells from 3 PDX spheroid models (LFS_MB_P, primary tumor, LFS_MB_1R, first relapse and RCMB18) were seeded in suspension culture flask (Greiner Bio-One, 690195) and treated with either vehicle control, BGB 290 (IC20), romidepsin (IC20) or BGB 290 (IC20) with romidepsin (IC20) for 24 hours. Pictures were taken using a bright-field microscope to determine the effect of the treatment on the spheroid growth. RNA was then isolated using the RNeasy mini kit (Qiagen, 74104). Expression data were generated using Affymetrix Clariom S arrays.

### Analysis of expression data and GSEA

Expression analysis and normalization of Affymetrix Clariom S arrays was performed by the Bioconductor package ‘oligo’^28^. Sample to sample correlation was illustrated by a heatmap, using the Pearson correlation of the top 1000 variable genes. Combination treated cell lines were compared with both the romidepsin treated cell lines, BGB 290 treated cell lines, and the normal human astrocytes. Likewise, romidepsin treated cell lines and BGB 290 treated cell lines were compared with the normal human astrocytes. Differential expression analysis was performed by limma^29^. The False Discovery Rate (FDR) adjustment of p-values were performed by the Benjamini-Hochberg method. The Gene Set Enrichment Analysis (GSEA) was performed using gene-sets from MSigDB5. Enrichment test on Oncogenic signature gene sets (C6) from MSigDB was performed by the signed t-statistic from the differential expression analysis. Gene-sets with False Discovery Rate (FDR) smaller than 5% were considered as significant.

### Statistical analysis

Statistical analyses and visualizations were performed using R, shiny app, Combenefit software^30^ and GraphPad PRISM 5.

## Supplementary Figures

**Supplementary Figure 1:**

**Molecular characterization of *in vitro* models included in the screen:** Copy number variant (CNV) plots of cell lines included in the screen generated using 450 K methylation array, EPIC methylation arrays or whole-genome sequencing.

**Supplementary Figure 2: Controls for the primary screen:** (a) 96-well plate showing raw ATPlite readout values for the primary screen with negative controls on the left and positive controls on the right of the plate. (b) Scatter plot of the inhibition of the metabolic activity values (percentage of the DMSO controls) for CUDC-907 (on the left) and entinostat (on the right), which were present in both drug libraries utilized for the screen. The plot was generated using the percentage of the metabolic inhibition values of the drug in TargetMol on the X-axis versus ApexBio on the Y-axis. (c) Scatter plot of the inhibition of the metabolic activity values (percentage of the DMSO controls) for UWB1.289 (on the left) and UWB1.289 + BRCA1 (on the right). The plot was generated using the percentage of the metabolic inhibition values at the start of the primary screen on the X-axis versus at the end of the primary screen on the Y-axis.

**Supplementary Figure 3:**

**Analysis of the primary screen data:** (a) Cluster dendrogram of all the cell lines and tumor entities included in the primary screen. The dendrogram was constructed based on the primary screen data. (b) Scatter plot of the inhibition of the metabolic activity values (percentage of the DMSO controls) from the primary screen. The plot was generated using the percentage of the metabolic inhibition values of the single agents on the X-axis versus the same agent with cisplatin (IC20) double treatment (left plot) or the same agent with BGB 290 (IC20) double treatment (right plot) on the Y-axis. (c) Bright-field images of spheroids (Sonic Hedgehog medulloblastoma model, first relapse) after 24 hours of drug treatment. Scale bar in each panel: 100 μm.

**Supplementary Figure 4:**

**Analysis of the secondary screen data:** Loewe surface plots generated using the combenefit software. Synergy and antagonism are mapped to dose response using the average of the percentage of the metabolic activity values of 3 replicates per data point. Technical replicates for patient derived xenograft spheroid models and HD-MB03 cell line. Biological replicates for all other cell lines.

**Supplementary Figure 5:**

**Apoptosis in RPE1 WT and RPE1 *TP53* KO cells upon combination treatment with romidepsin and BGB 290:** Apoptotic fractions in RPE1 WT and RPE1 *TP53* KO cells were analyzed by FITC-annexin V and 7AAD stains. Cells were treated with drugs at the indicated concentrations for 48 hours and subjected to flow cytometry analysis. Dot plots are representative of three biological replicates. Bar graphs (at the bottom) showing the quantitative distribution of RPE1 WT and RPE1 *TP53* KO cells as living, early apoptotic and late apoptotic/ necrotic. Statistical significance was tested by two-way ANOVA followed by Bonferroni post-test, *** p<0.001, ** p<0.01, * p<0.05, ns p>0.05. Error bars represent SEM, n=3 biological replicates (L = Live, EA = Early apoptosis and LA = Late apoptosis, N = necrosis, ns = non-significant).

**Supplementary Figure 6:**

**Sample to sample transcriptome correlation heatmap:**

The heatmap represents the Pearson correlation between the transcriptomes of spheroids isolated from patient-derived xenograft models subjected to different treatments. Pearson correlations were computed by cell-line corrected expression values of the top 1000 variable genes.

## Acknowledgements

We thank Thomas Höfer for discussions, Eugene Gbekor, Britta Jehle, Norman Mack, Sabrina Kirschner, Herman Stammer, and Manuel Fischer for technical support. We thank Kerstin Dell and her team for advice on animal experiments. The DKFZ Genomics Core facility is acknowledged for support with microarray experiments. The DKFZ Light Microscopy facility, and especially Damir Krunic, is acknowledged for help with macro design. Axel Benner is acknowledged for advice on statistical analyses. We thank the Wilhelm Sander Foundation and the Fritz Thyssen Foundation for funding.

## REFERENCES

1 Rausch, T. et al. Genome sequencing of pediatric medulloblastoma links catastrophic DNA rearrangements with TP53 mutations. Cell 148, 59–71, doi:10.1016/j.cell.2011.12.013 (2012).

2 Stephens, P. J. et al. Massive genomic rearrangement acquired in a single catastrophic event during cancer development. Cell 144, 27–40, doi:10.1016/j.cell.2010.11.055 (2011).

3 Voronina, N. et al. The landscape of chromothripsis across adult cancer types. Nature communications 11, 2320, doi:10.1038/s41467-020-16134-7 (2020).

4 Cortes-Ciriano, I. et al. Comprehensive analysis of chromothripsis in 2,658 human cancers using whole-genome sequencing. Nature genetics, doi:10.1038/s41588-019-0576-7 (2020).

5 Waszak, S. M. et al. Spectrum and prevalence of genetic predisposition in medulloblastoma: a retrospective genetic study and prospective validation in a clinical trial cohort. The Lancet. Oncology 19, 785–798, doi:10.1016/S1470-2045(18)30242-0 (2018).

6 Molenaar, J. J. et al. Sequencing of neuroblastoma identifies chromothripsis and defects in neuritogenesis genes. Nature 483, 589–593, doi:10.1038/nature10910 (2012).

7 Fontana, M. C. et al. Chromothripsis in acute myeloid leukemia: biological features and impact on survival. Leukemia 32, 1609–1620, doi:10.1038/s41375-018-0035-y (2018).

8 Ratnaparkhe, M. et al. Defective DNA damage repair leads to frequent catastrophic genomic events in murine and human tumors. Nature communications 9, 4760, doi:10.1038/s41467-018-06925-4 (2018).

9 Grobner, S. N. et al. The landscape of genomic alterations across childhood cancers. Nature, doi:10.1038/nature25480 (2018).

10 Evers, B., Helleday, T. & Jonkers, J. Targeting homologous recombination repair defects in cancer. Trends Pharmacol Sci 31, 372–380, doi:10.1016/j.tips.2010.06.001 (2010).

11 Bryant, H. E. et al. Specific killing of BRCA2-deficient tumours with inhibitors of poly(ADP-ribose) polymerase. Nature 434, 913–917, doi:10.1038/nature03443 (2005).

12 Kaufman, B. et al. Olaparib monotherapy in patients with advanced cancer and a germline BRCA1/2 mutation. Journal of clinical oncology: official journal of the American Society of Clinical Oncology 33, 244–250, doi:10.1200/JCO.2014.56.2728 (2015).

13 Bundred, N. et al. Evaluation of the pharmacodynamics and pharmacokinetics of the PARP inhibitor olaparib: a phase I multicentre trial in patients scheduled for elective breast cancer surgery. Invest New Drugs 31, 949–958, doi:10.1007/s10637-012-9922-7 (2013).

14 Pilie, P. G., Gay, C. M., Byers, L. A., O’Connor, M. J. & Yap, T. A. PARP Inhibitors: Extending Benefit Beyond BRCA-Mutant Cancers. Clinical cancer research: an official journal of the American Association for Cancer Research 25, 3759–3771, doi:10.1158/1078-0432.CCR-18-0968 (2019).

15 Heske, C. M. et al. Matrix Screen Identifies Synergistic Combination of PARP Inhibitors and Nicotinamide Phosphoribosyltransferase (NAMPT) Inhibitors in Ewing Sarcoma. Clinical cancer research: an official journal of the American Association for Cancer Research 23, 7301–7311, doi:10.1158/1078-0432.CCR-17-1121 (2017).

16 Henssen, A. G. et al. Therapeutic targeting of PGBD5-induced DNA repair dependency in pediatric solid tumors. Sci Transl Med 9, doi:10.1126/scitranslmed.aam9078 (2017).

17 Takagi, M. et al. Loss of DNA Damage Response in Neuroblastoma and Utility of a PARP Inhibitor. J Natl Cancer Inst 109, doi:10.1093/jnci/djx062 (2017).

18 Milde, T. et al. HD-MB03 is a novel Group 3 medulloblastoma model demonstrating sensitivity to histone deacetylase inhibitor treatment. J Neurooncol 110, 335–348, doi:10.1007/s11060-012-0978-1 (2012).

19 Pei, Y. et al. HDAC and PI3K Antagonists Cooperate to Inhibit Growth of MYC-Driven Medulloblastoma. Cancer cell 29, 311–323, doi:10.1016/j.ccell.2016.02.011 (2016).

20 Smolewski, P. & Robak, T. The discovery and development of romidepsin for the treatment of T-cell lymphoma. Expert Opin Drug Discov 12, 859–873, doi:10.1080/17460441.2017.1341487 (2017).

21 Ecker, J. et al. Reduced chromatin binding of MYC is a key effect of HDAC inhibition in MYC amplified medulloblastoma. Neuro Oncol 23, 226–239, doi:10.1093/neuonc/noaa191 (2021).

22 Team, R. C. R: A language and environment for statistical computing. (2013).

23 Chang, W., Cheng, J., Allaire, J., Xie, Y. & McPherson, J. Shiny: web application framework for R. R package version 1, 2017 (2017).

24 van der Maaten, L. & Hinton, G. Visualizing data using t-SNE. journal of Machine Learning Research 9. Nov (2008) (2008).

25 Gaujoux, R. & Seoighe, C. A flexible R package for nonnegative matrix factorization. BMC bioinformatics 11, 1–9 (2010).

26 Gu, Z., Eils, R. & Schlesner, M. Complex heatmaps reveal patterns and correlations in multidimensional genomic data. Bioinformatics 32, 2847–2849 (2016).

27 Sievert, C. Interactive web-based data visualization with R, plotly, and shiny. (CRC Press, 2020).

28 Carvalho, B. S. & Irizarry, R. A. A framework for oligonucleotide microarray preprocessing. Bioinformatics 26, 2363–2367, doi:10.1093/bioinformatics/btq431 (2010).

29 Ritchie, M. E. et al. limma powers differential expression analyses for RNA-sequencing and microarray studies. Nucleic acids research 43, e47, doi:10.1093/nar/gkv007 (2015).

30 Di Veroli, G. Y. et al. Combenefit: an interactive platform for the analysis and visualization of drug combinations. Bioinformatics 32, 2866–2868, doi:10.1093/bioinformatics/btw230 (2016).

