## Supplementary figures 1-6 for "A synergistic interaction between HDAC- and PARP inhibitors in childhood tumors with chromothripsis"

### Supplementary figure 1

KHOS-240S (Osteosarcoma cell line)

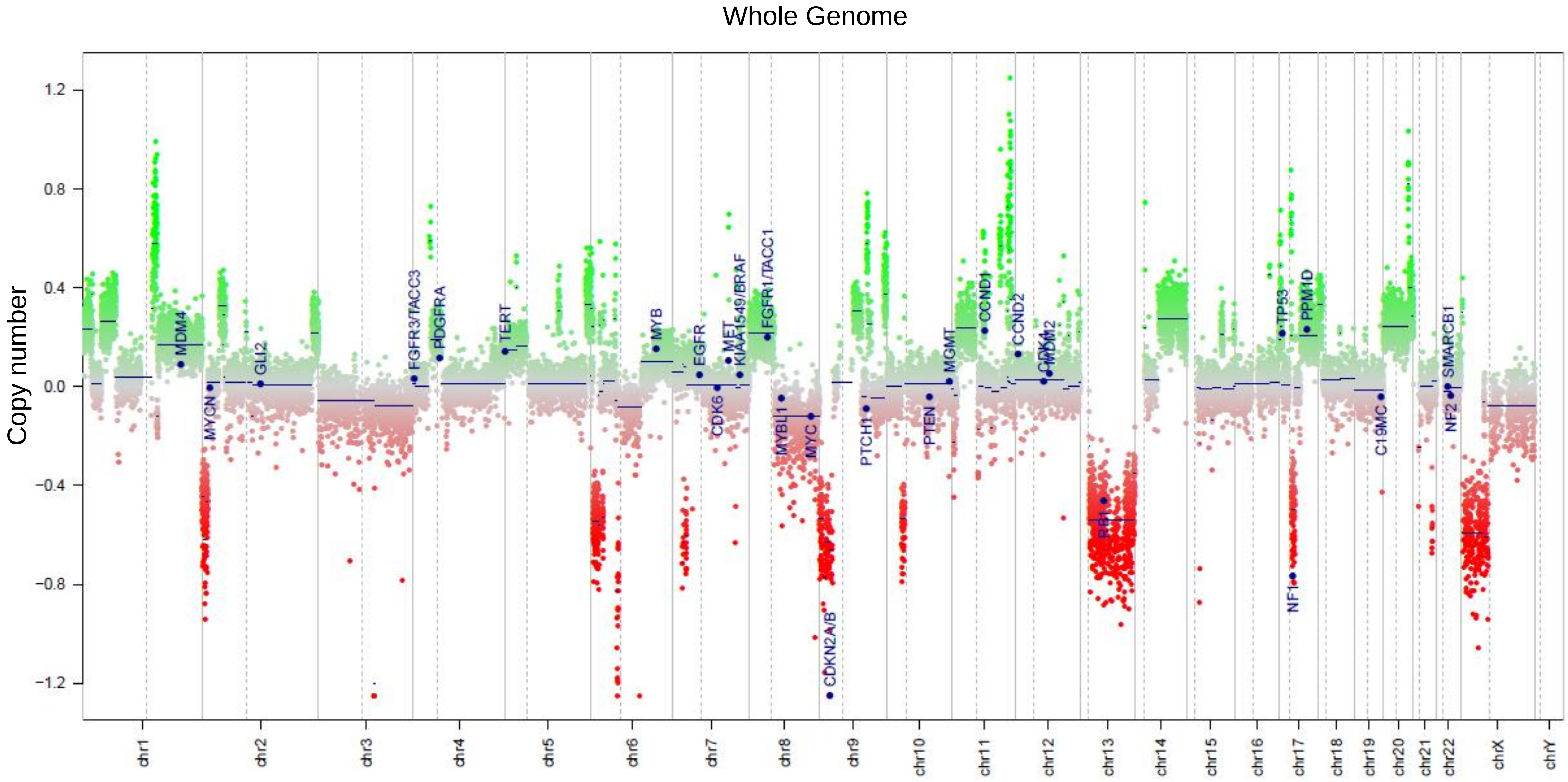


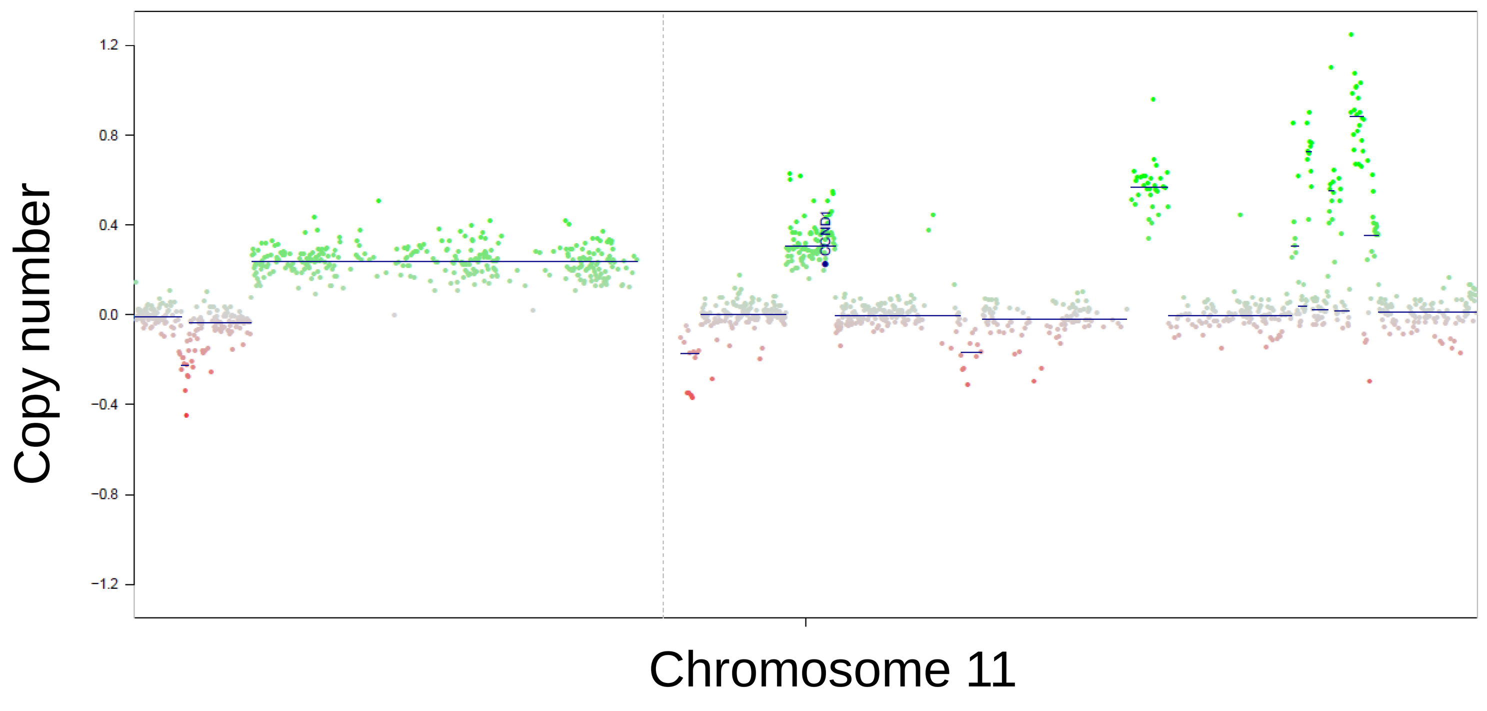


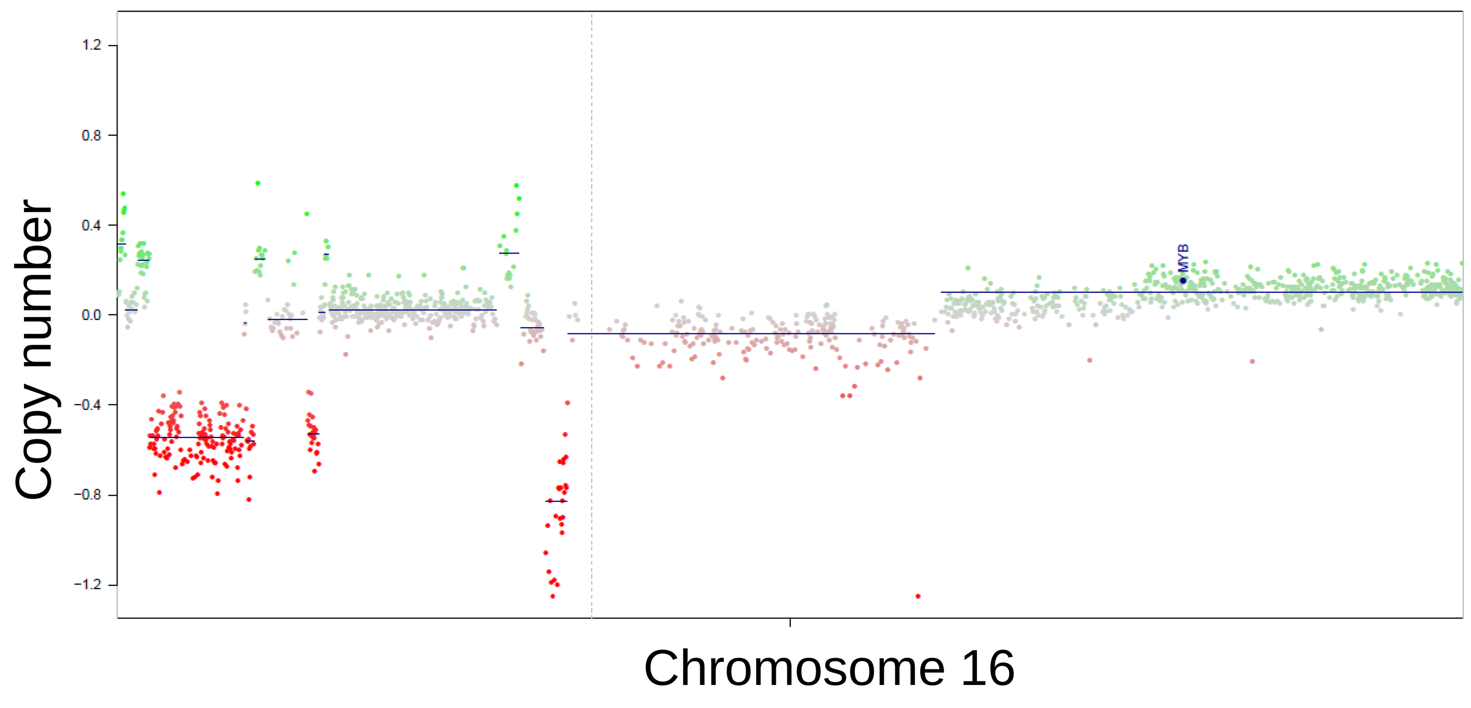


Saos-2 (Osteosarcoma cell line)

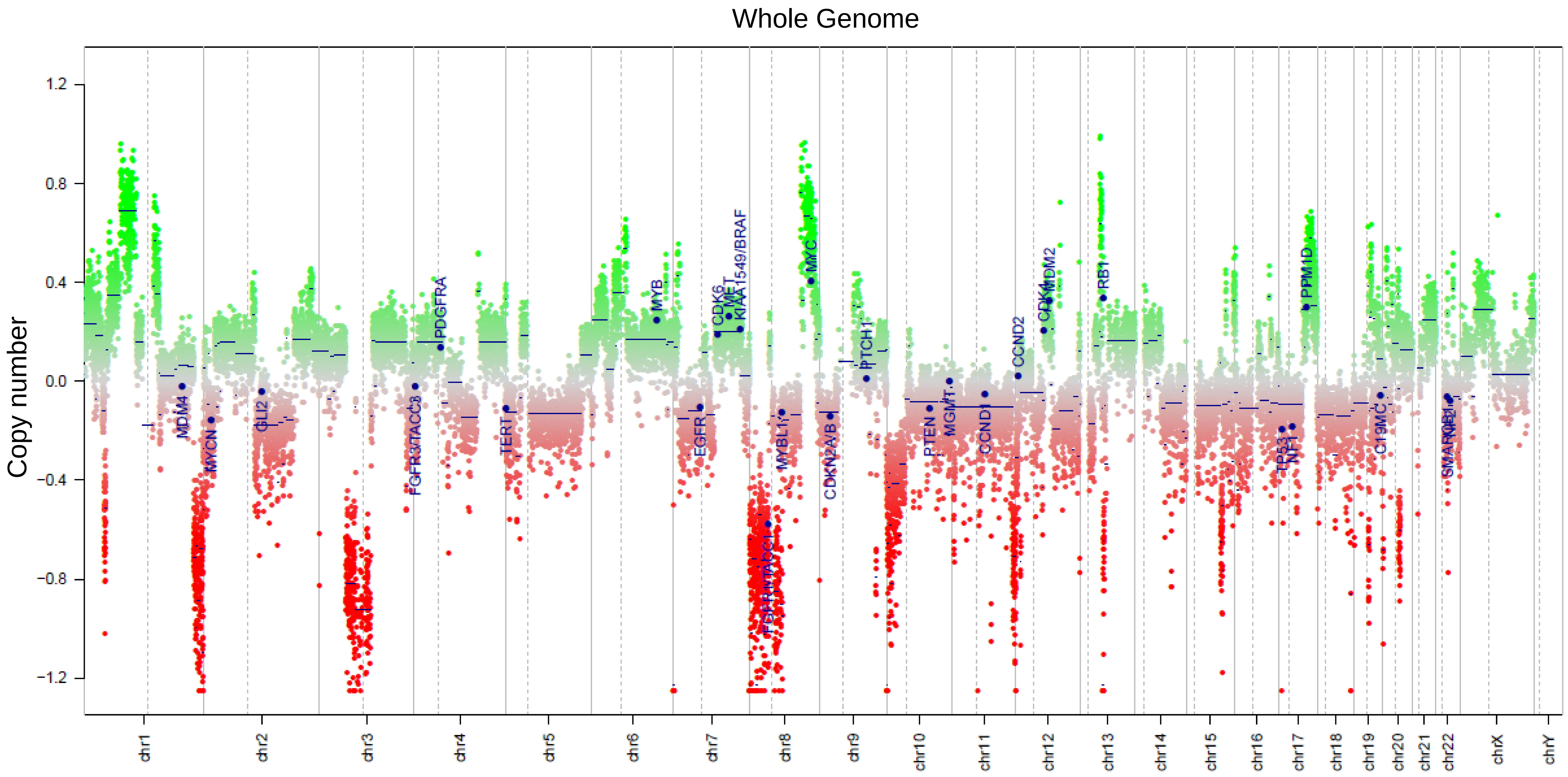


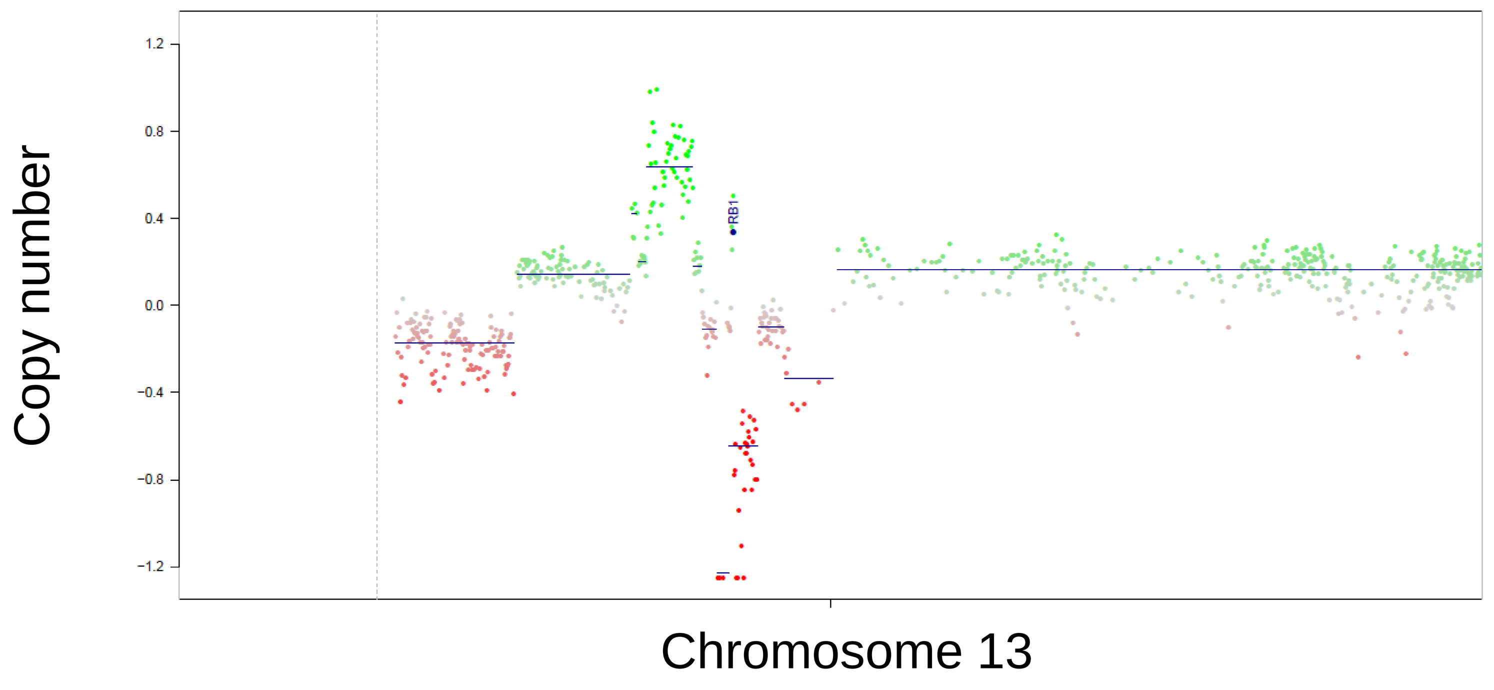


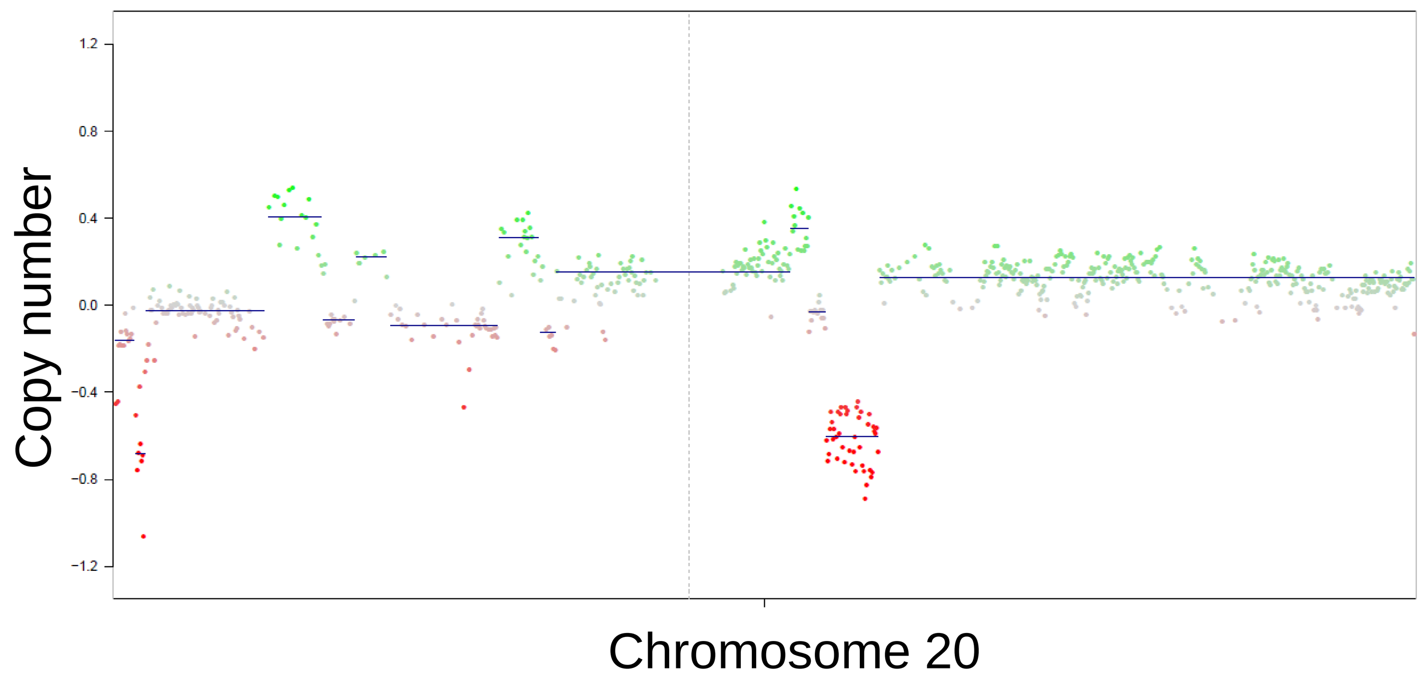

UWB1.289 (*BRCA1* mutant control line)

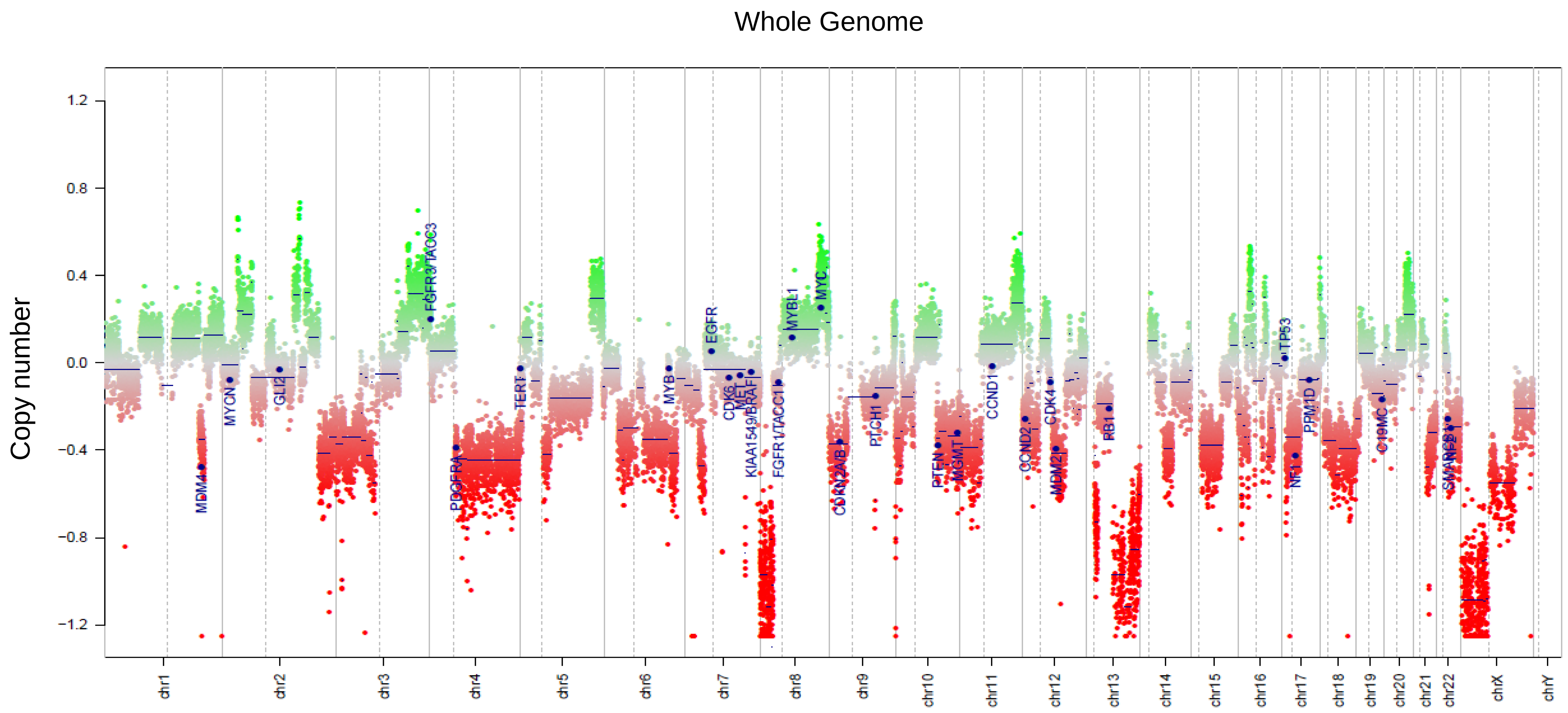


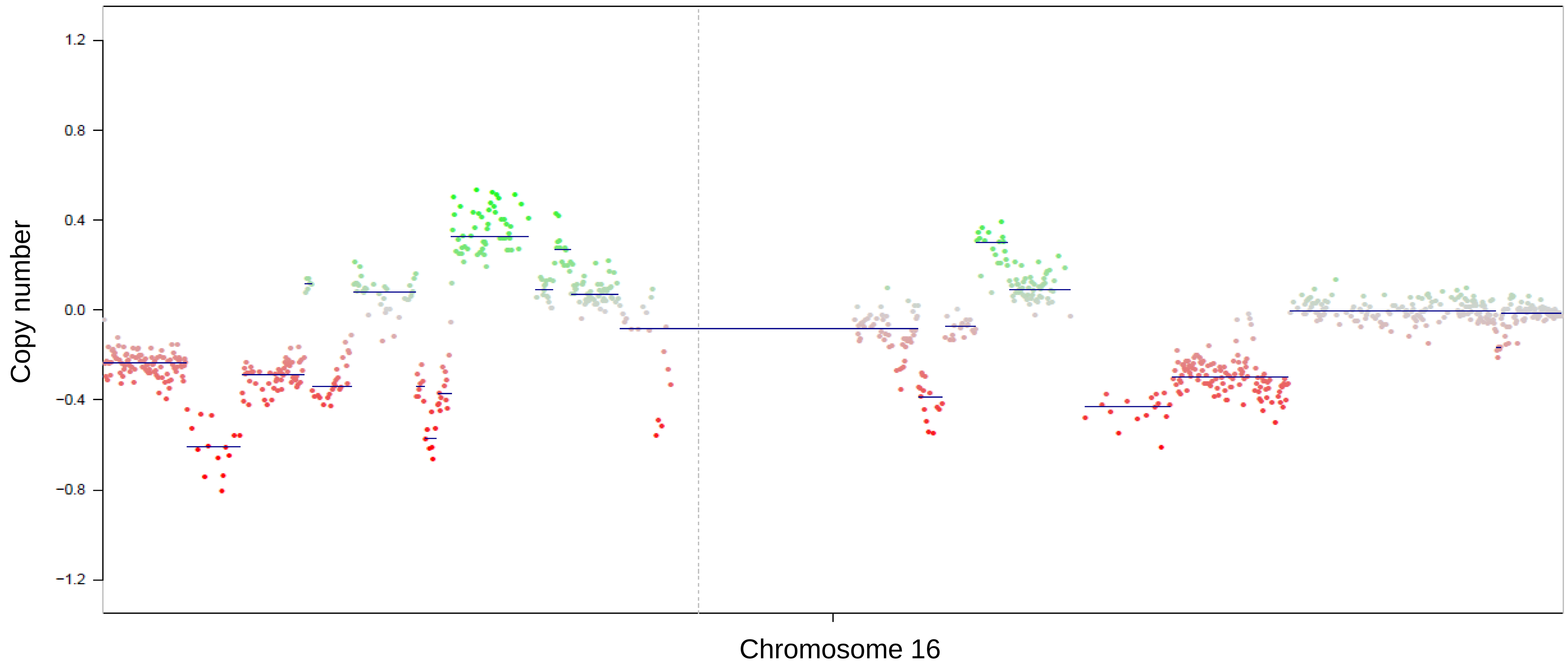


SJSA-1 (Osteosarcoma cell line)

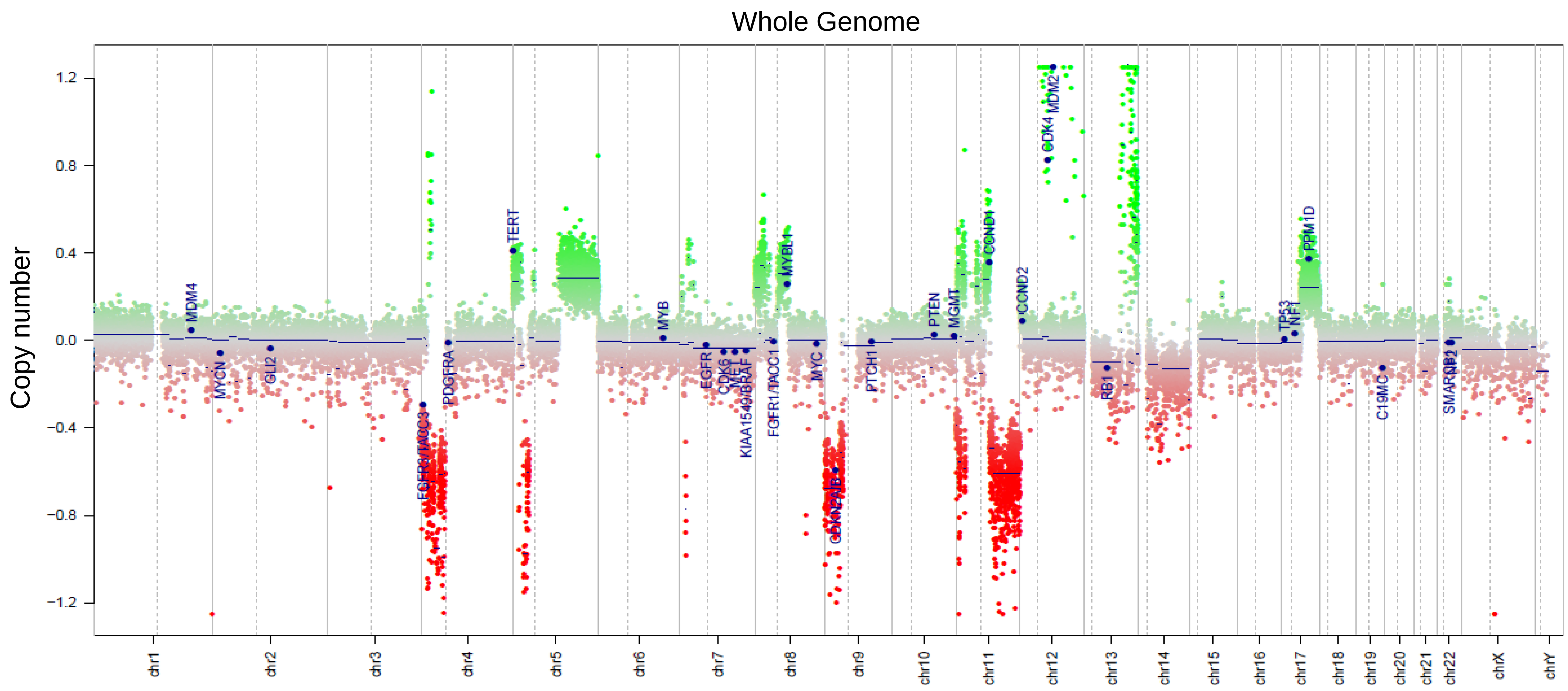


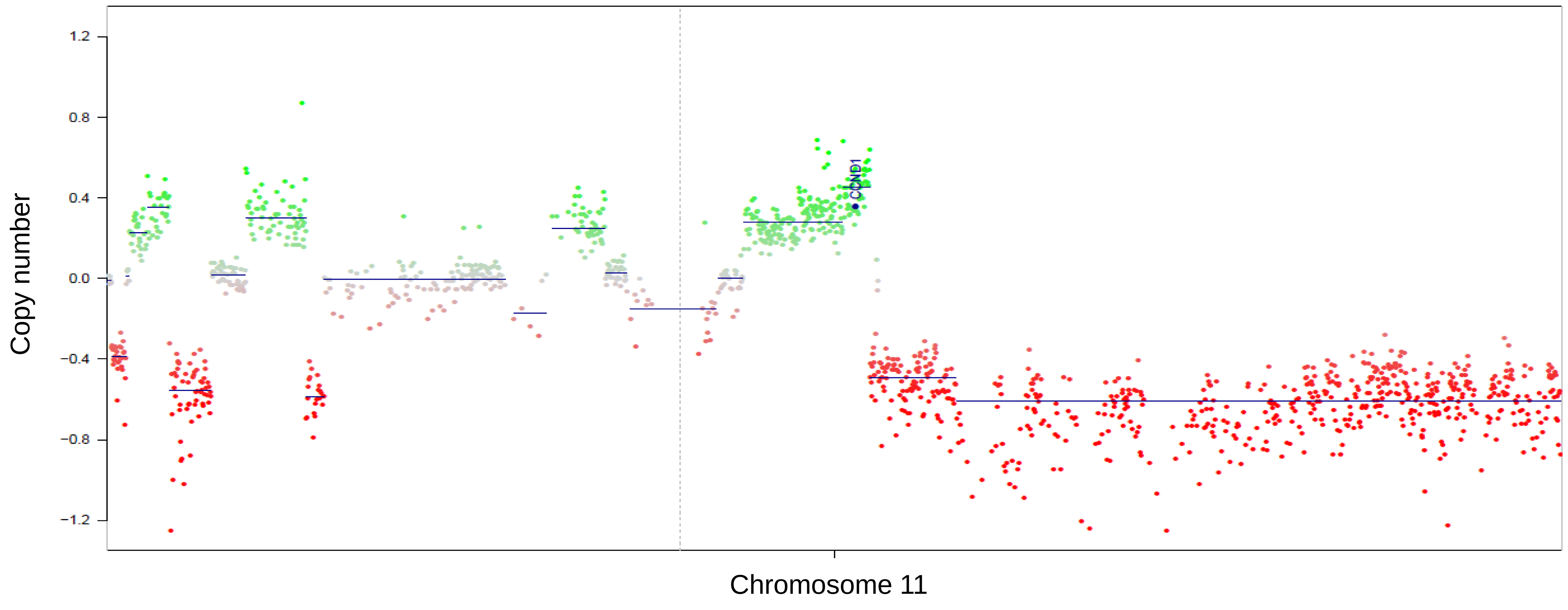

UWB1.289 (*BRCA1* rescue line)

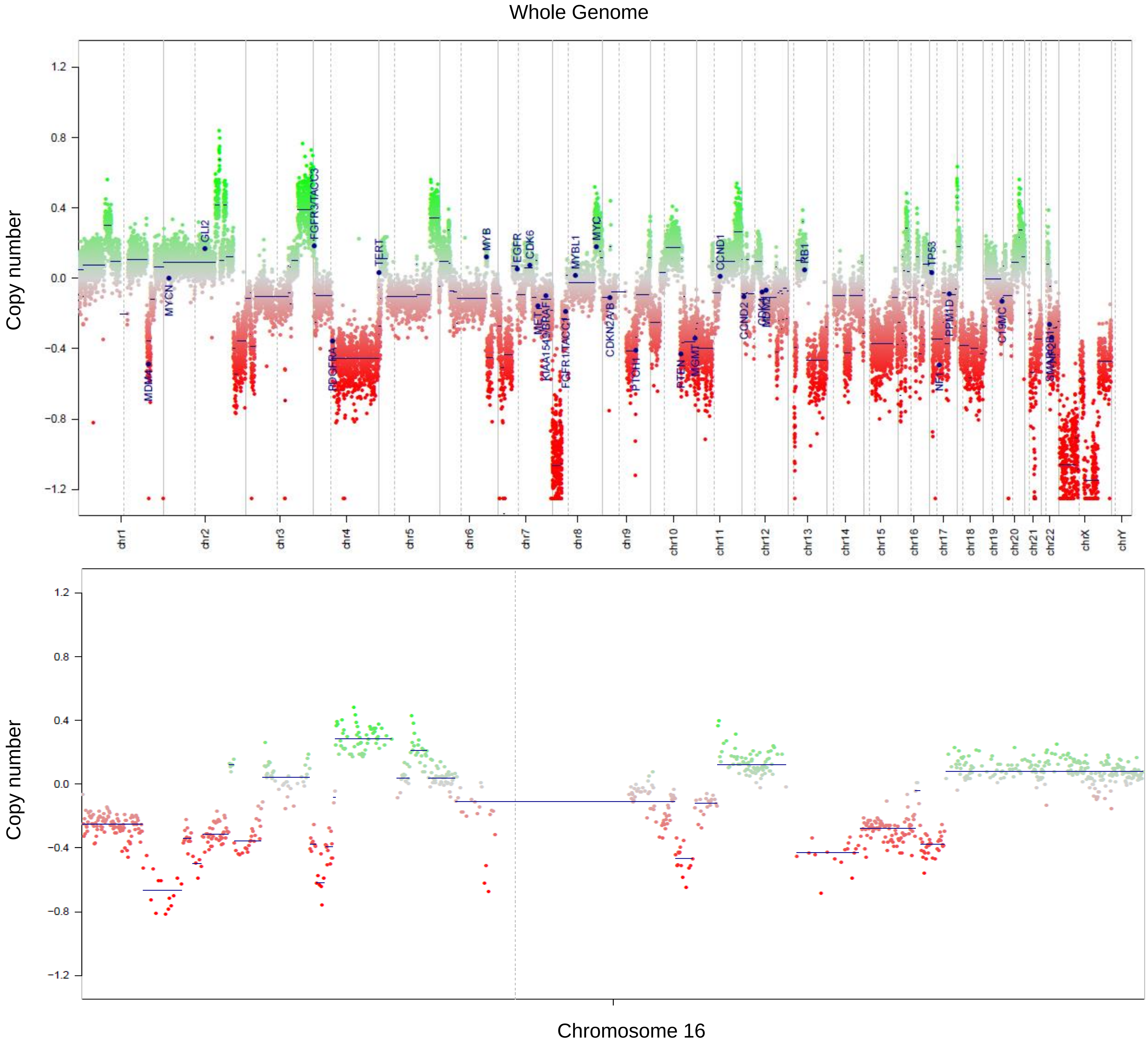


MDA-MB-436 (*BRCA 1* mutant line )

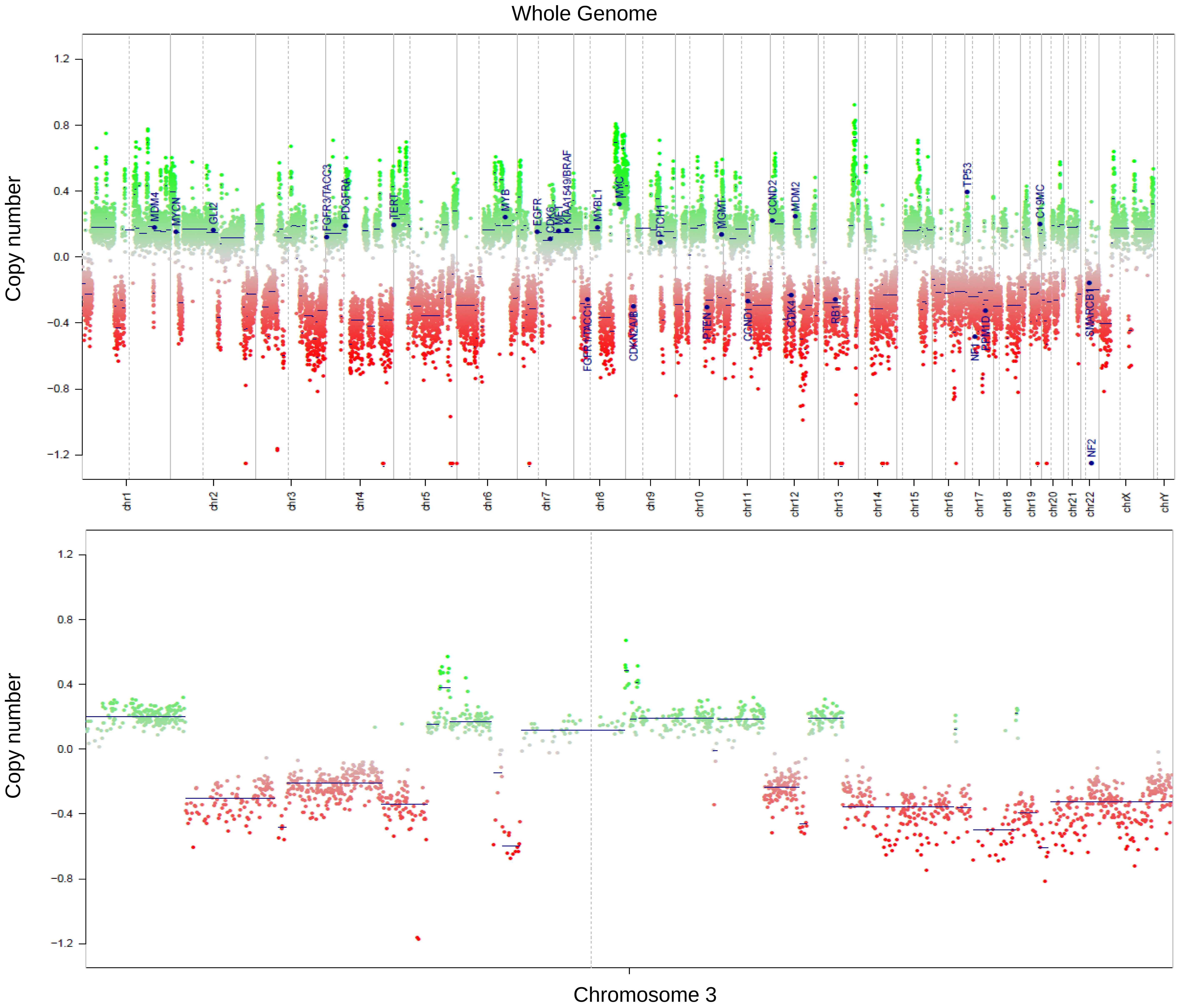

HD-N33 (Neuroblastoma cell line )

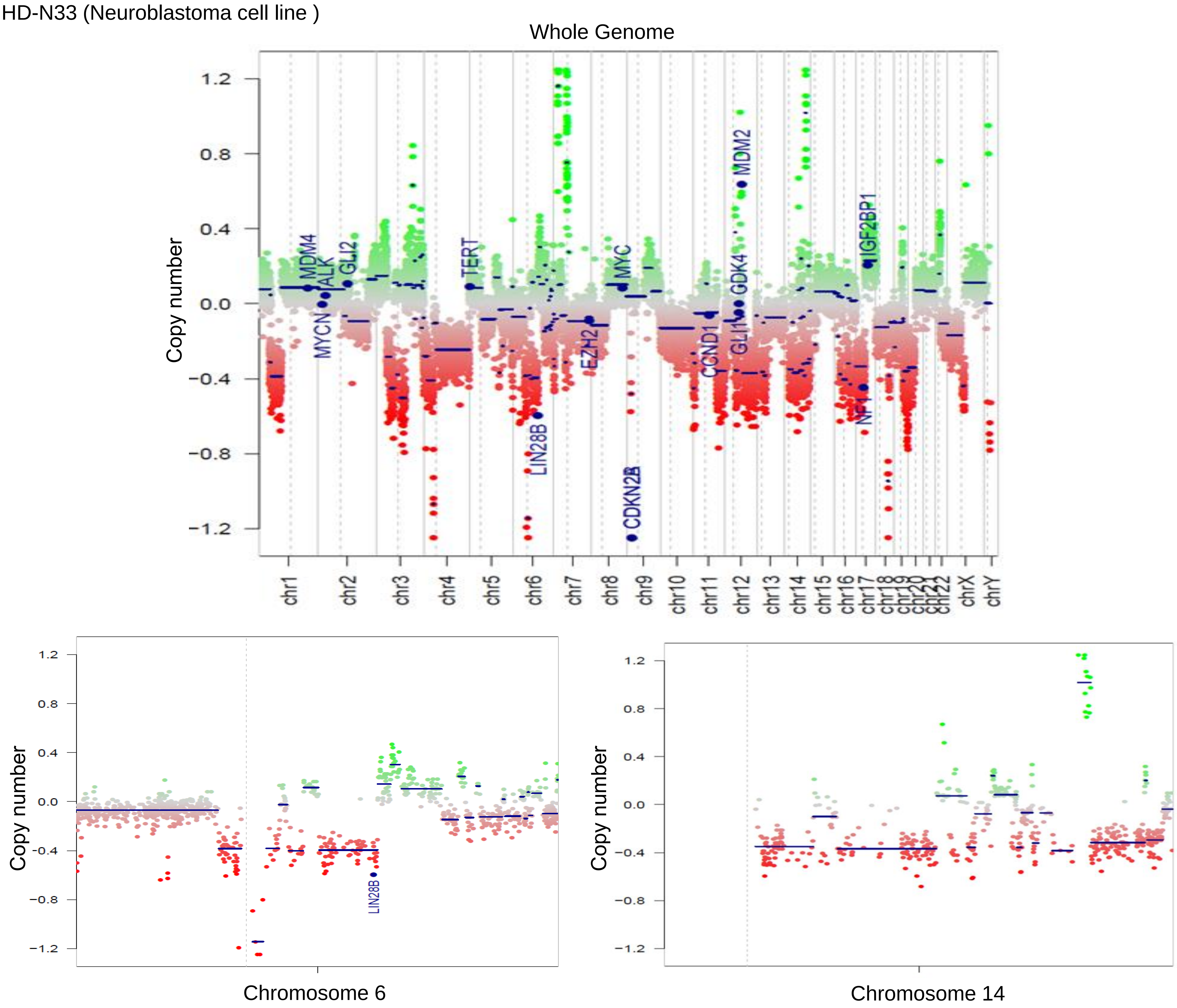


MB145 (medulloblastoma from which patient-derived xenograft model BT084 was established)

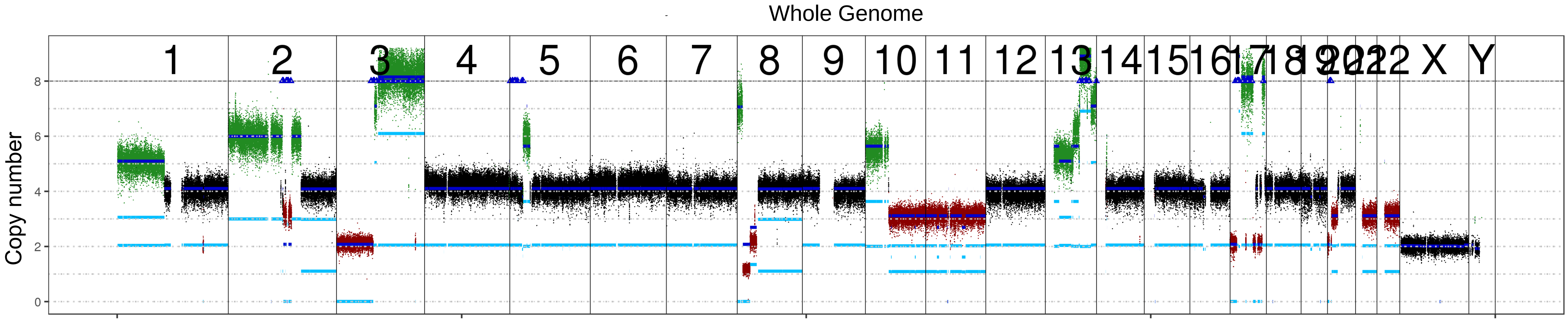


LFS_MB_P (medulloblastoma)

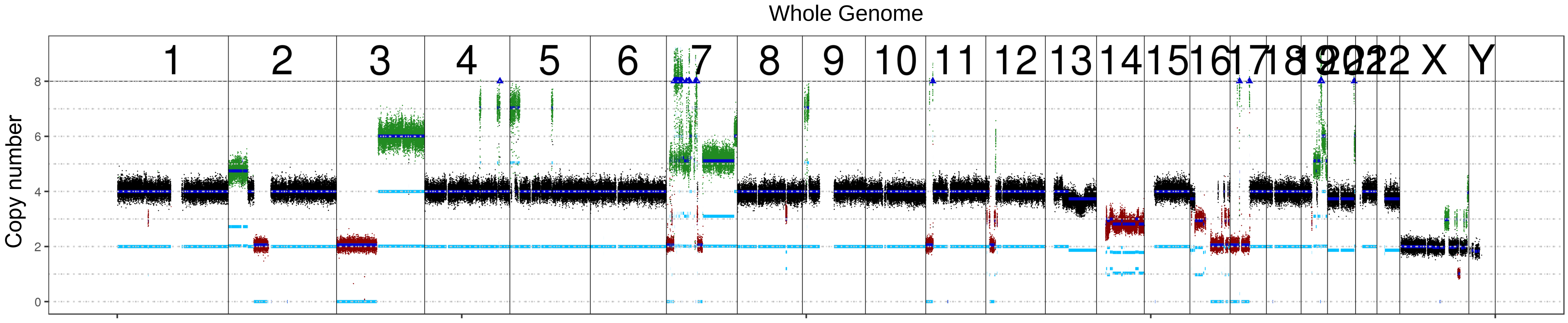

LFS_MB_1R (medulloblastoma)

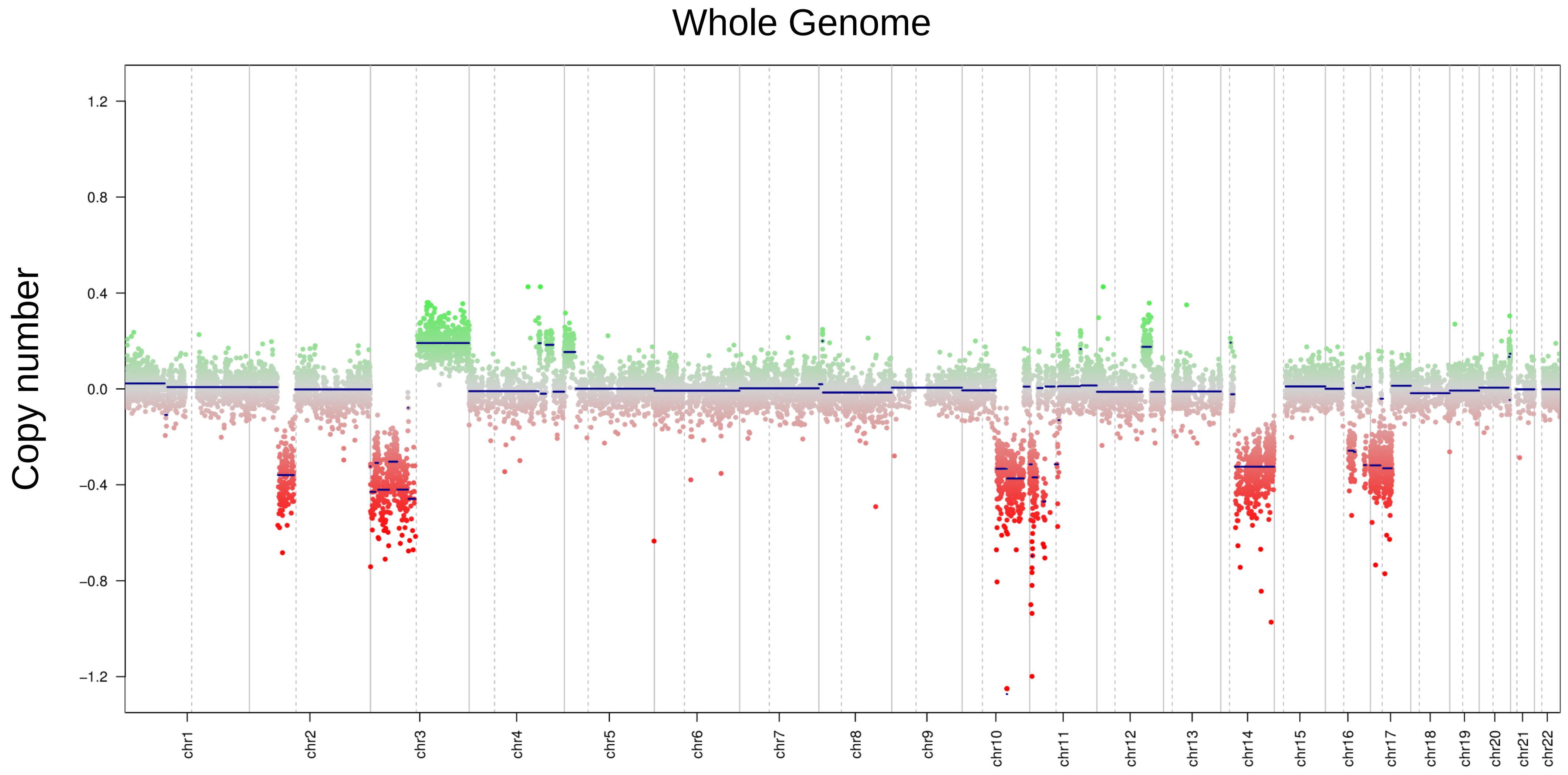


LFS_MB_2R (medulloblastoma)

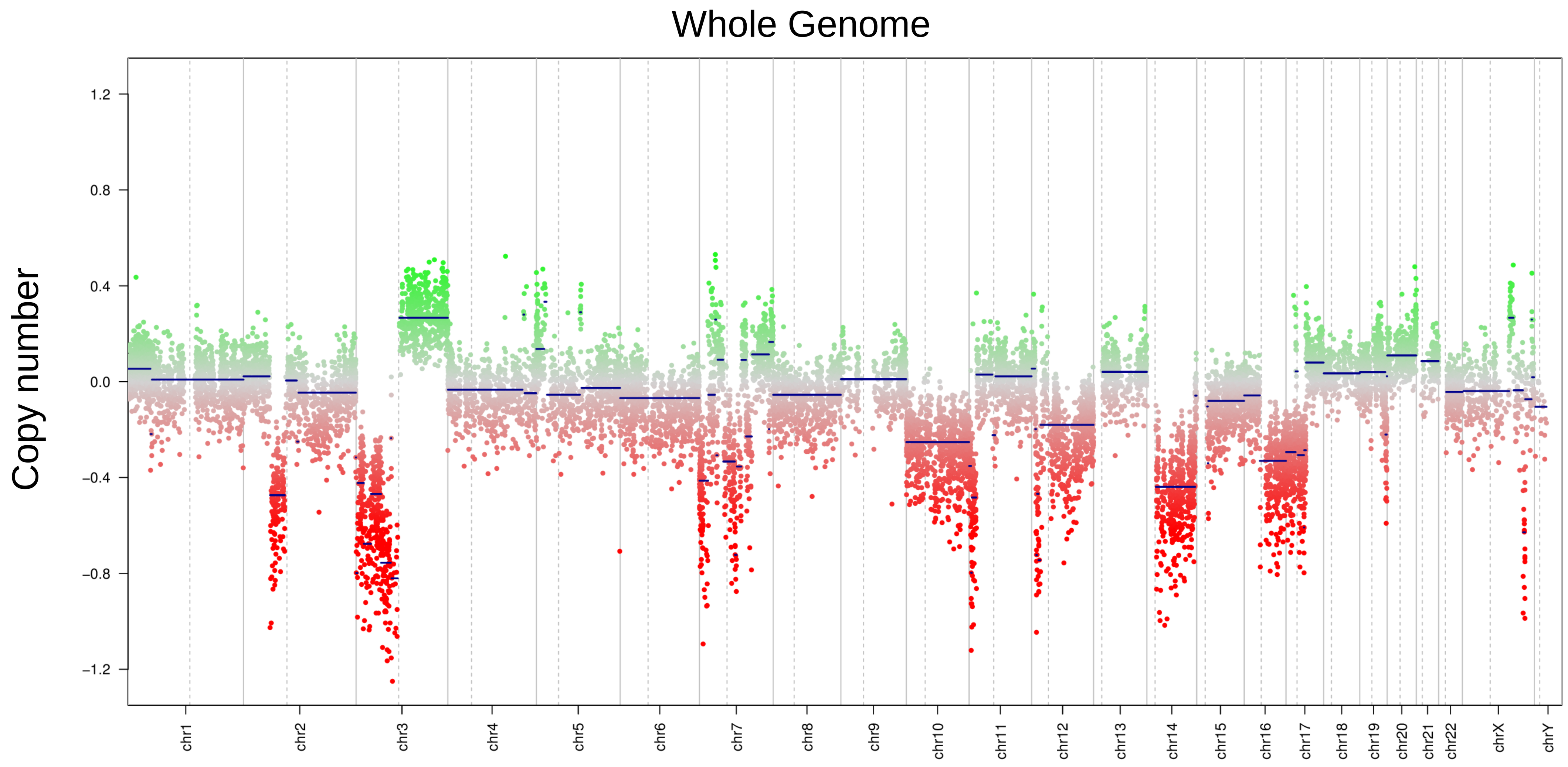

### Supplementary figure 2

## a.

UWB1.289

| | | | | | | | | | | | |
|---|---|---|---|---|---|---|---|---|---|---|---|
| 4450480 | 4475720 | 646640 | 4161560 | 4677920 | 4655440 | 1851080 | 4733520 | 4410200 | 3581240 | 4465960 | 65840 |
| 4855760 | 4830120 | 5083000 | 4424640 | 848160 | 4753440 | 4633120 | 4471520 | 5267840 | 4884320 | 4697200 | 98600 |
| 4200160 | 1569640 | 3725600 | 4710000 | 4490440 | 4377320 | 4160880 | 2016680 | 4248040 | 2187400 | 4319720 | 90480 |
| 5017840 | 4650320 | 4788240 | 4537600 | 1129720 | 5361000 | 4511760 | 4154120 | 4039880 | 4968480 | 4500440 | 99920 |
| 3962400 | 4246600 | 3966120 | 3789520 | 2744960 | 3941040 | 1983480 | 3870800 | 4185160 | 4062120 | 3931760 | 71200 |
| 4640360 | 3993480 | 3772640 | 3923640 | 2376480 | 3809800 | 4152640 | 851600 | 4290000 | 4068800 | 2237000 | 60680 |
| 3951680 | 4156640 | 4053400 | 4029400 | 4232200 | 4009720 | 3986120 | 3419240 | 3021240 | 472040 | 1802960 | 82560 |
| 4439440 | 4549760 | 4433400 | 4600280 | 4467200 | 4119120 | 4455840 | 4543160 | 4423160 | 4349720 | 4477840 | 83640 |

Negative control

Positive control

## b.

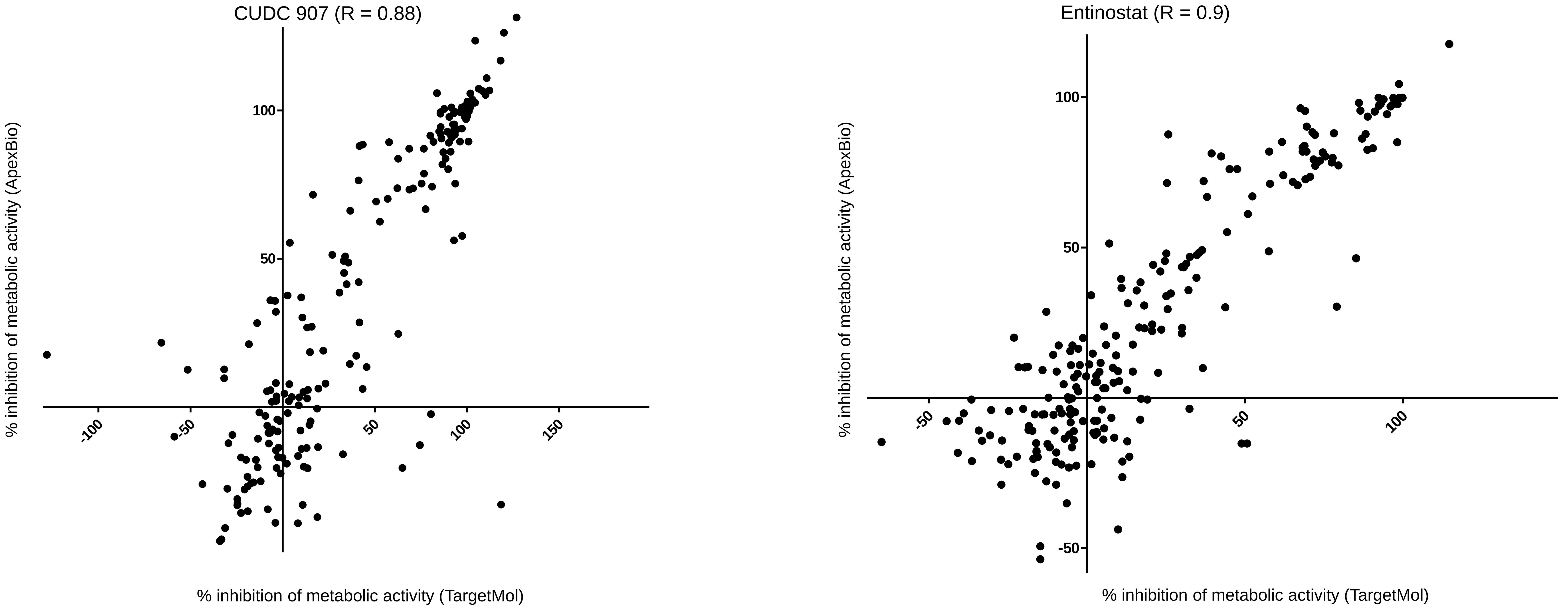


## c.

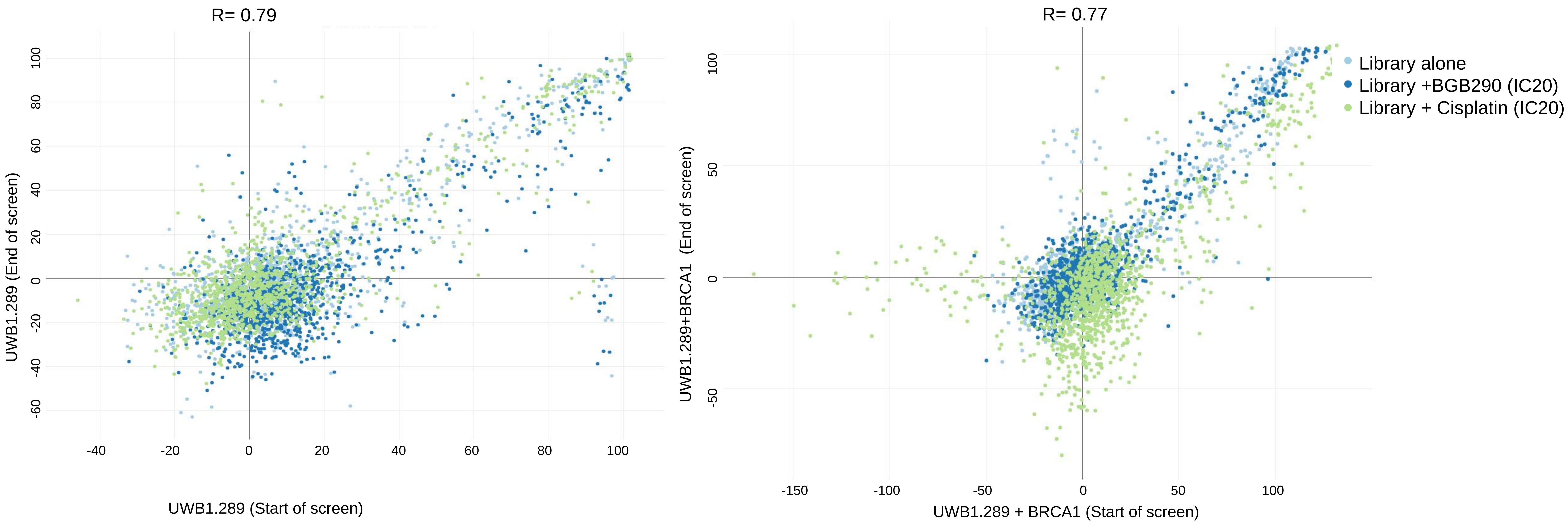

### Supplementary figure 3

**a.**

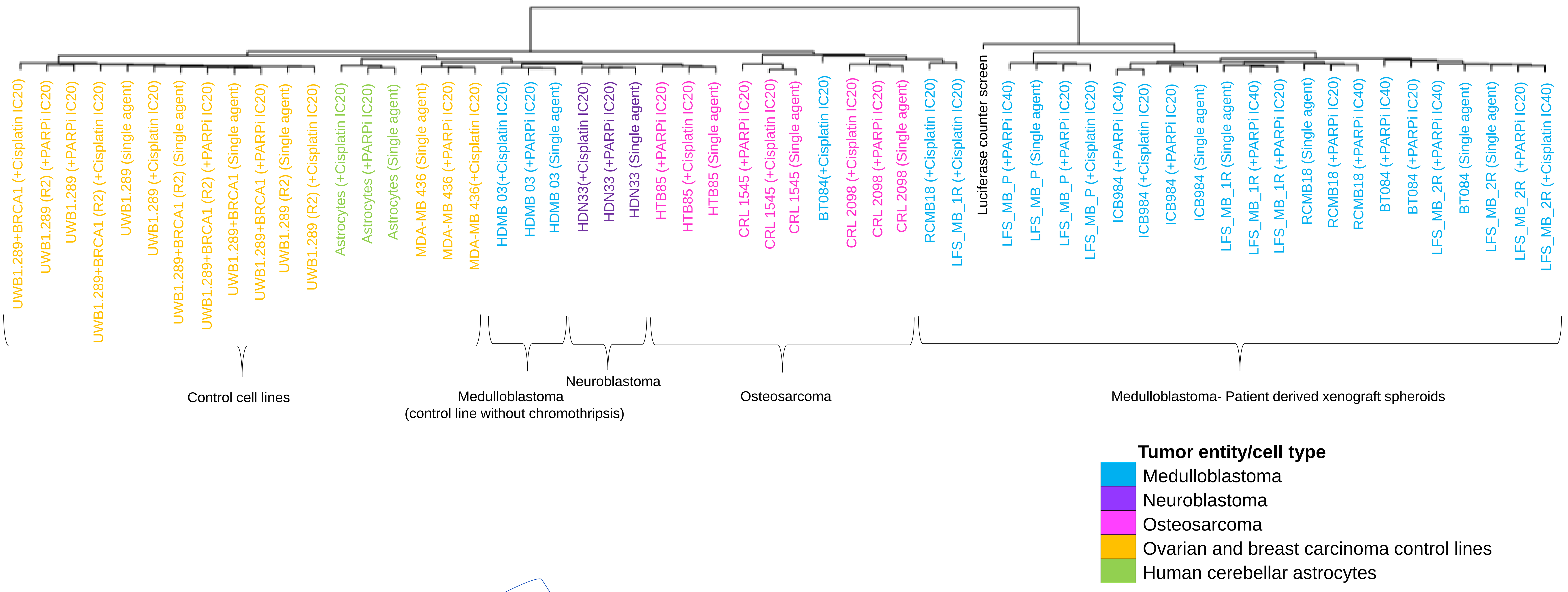


**b.**

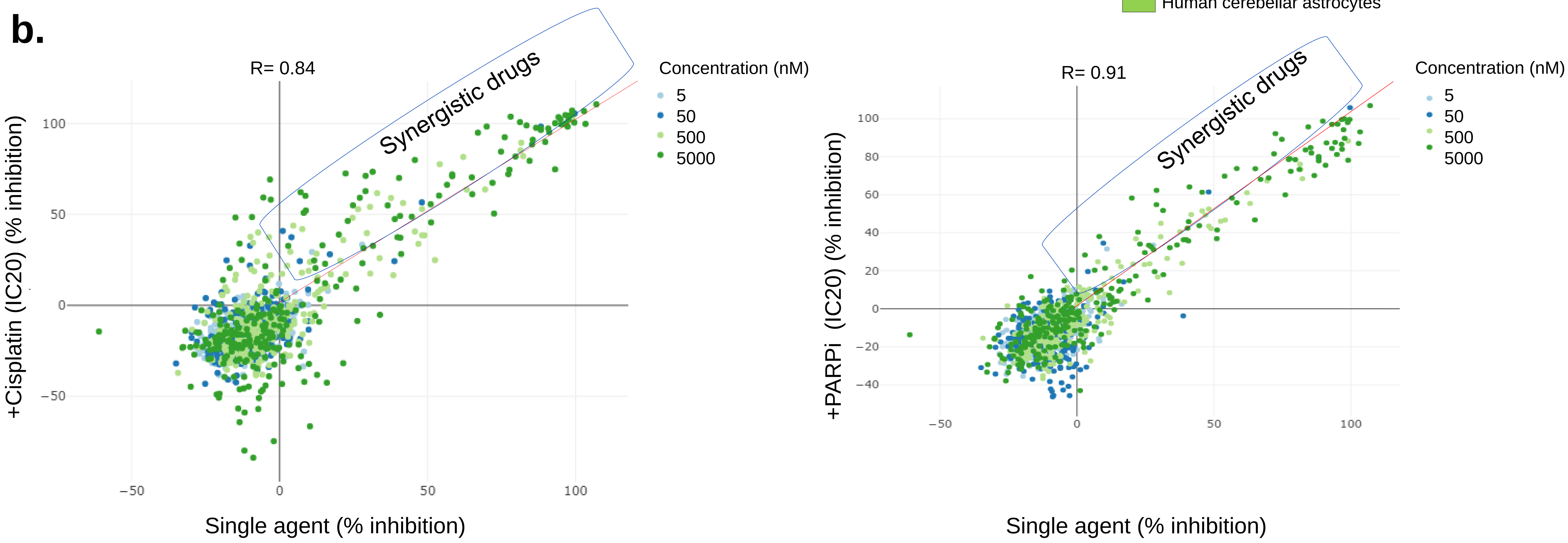


**c.**

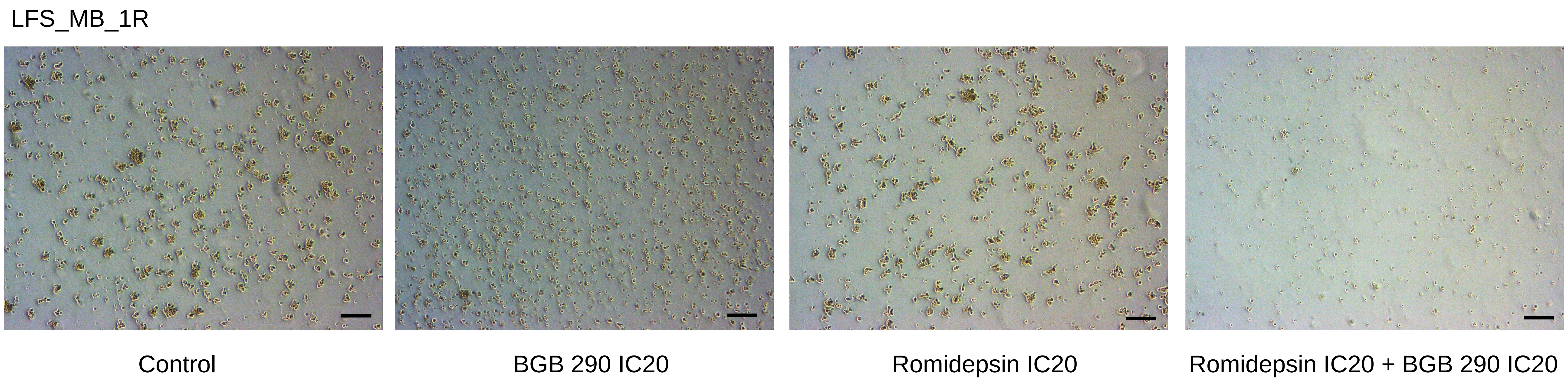

### Supplementary figure 4

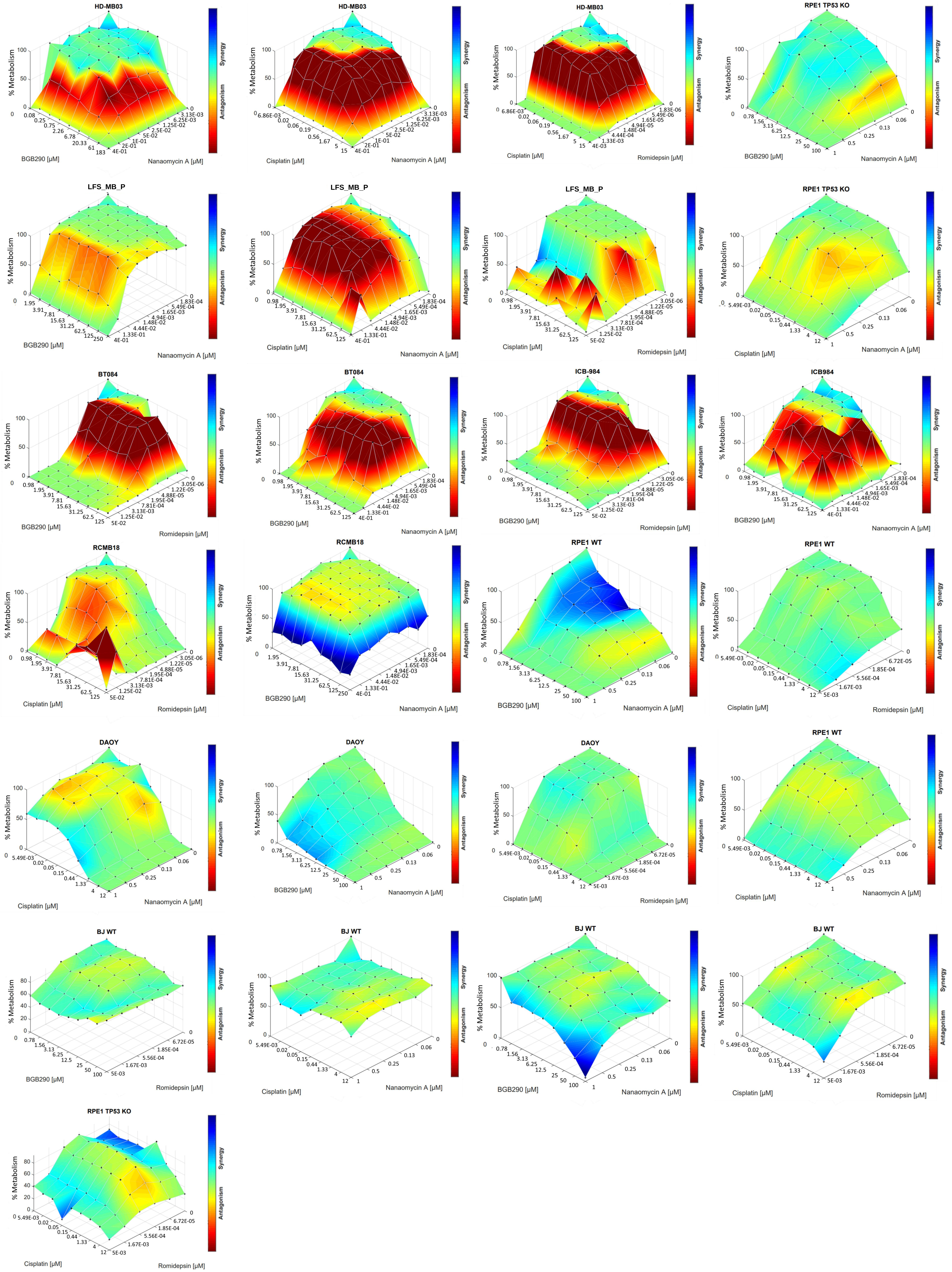

### Supplementary figure 5

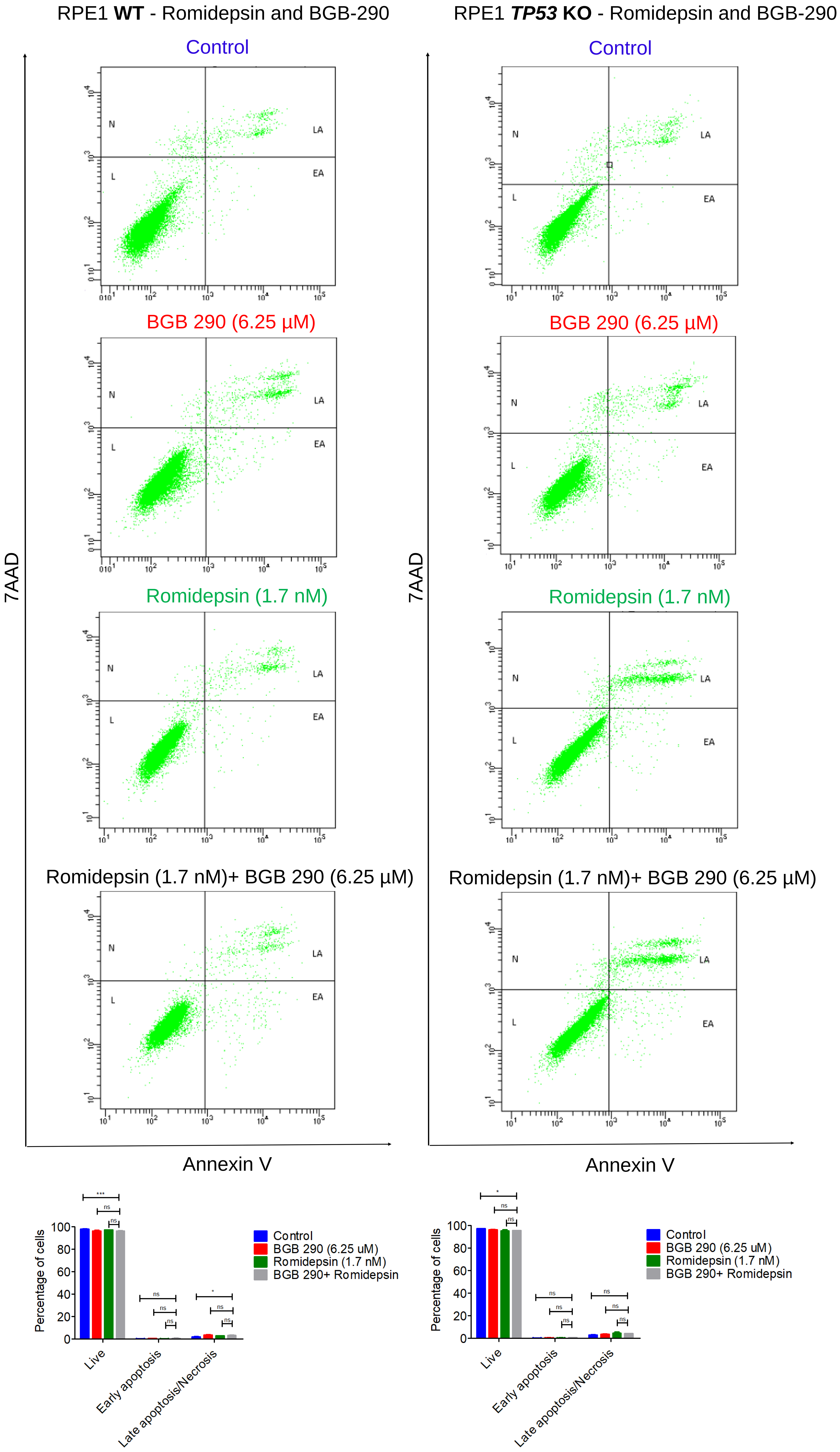

### Supplementary figure 6

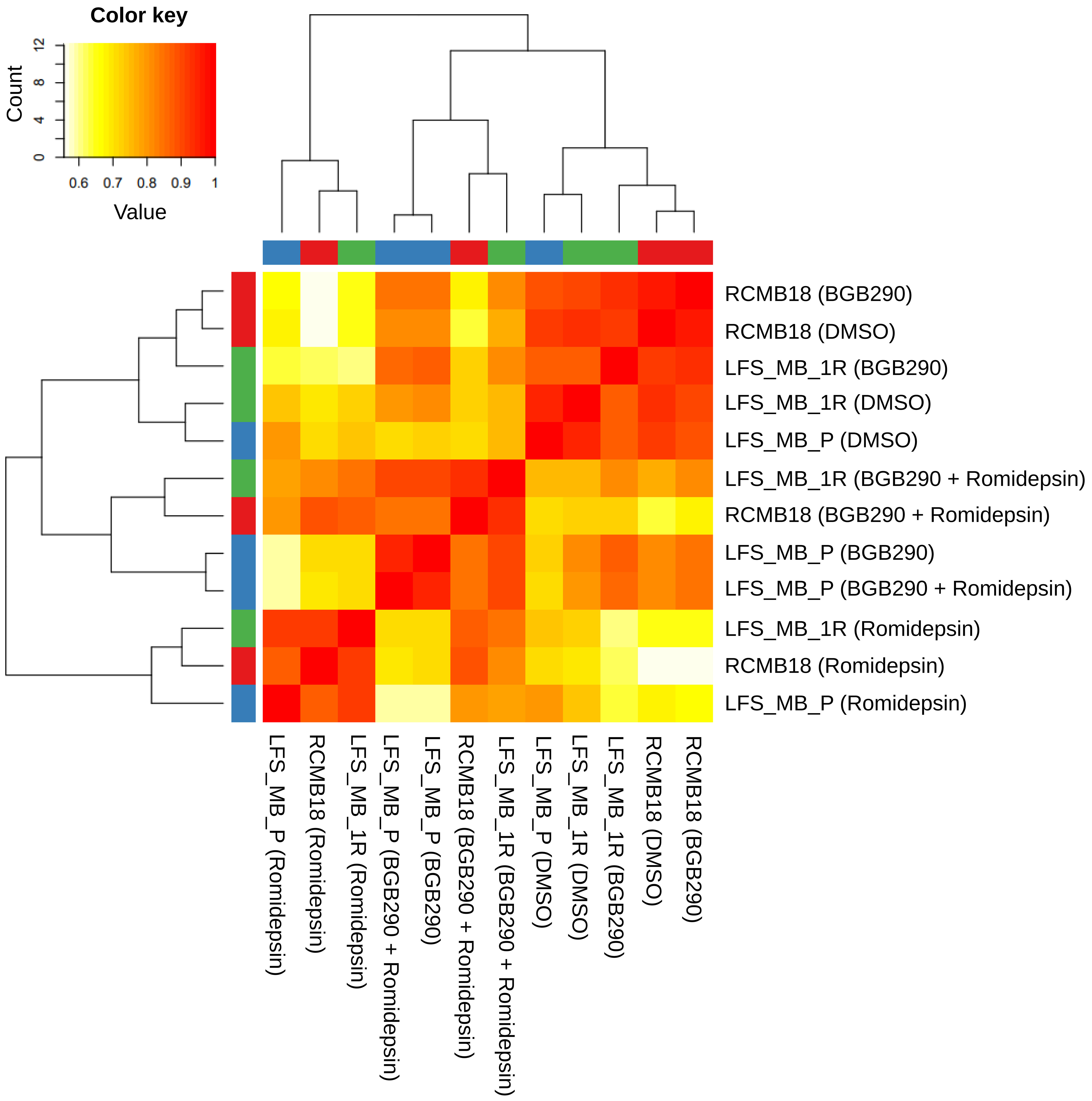
