## Supplementary table 1 for "A synergistic interaction between HDAC- and PARP inhibitors in childhood tumors with chromothripsis"

Supplementary table 1
Double treatment vs single treatment

| Probe ID | Symbol | Gene name | P value | Fold change |
|---|---|---|---|---|
| TC0500007292.hg.1 | NIM1K | NIM1 serine/threonine protein kinase | 1.05E-04 | 5.02 |
| HTA2-neg-47424007_st | NA | NA | 3.44E-03 | 4.11 |
| HTA2-pos-3475282_st | NA | NA | 3.30E-03 | 3.24 |
| TC0X00007013.hg.1 | MPC1L | mitochondrial pyruvate carrier 1-like | 5.22E-03 | 3.21 |
| TC0200010447.hg.1 | CASP8 | caspase 8, apoptosis-related cysteine peptidase | 3.54E-03 | 2.46 |
| TC0400008390.hg.1 | LRIT3 | leucine-rich repeat, immunoglobulin-like and transmembrane domains 3 | 1.86E-03 | 2.41 |
| TC1700011905.hg.1 | DNAH17 | dynein, axonemal, heavy chain 17 | 1.81E-04 | 2.40 |
| TC0600012064.hg.1 | GCM1 | glial cells missing homolog 1 (Drosophila) | 2.81E-03 | 2.39 |
| TC0100015789.hg.1 | POGZ | Transcript Identified by AceView, Entrez Gene ID(s) 23126 | 3.64E-04 | 2.38 |
| TC1300010039.hg.1 | NEK5 | NIMA-related kinase 5 | 3.39E-03 | 2.36 |
| TC0900008222.hg.1 | STX17 | syntaxin 17 | 1.08E-03 | 2.29 |
| TC1700012355.hg.1 | KRBA2 | KRAB-A domain containing 2 | 5.98E-03 | 2.28 |
| HTA2-neg-47424044_st | NA | NA | 5.94E-03 | 2.24 |
| HTA2-neg-47424360_st | NA | NA | 2.12E-03 | 2.22 |
| TC0800010802.hg.1 | C8orf89 | chromosome 8 open reading frame 89 | 6.51E-04 | 2.20 |
| TC1500010745.hg.1 | POLR2M | polymerase (RNA) II (DNA directed) polypeptide M | 5.19E-03 | 2.20 |
| TC1500007409.hg.1 | GCNT3 | glucosaminyl (N-acetyl) transferase 3, mucin type | 6.48E-03 | 2.17 |
| TC2200007132.hg.1 | RFPL3 | ret finger protein-like 3 | 5.91E-05 | 2.17 |
| HTA2-neg-47424024_st | NA | NA | 2.45E-03 | 2.16 |
| TC0200010474.hg.1 | KIAA2012 | KIAA2012 | 5.20E-03 | 2.16 |
| TC1100007216.hg.1 | PRRG4 | proline rich Gla (G-carboxyglutamic acid) 4 (transmembrane) | 7.43E-03 | 2.15 |
| TC0400012977.hg.1 | SH3D19 | SH3 domain containing 19 | 3.74E-03 | 2.09 |
| HTA2-neg-47422090_st | NA | NA | 1.57E-03 | 2.09 |
| TC2200008513.hg.1 | RFPL3S | RFPL3 antisense | 2.62E-04 | 2.07 |
| HTA2-neg-47420194_st | NA | NA | 2.88E-03 | 2.06 |
| TC0600007262.hg.1 | HIST1H3A | histone cluster 1, H3a | 3.81E-03 | 2.05 |
| TC0600007293.hg.1 | HIST1H2BI | histone cluster 1, H2bi | 1.27E-03 | 2.03 |
| TC0200011129.hg.1 | SAG | S-antigen; retina and pineal gland (arrestin) | 1.62E-03 | 2.02 |
| TC0200007038.hg.1 | DRC1 | dynein regulatory complex subunit 1 | 3.77E-04 | 2.02 |
| TC1700006813.hg.1 | ODF4 | outer dense fiber of sperm tails 4 | 3.75E-03 | 2.02 |
| TC0300012771.hg.1 | TM4SF18 | Transcript Identified by AceView, Entrez Gene ID(s) 116441 | 3.33E-03 | 2.01 |
| HTA2-neg-47422060_st | NA | NA | 2.74E-03 | 2.00 |
| TC1700007871.hg.1 | HSPB9 | heat shock protein, alpha-crystallin-related, B9 | 1.95E-03 | 1.99 |
| TC0300009772.hg.1 | AHSG | alpha-2-HS-glycoprotein | 2.72E-03 | 1.98 |
| TC1500009998.hg.1 | GOLGA6A | golgin A6 family, member A | 7.39E-04 | 1.98 |
| TC0800012350.hg.1 | MTBP | MDM2 binding protein | 1.41E-03 | 1.96 |
| TC1400010637.hg.1 | C14orf178 | chromosome 14 open reading frame 178 | 3.01E-03 | 1.95 |
| 23067837 | NA | NA | 1.30E-03 | 1.95 |
| TC0200007418.hg.1 | SLC3A1 | solute carrier family 3 (amino acid transporter heavy chain), member 1 | 1.95E-04 | 1.94 |
| TC1100010044.hg.1 | ST5 | Transcript Identified by AceView, Entrez Gene ID(s) 6764 | 1.83E-04 | 1.94 |
| TC1400010749.hg.1 | LINC01599 | long intergenic non-protein coding RNA 1599 | 8.21E-03 | 1.92 |
| TC0200016417.hg.1 | CENPA | centromere protein A | 9.44E-05 | 1.88 |
| TC1700010550.hg.1 | C17orf98 | chromosome 17 open reading frame 98 | 4.62E-03 | 1.86 |
| HTA2-neg-47424023_st | NA | NA | 3.33E-03 | 1.86 |
| HTA2-neg-47420912_st | NA | NA | 3.54E-03 | 1.84 |
| TC0900007069.hg.1 | C9orf131 | chromosome 9 open reading frame 131 | 6.88E-03 | 1.81 |
| TC0200008268.hg.1 | GNLY | granulysin | 3.47E-03 | 1.79 |
| TC0100015586.hg.1 | PDZK1 | PDZ domain containing 1 | 2.56E-04 | 1.79 |
| TC0100008773.hg.1 | MSH4 | mutS homolog 4 | 1.47E-03 | 1.78 |
| HTA2-neg-47423805_st | NA | NA | 3.64E-03 | 1.78 |
| HTA2-neg-47422085_st | NA | NA | 1.72E-03 | 1.78 |
| TC1500010769.hg.1 | GOLGA6C | golgin A6 family, member C | 9.92E-03 | 1.77 |
| TC0600014246.hg.1 | C6orf229 | chromosome 6 open reading frame 229 | 2.39E-03 | 1.77 |
| TC1700011117.hg.1 | LINC00483 | long intergenic non-protein coding RNA 483 | 6.29E-04 | 1.77 |
| TC1100012352.hg.1 | TMPRSS5 | transmembrane protease, serine 5 | 2.28E-03 | 1.76 |
| TC0900010370.hg.1 | TRPM3 | transient receptor potential cation channel, subfamily M, member 3 | 8.33E-03 | 1.76 |
| TC0700013525.hg.1 | FAM126A | family with sequence similarity 126, member A | 8.76E-04 | 1.76 |
| TC0200010445.hg.1 | CASP10 | caspase 10 | 4.70E-03 | 1.75 |
| TC0100009913.hg.1 | ADAMTSL4 | ADAMTS like 4 | 4.73E-03 | 1.74 |
| TC0300011997.hg.1 | BTLA | B and T lymphocyte associated | 5.15E-03 | 1.74 |
| TC1200012660.hg.1 | SLC35E3 | solute carrier family 35, member E3 | 8.44E-03 | 1.73 |
| HTA2-neg-47424704_st | NA | NA | 8.15E-03 | 1.72 |
| TC0500008654.hg.1 | LEAP2 | liver expressed antimicrobial peptide 2 | 2.06E-04 | 1.72 |
| TC1600007189.hg.1 | VWA3A | von Willebrand factor A domain containing 3A | 9.81E-04 | 1.71 |
| TC0100010526.hg.1 | TBX19 | T-box 19 | 3.99E-03 | 1.71 |
| TC1200009547.hg.1 | SLC6A13 | solute carrier family 6 (neurotransmitter transporter), member 13 | 1.03E-03 | 1.71 |
| TC0100018360.hg.1 | DISC1 | disrupted in schizophrenia 1 | 1.60E-03 | 1.70 |
| HTA2-neg-47424705_st | NA | NA | 2.69E-03 | 1.69 |
| TC0500011296.hg.1 | ACOT12 | acyl-CoA thioesterase 12 | 1.68E-04 | 1.69 |
| TC1200009220.hg.1 | CCDC62 | coiled-coil domain containing 62 | 8.64E-03 | 1.69 |
| TC0100009936.hg.1 | BNIPL | BCL2/adenovirus E1B 19kD interacting protein like | 5.80E-03 | 1.69 |
| TC1200008223.hg.1 | GLIPR1L2 | GLI pathogenesis-related 1 like 2 | 9.30E-03 | 1.68 |
| TC1100013093.hg.1 | BCO2 | beta-carotene oxygenase 2 | 8.52E-03 | 1.67 |
| TC0200016751.hg.1 | LOC100130691 | uncharacterized LOC100130691 | 5.53E-03 | 1.66 |
| TC0200016661.hg.1 | C2orf61 | chromosome 2 open reading frame 61 | 1.51E-03 | 1.66 |
| TC0200012071.hg.1 | UCN | urocortin | 4.36E-04 | 1.65 |
| HTA2-neg-47422095_st | NA | NA | 8.42E-04 | 1.65 |
| TC1700012241.hg.1 | TBC1D3L | TBC1 domain family, member 3L | 4.77E-03 | 1.65 |
| TSUnmapped00000145.hg.1 | SAG | S-antigen; retina and pineal gland (arrestin) | 1.00E-04 | 1.64 |
| TC0X00009089.hg.1 | FAM9C | family with sequence similarity 9, member C | 9.64E-03 | 1.64 |
| TC0600009623.hg.1 | ECT2L | epithelial cell transforming 2 like | 9.77E-03 | 1.63 |
| TC0X00007562.hg.1 | ITGB1BP2 | integrin beta 1 binding protein (melusin) 2 | 5.30E-04 | 1.63 |
| TC1900006602.hg.1 | ZNF555 | zinc finger protein 555 | 8.88E-03 | 1.63 |
| TC0200010837.hg.1 | ANKZF1 | ankyrin repeat and zinc finger domain containing 1 | 8.90E-03 | 1.61 |
| TC1500010712.hg.1 | GOLGA8N | golgin A8 family, member N | 9.53E-03 | 1.61 |
| HTA2-neg-47419220_st | NA | NA | 5.60E-04 | 1.59 |
| TC1200012744.hg.1 | C1R | complement component 1, r subcomponent | 5.41E-03 | 1.59 |
| TC0100016445.hg.1 | SERPINC1 | serpin peptidase inhibitor, clade C (antithrombin), member 1 | 5.03E-03 | 1.58 |
| TC1100010733.hg.1 | SPI1 | Spi-1 proto-oncogene | 2.27E-04 | 1.58 |
| TC0700012228.hg.1 | LAMB4 | laminin, beta 4 | 3.01E-03 | 1.58 |
| TC0300013941.hg.1 | MKRN2OS | MKRN2 opposite strand | 4.53E-04 | 1.58 |
| AFFX-r2-Ec-bioB-5_at | NA | NA | 6.75E-03 | 1.58 |
| TC0900007163.hg.1 | FRMPD1 | FERM and PDZ domain containing 1 | 4.72E-03 | 1.57 |
| TC1700009746.hg.1 | MYH3 | myosin, heavy chain 3, skeletal muscle, embryonic | 7.83E-03 | 1.56 |
| AFFX-r2-Ec-bioB-3_at | NA | NA | 7.40E-03 | 1.56 |
| TC1100011029.hg.1 | VWCE | von Willebrand factor C and EGF domains | 5.53E-04 | 1.56 |
| TSUnmapped00000176.hg.1 | SAG | S-antigen; retina and pineal gland (arrestin) | 8.78E-03 | 1.56 |
| AFFX-r2-Ec-bioC-3_at | NA | NA | 6.90E-03 | 1.56 |
| TC0500010592.hg.1 | C6 | complement component 6 | 2.34E-03 | 1.56 |
| TC0800008946.hg.1 | TG | thyroglobulin | 2.32E-03 | 1.56 |
| TC1000009916.hg.1 | CUBN | cubilin (intrinsic factor-cobalamin receptor) | 6.40E-03 | 1.56 |
| TC1500008271.hg.1 | WDR93 | WD repeat domain 93 | 4.63E-03 | 1.55 |
| TSUnmapped00000141.hg.1 | HMBS | hydroxymethylbilane synthase | 2.92E-03 | 1.55 |
| TC2200008588.hg.1 | APOL3 | apolipoprotein L, 3 | 5.59E-03 | 1.55 |
| TC1600009192.hg.1 | MTRNR2L4 | MT-RNR2-like 4 | 7.88E-03 | 1.55 |
| TC0600010071.hg.1 | SLC22A1 | solute carrier family 22 (organic cation transporter), member 1 | 1.50E-03 | 1.55 |
| 23075460 | NA | NA | 1.69E-04 | 1.55 |
| HTA2-pos-PSR0X007730.hg.1 | NA | NA | 4.51E-03 | 1.54 |
| TC0900008667.hg.1 | OR1N2 | olfactory receptor, family 1, subfamily N, member 2 | 2.34E-03 | 1.54 |
| TC0600014251.hg.1 | ZBED9 | zinc finger, BED-type containing 9 | 2.72E-04 | 1.54 |
| HTA2-neg-47422763_st | NA | NA | 2.70E-03 | 1.54 |
| TC0600011805.hg.1 | USP49 | ubiquitin specific peptidase 49 | 4.43E-03 | 1.54 |
| TC0X00011291.hg.1 | PAGE2 | P antigen family, member 2 (prostate associated) | 4.94E-03 | 1.54 |
| TC0700013471.hg.1 | MGAM | maltase-glucoamylase | 9.81E-03 | 1.54 |
| TC1600009545.hg.1 | ABCC6 | ATP binding cassette subfamily C member 6 | 7.14E-03 | 1.52 |

| | | | | |
|---|---|---|---|---|
| TC1700008328.hg.1 | STXBP4 | syntaxin binding protein 4 | 8.74E-03 | 1.52 |
| TC0100009274.hg.1 | AMY1C | amylase, alpha 1C (salivary) | 1.64E-03 | 1.52 |
| HTA2-neg-47420204_st | NA | NA | 7.74E-03 | 1.52 |
| TC2000007094.hg.1 | TTLL9 | tubulin tyrosine ligase-like family member 9 | 8.91E-03 | 1.52 |
| TC0200006537.hg.1 | COLEC11 | collectin subfamily member 11 | 4.33E-03 | 1.51 |
| HTA2-neg-47420566_st | NA | NA | 5.80E-03 | 1.51 |
| TC0700008063.hg.1 | SPDYE5 | speedy/RINGO cell cycle regulator family member E5 | 4.83E-03 | 1.51 |
| AFFX-BioB-3_at | NA | NA | 8.91E-03 | 1.50 |
| TC1400009924.hg.1 | EML5 | echinoderm microtubule associated protein like 5 | 1.01E-03 | 1.50 |
| TC0X00007371.hg.1 | PAGE2B | P antigen family, member 2B | 6.90E-03 | 1.50 |

Double treatment vs control

| Probe ID | Symbol | Gene name | P value | Fold change |
|---|---|---|---|---|
| TC1200011845.hg.1 | SELPLG | selectin P ligand | 3.20E-05 | 10.41 |
| TC0500007292.hg.1 | NIM1K | NIM1 serine/threonine protein kinase | 1.68E-04 | 7.65 |
| TC1200012643.hg.1 | ERBB3 | erb-b2 receptor tyrosine kinase 3 | 2.44E-04 | 6.27 |
| HTA2-neg-47422282_st | NA | NA | 2.93E-05 | 6.10 |
| HTA2-neg-47424007_st | NA | NA | 5.54E-03 | 5.81 |
| TC1400006697.hg.1 | DHRS2 | dehydrogenase/reductase (SDR family) member 2 | 1.08E-03 | 5.21 |
| TC0X00007013.hg.1 | MPC1L | mitochondrial pyruvate carrier 1-like | 1.95E-04 | 5.11 |
| TC0300014032.hg.1 | POPDC2 | popeye domain containing 2 | 3.09E-06 | 5.07 |
| TC0600014190.hg.1 | ZC2HC1B | zinc finger, C2HC-type containing 1B | 1.75E-03 | 4.98 |
| 23076543 | NA | NA | 3.55E-05 | 4.76 |
| HTA2-neg-47424020_st | NA | NA | 2.71E-03 | 4.64 |
| TC0600007262.hg.1 | HIST1H3A | histone cluster 1, H3a | 1.27E-05 | 4.35 |
| TC0700013472.hg.1 | MGAM | maltase-glucoamylase | 4.35E-07 | 4.35 |
| TC1900006977.hg.1 | ICAM1 | intercellular adhesion molecule 1 | 5.47E-04 | 4.34 |
| HTA2-neg-47424041_st | NA | NA | 6.56E-03 | 4.24 |
| TC0300012007.hg.1 | CD200R1 | CD200 receptor 1 | 2.64E-05 | 4.19 |
| TC1300008048.hg.1 | TEX29 | testis expressed 29 | 4.70E-05 | 4.02 |
| TC1000006911.hg.1 | TMEM236 | transmembrane protein 236 | 4.96E-04 | 4.02 |
| TC1700008673.hg.1 | CACNG5 | calcium channel, voltage-dependent, gamma subunit 5 | 5.13E-03 | 3.90 |
| TC0400008240.hg.1 | DAPP1 | dual adaptor of phosphotyrosine and 3-phosphoinositides | 1.20E-03 | 3.79 |
| TC0600012064.hg.1 | GCM1 | glial cells missing homolog 1 (Drosophila) | 9.66E-06 | 3.76 |
| TC0300014084.hg.1 | CRYGS | crystallin gamma S | 1.84E-04 | 3.70 |
| TC1100009075.hg.1 | NCAM1 | Transcript Identified by AceView, Entrez Gene ID(s) 4684 | 2.50E-03 | 3.60 |
| TC1200007053.hg.1 | SPX | spexin hormone | 1.37E-03 | 3.57 |
| TC1200008686.hg.1 | CHST11 | Transcript Identified by AceView, Entrez Gene ID(s) 50515 | 4.36E-04 | 3.55 |
| TC0700009411.hg.1 | MGAM2 | maltase-glucoamylase 2 (putative) | 8.58E-06 | 3.54 |
| TC1600006593.hg.1 | RAB26 | RAB26, member RAS oncogene family | 1.09E-04 | 3.49 |
| TC1600007147.hg.1 | TMEM159 | transmembrane protein 159 | 3.01E-06 | 3.45 |
| TC0700008495.hg.1 | BUD31 | Transcript Identified by AceView, Entrez Gene ID(s) 8896 | 1.86E-06 | 3.44 |
| TC1300010039.hg.1 | NEK5 | NIMA-related kinase 5 | 9.38E-05 | 3.44 |
| TC1900009240.hg.1 | GNG7 | guanine nucleotide binding protein (G protein), gamma 7 | 1.29E-03 | 3.43 |
| TC1100007216.hg.1 | PRRG4 | proline rich Gla (G-carboxyglutamic acid) 4 (transmembrane) | 5.05E-06 | 3.43 |
| TC0300009412.hg.1 | SERPINI1 | serpin peptidase inhibitor, clade I (neuroserpin), member 1 | 2.50E-04 | 3.43 |
| 23070242 | NA | NA | 2.16E-05 | 3.42 |
| TC1100010123.hg.1 | DKK3 | dickkopf WNT signaling pathway inhibitor 3 | 1.03E-05 | 3.40 |
| TC2000009964.hg.1 | SDCBP2 | syndecan binding protein (syntenin) 2 | 1.52E-06 | 3.38 |
| TC0600009597.hg.1 | TNFAIP3 | tumor necrosis factor, alpha-induced protein 3 | 2.28E-03 | 3.38 |
| TC1600008712.hg.1 | IRF8 | interferon regulatory factor 8 | 2.21E-04 | 3.36 |
| TC0600011133.hg.1 | HIST1H2BE | Memczak2013 ANTISENSE, CDS, coding, upstream_start, UTR3, UTR5 best transcript NM_003523 | 2.35E-03 | 3.32 |
| TC0500007077.hg.1 | NPR3 | natriuretic peptide receptor 3 | 2.45E-04 | 3.31 |
| HTA2-pos-47421925_st | NA | NA | 6.55E-04 | 3.29 |
| HTA2-neg-47422060_st | NA | NA | 1.27E-04 | 3.28 |
| TC2000009905.hg.1 | EFCAB8 | EF-hand calcium binding domain 8 | 2.75E-03 | 3.28 |
| TC0700010899.hg.1 | POLR2J4 | polymerase (RNA) II (DNA directed) polypeptide J4, pseudogene | 1.19E-05 | 3.25 |
| TC0700008506.hg.1 | CYP3A43 | cytochrome P450, family 3, subfamily A, polypeptide 43 | 5.94E-05 | 3.21 |
| TC1500010745.hg.1 | POLR2M | polymerase (RNA) II (DNA directed) polypeptide M | 1.38E-04 | 3.21 |
| TC0900008222.hg.1 | STX17 | syntaxin 17 | 5.78E-04 | 3.21 |
| HTA2-neg-47424360_st | NA | NA | 5.25E-03 | 3.20 |
| TC1900010016.hg.1 | ISYNA1 | inositol-3-phosphate synthase 1 | 7.99E-06 | 3.15 |
| TC0300009916.hg.1 | HES1 | hes family bHLH transcription factor 1 | 1.59E-05 | 3.15 |
| TC1300008813.hg.1 | KCTD4 | potassium channel tetramerization domain containing 4 | 3.33E-04 | 3.11 |
| TC0X00009025.hg.1 | ANOS1 | anosmin 1 | 3.78E-04 | 3.11 |
| TC0600007263.hg.1 | HIST1H4A | histone cluster 1, H4a | 3.23E-04 | 3.10 |
| TC0500012017.hg.1 | IRF1 | interferon regulatory factor 1 | 6.72E-03 | 3.08 |
| TC2100008297.hg.1 | SIK1 | salt-inducible kinase 1 | 7.91E-03 | 3.08 |
| TC1700010550.hg.1 | C17orf98 | chromosome 17 open reading frame 98 | 3.17E-05 | 3.07 |
| TC0600014151.hg.1 | SMIM8 | small integral membrane protein 8 | 2.24E-03 | 3.03 |
| HTA2-neg-47424044_st | NA | NA | 1.79E-03 | 3.02 |
| TC0800010802.hg.1 | C8orf89 | chromosome 8 open reading frame 89 | 1.26E-03 | 3.01 |
| TC0400012977.hg.1 | SH3D19 | SH3 domain containing 19 | 6.80E-05 | 3.00 |
| HTA2-pos-47421982_st | NA | NA | 1.39E-04 | 3.00 |
| HTA2-neg-47421192_st | NA | NA | 1.32E-04 | 2.97 |
| TC0700013525.hg.1 | FAM126A | family with sequence similarity 126, member A | 1.34E-04 | 2.97 |
| TC0600007274.hg.1 | HIST1H2BD | histone cluster 1, H2bd | 1.80E-06 | 2.96 |
| TC0700013424.hg.1 | GS1-259H13.2 | transmembrane protein 225-like | 7.98E-06 | 2.96 |
| HTA2-neg-47423838_st | NA | NA | 1.24E-04 | 2.94 |
| HTA2-pos-3475282_st | NA | NA | 4.40E-03 | 2.93 |
| TC1900008931.hg.1 | RFPL4A | ret finger protein-like 4A | 9.34E-03 | 2.89 |
| TC0100009442.hg.1 | WNT2B | wingless-type MMTV integration site family, member 2B | 3.41E-05 | 2.86 |
| TC0700013587.hg.1 | SHFM1 | split hand/foot malformation (ectrodactyly) type 1 | 5.20E-04 | 2.85 |
| TC1700008635.hg.1 | RGS9 | regulator of G-protein signaling 9 | 7.85E-03 | 2.84 |
| HTA2-pos-47421926_st | NA | NA | 5.49E-04 | 2.83 |
| HTA2-pos-47421981_st | NA | NA | 4.89E-04 | 2.83 |
| TC1300006676.hg.1 | RASL11A | RAS-like, family 11, member A | 4.69E-04 | 2.82 |
| TC2200008513.hg.1 | RFPL3S | RFPL3 antisense | 1.23E-05 | 2.80 |
| TSUnmapped00000401.hg.1 | INPP5D | inositol polyphosphate-5-phosphatase D | 4.09E-03 | 2.80 |
| TC0900008219.hg.1 | NR4A3 | nuclear receptor subfamily 4, group A, member 3 | 7.69E-04 | 2.79 |
| HTA2-pos-PSR02023992.hg.1 | NA | NA | 4.80E-04 | 2.78 |
| TC0100011378.hg.1 | RASSF5 | Ras association (RalGDS/AF-6) domain family member 5 | 4.44E-05 | 2.76 |
| TC0200009078.hg.1 | EPB41L5 | erythrocyte membrane protein band 4.1 like 5 | 4.98E-04 | 2.76 |
| TC0600007293.hg.1 | HIST1H2BI | histone cluster 1, H2bi | 5.82E-04 | 2.76 |
| TC0X00008747.hg.1 | GABRQ | gamma-aminobutyric acid (GABA) A receptor, theta | 7.57E-05 | 2.69 |
| TC1900006588.hg.1 | GADD45B | growth arrest and DNA-damage-inducible, beta | 5.25E-05 | 2.67 |
| TC1000007895.hg.1 | TSPAN15 | tetraspanin 15 | 9.53E-04 | 2.67 |
| TC0200016661.hg.1 | C2orf61 | chromosome 2 open reading frame 61 | 3.31E-04 | 2.67 |
| TC0700013442.hg.1 | LSMEM1 | leucine-rich single-pass membrane protein 1 | 1.54E-04 | 2.66 |
| HTA2-neg-47423602_st | NA | NA | 2.51E-03 | 2.66 |
| TC2000007202.hg.1 | ACSS2 | acyl-CoA synthetase short-chain family member 2 | 6.92E-03 | 2.64 |
| TC1900010782.hg.1 | ATP1A3 | ATPase, Na+/K+ transporting, alpha 3 polypeptide | 6.50E-06 | 2.64 |
| HTA2-pos-2960249_st | NA | NA | 1.40E-04 | 2.64 |
| TC0600010797.hg.1 | MAK | male germ cell-associated kinase | 2.98E-04 | 2.63 |
| TC0700013601.hg.1 | RASA4B | RAS p21 protein activator 4B | 1.38E-04 | 2.63 |
| TC0700013603.hg.1 | RASA4 | RAS p21 protein activator 4 | 3.72E-04 | 2.61 |
| TC0900012173.hg.1 | GARNL3 | GTPase activating Rap/RanGAP domain-like 3 | 7.49E-05 | 2.61 |
| TC0100009913.hg.1 | ADAMTSL4 | ADAMTS like 4 | 9.37E-05 | 2.60 |
| TC2200007132.hg.1 | RFPL3 | ret finger protein-like 3 | 7.23E-04 | 2.60 |
| TC0600012502.hg.1 | GJB7 | gap junction protein beta 7 | 2.05E-05 | 2.59 |
| TC1600008661.hg.1 | CRISPLD2 | cysteine-rich secretory protein LCCL domain containing 2 | 1.11E-04 | 2.59 |
| TC1400010749.hg.1 | LINC01599 | long intergenic non-protein coding RNA 1599 | 4.26E-04 | 2.58 |
| TC0400008389.hg.1 | RRH | retinal pigment epithelium-derived rhodopsin homolog | 4.16E-05 | 2.57 |
| TC1900011817.hg.1 | ZNF773 | zinc finger protein 773 | 6.35E-06 | 2.57 |
| TC0500007050.hg.1 | PDZD2 | PDZ domain containing 2 | 2.18E-05 | 2.57 |
| TC0100018207.hg.1 | TSSK3 | testis-specific serine kinase 3 | 6.36E-04 | 2.55 |

| | | | | |
|---|---|---|---|---|
| TC0200007458.hg.1 | EPAS1 | endothelial PAS domain protein 1 | 1.13E-03 | 2.55 |
| HTA2-neg-47424241_st | NA | NA | 1.99E-03 | 2.55 |
| TC0400008390.hg.1 | LRIT3 | leucine-rich repeat, immunoglobulin-like and transmembrane domains 3 | 5.28E-05 | 2.55 |
| TC0100014191.hg.1 | RAB3B | RAB3B, member RAS oncogene family | 3.08E-04 | 2.51 |
| TSUnmapped00000108.hg.1 | RPS6KA1 | ribosomal protein S6 kinase, 90kDa, polypeptide 1 | 3.97E-05 | 2.50 |
| TC0300012771.hg.1 | TM4SF18 | Transcript Identified by AceView, Entrez Gene ID(s) 116441 | 2.76E-03 | 2.50 |
| TC1700011905.hg.1 | DNAH17 | dynein, axonemal, heavy chain 17 | 2.70E-03 | 2.50 |
| TC2000009886.hg.1 | PANK2 | pantothenate kinase 2 | 2.55E-05 | 2.50 |
| TC0100009936.hg.1 | BNIPL | BCL2/adenovirus E1B 19kD interacting protein like | 3.58E-05 | 2.49 |
| TC0300007050.hg.1 | C3orf35 | chromosome 3 open reading frame 35 | 6.45E-05 | 2.49 |
| TC1500010744.hg.1 | GCOM1 | GRINL1A complex locus 1 | 6.20E-05 | 2.48 |
| TC0200013595.hg.1 | MGAT4A | mannosyl (alpha-1,3-)-glycoprotein beta-1,4-N-acetylglucosaminyltransferase, isozyme A | 3.55E-04 | 2.48 |
| TC0500008785.hg.1 | EGR1 | early growth response 1 | 3.38E-04 | 2.48 |
| TC0400007360.hg.1 | DCAF4L1 | DDB1 and CUL4 associated factor 4-like 1 | 1.24E-03 | 2.46 |
| TC0500008356.hg.1 | KCNN2 | potassium channel, calcium activated intermediate/small conductance subfamily N alpha, member 2 | 1.10E-03 | 2.46 |
| TC1500009606.hg.1 | MYO1E | Transcript Identified by AceView, Entrez Gene ID(s) 4643 | 1.80E-03 | 2.46 |
| TC0900007069.hg.1 | C9orf131 | chromosome 9 open reading frame 131 | 2.26E-03 | 2.46 |
| HTA2-pos-47421984_st | NA | NA | 1.59E-03 | 2.45 |
| TC0500013280.hg.1 | ZDHHC11B | zinc finger, DHHC-type containing 11B | 4.91E-04 | 2.45 |
| TC1900008826.hg.1 | CACNG7 | calcium channel, voltage-dependent, gamma subunit 7 | 2.17E-04 | 2.45 |
| TC0500007895.hg.1 | CMYA5 | cardiomyopathy associated 5 | 1.40E-05 | 2.44 |
| TC1700006763.hg.1 | ATP1B2 | ATPase, Na+/K+ transporting, beta 2 polypeptide | 7.63E-05 | 2.44 |
| TC1500007409.hg.1 | GCNT3 | glucosaminyl (N-acetyl) transferase 3, mucin type | 8.31E-04 | 2.44 |
| HTA2-pos-47421983_st | NA | NA | 1.04E-03 | 2.44 |
| TC1000009535.hg.1 | IDI2 | isopentenyl-diphosphate delta isomerase 2 | 4.19E-03 | 2.43 |
| HTA2-neg-47423726_st | NA | NA | 2.18E-04 | 2.42 |
| TC2000008910.hg.1 | EIF2S2 | Zhang2013 ALT_ACCEPTOR, ALT_DONOR, coding, INTERNAL, intronic best transcript NM_003908 | 6.47E-03 | 2.42 |
| TC1500009998.hg.1 | GOLGA6A | golgin A6 family, member A | 2.79E-04 | 2.42 |
| TC1400007443.hg.1 | HSPA2 | heat shock 70kDa protein 2 | 1.33E-03 | 2.42 |
| 23067837 | NA | NA | 3.87E-04 | 2.41 |
| TC0200015869.hg.1 | AP1S3 | adaptor-related protein complex 1 sigma 3 subunit | 4.36E-03 | 2.41 |
| TC0200006627.hg.1 | ID2 | inhibitor of DNA binding 2, dominant negative helix-loop-helix protein | 2.47E-03 | 2.41 |
| TC0100008334.hg.1 | ZYG11A | zyg-11 family member A, cell cycle regulator | 2.69E-03 | 2.39 |
| TC1400010637.hg.1 | C14orf178 | chromosome 14 open reading frame 178 | 4.00E-04 | 2.39 |
| TC0800010685.hg.1 | MYBL1 | v-myb avian myeloblastosis viral oncogene homolog-like 1 | 2.08E-04 | 2.39 |
| TC0700009472.hg.1 | EPHB6 | EPH receptor B6 | 9.75E-04 | 2.39 |
| TC1100010505.hg.1 | ABTB2 | ankyrin repeat and BTB (POZ) domain containing 2 | 2.00E-04 | 2.38 |
| TC1100007729.hg.1 | DTX4 | deltex 4, E3 ubiquitin ligase | 1.83E-05 | 2.38 |
| TC0200015226.hg.1 | HIBCH | Transcript Identified by AceView, Entrez Gene ID(s) 26275 | 5.24E-05 | 2.37 |
| TC0200007048.hg.1 | MAPRE3 | microtubule-associated protein, RP/EB family, member 3 | 1.04E-04 | 2.37 |
| TC0100018485.hg.1 | ADAMTSL4-AS1 | ADAMTSL4 antisense RNA 1 | 1.06E-04 | 2.37 |
| TC0300008853.hg.1 | TMEM108 | transmembrane protein 108 | 4.22E-04 | 2.37 |
| TC1500010369.hg.1 | MFGE8 | milk fat globule-EGF factor 8 protein | 3.05E-05 | 2.36 |
| TC0500008654.hg.1 | LEAP2 | liver expressed antimicrobial peptide 2 | 9.14E-05 | 2.36 |
| TC0900008793.hg.1 | ZBTB34 | zinc finger and BTB domain containing 34 | 1.57E-04 | 2.36 |
| TC0200015402.hg.1 | FAM126B | family with sequence similarity 126, member B | 4.93E-05 | 2.36 |
| TC0200006671.hg.1 | GRHL1 | grainyhead-like transcription factor 1 | 1.27E-03 | 2.35 |
| TC1000010432.hg.1 | RASGEF1A | RasGEF domain family member 1A | 7.47E-03 | 2.35 |
| TC1500007615.hg.1 | MAP2K1 | Transcript Identified by AceView, Entrez Gene ID(s) 5604 | 6.38E-05 | 2.35 |
| TC0500012588.hg.1 | HAVCR1 | hepatitis A virus cellular receptor 1 | 8.22E-04 | 2.35 |
| TC0X00006593.hg.1 | CLCN4 | chloride channel, voltage-sensitive 4 | 8.48E-06 | 2.35 |
| HTA2-neg-47423769_st | NA | NA | 6.85E-03 | 2.35 |
| TC0X00007562.hg.1 | ITGB1BP2 | integrin beta 1 binding protein (melusin) 2 | 5.22E-05 | 2.35 |
| HTA2-neg-47422979_st | NA | NA | 7.60E-03 | 2.34 |
| TC1900007399.hg.1 | TMEM59L | transmembrane protein 59-like | 7.47E-04 | 2.33 |
| TC1200012711.hg.1 | LINC00173 | long intergenic non-protein coding RNA 173 | 6.26E-04 | 2.33 |
| TC0200008870.hg.1 | BCL2L11 | BCL2-like 11 (apoptosis facilitator) | 1.18E-04 | 2.33 |
| TC1200008683.hg.1 | CHST11 | carbohydrate (chondroitin 4) sulfotransferase 11 | 2.58E-04 | 2.33 |
| TC0300009279.hg.1 | KCNAB1 | potassium channel, voltage gated subfamily A regulatory beta subunit 1 | 4.83E-04 | 2.33 |
| TC0600006870.hg.1 | SNRNP48 | small nuclear ribonucleoprotein, U11/U12 48KDa subunit | 3.14E-04 | 2.33 |
| TC2000007094.hg.1 | TTLL9 | tubulin tyrosine ligase-like family member 9 | 3.11E-05 | 2.33 |
| TC0100017844.hg.1 | NID1 | nidogen 1 | 1.98E-03 | 2.33 |
| TC0700011924.hg.1 | TMEM130 | transmembrane protein 130 | 6.06E-03 | 2.33 |
| TC0400011043.hg.1 | CDKL2 | cyclin-dependent kinase-like 2 (CDC2-related kinase) | 4.82E-04 | 2.32 |
| TC0300011525.hg.1 | RYBP | RING1 and YY1 binding protein | 2.65E-03 | 2.32 |
| TC1400008622.hg.1 | OR5AU1 | olfactory receptor, family 5, subfamily AU, member 1 | 3.22E-03 | 2.32 |
| TC0200008894.hg.1 | MERTK | MER proto-oncogene, tyrosine kinase | 9.13E-04 | 2.32 |
| TC1900006602.hg.1 | ZNF555 | zinc finger protein 555 | 5.75E-04 | 2.32 |
| TC0100009341.hg.1 | KIAA1324 | KIAA1324 | 1.97E-03 | 2.32 |
| TC1100009242.hg.1 | ABCG4 | ATP binding cassette subfamily G member 4 | 1.40E-04 | 2.32 |
| TC0500013282.hg.1 | ZDHHC11 | zinc finger, DHHC-type containing 11 | 3.27E-04 | 2.32 |
| TC0600014152.hg.1 | LINC01590 | long intergenic non-protein coding RNA 1590 | 5.75E-04 | 2.31 |
| TC1900010375.hg.1 | RHPN2 | rhophilin, Rho GTPase binding protein 2 | 9.80E-03 | 2.31 |
| TC0400011973.hg.1 | CLGN | calmegin | 2.06E-04 | 2.31 |
| TC1400008415.hg.1 | ZFYVE21 | zinc finger, FYVE domain containing 21 | 1.27E-05 | 2.31 |
| HTA2-neg-47422085_st | NA | NA | 1.19E-03 | 2.31 |
| TC0500011296.hg.1 | ACOT12 | acyl-CoA thioesterase 12 | 3.29E-05 | 2.31 |
| TC0100015436.hg.1 | SPAG17 | sperm associated antigen 17 | 5.37E-04 | 2.31 |
| TC1600009944.hg.1 | KCTD13 | potassium channel tetramerization domain containing 13 | 6.81E-05 | 2.29 |
| TC0200008268.hg.1 | GNLY | granulysin | 1.71E-04 | 2.29 |
| HTA2-pos-3295162_st | NA | NA | 8.25E-03 | 2.29 |
| TC2000007177.hg.1 | RALY | Jeck2013 ALT_DONOR, coding, INTERNAL, intronic best transcript NM_016732 | 3.72E-03 | 2.28 |
| TC1600008638.hg.1 | DNAAF1 | dynein, axonemal, assembly factor 1 | 2.92E-03 | 2.28 |
| TC1700011436.hg.1 | ERN1 | endoplasmic reticulum to nucleus signaling 1 | 3.01E-03 | 2.28 |
| TC0X00011174.hg.1 | PDZD4 | PDZ domain containing 4 | 1.18E-03 | 2.27 |
| TC0100015789.hg.1 | POGZ | Transcript Identified by AceView, Entrez Gene ID(s) 23126 | 2.33E-03 | 2.27 |
| HTA2-pos-PSR02023993.hg.1 | NA | NA | 1.05E-03 | 2.27 |
| TC1200009734.hg.1 | VAMP1 | vesicle associated membrane protein 1 | 1.08E-03 | 2.27 |
| TC0600011234.hg.1 | HIST1H4L | histone cluster 1, H4l | 4.44E-03 | 2.27 |
| TC1200012111.hg.1 | TAOK3 | TAO kinase 3 | 3.49E-03 | 2.27 |
| TC0700012099.hg.1 | RASA4 | Salzman2013 ALT_ACCEPTOR, ALT_DONOR, coding, INTERNAL, intronic best transcript NM_006989 | 6.29E-04 | 2.25 |
| TC1500010886.hg.1 | CALML4 | calmodulin-like 4 | 4.07E-05 | 2.24 |
| TC0400007876.hg.1 | THAP6 | THAP domain containing 6 | 3.51E-04 | 2.24 |
| TC2000009945.hg.1 | FAM209A | family with sequence similarity 209, member A | 8.35E-04 | 2.24 |
| TC0900010808.hg.1 | BICD2 | bicaudal D homolog 2 (Drosophila) | 8.64E-04 | 2.24 |
| TC0400010618.hg.1 | TEC | tec protein tyrosine kinase | 2.31E-04 | 2.24 |
| TC1700011117.hg.1 | LINC00483 | long intergenic non-protein coding RNA 483 | 6.76E-06 | 2.24 |
| TC1500010157.hg.1 | ADAMTS7 | ADAM metallopeptidase with thrombospondin type 1 motif 7 | 1.51E-04 | 2.23 |
| TC1700009746.hg.1 | MYH3 | myosin, heavy chain 3, skeletal muscle, embryonic | 3.64E-04 | 2.23 |
| TC1200007359.hg.1 | CNTN1 | contactin 1 | 4.86E-03 | 2.22 |
| TC0100013610.hg.1 | FAM229A | family with sequence similarity 229, member A | 5.26E-05 | 2.22 |
| TC0500009984.hg.1 | FLJ33360 | FLJ33360 protein | 2.39E-03 | 2.22 |
| TC1700009318.hg.1 | FAM101B | family with sequence similarity 101, member B | 2.16E-03 | 2.22 |
| TC1900009603.hg.1 | OLFM2 | olfactomedin 2 | 2.99E-04 | 2.21 |
| TC0200010837.hg.1 | ANKZF1 | ankyrin repeat and zinc finger domain containing 1 | 1.44E-03 | 2.21 |
| TC0100018418.hg.1 | IFFO2 | intermediate filament family orphan 2 | 3.33E-03 | 2.20 |
| TC0200010586.hg.1 | CPO | carboxypeptidase O | 1.32E-03 | 2.20 |
| TC0600008166.hg.1 | SLC25A27 | solute carrier family 25, member 27 | 3.61E-03 | 2.20 |
| TC1800008281.hg.1 | TMEM241 | transmembrane protein 241 | 1.14E-03 | 2.19 |
| 23076546 | NA | NA | 2.64E-03 | 2.19 |
| TC0500008541.hg.1 | TEX43 | testis expressed 43 | 1.53E-03 | 2.19 |
| TC0100012465.hg.1 | CPSF3L | cleavage and polyadenylation specific factor 3-like | 1.27E-04 | 2.19 |
| TSUnmapped00000109.hg.1 | ATG16L1 | autophagy related 16-like 1 | 7.09E-04 | 2.18 |
| TC0500009521.hg.1 | CPEB4 | cytoplasmic polyadenylation element binding protein 4 | 4.18E-04 | 2.18 |
| TC2100006659.hg.1 | CXADR | coxsackie virus and adenovirus receptor | 1.83E-04 | 2.18 |
| TC0X00009576.hg.1 | SYN1 | synapsin I | 2.07E-03 | 2.18 |
| TC2100007599.hg.1 | SAMSN1 | SAM domain, SH3 domain and nuclear localization signals 1 | 7.36E-04 | 2.17 |
| TC0100015586.hg.1 | PDZK1 | PDZ domain containing 1 | 6.45E-04 | 2.17 |
| TC1100012352.hg.1 | TMPRSS5 | transmembrane protease, serine 5 | 3.00E-03 | 2.17 |

| | | | | |
|---|---|---|---|---|
| TC1100012389.hg.1 | CADM1 | cell adhesion molecule 1 | 1.41E-04 | 2.17 |
| TC0200012071.hg.1 | UCN | urocortin | 1.71E-05 | 2.17 |
| HTA2-neg-47423694_st | NA | NA | 1.48E-03 | 2.17 |
| TC1200007108.hg.1 | KRAS | Memczak2013 ANTISENSE, coding, INTERNAL, intronic best transcript NM_033360 | 6.16E-05 | 2.16 |
| TC1400008615.hg.1 | NDRG2 | NDRG family member 2 | 2.52E-03 | 2.16 |
| TC1100006576.hg.1 | CD81 | CD81 molecule | 8.41E-05 | 2.16 |
| TC1500010712.hg.1 | GOLGA8N | golgin A8 family, member N | 1.07E-03 | 2.16 |
| TC0200012977.hg.1 | MXD1 | Memczak2013 ANTISENSE, coding, INTERNAL, intronic best transcript NM_001202514 | 4.22E-03 | 2.16 |
| TSUnmapped00000141.hg.1 | HMBS | hydroxymethylbilane synthase | 9.91E-04 | 2.15 |
| TC0900011501.hg.1 | NR6A1 | nuclear receptor subfamily 6, group A, member 1 | 7.18E-04 | 2.15 |
| TC1300010007.hg.1 | TMEM255B | transmembrane protein 255B | 4.25E-04 | 2.14 |
| TC0100013578.hg.1 | ADGRB2 | adhesion G protein-coupled receptor B2 | 1.01E-03 | 2.14 |
| TC2200007525.hg.1 | SERHL2 | serine hydrolase-like 2 | 2.06E-04 | 2.14 |
| HTA2-pos-3145790_st | NA | NA | 1.17E-03 | 2.14 |
| HTA2-pos-47422721_st | NA | NA | 1.02E-04 | 2.14 |
| TC0900008795.hg.1 | RALGPS1 | Ral GEF with PH domain and SH3 binding motif 1 | 4.71E-05 | 2.14 |
| TC0100010609.hg.1 | DNM3 | dynamin 3 | 1.99E-03 | 2.13 |
| TC1500007972.hg.1 | CRABP1 | cellular retinoic acid binding protein 1 | 1.07E-04 | 2.13 |
| TC1700006813.hg.1 | ODF4 | outer dense fiber of sperm tails 4 | 9.68E-03 | 2.13 |
| TC0100008052.hg.1 | CFAP57 | cilia and flagella associated protein 57 | 8.39E-04 | 2.13 |
| TC0600007375.hg.1 | HIST1H3H | histone cluster 1, H3h | 2.10E-03 | 2.12 |
| TC1500010769.hg.1 | GOLGA6C | golgin A6 family, member C | 1.56E-03 | 2.12 |
| TC1700010198.hg.1 | UNC119 | unc-119 lipid binding chaperone | 2.01E-05 | 2.12 |
| TC0700012296.hg.1 | C7orf60 | chromosome 7 open reading frame 60 | 1.05E-03 | 2.12 |
| TC0100015271.hg.1 | OVGP1 | oviductal glycoprotein 1 | 6.50E-04 | 2.12 |
| TC0400010519.hg.1 | APBB2 | amyloid beta (A4) precursor protein-binding, family B, member 2 | 2.36E-03 | 2.11 |
| TC0700010644.hg.1 | CRHR2 | corticotropin releasing hormone receptor 2 | 2.92E-04 | 2.11 |
| TC0200007418.hg.1 | SLC3A1 | solute carrier family 3 (amino acid transporter heavy chain), member 1 | 7.54E-04 | 2.11 |
| TC0600007792.hg.1 | PPARD | peroxisome proliferator-activated receptor delta | 1.33E-03 | 2.11 |
| TC0600014246.hg.1 | C6orf229 | chromosome 6 open reading frame 229 | 2.20E-03 | 2.11 |
| TC2000008242.hg.1 | RNF24 | ring finger protein 24 | 9.31E-04 | 2.10 |
| TC1500010770.hg.1 | GOLGA6D | golgin A6 family, member D | 9.51E-04 | 2.10 |
| TC0900011660.hg.1 | CCBL1 | cysteine conjugate-beta lyase, cytoplasmic | 5.60E-03 | 2.10 |
| TC2000007670.hg.1 | SNAI1 | snail family zinc finger 1 | 1.36E-04 | 2.10 |
| TC0500013088.hg.1 | MAPK9 | mitogen-activated protein kinase 9 | 8.50E-04 | 2.10 |
| TC1400009617.hg.1 | DPF3 | D4, zinc and double PHD fingers, family 3 | 3.20E-03 | 2.09 |
| HTA2-neg-47424023_st | NA | NA | 1.65E-04 | 2.09 |
| TC0100012089.hg.1 | GPR137B | G protein-coupled receptor 137B | 1.63E-03 | 2.09 |
| TC0600011125.hg.1 | HIST1H2AB | histone cluster 1, H2ab | 4.98E-03 | 2.09 |
| TC0400008331.hg.1 | NPNT | nephronectin | 3.91E-05 | 2.09 |
| TC0200007746.hg.1 | B3GNT2 | UDP-GlcNAc:betaGal beta-1,3-N-acetylglucosaminyltransferase 2 | 3.08E-04 | 2.08 |
| TC1000012525.hg.1 | ACBD7 | acyl-CoA binding domain containing 7 | 5.07E-03 | 2.08 |
| TC0800011861.hg.1 | LRRC6 | leucine rich repeat containing 6 | 2.56E-03 | 2.08 |
| TC0200011040.hg.1 | ITM2C | integral membrane protein 2C | 3.91E-04 | 2.08 |
| TC1200009547.hg.1 | SLC6A13 | solute carrier family 6 (neurotransmitter transporter), member 13 | 4.02E-05 | 2.08 |
| TC0900006559.hg.1 | CD274 | CD274 molecule | 5.66E-03 | 2.07 |
| TC1500008183.hg.1 | GOLGA6L3 | golgin A6 family-like 3 | 1.67E-03 | 2.07 |
| HTA2-neg-47422090_st | NA | NA | 3.20E-03 | 2.07 |
| TC1900010855.hg.1 | CADM4 | cell adhesion molecule 4 | 9.90E-05 | 2.07 |
| TC0500013260.hg.1 | MFAP3 | microfibrillar associated protein 3 | 1.31E-03 | 2.07 |
| TC0100014127.hg.1 | SPATA6 | spermatogenesis associated 6 | 6.65E-04 | 2.06 |
| TC0100018249.hg.1 | AMY2B | amylase, alpha 2B (pancreatic) | 1.28E-04 | 2.06 |
| TC0900012276.hg.1 | SCAI | suppressor of cancer cell invasion | 4.67E-05 | 2.06 |
| TC1900011814.hg.1 | ZNF17 | zinc finger protein 17 | 4.08E-05 | 2.06 |
| TC0200009829.hg.1 | GCA | grancalcin, EF-hand calcium binding protein | 4.39E-04 | 2.06 |
| TC1100008188.hg.1 | PPP6R3 | Memczak2013 ALT_ACCEPTOR, ALT_DONOR, coding, INTERNAL, intronic best transcript NM_001164162 | 5.65E-04 | 2.05 |
| TC2000008211.hg.1 | UBOX5 | U-box domain containing 5 | 1.73E-04 | 2.05 |
| TC0100018360.hg.1 | DISC1 | disrupted in schizophrenia 1 | 1.44E-04 | 2.05 |
| TC1900009756.hg.1 | SYCE2 | synaptonemal complex central element protein 2 | 9.40E-04 | 2.05 |
| 23076544 | NA | NA | 5.62E-05 | 2.04 |
| TC1100012382.hg.1 | NXPE4 | neurexophilin and PC-esterase domain family, member 4 | 1.26E-03 | 2.04 |
| TC0200007038.hg.1 | DRC1 | dynein regulatory complex subunit 1 | 3.53E-04 | 2.04 |
| TC1500010915.hg.1 | GOLGA6L5P | golgin A6 family-like 5, pseudogene | 1.15E-03 | 2.04 |
| TC1200009220.hg.1 | CCDC62 | coiled-coil domain containing 62 | 6.11E-05 | 2.03 |
| TC2000008268.hg.1 | SLC23A2 | solute carrier family 23 (ascorbic acid transporter), member 2 | 2.50E-03 | 2.03 |
| TC0500008619.hg.1 | IL3 | interleukin 3 | 6.71E-03 | 2.03 |
| AFFX-DapX-M_st | NA | NA | 7.90E-05 | 2.03 |
| TC0200011450.hg.1 | FAM150B | family with sequence similarity 150, member B | 2.77E-05 | 2.03 |
| TC1100011629.hg.1 | XRRA1 | X-ray radiation resistance associated 1 | 8.89E-04 | 2.03 |
| TC0100014910.hg.1 | TGFBR3 | transforming growth factor beta receptor III | 1.19E-03 | 2.03 |
| TC0800007529.hg.1 | SPIDR | scaffolding protein involved in DNA repair | 8.52E-04 | 2.03 |
| TC0100017018.hg.1 | ETNK2 | ethanolamine kinase 2 | 9.90E-03 | 2.02 |
| HTA2-neg-47424705_st | NA | NA | 5.11E-04 | 2.02 |
| TC1600006893.hg.1 | CLEC16A | C-type lectin domain family 16, member A | 4.47E-05 | 2.02 |
| TC0700013323.hg.1 | SUN1 | Sad1 and UNC84 domain containing 1 | 1.86E-03 | 2.02 |
| TC1600010000.hg.1 | STX1B | syntaxin 1B | 1.19E-03 | 2.01 |
| TC0500007231.hg.1 | PTGER4 | prostaglandin E receptor 4 (subtype EP4) | 5.20E-05 | 2.01 |
| TC1300008253.hg.1 | CRYL1 | crystallin lambda 1 | 1.34E-03 | 2.01 |
| TC0100018261.hg.1 | GSTM4 | glutathione S-transferase mu 4 | 5.69E-04 | 2.01 |
| AFFX-r2-Bs-dap-3_st | NA | NA | 5.12E-05 | 2.01 |
| TC1900008287.hg.1 | PVRL2 | poliovirus receptor-related 2 (herpesvirus entry mediator B) | 4.73E-04 | 2.00 |
| TC1200007594.hg.1 | ASIC1 | acid sensing ion channel 1 | 5.28E-04 | 2.00 |
| TC0100018262.hg.1 | GSTM2 | glutathione S-transferase mu 2 (muscle) | 1.37E-03 | 2.00 |
| AFFX-r2-Bs-phe-3_st | NA | NA | 6.06E-04 | 2.00 |
| TC0700008063.hg.1 | SPDYE5 | speedy/RINGO cell cycle regulator family member E5 | 8.93E-04 | 2.00 |
| TC1200011993.hg.1 | SLC8B1 | solute carrier family 8 (sodium/lithium/calcium exchanger), member B1 | 1.18E-03 | 1.99 |
| TC0200014620.hg.1 | CACNB4 | calcium channel, voltage-dependent, beta 4 subunit | 1.80E-03 | 1.99 |
| TC1900009584.hg.1 | ZNF560 | zinc finger protein 560 | 1.01E-03 | 1.98 |
| TC0700013604.hg.1 | UPK3BL | uroplakin 3B-like | 5.35E-05 | 1.98 |
| TC1900010005.hg.1 | RAB3A | RAB3A, member RAS oncogene family | 3.74E-04 | 1.98 |
| TC0100013205.hg.1 | ECE1 | endothelin converting enzyme 1 | 3.24E-04 | 1.98 |
| TC0800012447.hg.1 | AZIN1 | antizyme inhibitor 1 | 4.92E-04 | 1.98 |
| TC2200006827.hg.1 | GNAZ | guanine nucleotide binding protein (G protein), alpha z polypeptide | 5.38E-03 | 1.97 |
| 23075476 | NA | NA | 6.01E-04 | 1.97 |
| TC0200014361.hg.1 | LYPD1 | LY6/PLAUR domain containing 1 | 8.58E-03 | 1.97 |
| HTA2-pos-PSR01002164.hg.1 | NA | NA | 8.58E-04 | 1.97 |
| TC0100018210.hg.1 | SH3D21 | SH3 domain containing 21 | 4.44E-04 | 1.97 |
| TC0800010926.hg.1 | PAG1 | phosphoprotein membrane anchor with glycosphingolipid microdomains 1 | 5.04E-04 | 1.97 |
| TC0800012405.hg.1 | FUT10 | fucosyltransferase 10 (alpha (1,3) fucosyltransferase) | 7.53E-04 | 1.96 |
| TC0800009853.hg.1 | BIN3 | bridging integrator 3 | 1.19E-03 | 1.96 |
| TC0X00010643.hg.1 | 44080 | septin 6 | 8.32E-04 | 1.96 |
| 23075475 | NA | NA | 7.95E-04 | 1.96 |
| TC0300010821.hg.1 | ULK4 | unc-51 like kinase 4 | 1.81E-03 | 1.95 |
| TC1200012660.hg.1 | SLC35E3 | solute carrier family 35, member E3 | 7.54E-03 | 1.95 |
| TC0700009036.hg.1 | LEP | leptin | 6.76E-03 | 1.95 |
| AFFX-DapX-5_st | NA | NA | 1.71E-04 | 1.95 |
| TC0600013165.hg.1 | EPB41L2 | erythrocyte membrane protein band 4.1-like 2 | 1.64E-04 | 1.95 |
| TC0200011504.hg.1 | PXDN | Transcript Identified by AceView, Entrez Gene ID(s) 7837 | 7.39E-03 | 1.95 |
| TC2000009250.hg.1 | PLTP | phospholipid transfer protein | 1.11E-04 | 1.95 |
| TC1500010128.hg.1 | TBC1D2B | TBC1 domain family, member 2B | 1.59E-03 | 1.94 |
| TC0X00011275.hg.1 | TCEANC | transcription elongation factor A (SII) N-terminal and central domain containing | 1.18E-03 | 1.94 |
| TC1500008086.hg.1 | SAXO2 | stabilizer of axonemal microtubules 2 | 1.02E-04 | 1.94 |
| TC2000006791.hg.1 | PCSK2 | proprotein convertase subtilisin/kexin type 2 | 1.80E-03 | 1.94 |
| TC0100015401.hg.1 | IGSF3 | immunoglobulin superfamily, member 3 | 1.13E-03 | 1.93 |
| TC1900007096.hg.1 | JUNB | jun B proto-oncogene | 1.20E-04 | 1.93 |
| TC1700007918.hg.1 | AOC2 | amine oxidase, copper containing 2 (retina-specific) | 1.03E-03 | 1.93 |
| TC2200008196.hg.1 | MIF-AS1 | MIF antisense RNA 1 | 2.11E-03 | 1.93 |
| TC0500011334.hg.1 | TMEM167A | Transcript Identified by AceView, Entrez Gene ID(s) 153339 | 9.13E-03 | 1.93 |
| TC2200006643.hg.1 | ZDHHC8 | zinc finger, DHHC-type containing 8 | 1.76E-04 | 1.93 |

| | | | | |
|---|---|---|---|---|
| TC0700011477.hg.1 | TRIM74 | tripartite motif containing 74 | 4.45E-04 | 1.93 |
| TC1100010718.hg.1 | LRP4 | LDL receptor related protein 4 | 2.43E-03 | 1.93 |
| TC1400007145.hg.1 | FRMD6 | FERM domain containing 6 | 4.80E-03 | 1.92 |
| TC0800009856.hg.1 | EGR3 | early growth response 3 | 4.97E-03 | 1.92 |
| TC1100009302.hg.1 | TECTA | tectorin alpha | 5.23E-03 | 1.92 |
| TC2100007967.hg.1 | PAXBP1 | PAX3 and PAX7 binding protein 1 | 4.50E-05 | 1.92 |
| TSUnmapped00000085.hg.1 | CCDC84 | coiled-coil domain containing 84 | 6.48E-04 | 1.92 |
| HTA2-pos-2985880_st | NA | NA | 7.98E-03 | 1.92 |
| TC0100014502.hg.1 | WDR78 | WD repeat domain 78 | 9.66E-05 | 1.92 |
| TC1100011652.hg.1 | GDPD5 | glycerophosphodiester phosphodiesterase domain containing 5 | 6.42E-04 | 1.92 |
| TC0900010580.hg.1 | GKAP1 | G kinase anchoring protein 1 | 1.51E-03 | 1.92 |
| TC1900009947.hg.1 | MYO9B | Memczak2013 ANTISENSE, CDS, coding, INTERNAL best transcript NM_004145 | 4.21E-04 | 1.92 |
| TC1300006919.hg.1 | FREM2 | FRAS1 related extracellular matrix protein 2 | 7.09E-05 | 1.92 |
| TC2000010008.hg.1 | NFS1 | NFS1 cysteine desulfurase | 6.03E-05 | 1.92 |
| TC1900007888.hg.1 | APLP1 | amyloid beta (A4) precursor-like protein 1 | 8.23E-04 | 1.92 |
| HTA2-neg-47423221_st | NA | NA | 4.11E-03 | 1.92 |
| TC1900011734.hg.1 | MIA | melanoma inhibitory activity | 6.34E-05 | 1.92 |
| TC0300012482.hg.1 | RAB6B | RAB6B, member RAS oncogene family | 4.35E-03 | 1.91 |
| TC0100010526.hg.1 | TBX19 | T-box 19 | 3.14E-04 | 1.91 |
| TC1900009867.hg.1 | AKAP8L | A kinase (PRKA) anchor protein 8-like | 8.93E-05 | 1.91 |
| TC0100010281.hg.1 | ATP1A2 | ATPase, Na+/K+ transporting, alpha 2 polypeptide | 3.58E-03 | 1.91 |
| 23071851 | NA | NA | 9.36E-04 | 1.91 |
| 23071994 | NA | NA | 9.36E-04 | 1.91 |
| TC1500010780.hg.1 | GOLGA6L9 | golgin A6 family-like 9 | 1.62E-03 | 1.91 |
| TC1900009670.hg.1 | DOCK6 | dedicator of cytokinesis 6 | 6.12E-04 | 1.90 |
| TC1100010472.hg.1 | CCDC73 | coiled-coil domain containing 73 | 5.13E-03 | 1.90 |
| TC1000010982.hg.1 | ASCC1 | Transcript Identified by AceView, Entrez Gene ID(s) 51008 | 1.55E-03 | 1.90 |
| AFFX-DapX-3_st | NA | NA | 1.21E-04 | 1.90 |
| TC0100007876.hg.1 | RRAGC | Memczak2013 ANTISENSE, coding, INTERNAL, intronic best transcript NM_022157 | 3.90E-04 | 1.90 |
| TC0200011107.hg.1 | EFHD1 | EF-hand domain family member D1 | 1.68E-03 | 1.90 |
| TC1900007155.hg.1 | MIR1199 | microRNA 1199 | 9.07E-05 | 1.90 |
| TSUnmapped00000176.hg.1 | SAG | S-antigen; retina and pineal gland (arrestin) | 5.98E-03 | 1.90 |
| TC1700006768.hg.1 | EFNB3 | ephrin-B3 | 8.22E-03 | 1.90 |
| TC0100017241.hg.1 | TMEM206 | transmembrane protein 206 | 4.03E-03 | 1.90 |
| TC0900007547.hg.1 | C9orf85 | chromosome 9 open reading frame 85 | 3.72E-03 | 1.90 |
| TC1700007262.hg.1 | MAP2K3 | mitogen-activated protein kinase kinase 3 | 9.12E-04 | 1.90 |
| TC1600010987.hg.1 | CMIP | Memczak2013 ANTISENSE, coding, INTERNAL, intronic best transcript NM_198390 | 6.69E-04 | 1.90 |
| TC1600009192.hg.1 | MTRNR2L4 | MT-RNR2-like 4 | 1.07E-04 | 1.90 |
| TC0X00006799.hg.1 | SAT1 | spermidine/spermine N1-acetyltransferase 1 | 7.70E-04 | 1.90 |
| TC0600011227.hg.1 | HIST1H4K | histone cluster 1, H4k | 1.01E-03 | 1.89 |
| TC0100016121.hg.1 | KCNJ10 | potassium channel, inwardly rectifying subfamily J, member 10 | 1.62E-04 | 1.89 |
| TC0100017445.hg.1 | TLR5 | toll-like receptor 5 | 2.44E-03 | 1.89 |
| TC1000007461.hg.1 | RASSF4 | Ras association (RalGDS/AF-6) domain family member 4 | 3.43E-03 | 1.89 |
| HTA2-pos-2985881_st | NA | NA | 8.61E-04 | 1.89 |
| TC0100017716.hg.1 | FAM89A | family with sequence similarity 89, member A | 5.98E-04 | 1.88 |
| TC1900011874.hg.1 | ZNF878 | zinc finger protein 878 | 5.15E-03 | 1.88 |
| TC0200016571.hg.1 | OSBPL6 | oxysterol binding protein-like 6 | 2.36E-03 | 1.88 |
| TC0100017310.hg.1 | ESRRG | estrogen-related receptor gamma | 8.65E-04 | 1.88 |
| TC0700013602.hg.1 | POLR2J3 | polymerase (RNA) II (DNA directed) polypeptide J3 | 5.88E-05 | 1.88 |
| AFFX-r2-Bs-dap-M_st | NA | NA | 6.80E-05 | 1.88 |
| TC1700010560.hg.1 | PLXDC1 | plexin domain containing 1 | 9.40E-04 | 1.88 |
| TC1100010274.hg.1 | SPTY2D1 | SPT2 chromatin protein domain containing 1 | 4.24E-04 | 1.88 |
| TC1700007204.hg.1 | SPECC1 | sperm antigen with calponin homology and coiled-coil domains 1 | 5.08E-03 | 1.88 |
| HTA2-neg-47419415_st | NA | NA | 3.01E-04 | 1.88 |
| TC1700008769.hg.1 | KCNJ2 | potassium channel, inwardly rectifying subfamily J, member 2 | 2.89E-04 | 1.88 |
| TC0800009619.hg.1 | CTSB | cathepsin B | 8.67E-04 | 1.88 |
| TC0300008086.hg.1 | ARL6 | ADP-ribosylation factor like GTPase 6 | 9.26E-03 | 1.88 |
| TC1700008016.hg.1 | HIGD1B | HIG1 hypoxia inducible domain family, member 1B | 6.79E-03 | 1.88 |
| TC0X00011279.hg.1 | TSPAN7 | tetraspanin 7 | 2.35E-03 | 1.88 |
| TC1900007127.hg.1 | IER2 | immediate early response 2 | 2.58E-04 | 1.88 |
| 23067838 | NA | NA | 8.96E-04 | 1.87 |
| TC1400008058.hg.1 | PPP4R4 | protein phosphatase 4, regulatory subunit 4 | 8.96E-04 | 1.87 |
| TC1600011442.hg.1 | MAP1LC3B | microtubule-associated protein 1 light chain 3 beta | 1.55E-03 | 1.87 |
| HTA2-neg-47422232_st | NA | NA | 3.21E-03 | 1.87 |
| TC1900009158.hg.1 | RPS15 | Memczak2013 ANTISENSE, coding, INTERNAL, intronic best transcript NM_001018 | 8.78E-04 | 1.87 |
| TC1600010801.hg.1 | PKD1L3 | polycystic kidney disease 1-like 3 | 1.09E-03 | 1.87 |
| TC0100006861.hg.1 | FBXO44 | F-box protein 44 | 1.47E-04 | 1.87 |
| TC0200011419.hg.1 | ATG4B | autophagy related 4B, cysteine peptidase | 1.88E-03 | 1.87 |
| TC0600007613.hg.1 | HSPA1A | heat shock 70kDa protein 1A | 9.31E-05 | 1.87 |
| TC1600007448.hg.1 | CORO1A | coronin, actin binding protein, 1A | 5.30E-03 | 1.87 |
| TC1000012433.hg.1 | SPAG6 | sperm associated antigen 6 | 2.58E-03 | 1.86 |
| TC0600008255.hg.1 | EFHC1 | EF-hand domain (C-terminal) containing 1 | 2.11E-03 | 1.86 |
| TC1400010754.hg.1 | C14orf37 | chromosome 14 open reading frame 37 | 3.31E-04 | 1.86 |
| TC0700013421.hg.1 | ZNF789 | zinc finger protein 789 | 5.26E-04 | 1.86 |
| HTA2-pos-PSR0X007730.hg.1 | NA | NA | 9.11E-04 | 1.86 |
| TC1800008677.hg.1 | CFAP53 | cilia and flagella associated protein 53 | 5.31E-03 | 1.86 |
| TC0200015424.hg.1 | ALS2CR12 | amyotrophic lateral sclerosis 2 chromosome region candidate 12 | 4.16E-03 | 1.86 |
| TC1900011879.hg.1 | ZNF44 | zinc finger protein 44 | 6.64E-03 | 1.86 |
| TC0200016464.hg.1 | APLF | aprataxin and PNKP like factor | 2.29E-04 | 1.86 |
| TC1200007653.hg.1 | NR4A1 | nuclear receptor subfamily 4, group A, member 1 | 6.86E-03 | 1.86 |
| TC0900011669.hg.1 | SH3GLB2 | SH3-domain GRB2-like endophilin B2 | 2.40E-03 | 1.86 |
| TC1000011567.hg.1 | CRTAC1 | cartilage acidic protein 1 | 1.78E-03 | 1.85 |
| 23076588 | NA | NA | 1.17E-03 | 1.85 |
| TC0800009212.hg.1 | MAPK15 | mitogen-activated protein kinase 15 | 1.17E-03 | 1.85 |
| HTA2-neg-47422095_st | NA | NA | 1.45E-03 | 1.85 |
| TC0X00011236.hg.1 | RAB39B | Transcript Identified by AceView, Entrez Gene ID(s) 116442 | 4.09E-03 | 1.85 |
| TC1900011816.hg.1 | ZNF419 | zinc finger protein 419 | 8.08E-05 | 1.85 |
| HTA2-pos-JUC01000985.hg.1 | NA | NA | 8.13E-05 | 1.85 |
| HTA2-pos-JUC13008384.hg.1 | NA | NA | 8.13E-05 | 1.85 |
| HTA2-pos-JUC19003067.hg.1 | NA | NA | 8.13E-05 | 1.85 |
| TSUnmapped00000608.hg.1 | ZNF35 | zinc finger protein 35 | 1.07E-03 | 1.85 |
| TC2100007952.hg.1 | URB1 | URB1 ribosome biogenesis 1 homolog (S. cerevisiae) | 1.57E-03 | 1.85 |
| TC0600014371.hg.1 | RNASET2 | ribonuclease T2 | 7.99E-03 | 1.85 |
| TC0300011264.hg.1 | IL17RD | interleukin 17 receptor D | 1.83E-03 | 1.85 |
| TC1100010041.hg.1 | ST5 | Transcript Identified by AceView, Entrez Gene ID(s) 6764 | 9.67E-03 | 1.85 |
| TC0200015776.hg.1 | RESP18 | regulated endocrine-specific protein 18 | 1.69E-04 | 1.84 |
| TC1600009351.hg.1 | GRIN2A | Transcript Identified by AceView, Entrez Gene ID(s) 2903 | 7.26E-05 | 1.84 |
| TC0200006703.hg.1 | ATP6V1C2 | ATPase, H+ transporting, lysosomal 42kDa, V1 subunit C2 | 2.98E-03 | 1.84 |
| TC0800011597.hg.1 | EXT1 | Jeck2013 ALT_ACCEPTOR, ALT_DONOR, coding, INTERNAL, intronic best transcript NM_000127 | 1.81E-03 | 1.84 |
| TC1700008328.hg.1 | STXBP4 | syntaxin binding protein 4 | 5.78E-04 | 1.84 |
| TC0600011821.hg.1 | GUCA1B | guanylate cyclase activator 1B (retina) | 9.59E-04 | 1.84 |
| TC1600007240.hg.1 | PRKCB | protein kinase C, beta | 1.24E-03 | 1.84 |
| TC2100008534.hg.1 | PCBP3 | poly(rC) binding protein 3 | 8.57E-04 | 1.84 |
| TC1500009200.hg.1 | TTBK2 | tau tubulin kinase 2 | 1.93E-03 | 1.84 |
| TC0300009696.hg.1 | VWA5B2 | von Willebrand factor A domain containing 5B2 | 6.36E-03 | 1.84 |
| TC0200014728.hg.1 | LY75-CD302 | LY75-CD302 readthrough | 4.65E-04 | 1.84 |
| TC0100018464.hg.1 | PPM1J | protein phosphatase, Mg2+/Mn2+ dependent, 1J | 1.70E-03 | 1.84 |
| TC0800012351.hg.1 | WDYHV1 | WDYHV motif containing 1 | 1.88E-03 | 1.84 |
| TC0100015701.hg.1 | HIST2H3A | histone cluster 2, H3a | 4.97E-04 | 1.84 |
| TC0800009194.hg.1 | RHPN1 | rhophilin, Rho GTPase binding protein 1 | 2.83E-03 | 1.84 |
| TC0200010745.hg.1 | IGFBP2 | insulin like growth factor binding protein 2 | 1.27E-03 | 1.84 |
| TC1300006690.hg.1 | POLR1D | polymerase (RNA) I polypeptide D | 6.14E-05 | 1.83 |
| TC2200007903.hg.1 | LOC101929372 | uncharacterized LOC101929372 | 6.86E-04 | 1.83 |
| TC0200008536.hg.1 | ANKRD36 | Transcript Identified by AceView, Entrez Gene ID(s) 375248 | 6.70E-04 | 1.83 |
| TC1900008259.hg.1 | ZNF233 | zinc finger protein 233 | 4.95E-04 | 1.83 |
| TC0100008773.hg.1 | MSH4 | mutS homolog 4 | 3.81E-03 | 1.83 |
| TC0900007163.hg.1 | FRMPD1 | FERM and PDZ domain containing 1 | 9.84E-04 | 1.83 |
| TC1100012519.hg.1 | CBL | Memczak2013 ANTISENSE, coding, INTERNAL, UTR3 best transcript NM_005188 | 4.98E-03 | 1.83 |
| TC1100011101.hg.1 | NXF1 | nuclear RNA export factor 1 | 3.24E-03 | 1.83 |

| | | | | |
|---|---|---|---|---|
| TC1600009620.hg.1 | SMG1 | SMG1 phosphatidylinositol 3-kinase-related kinase | 5.65E-04 | 1.83 |
| HTA2-pos-JUC09010881.hg.1 | NA | NA | 4.98E-03 | 1.82 |
| TC0800011334.hg.1 | KLF10 | Kruppel-like factor 10 | 2.37E-04 | 1.82 |
| TC1800009203.hg.1 | PQLC1 | PQ loop repeat containing 1 | 2.22E-03 | 1.82 |
| TC1900011653.hg.1 | MCOLN1 | mucolipin 1 | 8.82E-04 | 1.82 |
| TC0800010234.hg.1 | SFRP1 | secreted frizzled-related protein 1 | 9.22E-05 | 1.82 |
| TC1900009269.hg.1 | TLE2 | transducin-like enhancer of split 2 | 7.46E-03 | 1.82 |
| TC1900008860.hg.1 | LENG8 | leukocyte receptor cluster (LRC) member 8 | 1.44E-03 | 1.82 |
| 23076569 | NA | NA | 2.70E-04 | 1.82 |
| TC1500010841.hg.1 | CHRFAM7A | CHRNA7 (cholinergic receptor, nicotinic, alpha 7, exons 5-10) and FAM7A (family with sequence similarity 7A, exons A-E) fusion | 3.19E-03 | 1.81 |
| TC0100009746.hg.1 | NUDT17 | nudix hydrolase 17 | 2.11E-03 | 1.81 |
| 23069220 | NA | NA | 2.42E-04 | 1.81 |
| TSUnmapped00000145.hg.1 | SAG | S-antigen; retina and pineal gland (arrestin) | 1.62E-04 | 1.81 |
| TC0700013388.hg.1 | STAG3L1 | stromal antigen 3-like 1 (pseudogene) | 2.66E-04 | 1.81 |
| TC1500010709.hg.1 | ARHGAP11B | Rho GTPase activating protein 11B | 5.45E-03 | 1.81 |
| TC0500011752.hg.1 | FEM1C | Jeck2013 ALT_ACCEPTOR, ALT_DONOR, coding, INTERNAL, intronic best transcript NM_020177 | 8.95E-03 | 1.81 |
| TC0X00007116.hg.1 | RP2 | retinitis pigmentosa 2 (X-linked recessive) | 1.62E-03 | 1.81 |
| TC1100010044.hg.1 | ST5 | Transcript Identified by AceView, Entrez Gene ID(s) 6764 | 2.21E-03 | 1.81 |
| TC0100016649.hg.1 | NMNAT2 | nicotinamide nucleotide adenylyltransferase 2 | 9.09E-03 | 1.81 |
| TC1800007218.hg.1 | C18orf25 | chromosome 18 open reading frame 25 | 2.41E-03 | 1.80 |
| TC0700010162.hg.1 | FBXL18 | F-box and leucine-rich repeat protein 18 | 2.80E-04 | 1.80 |
| 23072532 | NA | NA | 1.41E-03 | 1.80 |
| TC1600007189.hg.1 | VWA3A | von Willebrand factor A domain containing 3A | 7.04E-04 | 1.80 |
| TC0700012163.hg.1 | EFCAB10 | EF-hand calcium binding domain 10 | 4.82E-04 | 1.80 |
| TC0800011713.hg.1 | MTSS1 | metastasis suppressor 1 | 5.09E-03 | 1.80 |
| TC0300011834.hg.1 | FILIP1L | filamin A interacting protein 1-like | 9.33E-05 | 1.80 |
| TC0700013401.hg.1 | AC007566.10 | --- | 5.19E-03 | 1.80 |
| TC1500007805.hg.1 | BBS4 | Memczak2013 ALT_ACCEPTOR, ALT_DONOR, coding, INTERNAL, intronic best transcript NM_033028 | 5.19E-03 | 1.80 |
| TC0200016734.hg.1 | MAP3K19 | mitogen-activated protein kinase kinase kinase 19 | 2.53E-04 | 1.80 |
| TC0X00009117.hg.1 | FANCB | Fanconi anemia complementation group B | 1.23E-03 | 1.80 |
| TC2200007909.hg.1 | MICAL3 | microtubule associated monooxygenase, calponin and LIM domain containing 3 | 1.37E-04 | 1.80 |
| TC1000011040.hg.1 | CAMK2G | calcium/calmodulin-dependent protein kinase II gamma | 3.94E-03 | 1.80 |
| TC0900010737.hg.1 | DIRAS2 | DIRAS family, GTP-binding RAS-like 2 | 2.30E-03 | 1.79 |
| TC0100009274.hg.1 | AMY1C | amylase, alpha 1C (salivary) | 6.44E-04 | 1.79 |
| TC1300006658.hg.1 | WASF3 | WAS protein family, member 3 | 4.35E-03 | 1.79 |
| TC1900011765.hg.1 | PPP1R37 | protein phosphatase 1, regulatory subunit 37 | 2.45E-04 | 1.79 |
| TC1500008135.hg.1 | UBE2Q2L | ubiquitin conjugating enzyme E2Q family member 2-like | 5.37E-04 | 1.79 |
| TC1200009076.hg.1 | SUDS3 | SDS3 homolog, SIN3A corepressor complex component | 2.34E-03 | 1.79 |
| TC0200014717.hg.1 | WDSUB1 | WD repeat, sterile alpha motif and U-box domain containing 1 | 3.97E-03 | 1.79 |
| TC0200008263.hg.1 | USP39 | ubiquitin specific peptidase 39 | 2.81E-04 | 1.79 |
| TC0100015180.hg.1 | WDR47 | WD repeat domain 47 | 8.51E-03 | 1.79 |
| TC2000008587.hg.1 | RALGAPA2 | Ral GTPase activating protein, alpha subunit 2 (catalytic) | 2.12E-04 | 1.79 |
| AFFX-r2-Bs-phe-M_st | NA | NA | 2.98E-04 | 1.79 |
| TC1700011795.hg.1 | PRPSAP1 | phosphoribosyl pyrophosphate synthetase-associated protein 1 | 1.66E-04 | 1.78 |
| TSUnmapped00000149.hg.1 | ZNF35 | zinc finger protein 35 | 1.36E-03 | 1.78 |
| TC0700013586.hg.1 | C7orf76 | chromosome 7 open reading frame 76 | 9.00E-04 | 1.78 |
| TC2000009966.hg.1 | FKBP1A-SDCBP2 | FKBP1A-SDCBP2 readthrough (NMD candidate) | 2.71E-04 | 1.78 |
| TC0100013763.hg.1 | POU3F1 | POU class 3 homeobox 1 | 2.95E-03 | 1.78 |
| TC1100013020.hg.1 | PPP1R32 | protein phosphatase 1, regulatory subunit 32 | 7.72E-03 | 1.78 |
| TC0200012412.hg.1 | C1GALT1C1L | C1GALT1-specific chaperone 1 like | 1.19E-03 | 1.78 |
| TC0400008384.hg.1 | PLA2G12A | Memczak2013 ANTISENSE, CDS, coding, INTERNAL best transcript NM_030821 | 5.40E-03 | 1.78 |
| TC1500010845.hg.1 | OTUD7A | OTU deubiquitinase 7A | 9.60E-04 | 1.78 |
| TC0700012080.hg.1 | SPDYE6 | speedy/RINGO cell cycle regulator family member E6 | 9.98E-04 | 1.78 |
| TC1300006770.hg.1 | ALOX5AP | arachidonate 5-lipoxygenase-activating protein | 3.81E-04 | 1.78 |
| TC0800010450.hg.1 | ATP6V1H | ATPase, H+ transporting, lysosomal 50/57kDa, V1 subunit H | 5.08E-03 | 1.78 |
| TC0300013763.hg.1 | RUBCN | RUN domain and cysteine-rich domain containing, Beclin 1-interacting protein | 3.54E-03 | 1.78 |
| HTA2-neg-47420204_st | NA | NA | 4.40E-03 | 1.78 |
| TC0700013457.hg.1 | FSCN3 | fascin actin-bundling protein 3, testicular | 8.69E-04 | 1.78 |
| TC1100012712.hg.1 | ACRV1 | acrosomal vesicle protein 1 | 1.96E-03 | 1.78 |
| TC0500009824.hg.1 | ZDHHC11 | Transcript Identified by AceView, Entrez Gene ID(s) 79844 | 4.52E-03 | 1.78 |
| TC1900011819.hg.1 | ZNF211 | zinc finger protein 211 | 7.66E-05 | 1.78 |
| TC0400012882.hg.1 | ZNF732 | zinc finger protein 732 | 5.06E-04 | 1.78 |
| HTA2-pos-2960237_st | NA | NA | 8.76E-03 | 1.78 |
| TC1500008139.hg.1 | GOLGA6L4 | Homo sapiens golgin A6 family-like 4 (GOLGA6L4), mRNA. | 1.73E-03 | 1.77 |
| TC1600011047.hg.1 | HSDL1 | hydroxysteroid dehydrogenase like 1 | 5.01E-04 | 1.77 |
| HTA2-pos-47422722_st | NA | NA | 1.95E-04 | 1.77 |
| HTA2-pos-PSR02023970.hg.1 | NA | NA | 9.13E-04 | 1.77 |
| TC1200012700.hg.1 | C12orf75 | chromosome 12 open reading frame 75 | 8.51E-03 | 1.77 |
| TC0900008865.hg.1 | CERCAM | cerebral endothelial cell adhesion molecule | 1.93E-03 | 1.77 |
| TSUnmapped00000262.hg.1 | MLXIP | MLX interacting protein | 4.01E-04 | 1.77 |
| TC0700008639.hg.1 | SPDYE2 | speedy/RINGO cell cycle regulator family member E2 | 4.03E-04 | 1.77 |
| TC0700008647.hg.1 | SPDYE2B | speedy/RINGO cell cycle regulator family member E2B | 4.03E-04 | 1.77 |
| TC0700008000.hg.1 | LAT2 | linker for activation of T-cells family member 2 | 8.10E-04 | 1.77 |
| TC1500010291.hg.1 | NMB | neuromedin B | 2.99E-03 | 1.77 |
| TC0300013687.hg.1 | TFRC | Transcript Identified by AceView, Entrez Gene ID(s) 7037 | 4.98E-03 | 1.77 |
| 23076565 | NA | NA | 2.60E-03 | 1.76 |
| AFFX-PheX-3_st | NA | NA | 2.02E-04 | 1.76 |
| 23075412 | NA | NA | 9.95E-03 | 1.76 |
| TC0300009974.hg.1 | MUC20 | mucin 20, cell surface associated | 9.95E-03 | 1.76 |
| TC0700013119.hg.1 | XRCC2 | X-ray repair complementing defective repair in Chinese hamster cells 2 | 2.26E-03 | 1.76 |
| TC0400011180.hg.1 | SCD5 | stearoyl-CoA desaturase 5 | 9.76E-04 | 1.76 |
| TC2000009014.hg.1 | NDRG3 | NDRG family member 3 | 2.49E-04 | 1.76 |
| TC1400008583.hg.1 | CCNB1IP1 | cyclin B1 interacting protein 1, E3 ubiquitin protein ligase | 2.47E-03 | 1.76 |
| TC0700007364.hg.1 | SPDYE1 | speedy/RINGO cell cycle regulator family member E1 | 1.15E-03 | 1.76 |
| TC0700012998.hg.1 | ZNF767P | zinc finger family member 767, pseudogene | 8.42E-04 | 1.76 |
| TC1200010108.hg.1 | GYS2 | glycogen synthase 2 (liver) | 7.28E-03 | 1.76 |
| TSUnmapped00000268.hg.1 | CCDC84 | coiled-coil domain containing 84 | 5.70E-04 | 1.76 |
| TC0100011325.hg.1 | TMCC2 | transmembrane and coiled-coil domain family 2 | 6.30E-03 | 1.76 |
| TC1000009152.hg.1 | HTRA1 | HtrA serine peptidase 1 | 2.11E-03 | 1.76 |
| TC0100010284.hg.1 | PEA15 | phosphoprotein enriched in astrocytes 15 | 9.02E-05 | 1.76 |
| TC0800009326.hg.1 | TDRP | testis development related protein | 6.58E-03 | 1.76 |
| TC1100013053.hg.1 | RAD9A | RAD9 checkpoint clamp component A | 2.44E-04 | 1.76 |
| 23068514 | NA | NA | 7.12E-03 | 1.76 |
| TC0800007311.hg.1 | ADGRA2 | adhesion G protein-coupled receptor A2 | 2.86E-03 | 1.75 |
| TC0700013575.hg.1 | STAG3L3 | stromal antigen 3-like 3 (pseudogene) | 4.20E-04 | 1.75 |
| TC1500010909.hg.1 | STARD5 | StAR-related lipid transfer domain containing 5 | 3.70E-03 | 1.75 |
| TC0200015434.hg.1 | ALS2 | ALS2, alsin Rho guanine nucleotide exchange factor | 1.02E-03 | 1.75 |
| 23075635 | NA | NA | 3.33E-04 | 1.75 |
| TC1400010632.hg.1 | GPATCH2L | G-patch domain containing 2 like | 1.21E-04 | 1.75 |
| TC1700011919.hg.1 | CEP295NL | CEP295 N-terminal like | 1.76E-03 | 1.75 |
| TC0400011053.hg.1 | CXCL10 | chemokine (C-X-C motif) ligand 10 | 4.62E-03 | 1.75 |
| TC0100007184.hg.1 | OTUD3 | OTU deubiquitinase 3 | 3.81E-04 | 1.75 |
| TC0100013349.hg.1 | RSRP1 | arginine/serine-rich protein 1 | 3.00E-03 | 1.75 |
| 23068498 | NA | NA | 3.13E-03 | 1.75 |
| TC0200008734.hg.1 | C2orf49 | chromosome 2 open reading frame 49 | 5.91E-04 | 1.75 |
| TC0200015514.hg.1 | GPR1 | G protein-coupled receptor 1 | 1.46E-03 | 1.74 |
| TC1700011815.hg.1 | MXRA7 | matrix-remodelling associated 7 | 6.89E-04 | 1.74 |
| TC0100015698.hg.1 | HIST2H3D | histone cluster 2, H3d | 1.10E-03 | 1.74 |
| TC1400009710.hg.1 | ACYP1 | acylphosphatase 1, erythrocyte (common) type | 5.30E-03 | 1.74 |
| TC1100012994.hg.1 | CRY2 | cryptochrome circadian clock 2 | 1.15E-03 | 1.74 |
| TC0300009310.hg.1 | MLF1 | myeloid leukemia factor 1 | 1.57E-03 | 1.74 |
| TC1500010906.hg.1 | RP11-351M8.2 | --- | 2.17E-03 | 1.74 |
| TC0700012086.hg.1 | POLR2J | polymerase (RNA) II (DNA directed) polypeptide J, 13.3kDa | 1.73E-04 | 1.74 |
| TC0900008684.hg.1 | GPR21 | G protein-coupled receptor 21 | 2.87E-04 | 1.74 |
| TC1200006905.hg.1 | FAM234B | family with sequence similarity 234, member B | 1.17E-03 | 1.74 |
| HTA2-neg-47420908_st | NA | NA | 5.28E-03 | 1.74 |
| TC0100009877.hg.1 | HIST2H3A | histone cluster 2, H3a | 1.99E-03 | 1.74 |
| HTA2-neg-47420639_st | NA | NA | 1.52E-03 | 1.74 |
| TC0700011519.hg.1 | STAG3L2 | stromal antigen 3-like 2 (pseudogene) | 7.78E-04 | 1.74 |

| | | | | |
|---|---|---|---|---|
| TC2200008346.hg.1 | MN1 | meningioma (disrupted in balanced translocation) 1 | 1.03E-04 | 1.74 |
| TC0100018248.hg.1 | RNPC3 | RNA binding region (RNP1, RRM) containing 3 | 5.06E-04 | 1.74 |
| TC2000007083.hg.1 | ID1 | inhibitor of DNA binding 1, dominant negative helix-loop-helix protein | 7.09E-03 | 1.73 |
| TC0300007610.hg.1 | PXK | PX domain containing serine/threonine kinase | 1.77E-04 | 1.73 |
| TC0600010331.hg.1 | FAM120B | family with sequence similarity 120B | 6.21E-03 | 1.73 |
| TC1600008454.hg.1 | GABARAPL2 | GABA(A) receptor-associated protein like 2 | 3.94E-04 | 1.73 |
| TC0200014694.hg.1 | ACVR1C | activin A receptor type IC | 2.79E-03 | 1.73 |
| TC1100008038.hg.1 | SNX32 | sorting nexin 32 | 3.47E-03 | 1.73 |
| TC0400012964.hg.1 | SCLT1 | sodium channel and clathrin linker 1 | 1.07E-03 | 1.73 |
| TC1600007957.hg.1 | MT2A | metallothionein 2A | 5.30E-03 | 1.73 |
| TC0200014204.hg.1 | HS6ST1 | heparan sulfate 6-O-sulfotransferase 1 | 5.17E-03 | 1.73 |
| TC1900007040.hg.1 | CNN1 | calponin 1, basic, smooth muscle | 7.38E-03 | 1.73 |
| TC0100013747.hg.1 | SNIP1 | Smad nuclear interacting protein 1 | 2.02E-03 | 1.73 |
| TC0200013616.hg.1 | AFF3 | AF4/FMR2 family, member 3 | 8.05E-03 | 1.73 |
| TC1000008927.hg.1 | VTI1A | vesicle transport through interaction with t-SNAREs 1A | 4.44E-04 | 1.73 |
| HTA2-neg-47423802_st | NA | NA | 4.33E-03 | 1.73 |
| TC1200011991.hg.1 | IQCD | IQ motif containing D | 3.57E-03 | 1.73 |
| TC0700011050.hg.1 | FIGNL1 | fidgetin-like 1 | 3.24E-03 | 1.73 |
| TC0500013425.hg.1 | C5orf45 | chromosome 5 open reading frame 45 | 2.93E-03 | 1.73 |
| TC0200010165.hg.1 | PPP1R1C | protein phosphatase 1, regulatory (inhibitor) subunit 1C | 4.23E-03 | 1.72 |
| TC0100009215.hg.1 | CDC14A | cell division cycle 14A | 3.63E-04 | 1.72 |
| TC0800009726.hg.1 | MTMR7 | myotubularin related protein 7 | 2.60E-04 | 1.72 |
| TC1500010755.hg.1 | ANKDD1A | ankyrin repeat and death domain containing 1A | 3.08E-04 | 1.72 |
| TC1100009122.hg.1 | NXPE2 | neurexophilin and PC-esterase domain family, member 2 | 5.48E-03 | 1.72 |
| TC0700011977.hg.1 | GAL3ST4 | galactose-3-O-sulfotransferase 4 | 6.07E-03 | 1.72 |
| TC0400011188.hg.1 | LIN54 | lin-54 DREAM MuvB core complex component | 2.97E-03 | 1.72 |
| TC0200016417.hg.1 | CENPA | centromere protein A | 8.45E-04 | 1.72 |
| TC2000009813.hg.1 | EEF1A2 | eukaryotic translation elongation factor 1 alpha 2 | 1.10E-03 | 1.72 |
| TC0600013193.hg.1 | STX7 | syntaxin 7 | 1.80E-03 | 1.72 |
| TC2200008920.hg.1 | SULT4A1 | sulfotransferase family 4A member 1 | 1.98E-03 | 1.72 |
| TC0900012222.hg.1 | APTX | aprataxin | 1.02E-03 | 1.72 |
| TC1300008834.hg.1 | SIAH3 | siah E3 ubiquitin protein ligase family member 3 | 3.21E-03 | 1.72 |
| TC0700006977.hg.1 | NFE2L3 | nuclear factor, erythroid 2-like 3 | 1.01E-03 | 1.72 |
| TC0900012167.hg.1 | GSN | gelsolin | 6.44E-03 | 1.72 |
| TC1100009521.hg.1 | APLP2 | amyloid beta (A4) precursor-like protein 2 | 2.27E-03 | 1.72 |
| TC0700012728.hg.1 | PTN | pleiotrophin | 4.50E-04 | 1.72 |
| TC1900006819.hg.1 | TRIP10 | thyroid hormone receptor interactor 10 | 9.69E-04 | 1.72 |
| TC1200006771.hg.1 | CLEC2D | C-type lectin domain family 2, member D | 1.27E-03 | 1.71 |
| 23071946 | NA | NA | 9.31E-03 | 1.71 |
| TC0900011705.hg.1 | ASB6 | ankyrin repeat and SOCS box containing 6 | 8.72E-04 | 1.71 |
| TC1200006670.hg.1 | CLSTN3 | calsyntenin 3 | 8.89E-04 | 1.71 |
| TC0800008916.hg.1 | EFR3A | EFR3 homolog A | 1.40E-03 | 1.71 |
| TC1700008844.hg.1 | TTYH2 | tweety family member 2 | 6.13E-04 | 1.71 |
| TC0900011423.hg.1 | TTLL11 | tubulin tyrosine ligase-like family member 11 | 6.98E-04 | 1.71 |
| TC0500011049.hg.1 | NAIP | NLR family, apoptosis inhibitory protein | 4.92E-04 | 1.71 |
| TC1600009252.hg.1 | PPL | periplakin | 9.36E-03 | 1.71 |
| 23075636 | NA | NA | 4.07E-03 | 1.71 |
| 23076269 | NA | NA | 4.07E-03 | 1.71 |
| TC0200008534.hg.1 | ANKRD36 | ankyrin repeat domain 36 | 5.18E-03 | 1.71 |
| TC0200014775.hg.1 | KCNH7 | potassium channel, voltage gated eag related subfamily H, member 7 | 8.82E-03 | 1.71 |
| TC1100012528.hg.1 | THY1 | Thy-1 cell surface antigen | 6.33E-04 | 1.71 |
| 23066337 | NA | NA | 1.18E-03 | 1.70 |
| TC1600009545.hg.1 | ABCC6 | ATP binding cassette subfamily C member 6 | 7.30E-04 | 1.70 |
| TC0500009488.hg.1 | CREBRF | CREB3 regulatory factor | 4.44E-03 | 1.70 |
| TC1100012052.hg.1 | CCDC82 | coiled-coil domain containing 82 | 2.97E-03 | 1.70 |
| TC2000008267.hg.1 | RASSF2 | Ras association (RalGDS/AF-6) domain family member 2 | 6.30E-04 | 1.70 |
| TC2200007987.hg.1 | CLDN5 | claudin 5 | 5.99E-03 | 1.70 |
| TC0800007439.hg.1 | POLB | polymerase (DNA directed), beta | 4.13E-03 | 1.70 |
| TC0200014790.hg.1 | GRB14 | growth factor receptor bound protein 14 | 2.32E-03 | 1.70 |
| TC0700009240.hg.1 | AGBL3 | ATP/GTP binding protein-like 3 | 4.48E-03 | 1.70 |
| TC1200009550.hg.1 | KDM5A | lysine (K)-specific demethylase 5A | 6.02E-04 | 1.70 |
| TC0700013224.hg.1 | LMBR1 | Transcript Identified by AceView, Entrez Gene ID(s) 64327 | 5.46E-03 | 1.70 |
| TC0100015795.hg.1 | POGZ | Transcript Identified by AceView, Entrez Gene ID(s) 23126 | 1.56E-03 | 1.70 |
| TC0100016503.hg.1 | ASTN1 | astrotactin 1 | 9.62E-03 | 1.70 |
| TC1600006525.hg.1 | CACNA1H | calcium channel, voltage-dependent, T type, alpha 1H subunit | 3.66E-03 | 1.70 |
| TC0700006945.hg.1 | MPP6 | membrane protein, palmitoylated 6 | 2.69E-03 | 1.70 |
| TC1500009970.hg.1 | HIGD2B | HIG1 hypoxia inducible domain family, member 2B | 2.53E-04 | 1.69 |
| TC1800008235.hg.1 | ABHD3 | Transcript Identified by AceView, Entrez Gene ID(s) 171586 | 9.87E-03 | 1.69 |
| HTA2-pos-2930617_st | NA | NA | 1.55E-03 | 1.69 |
| HTA2-pos-PSR02023951.hg.1 | NA | NA | 2.52E-04 | 1.69 |
| TC0600014276.hg.1 | HLA-DMB | major histocompatibility complex, class II, DM beta | 5.84E-04 | 1.69 |
| TC1100006906.hg.1 | FAR1 | fatty acyl CoA reductase 1 | 4.73E-04 | 1.69 |
| TC0100011451.hg.1 | CAMK1G | calcium/calmodulin-dependent protein kinase IG | 3.44E-04 | 1.69 |
| TC1400007495.hg.1 | PLEKHH1 | pleckstrin homology domain containing, family H (with MyTH4 domain) member 1 | 7.26E-03 | 1.69 |
| TC1700012241.hg.1 | TBC1D3L | TBC1 domain family, member 3L | 7.64E-04 | 1.69 |
| TC0500012519.hg.1 | SPARC | secreted protein, acidic, cysteine-rich (osteonectin) | 4.89E-04 | 1.69 |
| TC1900008747.hg.1 | ZNF765 | zinc finger protein 765 | 9.58E-04 | 1.69 |
| TC0500013207.hg.1 | LIX1 | Memczak2013 ANTISENSE, coding, INTERNAL, intronic best transcript NM_153234 | 1.03E-03 | 1.69 |
| TC2000008940.hg.1 | PIGU | Salzman2013 ALT_ACCEPTOR, ALT_DONOR, coding, INTERNAL, intronic best transcript NM_080476 | 6.36E-03 | 1.69 |
| 23075808 | NA | NA | 1.57E-03 | 1.69 |
| TSUnmapped00000513.hg.1 | DGKD | diacylglycerol kinase, delta 130kDa | 8.29E-04 | 1.69 |
| TC1600010593.hg.1 | CMTM4 | CKLF-like MARVEL transmembrane domain containing 4 | 1.47E-03 | 1.69 |
| TC0800011137.hg.1 | RAD54B | RAD54 homolog B (S. cerevisiae) | 2.55E-03 | 1.69 |
| TC0900010370.hg.1 | TRPM3 | transient receptor potential cation channel, subfamily M, member 3 | 7.99E-03 | 1.69 |
| TC1400009579.hg.1 | ADAM20 | ADAM metallopeptidase domain 20 | 7.31E-04 | 1.68 |
| TC1500007034.hg.1 | SNAP23 | synaptosome associated protein 23kDa | 4.62E-03 | 1.68 |
| TC1800009215.hg.1 | ANKRD12 | ankyrin repeat domain 12 | 1.72E-03 | 1.68 |
| TC0800007065.hg.1 | CDCA2 | cell division cycle associated 2 | 2.82E-03 | 1.68 |
| TC0100006723.hg.1 | VAMP3 | vesicle associated membrane protein 3 | 6.00E-03 | 1.68 |
| TC2200007505.hg.1 | 44077 | septin 3 | 8.58E-03 | 1.68 |
| TC1900006766.hg.1 | HSD11B1L | hydroxysteroid (11-beta) dehydrogenase 1-like | 9.25E-04 | 1.68 |
| TC1600007317.hg.1 | KIAA0556 | KIAA0556 | 3.36E-03 | 1.68 |
| TC0500006479.hg.1 | NKD2 | naked cuticle homolog 2 (Drosophila) | 1.28E-03 | 1.68 |
| TC0900011938.hg.1 | C9orf116 | chromosome 9 open reading frame 116 | 8.42E-04 | 1.68 |
| TC0400008329.hg.1 | GSTCD | glutathione S-transferase, C-terminal domain containing | 2.52E-03 | 1.68 |
| TC0X00009972.hg.1 | SLC7A3 | solute carrier family 7 (cationic amino acid transporter, y+ system), member 3 | 5.36E-03 | 1.68 |
| TC0100018361.hg.1 | TSNAX-DISC1 | TSNAX-DISC1 readthrough (NMD candidate) | 3.65E-04 | 1.68 |
| TC2000008107.hg.1 | TCEA2 | transcription elongation factor A (SII), 2 | 2.12E-04 | 1.68 |
| TC0700011982.hg.1 | GATS | GATS, stromal antigen 3 opposite strand | 9.81E-03 | 1.67 |
| TC0400009088.hg.1 | GUCY1B3 | guanylate cyclase 1, soluble, beta 3 | 2.22E-03 | 1.67 |
| TC1500008545.hg.1 | ASB7 | ankyrin repeat and SOCS box containing 7 | 8.23E-03 | 1.67 |
| TC0700011784.hg.1 | FAM133B | Transcript Identified by AceView, Entrez Gene ID(s) 257415 | 1.49E-03 | 1.67 |
| TC1000008743.hg.1 | SUFU | Transcript Identified by AceView, Entrez Gene ID(s) 51684 | 4.13E-04 | 1.67 |
| TC1200008734.hg.1 | RIC8B | RIC8 guanine nucleotide exchange factor B | 2.81E-04 | 1.67 |
| TC2000008666.hg.1 | NAPB | N-ethylmaleimide-sensitive factor attachment protein, beta | 2.27E-03 | 1.67 |
| 23076268 | NA | NA | 8.48E-04 | 1.67 |
| TC0300010035.hg.1 | SENP5 | SUMO1/sentrin specific peptidase 5 | 2.88E-04 | 1.67 |
| TSUnmapped00000410.hg.1 | ATG16L1 | autophagy related 16-like 1 | 6.59E-03 | 1.67 |
| TC1700009049.hg.1 | PGS1 | phosphatidylglycerophosphate synthase 1 | 7.47E-04 | 1.67 |
| TC0300013520.hg.1 | CLDN1 | claudin 1 | 4.69E-03 | 1.67 |
| TC1700009417.hg.1 | SMG6 | SMG6 nonsense mediated mRNA decay factor | 5.53E-04 | 1.67 |
| TC1600009580.hg.1 | XYLT1 | xylosyltransferase I | 1.78E-03 | 1.67 |
| TC1500010236.hg.1 | GOLGA6L10 | golgin A6 family-like 10 | 1.04E-03 | 1.67 |
| TC1200012593.hg.1 | CLEC4A | C-type lectin domain family 4, member A | 4.14E-04 | 1.66 |
| TC0500013215.hg.1 | LVRN | laeverin | 6.04E-03 | 1.66 |
| TC2200007021.hg.1 | RFPL1 | ret finger protein-like 1 | 6.19E-04 | 1.66 |
| TC0500012285.hg.1 | FCHSD1 | FCH and double SH3 domains 1 | 5.45E-03 | 1.66 |
| 23072563 | NA | NA | 1.96E-04 | 1.66 |
| TC2000009447.hg.1 | DPM1 | dolichyl-phosphate mannosyltransferase polypeptide 1, catalytic subunit | 2.39E-04 | 1.66 |

| | | | | |
|---|---|---|---|---|
| TC1600008656.hg.1 | KLHL36 | kelch-like family member 36 | 2.36E-03 | 1.66 |
| TC0100018285.hg.1 | NBPF19 | neuroblastoma breakpoint family, member 19 | 1.72E-03 | 1.66 |
| TC0X00011291.hg.1 | PAGE2 | P antigen family, member 2 (prostate associated) | 2.08E-03 | 1.66 |
| TC1900007043.hg.1 | ZNF833P | zinc finger protein 833, pseudogene | 4.23E-03 | 1.66 |
| TC1600010685.hg.1 | SMPD3 | sphingomyelin phosphodiesterase 3, neutral membrane (neutral sphingomyelinase II) | 2.07E-03 | 1.66 |
| TC1200010865.hg.1 | ITGA7 | integrin alpha 7 | 7.30E-03 | 1.66 |
| TC0700009079.hg.1 | TSPAN33 | tetraspanin 33 | 4.43E-04 | 1.66 |
| 23075460 | NA | NA | 3.20E-04 | 1.66 |
| TC1500009458.hg.1 | ARPP19 | cAMP-regulated phosphoprotein 19kDa | 4.55E-03 | 1.66 |
| 23070237 | NA | NA | 3.48E-03 | 1.66 |
| TC0800008371.hg.1 | SPAG1 | sperm associated antigen 1 | 3.49E-03 | 1.66 |
| TC0300012164.hg.1 | POLQ | polymerase (DNA directed), theta | 1.73E-03 | 1.66 |
| TSUnmapped00000105.hg.1 | ZNF501 | zinc finger protein 501 [Source:HGNC Symbol;Acc:HGNC:23717] | 7.60E-03 | 1.65 |
| TC2200008862.hg.1 | CYP2D6 | cytochrome P450, family 2, subfamily D, polypeptide 6 | 2.94E-04 | 1.65 |
| TC0100015896.hg.1 | NUP210L | nucleoporin 210kDa like | 8.96E-03 | 1.65 |
| TC1900011852.hg.1 | RFX2 | regulatory factor X, 2 (influences HLA class II expression) | 5.98E-04 | 1.65 |
| 23075463 | NA | NA | 8.86E-03 | 1.65 |
| TC0500010998.hg.1 | CCDC125 | coiled-coil domain containing 125 | 8.86E-03 | 1.65 |
| TC0800012325.hg.1 | SLC26A7 | solute carrier family 26 (anion exchanger), member 7 | 3.18E-03 | 1.65 |
| TC2000006464.hg.1 | FAM110A | family with sequence similarity 110, member A | 2.53E-03 | 1.65 |
| TC0300012720.hg.1 | PLSCR4 | phospholipid scramblase 4 | 1.71E-03 | 1.65 |
| TC1000011743.hg.1 | CFAP43 | cilia and flagella associated protein 43 | 8.37E-03 | 1.65 |
| TC0200014703.hg.1 | CCDC148 | coiled-coil domain containing 148 | 4.09E-04 | 1.65 |
| TC0200016601.hg.1 | GIGYF2 | GRB10 interacting GYF protein 2 | 4.09E-03 | 1.65 |
| TC0700010189.hg.1 | CYTH3 | cytohesin 3 | 3.28E-03 | 1.65 |
| TC2000009178.hg.1 | JPH2 | junctophilin 2 | 4.77E-03 | 1.65 |
| TC1000008891.hg.1 | DUSP5 | dual specificity phosphatase 5 | 6.90E-03 | 1.65 |
| TC0500011495.hg.1 | SPATA9 | spermatogenesis associated 9 | 7.35E-03 | 1.65 |
| TC0100015707.hg.1 | HIST2H4A | histone cluster 2, H4a | 4.38E-03 | 1.65 |
| TC0100018208.hg.1 | ZBTB8B | zinc finger and BTB domain containing 8B | 6.37E-04 | 1.64 |
| 23074047 | NA | NA | 9.72E-03 | 1.64 |
| TC1900011892.hg.1 | RTBDN | retbindin | 4.46E-03 | 1.64 |
| TC0100006675.hg.1 | KCNAB2 | potassium channel, voltage gated subfamily A regulatory beta subunit 2 | 2.02E-03 | 1.64 |
| TC0600013519.hg.1 | ZC3H12D | zinc finger CCCH-type containing 12D | 3.27E-04 | 1.64 |
| TC0600014240.hg.1 | LOC100130357 | uncharacterized LOC100130357 | 7.58E-03 | 1.64 |
| TC2000010007.hg.1 | RBM12 | RNA binding motif protein 12 | 1.11E-03 | 1.64 |
| TC0X00007371.hg.1 | PAGE2B | P antigen family, member 2B | 6.57E-03 | 1.64 |
| TC0600007266.hg.1 | HIST1H1C | Memczak2013 ANTISENSE, CDS, coding, INTERNAL best transcript NM_005319 | 1.71E-03 | 1.64 |
| TC0600007887.hg.1 | ZFAND3 | zinc finger, AN1-type domain 3 | 1.22E-03 | 1.64 |
| TC1100010733.hg.1 | SPI1 | Spi-1 proto-oncogene | 7.84E-04 | 1.64 |
| TC0100018264.hg.1 | GSTM2 | glutathione S-transferase mu 2 (muscle) | 5.64E-03 | 1.64 |
| TC0600007869.hg.1 | CMTR1 | cap methyltransferase 1 | 4.46E-03 | 1.64 |
| TC1100011029.hg.1 | VWCE | von Willebrand factor C and EGF domains | 7.74E-04 | 1.64 |
| TC1400010331.hg.1 | XRCC3 | X-ray repair complementing defective repair in Chinese hamster cells 3 | 4.48E-03 | 1.63 |
| AFFX-PheX-M_st | NA | NA | 3.34E-03 | 1.63 |
| TC0200009687.hg.1 | ARL6IP6 | ADP-ribosylation factor like GTPase 6 interacting protein 6 | 4.62E-03 | 1.63 |
| TC0800008946.hg.1 | TG | thyroglobulin | 1.89E-03 | 1.63 |
| TC0100011219.hg.1 | PPP1R12B | protein phosphatase 1, regulatory subunit 12B | 7.58E-04 | 1.63 |
| TC0900009076.hg.1 | STKLD1 | serine/threonine kinase-like domain containing 1 | 6.59E-03 | 1.63 |
| TC1600011560.hg.1 | MTSS1L | metastasis suppressor 1-like | 2.06E-03 | 1.63 |
| TC1000007678.hg.1 | MTRNR2L5 | MT-RNR2-like 5 | 4.87E-03 | 1.63 |
| TC1400008056.hg.1 | IFI27 | interferon, alpha-inducible protein 27 | 6.45E-04 | 1.63 |
| TC1900009302.hg.1 | PIP5K1C | phosphatidylinositol-4-phosphate 5-kinase, type I, gamma | 6.93E-03 | 1.63 |
| TC0600009623.hg.1 | ECT2L | epithelial cell transforming 2 like | 8.03E-03 | 1.63 |
| TC0600008515.hg.1 | KCNQ5 | Jeck2013 ALT_ACCEPTOR, ALT_DONOR, coding, INTERNAL, intronic best transcript NM_001160133 | 8.25E-03 | 1.63 |
| TC1900011891.hg.1 | HOOK2 | hook microtubule-tethering protein 2 | 7.60E-04 | 1.63 |
| TC0300006618.hg.1 | SYN2 | synapsin II | 7.70E-03 | 1.63 |
| TC1500010319.hg.1 | KLHL25 | kelch-like family member 25 | 9.90E-03 | 1.63 |
| TC0100015102.hg.1 | COL11A1 | collagen, type XI, alpha 1 | 4.27E-03 | 1.63 |
| TC0100013634.hg.1 | TMEM54 | transmembrane protein 54 | 1.21E-03 | 1.62 |
| HTA2-pos-PSR02004962.hg.1 | NA | NA | 1.16E-03 | 1.62 |
| TC0200011890.hg.1 | LAPTM4A | Memczak2013 ALT_ACCEPTOR, ALT_DONOR, coding, INTERNAL, intronic best transcript NM_014713 | 7.18E-03 | 1.62 |
| TC1100013061.hg.1 | NADSYN1 | NAD synthetase 1 | 5.61E-03 | 1.62 |
| TC0700013526.hg.1 | FAM126A | family with sequence similarity 126, member A | 5.43E-04 | 1.62 |
| TC1700010515.hg.1 | TBC1D3 | TBC1 domain family, member 3 | 7.04E-04 | 1.62 |
| 23074024 | NA | NA | 9.14E-03 | 1.62 |
| TC0100007638.hg.1 | SERINC2 | serine incorporator 2 | 1.46E-03 | 1.62 |
| 23068496 | NA | NA | 1.82E-03 | 1.62 |
| TC0100009897.hg.1 | CA14 | carbonic anhydrase XIV | 8.51E-03 | 1.62 |
| TC0800007414.hg.1 | GOLGA7 | golgin A7 | 6.33E-04 | 1.62 |
| TC1700008539.hg.1 | MRC2 | mannose receptor, C type 2 | 8.96E-04 | 1.62 |
| TC2000007476.hg.1 | STK4 | serine/threonine kinase 4 | 6.38E-04 | 1.62 |
| TC0700010355.hg.1 | AGR2 | anterior gradient 2, protein disulphide isomerase family member | 9.06E-04 | 1.62 |
| TC0500011408.hg.1 | TMEM161B | transmembrane protein 161B | 7.29E-04 | 1.62 |
| TC0X00006477.hg.1 | CD99 | CD99 molecule | 8.11E-04 | 1.62 |
| TC0700012547.hg.1 | OPN1SW | opsin 1 (cone pigments), short-wave-sensitive | 7.17E-04 | 1.62 |
| TC0500009580.hg.1 | SIMC1 | SUMO-interacting motifs containing 1 | 2.02E-03 | 1.62 |
| TC1200009464.hg.1 | EP400NL | EP400 N-terminal like | 5.43E-03 | 1.62 |
| HTA2-neg-47420566_st | NA | NA | 4.93E-03 | 1.62 |
| TC1300006966.hg.1 | WBP4 | Transcript Identified by AceView, Entrez Gene ID(s) 11193 | 5.59E-04 | 1.61 |
| HTA2-neg-47422462_st | NA | NA | 1.30E-03 | 1.61 |
| TC1900010067.hg.1 | PBX4 | pre-B-cell leukemia homeobox 4 | 6.22E-04 | 1.61 |
| TC0800012306.hg.1 | MTFR1 | mitochondrial fission regulator 1 | 7.88E-03 | 1.61 |
| TC0200016454.hg.1 | KIAA1841 | KIAA1841 | 8.37E-03 | 1.61 |
| TC1900008999.hg.1 | ZNF549 | zinc finger protein 549 | 4.74E-03 | 1.61 |
| 23070220 | NA | NA | 8.77E-03 | 1.61 |
| TC0700009770.hg.1 | DPP6 | dipeptidyl-peptidase 6 | 7.32E-03 | 1.61 |
| TC0700012228.hg.1 | LAMB4 | laminin, beta 4 | 5.00E-03 | 1.61 |
| TC1900011766.hg.1 | MARK4 | MAP/microtubule affinity-regulating kinase 4 | 1.40E-03 | 1.61 |
| TC1600008869.hg.1 | ZNF778 | zinc finger protein 778 | 4.89E-03 | 1.61 |
| TC0500012980.hg.1 | FAM193B | family with sequence similarity 193, member B | 1.39E-03 | 1.61 |
| TC0X00008836.hg.1 | PLXNA3 | plexin A3 | 2.87E-03 | 1.61 |
| TC0100018224.hg.1 | FAM183A | family with sequence similarity 183, member A | 8.58E-04 | 1.61 |
| TC0400006487.hg.1 | TMEM175 | transmembrane protein 175 | 5.56E-03 | 1.61 |
| TC2100008533.hg.1 | PCBP3 | poly(rC) binding protein 3 | 1.59E-03 | 1.61 |
| 23064098 | NA | NA | 4.63E-03 | 1.61 |
| TC0500013322.hg.1 | NAIP | NLR family, apoptosis inhibitory protein | 1.45E-03 | 1.61 |
| TC0200008811.hg.1 | SH3RF3 | SH3 domain containing ring finger 3 | 8.31E-04 | 1.61 |
| TC1100006495.hg.1 | TSPAN4 | tetraspanin 4 | 4.67E-03 | 1.61 |
| TC1700011910.hg.1 | CYTH1 | cytohesin 1 | 3.38E-03 | 1.61 |
| TC0300009773.hg.1 | FETUB | fetuin B | 8.30E-04 | 1.61 |
| TC0100008066.hg.1 | PTPRF | protein tyrosine phosphatase, receptor type, F | 2.95E-03 | 1.61 |
| AFFX-r2-Bs-dap-5_st | NA | NA | 1.70E-03 | 1.60 |
| HTA2-neg-47421950_st | NA | NA | 7.68E-03 | 1.60 |
| TC0500012659.hg.1 | FABP6 | Memczak2013 ANTISENSE, coding, INTERNAL, intronic best transcript NM_001040442 | 1.62E-03 | 1.60 |
| TC0400007495.hg.1 | DCUN1D4 | DCN1, defective in cullin neddylation 1, domain containing 4 | 7.84E-03 | 1.60 |
| TC1100007786.hg.1 | TMEM132A | transmembrane protein 132A | 8.64E-04 | 1.60 |
| 23076534 | NA | NA | 4.76E-03 | 1.60 |
| TC1300008918.hg.1 | RCBTB1 | regulator of chromosome condensation (RCC1) and BTB (POZ) domain containing protein 1 | 3.53E-03 | 1.60 |
| TC1700012027.hg.1 | ENTHD2 | ENTH domain containing 2 | 9.31E-03 | 1.60 |
| TC1200010702.hg.1 | SMAGP | small cell adhesion glycoprotein | 1.43E-03 | 1.60 |
| TC0X00010445.hg.1 | ESX1 | ESX homeobox 1 | 9.05E-04 | 1.60 |
| TC0400008686.hg.1 | C4orf33 | chromosome 4 open reading frame 33 | 5.01E-03 | 1.60 |
| TC1500008346.hg.1 | PRC1-AS1 | PRC1 antisense RNA 1 | 2.75E-03 | 1.60 |
| TC0100007678.hg.1 | HDAC1 | histone deacetylase 1 | 1.02E-03 | 1.60 |
| TC1100007943.hg.1 | PLCB3 | phospholipase C, beta 3 (phosphatidylinositol-specific) | 1.98E-03 | 1.60 |
| TC0500009423.hg.1 | NPM1 | Zhang2013 ALT_ACCEPTOR, ALT_DONOR, coding, INTERNAL, intronic best transcript NM_002520 | 1.83E-03 | 1.60 |
| TC0700009680.hg.1 | TMEM176A | transmembrane protein 176A | 4.31E-03 | 1.60 |
| TC0800007119.hg.1 | ESCO2 | establishment of sister chromatid cohesion N-acetyltransferase 2 | 4.99E-03 | 1.60 |

| | | | | |
|---|---|---|---|---|
| TC1200010156.hg.1 | CASC1 | cancer susceptibility candidate 1 | 1.04E-03 | 1.59 |
| TC1900006684.hg.1 | SHD | Src homology 2 domain containing transforming protein D | 8.82E-03 | 1.59 |
| TC0100011316.hg.1 | NFASC | neurofascin | 6.75E-04 | 1.59 |
| TC0900010925.hg.1 | ZNF782 | zinc finger protein 782 | 8.86E-03 | 1.59 |
| TC0100016035.hg.1 | ETV3 | ets variant 3 | 3.54E-03 | 1.59 |
| 23076202 | NA | NA | 2.90E-03 | 1.59 |
| TC0600011590.hg.1 | SPDEF | SAM pointed domain containing ETS transcription factor | 1.32E-03 | 1.59 |
| TC1400009329.hg.1 | RTN1 | reticulon 1 | 5.72E-03 | 1.59 |
| TC1700011083.hg.1 | SAMD14 | sterile alpha motif domain containing 14 | 7.29E-04 | 1.59 |
| TC1100011052.hg.1 | FADS3 | fatty acid desaturase 3 | 7.82E-03 | 1.59 |
| TC2000006627.hg.1 | CHGB | chromogranin B | 6.86E-03 | 1.59 |
| 23075583 | NA | NA | 1.34E-03 | 1.59 |
| TC0100018081.hg.1 | ZNF670 | zinc finger protein 670 | 3.55E-03 | 1.59 |
| TC0200008223.hg.1 | DNAH6 | dynein, axonemal, heavy chain 6 | 1.60E-03 | 1.59 |
| TC0700011574.hg.1 | SPDYE16 | speedy/RINGO cell cycle regulator family member E16 [Source:HGNC Symbol;Acc:HGNC:51512] | 4.79E-03 | 1.59 |
| TC0100016122.hg.1 | IGSF8 | immunoglobulin superfamily, member 8 | 6.32E-03 | 1.59 |
| TC2200008257.hg.1 | LRP5L | LDL receptor related protein 5 like | 4.65E-03 | 1.59 |
| TC1700008014.hg.1 | ADAM11 | ADAM metallopeptidase domain 11 | 7.16E-03 | 1.59 |
| TC2100007327.hg.1 | RRP1 | ribosomal RNA processing 1 | 4.18E-03 | 1.59 |
| 23069208 | NA | NA | 1.24E-03 | 1.58 |
| TC1700011208.hg.1 | TRIM25 | tripartite motif containing 25 | 3.68E-03 | 1.58 |
| HTA2-neg-47422105_st | NA | NA | 3.87E-03 | 1.58 |
| TC0200013008.hg.1 | CD207 | CD207 molecule, langerin | 8.14E-03 | 1.58 |
| TC1100008085.hg.1 | PELI3 | pellino E3 ubiquitin protein ligase family member 3 | 1.21E-03 | 1.58 |
| HTA2-pos-PSR09020086.hg.1 | NA | NA | 8.78E-04 | 1.58 |
| TC1800007969.hg.1 | L3MBTL4 | l(3)mbt-like 4 (Drosophila) | 3.79E-03 | 1.58 |
| TC1700007645.hg.1 | CTB-75G16.3 | --- | 1.10E-03 | 1.58 |
| TC1300009012.hg.1 | THSD1 | thrombospondin type 1 domain containing 1 | 7.24E-03 | 1.58 |
| TC0700012584.hg.1 | UBE2H | ubiquitin conjugating enzyme E2H | 5.24E-04 | 1.58 |
| TC1900007090.hg.1 | BEST2 | bestrophin 2 | 2.75E-03 | 1.58 |
| TC0600008622.hg.1 | SH3BGRL2 | SH3 domain binding glutamate-rich protein like 2 | 4.89E-04 | 1.58 |
| TC1800008162.hg.1 | FAM210A | family with sequence similarity 210, member A | 4.13E-03 | 1.58 |
| 23070298 | NA | NA | 3.05E-03 | 1.58 |
| TC0X00010877.hg.1 | FAM122B | family with sequence similarity 122B | 7.01E-04 | 1.58 |
| TC0600011805.hg.1 | USP49 | ubiquitin specific peptidase 49 | 3.36E-03 | 1.58 |
| TC0900008197.hg.1 | GALNT12 | polypeptide N-acetylgalactosaminyltransferase 12 | 9.49E-04 | 1.58 |
| TC0100018553.hg.1 | PLEKHA6 | pleckstrin homology domain containing, family A member 6 | 2.25E-03 | 1.58 |
| TC1100012681.hg.1 | SIAE | sialic acid acetylesterase | 2.37E-03 | 1.58 |
| TC1200012248.hg.1 | HCAR3 | hydroxycarboxylic acid receptor 3 | 1.53E-03 | 1.58 |
| TC0600014251.hg.1 | ZBED9 | zinc finger, BED-type containing 9 | 9.69E-04 | 1.58 |
| TC1500010134.hg.1 | CIB2 | calcium and integrin binding family member 2 | 1.89E-03 | 1.58 |
| TC0900009776.hg.1 | MOB3B | MOB kinase activator 3B | 7.12E-03 | 1.58 |
| TC1700009402.hg.1 | SMYD4 | SET and MYND domain containing 4 | 3.20E-03 | 1.58 |
| 23066358 | NA | NA | 2.49E-03 | 1.57 |
| TC2000009748.hg.1 | CABLES2 | Cdk5 and Abl enzyme substrate 2 | 1.80E-03 | 1.57 |
| TC1400007706.hg.1 | FOS | FBJ murine osteosarcoma viral oncogene homolog | 2.79E-03 | 1.57 |
| TC1100011609.hg.1 | P4HA3 | prolyl 4-hydroxylase, alpha polypeptide III | 4.74E-03 | 1.57 |
| TC0900011432.hg.1 | RBM18 | RNA binding motif protein 18 | 5.52E-03 | 1.57 |
| TC1100008010.hg.1 | CDC42EP2 | CDC42 effector protein (Rho GTPase binding) 2 | 3.40E-03 | 1.57 |
| TC1700008661.hg.1 | PRKCA | protein kinase C, alpha | 1.98E-03 | 1.57 |
| TC1200011599.hg.1 | LTA4H | leukotriene A4 hydrolase | 2.51E-03 | 1.57 |
| TC1700009383.hg.1 | PITPNA | Memczak2013 ALT_ACCEPTOR, ALT_DONOR, coding, INTERNAL, intronic best transcript NM_006224 | 5.55E-03 | 1.57 |
| TC0200016746.hg.1 | TTC21B | tetratricopeptide repeat domain 21B | 4.51E-03 | 1.57 |
| 23075424 | NA | NA | 9.53E-03 | 1.57 |
| 23070118 | NA | NA | 2.35E-03 | 1.57 |
| TC0700012712.hg.1 | LUZP6 | leucine zipper protein 6 | 1.21E-03 | 1.57 |
| 23076310 | NA | NA | 7.70E-03 | 1.57 |
| 23076464 | NA | NA | 7.70E-03 | 1.57 |
| AFFX-r2-Ec-bioB-5_at | NA | NA | 5.69E-04 | 1.57 |
| TC0900007869.hg.1 | SEMA4D | Memczak2013 ANTISENSE, coding, INTERNAL, UTR3 best transcript NM_006378 | 1.67E-03 | 1.57 |
| TC0900007050.hg.1 | DNAI1 | Transcript Identified by AceView, Entrez Gene ID(s) 27019 | 2.07E-03 | 1.57 |
| TC1600011286.hg.1 | CHMP1A | charged multivesicular body protein 1A | 9.92E-04 | 1.57 |
| TC1100009068.hg.1 | NCAM1 | neural cell adhesion molecule 1 | 4.48E-04 | 1.57 |
| TC1600010122.hg.1 | LINC00273 | long intergenic non-protein coding RNA 273 | 1.81E-03 | 1.57 |
| TC0700011779.hg.1 | ERVW-1 | endogenous retrovirus group W, member 1 | 7.25E-03 | 1.57 |
| 23070246 | NA | NA | 8.37E-03 | 1.56 |
| TC1100010990.hg.1 | MRPL16 | mitochondrial ribosomal protein L16 | 2.06E-03 | 1.56 |
| TC0100016316.hg.1 | ADCY10 | adenylate cyclase 10 (soluble) | 8.20E-04 | 1.56 |
| TC1600008231.hg.1 | NFAT5 | nuclear factor of activated T-cells 5, tonicity-responsive | 3.06E-03 | 1.56 |
| TC0500013140.hg.1 | AHRR | aryl-hydrocarbon receptor repressor | 4.82E-03 | 1.56 |
| TC1600007967.hg.1 | MT1IP | metallothionein 1I, pseudogene | 3.13E-03 | 1.56 |
| TC2000009180.hg.1 | OSER1 | oxidative stress responsive serine-rich 1 | 2.75E-03 | 1.56 |
| TC0200008516.hg.1 | CNNM3 | Zhang2013 ALT_ACCEPTOR, ALT_DONOR, coding, INTERNAL, intronic, OVERLAPTX best transcript NM_017623 | 1.12E-03 | 1.56 |
| TC0500008467.hg.1 | SRFBP1 | serum response factor binding protein 1 | 1.59E-03 | 1.56 |
| TC0300009724.hg.1 | VPS8 | vacuolar protein sorting 8 homolog (S. cerevisiae) | 5.12E-03 | 1.56 |
| TC1900011908.hg.1 | PDE4C | phosphodiesterase 4C, cAMP-specific | 1.64E-03 | 1.56 |
| TSUnmapped00000213.hg.1 | DGKD | diacylglycerol kinase, delta 130kDa | 5.15E-03 | 1.56 |
| TC0700009398.hg.1 | TAS2R3 | taste receptor, type 2, member 3 | 5.01E-03 | 1.56 |
| TC1900009743.hg.1 | TNPO2 | transportin 2 | 4.02E-03 | 1.56 |
| TC1700008440.hg.1 | PPM1E | protein phosphatase, Mg2+/Mn2+ dependent, 1E | 3.13E-03 | 1.56 |
| TC1400009563.hg.1 | SLC10A1 | solute carrier family 10 (sodium/bile acid cotransporter), member 1 | 7.30E-03 | 1.56 |
| TC0700011344.hg.1 | GUSB | glucuronidase, beta | 2.09E-03 | 1.56 |
| TC0900009307.hg.1 | ARRDC1 | arrestin domain containing 1 | 3.40E-03 | 1.56 |
| TC0100009658.hg.1 | SRGAP2C | SLIT-ROBO Rho GTPase activating protein 2C | 5.94E-04 | 1.56 |
| TC1900011596.hg.1 | C19orf18 | chromosome 19 open reading frame 18 | 1.03E-03 | 1.56 |
| TC0200013765.hg.1 | UXS1 | UDP-glucuronate decarboxylase 1 | 1.11E-03 | 1.56 |
| TC0200014029.hg.1 | C2orf76 | chromosome 2 open reading frame 76 | 1.67E-03 | 1.56 |
| TC0X00007558.hg.1 | NLGN3 | neuroligin 3 | 5.15E-03 | 1.56 |
| TC1900010997.hg.1 | DACT3 | dishevelled-binding antagonist of beta-catenin 3 | 3.73E-03 | 1.56 |
| TC1200009959.hg.1 | LRP6 | LDL receptor related protein 6 | 2.00E-03 | 1.56 |
| 23075524 | NA | NA | 1.06E-03 | 1.56 |
| TC0100010341.hg.1 | FCER1G | Fc fragment of IgE, high affinity I, receptor for; gamma polypeptide | 2.49E-03 | 1.56 |
| TC0200012009.hg.1 | DTNB | dystrobrevin beta | 4.27E-03 | 1.56 |
| TC0900010587.hg.1 | C9orf64 | Transcript Identified by AceView, Entrez Gene ID(s) 84267 | 4.18E-03 | 1.55 |
| TCUn_GL000218v100006433.hg.1 | LOC389834 | ankyrin repeat domain 57 pseudogene | 5.24E-03 | 1.55 |
| AFFX-BioB-M_at | NA | NA | 5.46E-04 | 1.55 |
| TC0500009863.hg.1 | LPCAT1 | lysophosphatidylcholine acyltransferase 1 | 2.56E-03 | 1.55 |
| TC1600010778.hg.1 | HYDIN | HYDIN, axonemal central pair apparatus protein | 2.05E-03 | 1.55 |
| 23075525 | NA | NA | 2.57E-03 | 1.55 |
| HTA2-neg-47422968_st | NA | NA | 6.28E-03 | 1.55 |
| AFFX-r2-Ec-bioB-M_at | NA | NA | 8.83E-04 | 1.55 |
| TC0300008375.hg.1 | ZDHHC23 | zinc finger, DHHC-type containing 23 | 2.31E-03 | 1.55 |
| TC0X00010661.hg.1 | RHOXF2B | Rhox homeobox family, member 2B | 8.06E-03 | 1.55 |
| TC0X00008080.hg.1 | VSIG1 | V-set and immunoglobulin domain containing 1 | 2.85E-03 | 1.55 |
| TC1000012117.hg.1 | CHST15 | carbohydrate (N-acetylgalactosamine 4-sulfate 6-O) sulfotransferase 15 | 2.33E-03 | 1.55 |
| TC0500008684.hg.1 | TCF7 | transcription factor 7 (T-cell specific, HMG-box) | 1.95E-03 | 1.55 |
| TC1200008678.hg.1 | EID3 | EP300 interacting inhibitor of differentiation 3 | 7.50E-03 | 1.55 |
| TC1900011573.hg.1 | ZNF416 | zinc finger protein 416 | 1.19E-03 | 1.55 |
| TC0100007091.hg.1 | CROCC | Zhang2013 ALT_ACCEPTOR, ALT_DONOR, coding, INTERNAL, intronic best transcript NM_014675 | 1.63E-03 | 1.55 |
| TC1100007833.hg.1 | INCENP | inner centromere protein | 2.37E-03 | 1.55 |
| TC0500007552.hg.1 | LOC100421561 | family with sequence similarity 133, member A pseudogene | 7.93E-03 | 1.55 |
| TC0700013435.hg.1 | CDHR3 | cadherin-related family member 3 | 2.35E-03 | 1.55 |
| TC0100013837.hg.1 | RIMS3 | regulating synaptic membrane exocytosis 3 | 5.07E-03 | 1.55 |
| TC1900011593.hg.1 | ZNF418 | zinc finger protein 418 | 3.17E-03 | 1.55 |
| AFFX-BioB-5_at | NA | NA | 9.17E-04 | 1.55 |
| TC1700006877.hg.1 | ADPRM | ADP-ribose/CDP-alcohol diphosphatase, manganese-dependent | 2.54E-03 | 1.55 |
| TC0300009280.hg.1 | KCNAB1 | Transcript Identified by AceView, Entrez Gene ID(s) 7881 | 2.82E-03 | 1.55 |
| TC0900008959.hg.1 | FUBP3 | far upstream element (FUSE) binding protein 3 | 5.43E-04 | 1.54 |
| ERCCmix2step23 | NA | NA | 9.50E-03 | 1.54 |

| | | | | |
|---|---|---|---|---|
| TC1600010172.hg.1 | RP11-93O14.2 | --- | 1.02E-03 | 1.54 |
| TC2200008864.hg.1 | CYP2D6 | cytochrome P450, family 2, subfamily D, polypeptide 6 | 1.03E-03 | 1.54 |
| TC0800007370.hg.1 | ADAM32 | ADAM metallopeptidase domain 32 | 5.80E-03 | 1.54 |
| 23075807 | NA | NA | 1.69E-03 | 1.54 |
| TC1200011968.hg.1 | HECTD4 | HECT domain containing E3 ubiquitin protein ligase 4 | 4.70E-03 | 1.54 |
| TC1600007262.hg.1 | TNRC6A | trinucleotide repeat containing 6A | 6.92E-03 | 1.54 |
| TC2000008095.hg.1 | TPD52L2 | tumor protein D52-like 2 | 9.62E-03 | 1.54 |
| HTA2-pos-PSR14009921.hg.1 | NA | NA | 2.83E-03 | 1.54 |
| 23075679 | NA | NA | 8.01E-03 | 1.54 |
| 23076144 | NA | NA | 8.01E-03 | 1.54 |
| TC0700008584.hg.1 | AP1S1 | Transcript Identified by AceView, Entrez Gene ID(s) 1174 | 4.22E-03 | 1.54 |
| TC0800011259.hg.1 | ANKRD46 | ankyrin repeat domain 46 | 7.10E-03 | 1.54 |
| TC1000012586.hg.1 | SEC31B | SEC31 homolog B, COPII coat complex component | 2.67E-03 | 1.54 |
| TC0700008011.hg.1 | NCF1 | neutrophil cytosolic factor 1 | 2.26E-03 | 1.54 |
| TC1100010907.hg.1 | BTBD18 | BTB (POZ) domain containing 18 | 3.49E-03 | 1.54 |
| TC1900010521.hg.1 | ZFP14 | ZFP14 zinc finger protein | 5.55E-04 | 1.54 |
| TC0X00008394.hg.1 | BCORL1 | BCL6 corepressor-like 1 | 2.84E-03 | 1.54 |
| TC0100010282.hg.1 | ATP1A4 | ATPase, Na+/K+ transporting, alpha 4 polypeptide | 5.46E-04 | 1.54 |
| TC0100018398.hg.1 | TNFRSF25 | tumor necrosis factor receptor superfamily, member 25 | 8.29E-03 | 1.54 |
| TC1500010945.hg.1 | LINS1 | lines homolog 1 | 2.15E-03 | 1.54 |
| TC0500010592.hg.1 | C6 | complement component 6 | 1.38E-03 | 1.54 |
| TC0600009262.hg.1 | SLC35F1 | solute carrier family 35, member F1 | 7.79E-03 | 1.54 |
| TC1700008846.hg.1 | KIF19 | kinesin family member 19 | 1.29E-03 | 1.53 |
| TC1200010105.hg.1 | RECQL | RecQ helicase-like | 3.73E-03 | 1.53 |
| TC1900006968.hg.1 | C19orf66 | chromosome 19 open reading frame 66 | 8.48E-03 | 1.53 |
| TC2200009259.hg.1 | SMTN | smoothelin | 1.64E-03 | 1.53 |
| TC0100015891.hg.1 | CRTC2 | CREB regulated transcription coactivator 2 | 4.21E-03 | 1.53 |
| TC1900007328.hg.1 | SLC27A1 | solute carrier family 27 (fatty acid transporter), member 1 | 2.29E-03 | 1.53 |
| TC1600011358.hg.1 | C16orf45 | chromosome 16 open reading frame 45 | 1.85E-03 | 1.53 |
| TC1100008090.hg.1 | ACTN3 | actinin, alpha 3 (gene/pseudogene) | 9.54E-04 | 1.53 |
| 23076398 | NA | NA | 1.12E-03 | 1.53 |
| 23070245 | NA | NA | 2.04E-03 | 1.53 |
| TC0800009621.hg.1 | CTSB | Jeck2013 ALT_DONOR, coding, INTERNAL, intronic best transcript NM_147780 | 5.48E-03 | 1.53 |
| TC0200012417.hg.1 | ABCG5 | ATP binding cassette subfamily G member 5 | 8.74E-03 | 1.53 |
| TC0200016446.hg.1 | ACYP2 | acylphosphatase 2, muscle type | 3.21E-03 | 1.53 |
| TC0300012190.hg.1 | KPNA1 | Transcript Identified by AceView, Entrez Gene ID(s) 3836 | 9.83E-03 | 1.53 |
| TC0500008342.hg.1 | DCP2 | decapping mRNA 2 | 2.32E-03 | 1.53 |
| TC0200011379.hg.1 | SNED1 | sushi, nidogen and EGF-like domains 1 | 5.35E-03 | 1.53 |
| TC1600008329.hg.1 | HP | haptoglobin | 8.99E-03 | 1.53 |
| TC0X00008276.hg.1 | ATP1B4 | ATPase, Na+/K+ transporting, beta 4 polypeptide | 7.05E-04 | 1.53 |
| TC0700013429.hg.1 | PILRA | paired immunoglobin-like type 2 receptor alpha | 6.98E-03 | 1.53 |
| TC0700011554.hg.1 | PMS2P3 | PMS1 homolog 2, mismatch repair system component pseudogene 3 | 2.85E-03 | 1.53 |
| TC2100008459.hg.1 | FTCD | formimidoyltransferase cyclodeaminase | 8.00E-03 | 1.53 |
| TC0300011183.hg.1 | NEK4 | NIMA-related kinase 4 | 6.41E-03 | 1.53 |
| TC0500007586.hg.1 | RGS7BP | regulator of G-protein signaling 7 binding protein | 9.35E-03 | 1.53 |
| TC1600006941.hg.1 | SNX29 | sorting nexin 29 | 5.23E-03 | 1.53 |
| HTA2-pos-PSR02004963.hg.1 | NA | NA | 4.02E-03 | 1.53 |
| TC0700009400.hg.1 | TAS2R5 | taste receptor, type 2, member 5 | 6.43E-03 | 1.53 |
| TC1600011494.hg.1 | ARL6IP1 | ADP-ribosylation factor like GTPase 6 interacting protein 1 | 4.18E-03 | 1.53 |
| TC1900007688.hg.1 | CCNE1 | cyclin E1 | 5.89E-03 | 1.53 |
| TC1600010680.hg.1 | SLC7A6OS | solute carrier family 7, member 6 opposite strand | 2.74E-03 | 1.53 |
| TC1000012549.hg.1 | PARG | poly (ADP-ribose) glycohydrolase | 1.87E-03 | 1.53 |
| TC0300013350.hg.1 | YEATS2 | Memczak2013 ANTISENSE, coding, INTERNAL, intronic best transcript NM_018023 | 1.28E-03 | 1.52 |
| TC0700011961.hg.1 | GJC3 | gap junction protein gamma 3 | 1.38E-03 | 1.52 |
| AFFX-r2-Ec-bioB-3_at | NA | NA | 1.21E-03 | 1.52 |
| TC0300014085.hg.1 | TBCCD1 | TBCC domain containing 1 | 6.94E-03 | 1.52 |
| AFFX-BioC-3_at | NA | NA | 1.36E-03 | 1.52 |
| TC1000006796.hg.1 | SEC61A2 | Sec61 translocon alpha 2 subunit | 9.58E-03 | 1.52 |
| TC1200009136.hg.1 | CABP1 | calcium binding protein 1 | 3.97E-03 | 1.52 |
| TC1700012283.hg.1 | LUC7L3 | LUC7-like 3 pre-mRNA splicing factor | 3.86E-03 | 1.52 |
| TC1900010632.hg.1 | SIRT2 | sirtuin 2 | 9.22E-03 | 1.52 |
| TC1200012707.hg.1 | ALDH2 | aldehyde dehydrogenase 2 family (mitochondrial) | 4.78E-03 | 1.52 |
| TC0300006564.hg.1 | IL17RE | interleukin 17 receptor E | 7.02E-03 | 1.52 |
| TC1800006635.hg.1 | TWSG1 | twisted gastrulation BMP signaling modulator 1 | 3.00E-03 | 1.52 |
| TC0400007128.hg.1 | TBC1D19 | TBC1 domain family, member 19 | 4.82E-03 | 1.52 |
| TC1900008600.hg.1 | GPR32 | G protein-coupled receptor 32 | 2.00E-03 | 1.52 |
| TC0700009395.hg.1 | WEE2 | WEE1 homolog 2 (S. pombe) | 6.94E-03 | 1.52 |
| TC1200012561.hg.1 | ZNF891 | zinc finger protein 891 | 7.73E-03 | 1.52 |
| TC0200009902.hg.1 | NOSTRIN | nitric oxide synthase trafficking | 6.78E-03 | 1.52 |
| TC0X00011126.hg.1 | GABRA3 | gamma-aminobutyric acid (GABA) A receptor, alpha 3 | 9.19E-03 | 1.52 |
| TC0100015572.hg.1 | SRGAP2B | SLIT-ROBO Rho GTPase activating protein 2B | 1.18E-03 | 1.52 |
| 23069198 | NA | NA | 6.97E-03 | 1.52 |
| TC0300013670.hg.1 | TNK2 | tyrosine kinase, non-receptor, 2 | 1.06E-03 | 1.52 |
| TC1900011889.hg.1 | MAN2B1 | mannosidase, alpha, class 2B, member 1 | 5.51E-03 | 1.52 |
| TC1300008426.hg.1 | USP12 | Transcript Identified by AceView, Entrez Gene ID(s) 219333 | 2.12E-03 | 1.52 |
| 23071966 | NA | NA | 7.67E-03 | 1.52 |
| TC1100013065.hg.1 | KRTAP5-9 | keratin associated protein 5-9 | 1.27E-03 | 1.52 |
| TC2200007904.hg.1 | ATP6V1E1 | ATPase, H+ transporting, lysosomal 31kDa, V1 subunit E1 | 7.46E-04 | 1.52 |
| TC0100018348.hg.1 | CNIH3 | cornichon family AMPA receptor auxiliary protein 3 | 5.63E-03 | 1.52 |
| 23075438 | NA | NA | 5.49E-03 | 1.52 |
| TC1100012930.hg.1 | B3GAT1 | beta-1,3-glucuronyltransferase 1 | 6.47E-03 | 1.52 |
| TC1700010731.hg.1 | PSMC3IP | PSMC3 interacting protein | 1.54E-03 | 1.52 |
| TC1700012242.hg.1 | LOC101060389 | TBC1 domain family member-like | 7.50E-03 | 1.52 |
| TC0Y00006476.hg.1 | CD99 | Homo sapiens CD99 molecule (CD99), transcript variant 2, mRNA. | 1.13E-03 | 1.52 |
| TC0200012068.hg.1 | SLC30A3 | solute carrier family 30 (zinc transporter), member 3 | 3.44E-03 | 1.52 |
| HTA2-pos-2930600_st | NA | NA | 7.24E-04 | 1.52 |
| 23075427 | NA | NA | 8.45E-03 | 1.52 |
| AFFX-r2-Ec-bioC-5_at | NA | NA | 1.25E-03 | 1.52 |
| TC0100016053.hg.1 | CD5L | CD5 molecule-like | 2.84E-03 | 1.51 |
| TC1700011448.hg.1 | POLG2 | polymerase (DNA directed), gamma 2, accessory subunit | 2.66E-03 | 1.51 |
| TC1900006730.hg.1 | KDM4B | lysine (K)-specific demethylase 4B | 2.87E-03 | 1.51 |
| TC0600008870.hg.1 | FUT9 | fucosyltransferase 9 (alpha (1,3) fucosyltransferase) | 4.79E-03 | 1.51 |
| TC0Y00006858.hg.1 | VAMP7 | Homo sapiens vesicle-associated membrane protein 7 (VAMP7), transcript variant 2, mRNA. | 2.67E-03 | 1.51 |
| HTA2-pos-PSR01002166.hg.1 | NA | NA | 5.01E-03 | 1.51 |
| TC1700006638.hg.1 | ARRB2 | arrestin, beta 2 | 7.20E-03 | 1.51 |
| TC1300007045.hg.1 | GPALPP1 | GPALPP motifs containing 1 | 2.81E-03 | 1.51 |
| TC0600013431.hg.1 | PLAGL1 | pleiomorphic adenoma gene-like 1 | 1.91E-03 | 1.51 |
| TC0200006539.hg.1 | ALLC | allantoicase | 2.87E-03 | 1.51 |
| TC1500009224.hg.1 | STRC | stereocilin | 4.60E-03 | 1.51 |
| TC2000009907.hg.1 | EFCAB8 | EF-hand calcium binding domain 8 | 4.24E-03 | 1.51 |
| TC2000007114.hg.1 | KIF3B | kinesin family member 3B | 3.14E-03 | 1.51 |
| TC0500006822.hg.1 | OTULIN | OTU deubiquitinase with linear linkage specificity | 1.91E-03 | 1.51 |
| TC0300008369.hg.1 | ATP6V1A | ATPase, H+ transporting, lysosomal 70kDa, V1 subunit A | 8.71E-03 | 1.51 |
| TC1900006999.hg.1 | DNM2 | Transcript Identified by AceView, Entrez Gene ID(s) 1785 | 1.08E-03 | 1.51 |
| TC0200016413.hg.1 | FKBP1B | FK506 binding protein 1B | 1.84E-03 | 1.51 |
| TC0900008463.hg.1 | HSDL2 | hydroxysteroid dehydrogenase like 2 | 7.29E-03 | 1.51 |
| TC1100010741.hg.1 | CELF1 | CUGBP, Elav-like family member 1 | 1.37E-03 | 1.51 |
| TC1500007980.hg.1 | CHRNA5 | cholinergic receptor, nicotinic alpha 5 | 6.50E-03 | 1.51 |
| TC0900012258.hg.1 | MFSD14C | major facilitator superfamily domain containing 14C | 1.11E-03 | 1.51 |
| TC1900007325.hg.1 | MVB12A | multivesicular body subunit 12A | 4.20E-03 | 1.51 |
| TC0600009167.hg.1 | WISP3 | WNT1 inducible signaling pathway protein 3 | 6.60E-03 | 1.51 |
| TC0600008722.hg.1 | ZNF292 | zinc finger protein 292 | 2.57E-03 | 1.51 |
| TC0200010897.hg.1 | SGPP2 | sphingosine-1-phosphate phosphatase 2 | 8.14E-03 | 1.51 |
| HTA2-neg-47420332_st | NA | NA | 8.54E-03 | 1.51 |
| TC0800007738.hg.1 | SDCBP | syndecan binding protein | 1.30E-03 | 1.51 |
| TC0700010182.hg.1 | PMS2 | PMS1 homolog 2, mismatch repair system component | 1.57E-03 | 1.51 |
| TC1600007739.hg.1 | PHKB | phosphorylase kinase, beta | 6.26E-03 | 1.51 |
| TC0600007616.hg.1 | HSPA1B | heat shock 70kDa protein 1B | 1.99E-03 | 1.51 |
| TC0400007456.hg.1 | ZAR1 | zygote arrest 1 | 2.35E-03 | 1.51 |

| | | | | |
|---|---|---|---|---|
| TC1900006804.hg.1 | TNFSF9 | tumor necrosis factor (ligand) superfamily, member 9 | 1.05E-03 | 1.51 |
| TC1600009958.hg.1 | NPIPB4 | nuclear pore complex interacting protein family, member B4 | 1.24E-03 | 1.51 |
| TSUnmapped00000460.hg.1 | DGKD | diacylglycerol kinase, delta 130kDa | 6.01E-03 | 1.51 |
| TC1900011931.hg.1 | SYNE4 | spectrin repeat containing, nuclear envelope family member 4 | 4.40E-03 | 1.51 |
| TC0100007943.hg.1 | RLF | rearranged L-myc fusion | 2.05E-03 | 1.51 |
| TC0300012980.hg.1 | SPTSSB | serine palmitoyltransferase, small subunit B | 9.29E-03 | 1.51 |
| 23074924 | NA | NA | 1.12E-03 | 1.51 |
| TC1000008088.hg.1 | SAMD8 | sterile alpha motif domain containing 8 | 3.47E-03 | 1.50 |
| TC1400010615.hg.1 | MNAT1 | MNAT CDK-activating kinase assembly factor 1 | 8.21E-04 | 1.50 |
| TC1600009042.hg.1 | CLCN7 | chloride channel, voltage-sensitive 7 | 7.50E-03 | 1.50 |
| TC1000007564.hg.1 | MAPK8 | mitogen-activated protein kinase 8 | 3.72E-03 | 1.50 |
| TC0400010141.hg.1 | LDB2 | LIM domain binding 2 | 5.21E-03 | 1.50 |
| TC1300007707.hg.1 | GPC5 | glypican 5 | 4.22E-03 | 1.50 |
| TC1900007100.hg.1 | MAST1 | microtubule associated serine/threonine kinase 1 | 2.01E-03 | 1.50 |
| TC0500007120.hg.1 | TTC23L | tetratricopeptide repeat domain 23-like | 4.23E-03 | 1.50 |
| TC0600014006.hg.1 | WDR27 | WD repeat domain 27 | 3.41E-03 | 1.50 |
| TC0500011754.hg.1 | TMED7-TICAM2 | TMED7-TICAM2 readthrough | 8.21E-03 | 1.50 |
| TC0200009071.hg.1 | CFAP221 | cilia and flagella associated protein 221 | 1.40E-03 | 1.50 |
| 23075740 | NA | NA | 2.79E-03 | 1.50 |
| TC0100011534.hg.1 | FAM71A | family with sequence similarity 71, member A | 3.44E-03 | 1.50 |
| TC0200016660.hg.1 | C2orf61 | chromosome 2 open reading frame 61 | 2.42E-03 | 1.50 |

BGB290 treatment vs control

| Probe ID | Symbol | Gene name | P value | Fold change |
|---|---|---|---|---|
| TC0600014190.hg.1 | ZC2HC1B | zinc finger, C2HC-type containing 1B | 4.28E-03 | 5.60 |
| TC1200011845.hg.1 | SELPLG | selectin P ligand | 5.33E-03 | 5.09 |
| 23076543 | NA | NA | 5.03E-05 | 4.67 |
| TC0700013472.hg.1 | MGAM | maltase-glucoamylase | 4.09E-05 | 4.04 |
| TC0300014032.hg.1 | POPDC2 | popeye domain containing 2 | 7.87E-05 | 3.85 |
| HTA2-pos-3295162_st | NA | NA | 1.04E-04 | 3.80 |
| TC1200008686.hg.1 | CHST11 | Transcript Identified by AceView, Entrez Gene ID(s) 50515 | 2.18E-03 | 3.68 |
| TC0600009597.hg.1 | TNFAIP3 | tumor necrosis factor, alpha-induced protein 3 | 1.14E-03 | 3.40 |
| TC0500012017.hg.1 | IRF1 | interferon regulatory factor 1 | 6.32E-03 | 3.13 |
| TC1900006977.hg.1 | ICAM1 | intercellular adhesion molecule 1 | 8.16E-04 | 3.07 |
| HTA2-neg-47424041_st | NA | NA | 9.05E-03 | 3.03 |
| TC2100007599.hg.1 | SAMSN1 | SAM domain, SH3 domain and nuclear localization signals 1 | 7.08E-04 | 2.97 |
| TC2100008297.hg.1 | SIK1 | salt-inducible kinase 1 | 7.78E-03 | 2.92 |
| TC1200007053.hg.1 | SPX | spexin hormone | 3.18E-03 | 2.89 |
| HTA2-neg-47422282_st | NA | NA | 8.84E-03 | 2.88 |
| TC0700013442.hg.1 | LSMEM1 | leucine-rich single-pass membrane protein 1 | 1.03E-03 | 2.86 |
| HTA2-neg-47424062_st | NA | NA | 1.73E-03 | 2.84 |
| TC0500007077.hg.1 | NPR3 | natriuretic peptide receptor 3 | 5.08E-04 | 2.80 |
| TC2000009964.hg.1 | SDCBP2 | syndecan binding protein (syntenin) 2 | 3.53E-04 | 2.73 |
| TC0300012007.hg.1 | CD200R1 | CD200 receptor 1 | 2.66E-03 | 2.71 |
| TC1000006911.hg.1 | TMEM236 | transmembrane protein 236 | 2.36E-03 | 2.70 |
| TC0700009411.hg.1 | MGAM2 | maltase-glucoamylase 2 (putative) | 3.78E-04 | 2.67 |
| TC0700008495.hg.1 | BUD31 | Transcript Identified by AceView, Entrez Gene ID(s) 8896 | 4.62E-05 | 2.65 |
| HTA2-pos-47421926_st | NA | NA | 2.02E-03 | 2.62 |
| TC0400008240.hg.1 | DAPP1 | dual adaptor of phosphotyrosine and 3-phosphoinositides | 8.72E-03 | 2.58 |
| HTA2-pos-47421925_st | NA | NA | 1.05E-03 | 2.57 |
| TC1300008813.hg.1 | KCTD4 | potassium channel tetramerization domain containing 4 | 1.99E-03 | 2.54 |
| TC0800011566.hg.1 | RAD21 | Transcript Identified by AceView, Entrez Gene ID(s) 5885 | 4.45E-03 | 2.52 |
| TC0900010737.hg.1 | DIRAS2 | DIRAS family, GTP-binding RAS-like 2 | 1.23E-03 | 2.51 |
| HTA2-pos-2960249_st | NA | NA | 9.75E-05 | 2.44 |
| TC0700013424.hg.1 | GS1-259H13.2 | transmembrane protein 225-like | 2.16E-03 | 2.37 |
| TC0600013231.hg.1 | SGK1 | serum/glucocorticoid regulated kinase 1 | 2.73E-03 | 2.36 |
| TC0400007360.hg.1 | DCAF4L1 | DDB1 and CUL4 associated factor 4-like 1 | 1.79E-03 | 2.35 |
| TC1100008188.hg.1 | PPP6R3 | Memczak2013 ALT_ACCEPTOR, ALT_DONOR, coding, INTERNAL, intronic best transcript NM_001164162 | 2.79E-04 | 2.35 |
| TC1300008048.hg.1 | TEX29 | testis expressed 29 | 1.50E-03 | 2.32 |
| TC2000009945.hg.1 | FAM209A | family with sequence similarity 209, member A | 6.42E-05 | 2.24 |
| TC0900008219.hg.1 | NR4A3 | nuclear receptor subfamily 4, group A, member 3 | 6.97E-03 | 2.22 |
| HTA2-neg-47422433_st | NA | NA | 3.59E-04 | 2.22 |
| HTA2-pos-47421982_st | NA | NA | 1.81E-03 | 2.20 |
| TC1600006593.hg.1 | RAB26 | RAB26, member RAS oncogene family | 8.76E-03 | 2.17 |
| TC1900011817.hg.1 | ZNF773 | zinc finger protein 773 | 5.17E-05 | 2.14 |
| HTA2-pos-2985881_st | NA | NA | 1.12E-03 | 2.13 |
| TC0200014361.hg.1 | LYPD1 | LY6/PLAUR domain containing 1 | 5.06E-03 | 2.13 |
| HTA2-neg-47422979_st | NA | NA | 8.83E-03 | 2.13 |
| TC0500011334.hg.1 | TMEM167A | Transcript Identified by AceView, Entrez Gene ID(s) 153339 | 2.52E-03 | 2.11 |
| TC0300007050.hg.1 | C3orf35 | chromosome 3 open reading frame 35 | 2.64E-04 | 2.10 |
| TC0200015402.hg.1 | FAM126B | family with sequence similarity 126, member B | 1.10E-03 | 2.09 |
| HTA2-neg-47423115_st | NA | NA | 4.14E-05 | 2.06 |
| TC1100007216.hg.1 | PRRG4 | proline rich Gla (G-carboxyglutamic acid) 4 (transmembrane) | 8.31E-03 | 2.05 |
| TC0600007263.hg.1 | HIST1H4A | histone cluster 1, H4a | 3.20E-03 | 2.05 |
| TC1500010745.hg.1 | POLR2M | polymerase (RNA) II (DNA directed) polypeptide M | 5.61E-03 | 2.05 |
| HTA2-pos-2960237_st | NA | NA | 2.45E-03 | 2.04 |
| TC1100010505.hg.1 | ABTB2 | ankyrin repeat and BTB (POZ) domain containing 2 | 6.09E-04 | 2.03 |
| HTA2-neg-47423121_st | NA | NA | 2.61E-03 | 2.01 |
| HTA2-neg-47422060_st | NA | NA | 2.04E-03 | 2.01 |
| TC2000008211.hg.1 | UBOX5 | U-box domain containing 5 | 1.95E-03 | 2.01 |
| TC2100007140.hg.1 | ETS2 | v-ets avian erythroblastosis virus E26 oncogene homolog 2 | 7.29E-03 | 2.00 |
| TC0500012870.hg.1 | STC2 | stanniocalcin 2 | 1.20E-04 | 2.00 |
| TC0700013587.hg.1 | SHFM1 | split hand/foot malformation (ectrodactyly) type 1 | 1.32E-03 | 1.99 |
| TC1500009458.hg.1 | ARPP19 | cAMP-regulated phosphoprotein 19kDa | 7.19E-04 | 1.98 |
| TC0100008631.hg.1 | SGIP1 | SH3-domain GRB2-like (endophilin) interacting protein 1 | 7.52E-03 | 1.97 |
| 23076544 | NA | NA | 1.07E-03 | 1.97 |
| HTA2-pos-2985880_st | NA | NA | 4.20E-03 | 1.96 |
| TC1100007729.hg.1 | DTX4 | deltex 4, E3 ubiquitin ligase | 9.09E-04 | 1.96 |
| TC1400008940.hg.1 | NFKBIA | nuclear factor of kappa light polypeptide gene enhancer in B-cells inhibitor, alpha | 2.22E-03 | 1.95 |
| TC0600012064.hg.1 | GCM1 | glial cells missing homolog 1 (Drosophila) | 6.78E-03 | 1.95 |
| TC1700010560.hg.1 | PLXDC1 | plexin domain containing 1 | 1.89E-03 | 1.94 |
| TC0500008541.hg.1 | TEX43 | testis expressed 43 | 7.95E-04 | 1.94 |
| TC0700010899.hg.1 | POLR2J4 | polymerase (RNA) II (DNA directed) polypeptide J4, pseudogene | 2.29E-03 | 1.93 |
| TC1700006763.hg.1 | ATP1B2 | ATPase, Na+/K+ transporting, beta 2 polypeptide | 9.42E-03 | 1.93 |
| TC0500009984.hg.1 | FLJ33360 | FLJ33360 protein | 4.69E-03 | 1.92 |
| TC0900007877.hg.1 | GADD45G | growth arrest and DNA-damage-inducible, gamma | 5.52E-04 | 1.92 |
| TC0300009916.hg.1 | HES1 | hes family bHLH transcription factor 1 | 1.91E-03 | 1.91 |
| TC0200010586.hg.1 | CPO | carboxypeptidase O | 3.44E-03 | 1.91 |
| TC0100009417.hg.1 | C1orf162 | chromosome 1 open reading frame 162 | 3.86E-04 | 1.90 |
| TC0600012502.hg.1 | GJB7 | gap junction protein beta 7 | 4.44E-03 | 1.89 |
| TC0X00006585.hg.1 | SHROOM2 | shroom family member 2 | 5.43E-03 | 1.89 |
| TSUnmapped00000354.hg.1 | EIF3F | Eukaryotic translation initiation factor 3 subunit F [Source:UniProtKB/Swiss-Prot;Acc:O00303] | 7.91E-03 | 1.89 |
| TC0400008389.hg.1 | RRH | retinal pigment epithelium-derived rhodopsin homolog | 8.86E-04 | 1.88 |
| TC0200014620.hg.1 | CACNB4 | calcium channel, voltage-dependent, beta 4 subunit | 8.99E-04 | 1.88 |
| TC0600010056.hg.1 | WTAP | Wilms tumor 1 associated protein | 3.81E-03 | 1.87 |
| TC1800006733.hg.1 | PRELID3A | PRELI domain containing 3A | 1.32E-03 | 1.85 |
| TC0500009059.hg.1 | GRPEL2 | GrpE-like 2, mitochondrial (E. coli) | 7.59E-05 | 1.85 |
| TC0200012977.hg.1 | MXD1 | Memczak2013 ANTISENSE, coding, INTERNAL, intronic best transcript NM_001202514 | 8.19E-03 | 1.85 |
| TC0400011421.hg.1 | LAMTOR3 | late endosomal/lysosomal adaptor, MAPK and MTOR activator 3 | 5.85E-03 | 1.85 |
| HTA2-neg-47423502_st | NA | NA | 2.51E-03 | 1.84 |
| HTA2-neg-47421633_st | NA | NA | 8.96E-04 | 1.84 |
| TC1200009192.hg.1 | BCL7A | B-cell CLL/lymphoma 7A | 8.29E-03 | 1.84 |
| TC0200013638.hg.1 | LONRF2 | LON peptidase N-terminal domain and ring finger 2 | 2.64E-03 | 1.84 |
| TC0200016661.hg.1 | C2orf61 | chromosome 2 open reading frame 61 | 4.73E-03 | 1.84 |
| TC0200007746.hg.1 | B3GNT2 | UDP-GlcNAc:betaGal beta-1,3-N-acetylglucosaminyltransferase 2 | 4.90E-03 | 1.83 |
| HTA2-neg-47422334_st | NA | NA | 5.45E-03 | 1.82 |

| | | | | |
|---|---|---|---|---|
| HTA2-neg-47424061_st | NA | NA | 5.73E-03 | 1.82 |
| TC0100010341.hg.1 | FCER1G | Fc fragment of IgE, high affinity I, receptor for; gamma polypeptide | 1.36E-04 | 1.82 |
| TC0500007050.hg.1 | PDZD2 | PDZ domain containing 2 | 1.38E-03 | 1.81 |
| TC0500007895.hg.1 | CMYA5 | cardiomyopathy associated 5 | 2.21E-03 | 1.81 |
| TC1300009810.hg.1 | ANKRD10 | ankyrin repeat domain 10 | 3.94E-03 | 1.81 |
| TC2000009886.hg.1 | PANK2 | pantothenate kinase 2 | 5.91E-04 | 1.80 |
| TC0100015271.hg.1 | OVGP1 | oviductal glycoprotein 1 | 2.40E-03 | 1.80 |
| TC1200012737.hg.1 | ZNF268 | zinc finger protein 268 | 1.43E-04 | 1.80 |
| TC0200008904.hg.1 | FBLN7 | fibulin 7 | 6.57E-03 | 1.80 |
| TC1000012117.hg.1 | CHST15 | carbohydrate (N-acetylgalactosamine 4-sulfate 6-O) sulfotransferase 15 | 4.12E-04 | 1.79 |
| TC2000006791.hg.1 | PCSK2 | proprotein convertase subtilisin/kexin type 2 | 4.40E-03 | 1.79 |
| TC2100007967.hg.1 | PAXBP1 | PAX3 and PAX7 binding protein 1 | 2.31E-04 | 1.78 |
| TC0200008894.hg.1 | MERTK | MER proto-oncogene, tyrosine kinase | 4.14E-03 | 1.77 |
| TC0700011458.hg.1 | CALN1 | calneuron 1 | 4.88E-03 | 1.77 |
| TC0100018248.hg.1 | RNPC3 | RNA binding region (RNP1, RRM) containing 3 | 8.48E-04 | 1.76 |
| TC2000007094.hg.1 | TTLL9 | tubulin tyrosine ligase-like family member 9 | 3.77E-03 | 1.76 |
| TC1100012389.hg.1 | CADM1 | cell adhesion molecule 1 | 2.14E-03 | 1.76 |
| TC1400008415.hg.1 | ZFYVE21 | zinc finger, FYVE domain containing 21 | 8.02E-04 | 1.76 |
| AFFX-DapX-5_st | NA | NA | 4.53E-03 | 1.75 |
| TC0800007529.hg.1 | SPIDR | scaffolding protein involved in DNA repair | 5.32E-03 | 1.75 |
| TC0900008790.hg.1 | ZBTB43 | zinc finger and BTB domain containing 43 | 2.59E-03 | 1.75 |
| TC0800011713.hg.1 | MTSS1 | metastasis suppressor 1 | 4.17E-03 | 1.75 |
| TC0600006870.hg.1 | SNRNP48 | small nuclear ribonucleoprotein, U11/U12 48KDa subunit | 2.81E-03 | 1.75 |
| TC1300008938.hg.1 | DLEU2 | deleted in lymphocytic leukemia 2 (non-protein coding) | 6.60E-03 | 1.74 |
| TC2000006756.hg.1 | MACROD2 | MACRO domain containing 2 | 6.45E-03 | 1.74 |
| TC0700006945.hg.1 | MPP6 | membrane protein, palmitoylated 6 | 2.65E-04 | 1.73 |
| TC1400007145.hg.1 | FRMD6 | FERM domain containing 6 | 6.53E-03 | 1.73 |
| HTA2-pos-PSR02023970.hg.1 | NA | NA | 1.10E-03 | 1.72 |
| TC0500009488.hg.1 | CREBRF | CREB3 regulatory factor | 6.98E-03 | 1.72 |
| TC2000010008.hg.1 | NFS1 | NFS1 cysteine desulfurase | 3.54E-03 | 1.72 |
| TC1900009588.hg.1 | ZNF121 | zinc finger protein 121 | 9.74E-04 | 1.72 |
| TC0X00011275.hg.1 | TCEANC | transcription elongation factor A (SII) N-terminal and central domain containing | 3.61E-03 | 1.71 |
| TC1100010531.hg.1 | SLC1A2 | solute carrier family 1 (glial high affinity glutamate transporter), member 2 | 3.53E-03 | 1.71 |
| TC1400008622.hg.1 | OR5AU1 | olfactory receptor, family 5, subfamily AU, member 1 | 7.36E-03 | 1.71 |
| TC1600007147.hg.1 | TMEM159 | transmembrane protein 159 | 6.65E-03 | 1.71 |
| 23071851 | NA | NA | 7.38E-03 | 1.71 |
| 23071994 | NA | NA | 7.38E-03 | 1.71 |
| AFFX-DapX-M_st | NA | NA | 5.66E-03 | 1.71 |
| TC1500008986.hg.1 | LPCAT4 | lysophosphatidylcholine acyltransferase 4 | 9.88E-03 | 1.70 |
| TC1000009612.hg.1 | KLF6 | Kruppel-like factor 6 | 3.16E-03 | 1.70 |
| TC0900008793.hg.1 | ZBTB34 | zinc finger and BTB domain containing 34 | 1.02E-03 | 1.70 |
| TC0500010635.hg.1 | HMGCS1 | 3-hydroxy-3-methylglutaryl-CoA synthase 1 (soluble) | 3.41E-03 | 1.70 |
| TC0700011782.hg.1 | FAM133B | family with sequence similarity 133, member B | 8.36E-03 | 1.70 |
| TC1200011965.hg.1 | NAA25 | N(alpha)-acetyltransferase 25, NatB auxiliary subunit | 8.21E-04 | 1.70 |
| TC1200008255.hg.1 | ZDHHC17 | zinc finger, DHHC-type containing 17 | 5.39E-03 | 1.70 |
| HTA2-pos-PSR02023951.hg.1 | NA | NA | 2.51E-03 | 1.69 |
| AFFX-r2-Bs-phe-M_st | NA | NA | 2.05E-03 | 1.69 |
| TC1500010886.hg.1 | CALML4 | calmodulin-like 4 | 1.80E-03 | 1.69 |
| HTA2-neg-47420666_st | NA | NA | 5.81E-03 | 1.69 |
| TC0300006462.hg.1 | TRNT1 | tRNA nucleotidyl transferase, CCA-adding, 1 | 4.92E-04 | 1.69 |
| TC0200015424.hg.1 | ALS2CR12 | amyotrophic lateral sclerosis 2 chromosome region candidate 12 | 4.45E-03 | 1.69 |
| TC1200007108.hg.1 | KRAS | Memczak2013 ANTISENSE, coding, INTERNAL, intronic best transcript NM_033360 | 1.17E-03 | 1.69 |
| TC1500010723.hg.1 | CHAC1 | ChaC glutathione-specific gamma-glutamylcyclotransferase 1 | 1.31E-03 | 1.69 |
| TC0200014362.hg.1 | NCKAP5 | NCK-associated protein 5 | 1.66E-03 | 1.69 |
| TC0800012447.hg.1 | AZIN1 | antizyme inhibitor 1 | 2.89E-03 | 1.69 |
| TC0200009428.hg.1 | CCNT2 | cyclin T2 | 3.01E-03 | 1.68 |
| TC0600007262.hg.1 | HIST1H3A | histone cluster 1, H3a | 7.11E-03 | 1.68 |
| TC0400008331.hg.1 | NPNT | nephronectin | 3.15E-03 | 1.68 |
| TC1600006533.hg.1 | UBE2I | ubiquitin conjugating enzyme E2I | 2.16E-03 | 1.68 |
| TC1800008456.hg.1 | ZSCAN30 | zinc finger and SCAN domain containing 30 | 3.95E-03 | 1.67 |
| TC0100007876.hg.1 | RRAGC | Memczak2013 ANTISENSE, coding, INTERNAL, intronic best transcript NM_022157 | 4.99E-03 | 1.67 |
| TC0700011876.hg.1 | ASNS | asparagine synthetase (glutamine-hydrolyzing) | 8.07E-03 | 1.67 |
| TC1900010782.hg.1 | ATP1A3 | ATPase, Na+/K+ transporting, alpha 3 polypeptide | 2.62E-03 | 1.67 |
| TC1600010291.hg.1 | SALL1 | spalt-like transcription factor 1 | 8.52E-04 | 1.67 |
| TC1500010157.hg.1 | ADAMTS7 | ADAM metallopeptidase with thrombospondin type 1 motif 7 | 3.84E-03 | 1.67 |
| TC0600014151.hg.1 | SMIM8 | small integral membrane protein 8 | 5.20E-03 | 1.67 |
| TC1900011596.hg.1 | C19orf18 | chromosome 19 open reading frame 18 | 3.76E-04 | 1.67 |
| TC1700007918.hg.1 | AOC2 | amine oxidase, copper containing 2 (retina-specific) | 4.38E-03 | 1.66 |
| TC1200012668.hg.1 | TRHDE | thyrotropin-releasing hormone degrading enzyme | 3.11E-03 | 1.66 |
| HTA2-neg-47422968_st | NA | NA | 1.18E-03 | 1.66 |
| TC1900009743.hg.1 | TNPO2 | transportin 2 | 1.28E-03 | 1.66 |
| TC1900009867.hg.1 | AKAP8L | A kinase (PRKA) anchor protein 8-like | 1.03E-03 | 1.66 |
| TC1500010788.hg.1 | CHD2 | chromodomain helicase DNA binding protein 2 | 3.95E-03 | 1.66 |
| TC0100015976.hg.1 | KIAA0907 | KIAA0907 | 4.52E-03 | 1.66 |
| TC1000012565.hg.1 | DNAJC9 | DnaJ (Hsp40) homolog, subfamily C, member 9 | 3.83E-03 | 1.66 |
| TC1700011919.hg.1 | CEP295NL | CEP295 N-terminal like | 7.12E-03 | 1.66 |
| HTA2-neg-47419608_st | NA | NA | 4.81E-03 | 1.65 |
| TC0400007495.hg.1 | DCUN1D4 | DCN1, defective in cullin neddylation 1, domain containing 4 | 3.33E-03 | 1.65 |
| TSUnmapped00000105.hg.1 | ZNF501 | zinc finger protein 501 [Source:HGNC Symbol;Acc:HGNC:23717] | 3.72E-03 | 1.65 |
| TC0200010374.hg.1 | MARS2 | methionyl-tRNA synthetase 2, mitochondrial | 5.47E-03 | 1.64 |
| TC0200016601.hg.1 | GIGYF2 | GRB10 interacting GYF protein 2 | 5.43E-03 | 1.64 |
| HTA2-neg-47423547_st | NA | NA | 4.56E-03 | 1.64 |
| TC1900011593.hg.1 | ZNF418 | zinc finger protein 418 | 3.77E-03 | 1.64 |
| TC1700007557.hg.1 | CCL2 | chemokine (C-C motif) ligand 2 | 5.43E-03 | 1.64 |
| TC0600009262.hg.1 | SLC35F1 | solute carrier family 35, member F1 | 8.17E-04 | 1.64 |
| TC0700013525.hg.1 | FAM126A | family with sequence similarity 126, member A | 6.00E-03 | 1.64 |
| TC0300007044.hg.1 | GOLGA4 | golgin A4 | 1.09E-03 | 1.64 |
| TC1200012775.hg.1 | ST8SIA1 | ST8 alpha-N-acetyl-neuraminide alpha-2,8-sialyltransferase 1 | 2.48E-03 | 1.63 |
| HTA2-pos-47422722_st | NA | NA | 1.37E-03 | 1.63 |
| TC0700009240.hg.1 | AGBL3 | ATP/GTP binding protein-like 3 | 8.22E-03 | 1.63 |
| TC0100010926.hg.1 | OCLM | Transcript Identified by AceView, Entrez Gene ID(s) 10896 | 4.15E-03 | 1.63 |
| TC1700009746.hg.1 | MYH3 | myosin, heavy chain 3, skeletal muscle, embryonic | 4.40E-03 | 1.63 |
| TC0500008467.hg.1 | SRFBP1 | serum response factor binding protein 1 | 1.92E-03 | 1.63 |
| AFFX-r2-Bs-dap-M_st | NA | NA | 7.13E-03 | 1.63 |
| TC1500007975.hg.1 | IREB2 | iron responsive element binding protein 2 | 1.62E-03 | 1.63 |
| TC2000009966.hg.1 | FKBP1A-SDCBP2 | FKBP1A-SDCBP2 readthrough (NMD candidate) | 2.51E-03 | 1.63 |
| TC1000007217.hg.1 | KIF5B | Memczak2013 ANTISENSE, CDS, coding, INTERNAL, UTR3 best transcript NM_004521 | 8.03E-03 | 1.63 |
| AFFX-r2-Bs-dap-3_st | NA | NA | 8.00E-03 | 1.63 |
| HTA2-pos-47422721_st | NA | NA | 6.00E-03 | 1.63 |
| TC0500009423.hg.1 | NPM1 | Zhang2013 ALT_ACCEPTOR, ALT_DONOR, coding, INTERNAL, intronic best transcript NM_002520 | 1.53E-03 | 1.63 |
| TC0100009900.hg.1 | CIART | circadian associated repressor of transcription | 3.69E-03 | 1.62 |
| TC1000008927.hg.1 | VTI1A | vesicle transport through interaction with t-SNAREs 1A | 3.07E-03 | 1.62 |
| TC0100010925.hg.1 | OCLM | oculomedin | 3.80E-03 | 1.62 |
| TC1600011351.hg.1 | CARHSP1 | Jeck2013 ANTISENSE, CDS, coding, INTERNAL, OVCODE, OVEXON best transcript NM_001042476 | 4.12E-03 | 1.61 |
| 23072532 | NA | NA | 9.39E-03 | 1.61 |
| TC0500009012.hg.1 | TCERG1 | Transcript Identified by AceView, Entrez Gene ID(s) 10915 | 7.79E-03 | 1.61 |
| TC1100012712.hg.1 | ACRV1 | acrosomal vesicle protein 1 | 9.97E-03 | 1.61 |
| TC0100007690.hg.1 | RBBP4 | retinoblastoma binding protein 4 | 6.66E-04 | 1.61 |
| TC2000006627.hg.1 | CHGB | chromogranin B | 1.59E-03 | 1.61 |
| TC1900006578.hg.1 | SF3A2 | splicing factor 3a subunit 2 | 1.11E-03 | 1.61 |
| HTA2-neg-47422985_st | NA | NA | 8.52E-03 | 1.60 |
| TC1900011814.hg.1 | ZNF17 | zinc finger protein 17 | 1.35E-03 | 1.60 |
| TC0X00007562.hg.1 | ITGB1BP2 | integrin beta 1 binding protein (melusin) 2 | 2.13E-03 | 1.60 |
| TC0600014276.hg.1 | HLA-DMB | major histocompatibility complex, class II, DM beta | 8.96E-03 | 1.60 |
| TC0400008103.hg.1 | RP11-10L7.1 | --- | 5.40E-03 | 1.60 |
| TC0400008335.hg.1 | AIMP1 | aminoacyl tRNA synthetase complex-interacting multifunctional protein 1 | 1.79E-03 | 1.59 |
| TC0200015876.hg.1 | SERPINE2 | serpin peptidase inhibitor, clade E (nexin, plasminogen activator inhibitor type 1), member 2 | 1.37E-03 | 1.59 |
| TC1400010632.hg.1 | GPATCH2L | G-patch domain containing 2 like | 1.39E-03 | 1.59 |
| TC1600010711.hg.1 | CHTF8 | chromosome transmission fidelity factor 8 | 3.84E-03 | 1.59 |

| | | | | |
|---|---|---|---|---|
| TC0500012030.hg.1 | GDF9 | growth differentiation factor 9 | 8.06E-03 | 1.59 |
| TC0400010141.hg.1 | LDB2 | LIM domain binding 2 | 4.88E-03 | 1.59 |
| TC1100007293.hg.1 | LDLRAD3 | low density lipoprotein receptor class A domain containing 3 | 9.11E-03 | 1.59 |
| TC1800009215.hg.1 | ANKRD12 | ankyrin repeat domain 12 | 6.13E-03 | 1.58 |
| TC1300009714.hg.1 | ARGLU1 | arginine and glutamate rich 1 | 1.85E-03 | 1.58 |
| TC1000008806.hg.1 | CFAP58 | cilia and flagella associated protein 58 | 3.15E-03 | 1.58 |
| TC0300009280.hg.1 | KCNAB1 | Transcript Identified by AceView, Entrez Gene ID(s) 7881 | 7.09E-03 | 1.58 |
| TC0500010920.hg.1 | SREK1IP1 | SREK1-interacting protein 1 | 1.64E-03 | 1.58 |
| HTA2-pos-PSR02004962.hg.1 | NA | NA | 3.20E-03 | 1.58 |
| TC1500009466.hg.1 | ONECUT1 | one cut homeobox 1 | 7.20E-03 | 1.58 |
| HTA2-pos-3183407_st | NA | NA | 7.19E-03 | 1.57 |
| TC2100007534.hg.1 | TEKT4P2 | tektin 4 pseudogene 2 | 5.36E-03 | 1.57 |
| TC0400009221.hg.1 | MSMO1 | methylsterol monooxygenase 1 | 1.83E-03 | 1.57 |
| TC0700009827.hg.1 | RBM33 | RNA binding motif protein 33 | 1.16E-03 | 1.57 |
| AFFX-DapX-3_st | NA | NA | 7.66E-03 | 1.57 |
| TC1200009881.hg.1 | CLEC2B | C-type lectin domain family 2, member B | 1.64E-03 | 1.57 |
| TC1700008769.hg.1 | KCNJ2 | potassium channel, inwardly rectifying subfamily J, member 2 | 5.66E-03 | 1.57 |
| TC0100009112.hg.1 | LOC729970 | hCG2028352-like | 6.88E-03 | 1.56 |
| HTA2-pos-PSR01002164.hg.1 | NA | NA | 4.10E-03 | 1.56 |
| TC1900011816.hg.1 | ZNF419 | zinc finger protein 419 | 9.41E-04 | 1.56 |
| TC1700012483.hg.1 | MAFG | v-maf avian musculoaponeurotic fibrosarcoma oncogene homolog G | 5.22E-03 | 1.56 |
| HTA2-pos-PSR10021746.hg.1 | NA | NA | 2.63E-03 | 1.56 |
| TC1800008891.hg.1 | BCL2 | B-cell CLL/lymphoma 2 | 7.63E-03 | 1.56 |
| TC0100009383.hg.1 | RBM15 | RNA binding motif protein 15 | 3.10E-03 | 1.56 |
| TC0900010177.hg.1 | CNTNAP3B | Salzman2013 ANNOTATED, CDS, coding, INTERNAL, OVCODE, OVEXON best transcript NM_001201380 | 9.32E-03 | 1.55 |
| AFFX-PheX-M_st | NA | NA | 1.30E-03 | 1.55 |
| TC1100010990.hg.1 | MRPL16 | mitochondrial ribosomal protein L16 | 3.80E-03 | 1.55 |
| TC0600014366.hg.1 | PDE10A | phosphodiesterase 10A | 7.59E-03 | 1.55 |
| TC0600008379.hg.1 | MTRNR2L9 | MT-RNR2-like 9 | 4.53E-03 | 1.55 |
| TC0100008066.hg.1 | PTPRF | protein tyrosine phosphatase, receptor type, F | 5.67E-03 | 1.55 |
| TC1300006690.hg.1 | POLR1D | polymerase (RNA) I polypeptide D | 4.94E-03 | 1.55 |
| TC1600011560.hg.1 | MTSS1L | metastasis suppressor 1-like | 3.83E-03 | 1.55 |
| TSUnmapped00000057.hg.1 | SURF4 | surfeit 4 | 3.64E-03 | 1.54 |
| TC1800009306.hg.1 | RNF152 | ring finger protein 152 | 3.77E-03 | 1.54 |
| HTA2-pos-PSR02004963.hg.1 | NA | NA | 3.91E-03 | 1.54 |
| TC0200008734.hg.1 | C2orf49 | chromosome 2 open reading frame 49 | 1.58E-03 | 1.54 |
| TC0X00011306.hg.1 | NHSL2 | NHS-like 2 | 9.23E-03 | 1.54 |
| TC1200009997.hg.1 | GRIN2B | glutamate receptor, ionotropic, N-methyl D-aspartate 2B | 6.80E-03 | 1.53 |
| TC0500012519.hg.1 | SPARC | secreted protein, acidic, cysteine-rich (osteonectin) | 2.93E-03 | 1.53 |
| TC0300006714.hg.1 | EAF1 | ELL associated factor 1 | 5.28E-03 | 1.53 |
| TC0200010745.hg.1 | IGFBP2 | insulin like growth factor binding protein 2 | 7.32E-03 | 1.53 |
| HTA2-pos-PSR02023963.hg.1 | NA | NA | 3.21E-03 | 1.53 |
| TC1900011734.hg.1 | MIA | melanoma inhibitory activity | 2.21E-03 | 1.53 |
| TC1100012930.hg.1 | B3GAT1 | beta-1,3-glucuronyltransferase 1 | 1.41E-03 | 1.53 |
| TC0600012540.hg.1 | GABRR2 | gamma-aminobutyric acid (GABA) A receptor, rho 2 | 4.21E-03 | 1.53 |
| TC1900011573.hg.1 | ZNF416 | zinc finger protein 416 | 3.80E-03 | 1.53 |
| TC1000006464.hg.1 | GTPBP4 | GTP binding protein 4 | 5.66E-03 | 1.53 |
| TC0400011180.hg.1 | SCD5 | stearoyl-CoA desaturase 5 | 4.21E-03 | 1.53 |
| TC2000009247.hg.1 | SPATA25 | spermatogenesis associated 25 | 8.33E-03 | 1.52 |
| HTA2-neg-47424009_st | NA | NA | 7.88E-03 | 1.52 |
| TC0400010242.hg.1 | PPARGC1A | peroxisome proliferator-activated receptor gamma, coactivator 1 alpha | 6.07E-03 | 1.52 |
| TC0500010540.hg.1 | LIFR | leukemia inhibitory factor receptor alpha | 4.26E-03 | 1.52 |
| TC1800008550.hg.1 | SYT4 | synaptotagmin IV | 2.91E-03 | 1.52 |
| TC0400011014.hg.1 | CXCL5 | chemokine (C-X-C motif) ligand 5 | 9.08E-03 | 1.52 |
| TC0700009222.hg.1 | LRGUK | leucine-rich repeats and guanylate kinase domain containing | 3.09E-03 | 1.52 |
| TC2100008533.hg.1 | PCBP3 | poly(rC) binding protein 3 | 2.29E-03 | 1.52 |
| TC0600011868.hg.1 | SRF | Jeck2013 ANTISENSE, CDS, coding, INTERNAL, intronic, OVCODE, OVEXON best transcript NM_003131 | 9.18E-03 | 1.52 |
| TC1100013229.hg.1 | FXYD6 | FXYD domain containing ion transport regulator 6 | 6.27E-03 | 1.52 |
| TC0900012222.hg.1 | APTX | aprataxin | 2.12E-03 | 1.52 |
| TC2000006872.hg.1 | INSM1 | insulinoma-associated 1 | 5.83E-03 | 1.52 |
| TC0X00009356.hg.1 | TMEM47 | transmembrane protein 47 | 5.78E-03 | 1.51 |
| TC1200012667.hg.1 | TPH2 | tryptophan hydroxylase 2 | 9.51E-03 | 1.51 |
| TC2100007082.hg.1 | DYRK1A | dual specificity tyrosine-(Y)-phosphorylation regulated kinase 1A | 6.93E-03 | 1.51 |
| HTA2-pos-JUC01000985.hg.1 | NA | NA | 4.16E-03 | 1.51 |
| HTA2-pos-JUC13008384.hg.1 | NA | NA | 4.16E-03 | 1.51 |
| HTA2-pos-JUC19003067.hg.1 | NA | NA | 4.16E-03 | 1.51 |
| TC1100011182.hg.1 | PYGM | phosphorylase, glycogen, muscle | 2.91E-03 | 1.51 |
| TC0200009806.hg.1 | TANK | TRAF family member-associated NFKB activator | 1.89E-03 | 1.51 |
| TC1800007969.hg.1 | L3MBTL4 | l(3)mbt-like 4 (Drosophila) | 3.64E-03 | 1.51 |
| TC0500008540.hg.1 | PHAX | phosphorylated adaptor for RNA export | 5.34E-03 | 1.51 |
| TC1500010890.hg.1 | HEXA | hexosaminidase A (alpha polypeptide) | 8.70E-03 | 1.51 |
| TC0600014236.hg.1 | TXNDC5 | thioredoxin domain containing 5 (endoplasmic reticulum) | 4.18E-03 | 1.51 |
| TC1000010515.hg.1 | NCOA4 | nuclear receptor coactivator 4 | 3.85E-03 | 1.51 |
| AFFX-PheX-3_st | NA | NA | 2.06E-03 | 1.50 |
| TC0100010284.hg.1 | PEA15 | phosphoprotein enriched in astrocytes 15 | 1.79E-03 | 1.50 |
| TC1200012579.hg.1 | NTF3 | neurotrophin 3 | 4.32E-03 | 1.50 |
| TC0200007446.hg.1 | PRKCE | protein kinase C, epsilon | 4.91E-03 | 1.50 |

Romidepsin treatment vs control

| Probe ID | Symbol | Gene name | P value | Fold change |
|---|---|---|---|---|
| TC1400006697.hg.1 | DHRS2 | dehydrogenase/reductase (SDR family) member 2 | 2.20E-07 | 19.43 |
| TC1900008057.hg.1 | ZFP36 | ZFP36 ring finger protein | 7.70E-05 | 9.50 |
| TC0700008582.hg.1 | SERPINE1 | serpin peptidase inhibitor, clade E (nexin, plasminogen activator inhibitor type 1), member 1 | 2.71E-04 | 8.90 |
| TC0800009891.hg.1 | STC1 | stanniocalcin 1 | 1.76E-06 | 7.69 |
| TC0500006744.hg.1 | ROPN1L | rhophilin associated tail protein 1-like | 1.69E-05 | 7.10 |
| TC1100010054.hg.1 | NRIP3 | nuclear receptor interacting protein 3 | 9.02E-05 | 6.36 |
| TC1700006763.hg.1 | ATP1B2 | ATPase, Na+/K+ transporting, beta 2 polypeptide | 3.35E-07 | 6.20 |
| TC0700007977.hg.1 | FZD9 | frizzled class receptor 9 | 6.76E-07 | 6.14 |
| TC1900006537.hg.1 | REEP6 | receptor accessory protein 6 | 7.51E-06 | 5.75 |
| TC1600007931.hg.1 | LPCAT2 | lysophosphatidylcholine acyltransferase 2 | 2.65E-06 | 5.63 |
| HTA2-pos-2978683_st | NA | NA | 1.84E-04 | 5.59 |
| TC0300009412.hg.1 | SERPINI1 | serpin peptidase inhibitor, clade I (neuroserpin), member 1 | 6.92E-06 | 5.51 |
| TC1900009240.hg.1 | GNG7 | guanine nucleotide binding protein (G protein), gamma 7 | 4.76E-04 | 5.30 |
| TC0600006873.hg.1 | BMP6 | bone morphogenetic protein 6 | 6.40E-06 | 5.23 |
| TSUnmapped00000485.hg.1 | RPS6KA1 | ribosomal protein S6 kinase, 90kDa, polypeptide 1 | 9.84E-08 | 5.10 |
| TC1700008635.hg.1 | RGS9 | regulator of G-protein signaling 9 | 6.22E-04 | 5.06 |
| TC1000007895.hg.1 | TSPAN15 | tetraspanin 15 | 1.59E-05 | 5.06 |
| TC0100007705.hg.1 | AZIN2 | antizyme inhibitor 2 | 6.05E-04 | 5.01 |
| TC0200015869.hg.1 | AP1S3 | adaptor-related protein complex 1 sigma 3 subunit | 3.18E-04 | 5.01 |
| TC0800011620.hg.1 | ENPP2 | ectonucleotide pyrophosphatase/phosphodiesterase 2 | 7.80E-04 | 4.94 |
| TC1600009677.hg.1 | CRYM | crystallin mu | 2.06E-05 | 4.76 |
| TC1000010768.hg.1 | EGR2 | early growth response 2 | 2.63E-04 | 4.76 |
| TC1700009318.hg.1 | FAM101B | family with sequence similarity 101, member B | 9.04E-06 | 4.75 |
| TC1700008673.hg.1 | CACNG5 | calcium channel, voltage-dependent, gamma subunit 5 | 1.05E-03 | 4.69 |
| TC0X00008747.hg.1 | GABRQ | gamma-aminobutyric acid (GABA) A receptor, theta | 3.41E-06 | 4.64 |
| TC1500009606.hg.1 | MYO1E | Transcript Identified by AceView, Entrez Gene ID(s) 4643 | 2.72E-04 | 4.51 |
| TC2000007202.hg.1 | ACSS2 | acyl-CoA synthetase short-chain family member 2 | 3.34E-05 | 4.45 |
| TC0100017018.hg.1 | ETNK2 | ethanolamine kinase 2 | 7.31E-06 | 4.40 |
| TC0500008785.hg.1 | EGR1 | early growth response 1 | 1.15E-06 | 4.39 |
| TC1500007972.hg.1 | CRABP1 | cellular retinoic acid binding protein 1 | 1.87E-04 | 4.33 |
| TC0800010926.hg.1 | PAG1 | phosphoprotein membrane anchor with glycosphingolipid microdomains 1 | 2.49E-07 | 4.29 |
| TC0700007105.hg.1 | ADCYAP1R1 | adenylate cyclase activating polypeptide 1 (pituitary) receptor type I | 7.67E-03 | 4.23 |
| TC0200008894.hg.1 | MERTK | MER proto-oncogene, tyrosine kinase | 3.81E-06 | 4.22 |
| TC1900010016.hg.1 | ISYNA1 | inositol-3-phosphate synthase 1 | 2.81E-05 | 4.18 |
| TC0800012076.hg.1 | ARC | activity-regulated cytoskeleton-associated protein | 5.99E-03 | 4.14 |
| TC0400012078.hg.1 | NR3C2 | nuclear receptor subfamily 3, group C, member 2 | 5.37E-04 | 4.12 |

| | | | | |
|---|---|---|---|---|
| TC0100010609.hg.1 | DNM3 | dynamin 3 | 6.04E-06 | 4.11 |
| TC1400007443.hg.1 | HSPA2 | heat shock 70kDa protein 2 | 1.33E-04 | 4.11 |
| TC1200010598.hg.1 | RND1 | Rho family GTPase 1 | 5.21E-04 | 4.10 |
| TC1000012433.hg.1 | SPAG6 | sperm associated antigen 6 | 1.16E-06 | 4.07 |
| TC0100018464.hg.1 | PPM1J | protein phosphatase, Mg2+/Mn2+ dependent, 1J | 1.72E-06 | 3.96 |
| TC1600011060.hg.1 | COTL1 | coactosin-like F-actin binding protein 1 | 1.06E-06 | 3.94 |
| TC0100018307.hg.1 | ACKR1 | atypical chemokine receptor 1 (Duffy blood group) | 6.99E-07 | 3.91 |
| TC0300009696.hg.1 | VWA5B2 | von Willebrand factor A domain containing 5B2 | 4.20E-06 | 3.89 |
| TC0700009472.hg.1 | EPHB6 | EPH receptor B6 | 6.21E-04 | 3.89 |
| TC1600010239.hg.1 | CBLN1 | cerebellin 1 precursor | 1.01E-03 | 3.87 |
| TC1100009075.hg.1 | NCAM1 | Transcript Identified by AceView, Entrez Gene ID(s) 4684 | 4.08E-03 | 3.85 |
| TC1800008281.hg.1 | TMEM241 | transmembrane protein 241 | 7.57E-07 | 3.84 |
| TC0300009033.hg.1 | PLS1 | plastin 1 | 9.89E-05 | 3.78 |
| TC0800010138.hg.1 | RAB11FIP1 | RAB11 family interacting protein 1 (class I) | 4.77E-05 | 3.78 |
| TC0200013021.hg.1 | PAIP2B | poly(A) binding protein interacting protein 2B | 2.45E-06 | 3.76 |
| TC1900008287.hg.1 | PVRL2 | poliovirus receptor-related 2 (herpesvirus entry mediator B) | 2.26E-06 | 3.68 |
| TC1100013045.hg.1 | SIPA1 | signal-induced proliferation-associated 1 | 3.28E-05 | 3.67 |
| TC1200011207.hg.1 | CPM | carboxypeptidase M | 9.44E-04 | 3.66 |
| TC0X00009025.hg.1 | ANOS1 | anosmin 1 | 7.82E-05 | 3.66 |
| TC1600007147.hg.1 | TMEM159 | transmembrane protein 159 | 6.32E-07 | 3.65 |
| TC0800010224.hg.1 | ZMAT4 | zinc finger, matrin-type 4 | 6.88E-04 | 3.64 |
| TC0800011861.hg.1 | LRRC6 | leucine rich repeat containing 6 | 1.02E-05 | 3.64 |
| TC2100008508.hg.1 | CHAF1B | chromatin assembly factor 1, subunit B (p60) | 4.71E-04 | 3.63 |
| TC0X00010382.hg.1 | BEX5 | brain expressed X-linked 5 | 1.54E-03 | 3.61 |
| TC1900010375.hg.1 | RHPN2 | rhophilin, Rho GTPase binding protein 2 | 2.44E-05 | 3.58 |
| TC0100018519.hg.1 | F11R | F11 receptor | 2.83E-04 | 3.57 |
| TSUnmapped00000108.hg.1 | RPS6KA1 | ribosomal protein S6 kinase, 90kDa, polypeptide 1 | 2.18E-06 | 3.56 |
| TC0400011973.hg.1 | CLGN | calmegin | 1.54E-06 | 3.54 |
| TC0900012173.hg.1 | GARNL3 | GTPase activating Rap/RanGAP domain-like 3 | 1.37E-06 | 3.53 |
| TC1200009800.hg.1 | SLC2A3 | solute carrier family 2 (facilitated glucose transporter), member 3 | 4.16E-04 | 3.50 |
| 23076546 | NA | NA | 5.93E-04 | 3.46 |
| TC0300008086.hg.1 | ARL6 | ADP-ribosylation factor like GTPase 6 | 1.99E-05 | 3.46 |
| TC0X00009576.hg.1 | SYN1 | synapsin I | 1.06E-05 | 3.45 |
| TC0400012288.hg.1 | FSTL5 | follistatin-like 5 | 1.23E-03 | 3.45 |
| TC0200006831.hg.1 | VSNL1 | visinin like 1 | 2.05E-03 | 3.44 |
| TC1600007448.hg.1 | CORO1A | coronin, actin binding protein, 1A | 7.49E-06 | 3.44 |
| TC1200009796.hg.1 | SLC2A14 | solute carrier family 2 (facilitated glucose transporter), member 14 | 4.13E-04 | 3.43 |
| TC0100007469.hg.1 | RPS6KA1 | ribosomal protein S6 kinase, 90kDa, polypeptide 1 | 3.19E-06 | 3.42 |
| TC0300014084.hg.1 | CRYGS | crystallin gamma S | 2.51E-05 | 3.41 |
| TC0800007051.hg.1 | NEFM | neurofilament, medium polypeptide | 8.73E-05 | 3.41 |
| TC1900007173.hg.1 | ADGRE5 | adhesion G protein-coupled receptor E5 | 2.36E-04 | 3.40 |
| TC0100010855.hg.1 | NPL | N-acetylneuraminate pyruvate lyase (dihydrodipicolinate synthase) | 5.47E-05 | 3.40 |
| TC0200006671.hg.1 | GRHL1 | grainyhead-like transcription factor 1 | 7.22E-05 | 3.38 |
| TC0400010519.hg.1 | APBB2 | amyloid beta (A4) precursor protein-binding, family B, member 2 | 3.47E-05 | 3.38 |
| TC1200007804.hg.1 | METTL7B | methyltransferase like 7B | 5.77E-03 | 3.35 |
| TC0100007584.hg.1 | PTPRU | protein tyrosine phosphatase, receptor type, U | 3.00E-06 | 3.34 |
| TC0400009958.hg.1 | ABLIM2 | actin binding LIM protein family, member 2 | 9.88E-06 | 3.32 |
| TC1100012722.hg.1 | CDON | cell adhesion associated, oncogene regulated | 5.92E-04 | 3.30 |
| TC0X00006799.hg.1 | SAT1 | spermidine/spermine N1-acetyltransferase 1 | 6.37E-04 | 3.30 |
| TC1400010071.hg.1 | CLMN | calmin (calponin-like, transmembrane) | 6.30E-07 | 3.30 |
| TC0600014220.hg.1 | SERPINB9 | serpin peptidase inhibitor, clade B (ovalbumin), member 9 | 2.17E-04 | 3.29 |
| TC1500010942.hg.1 | LYSMD4 | LysM, putative peptidoglycan-binding, domain containing 4 | 2.65E-05 | 3.27 |
| TC0200014703.hg.1 | CCDC148 | coiled-coil domain containing 148 | 1.32E-06 | 3.26 |
| TC0300006477.hg.1 | ITPR1 | inositol 1,4,5-trisphosphate receptor, type 1 | 3.33E-04 | 3.26 |
| TC1900006603.hg.1 | ZNF556 | zinc finger protein 556 | 1.03E-05 | 3.25 |
| TC0200009078.hg.1 | EPB41L5 | erythrocyte membrane protein band 4.1 like 5 | 5.78E-05 | 3.23 |
| TC1200008667.hg.1 | HSP90B1 | Transcript Identified by AceView, Entrez Gene ID(s) 7184 | 6.37E-06 | 3.23 |
| TC0100015401.hg.1 | IGSF3 | immunoglobulin superfamily, member 3 | 2.07E-05 | 3.23 |
| TC0100010674.hg.1 | GPR52 | G protein-coupled receptor 52 | 2.36E-04 | 3.21 |
| TC0400009088.hg.1 | GUCY1B3 | guanylate cyclase 1, soluble, beta 3 | 2.01E-05 | 3.17 |
| TC0200007399.hg.1 | PLEKHH2 | pleckstrin homology domain containing, family H (with MyTH4 domain) member 2 | 3.21E-06 | 3.17 |
| TC1300010007.hg.1 | TMEM255B | transmembrane protein 255B | 2.84E-05 | 3.17 |
| TC0300009310.hg.1 | MLF1 | myeloid leukemia factor 1 | 3.10E-06 | 3.17 |
| TC0600013165.hg.1 | EPB41L2 | erythrocyte membrane protein band 4.1-like 2 | 9.56E-07 | 3.17 |
| TC1100010123.hg.1 | DKK3 | dickkopf WNT signaling pathway inhibitor 3 | 3.26E-04 | 3.17 |
| TC0300013520.hg.1 | CLDN1 | claudin 1 | 5.83E-05 | 3.16 |
| TC0100018306.hg.1 | CADM3 | cell adhesion molecule 3 | 5.34E-07 | 3.15 |
| TC0600014055.hg.1 | NQO2 | NAD(P)H dehydrogenase, quinone 2 | 1.05E-03 | 3.12 |
| TC0100008779.hg.1 | ST6GALNAC3 | ST6 (alpha-N-acetyl-neuraminyl-2,3-beta-galactosyl-1,3)-N-acetylgalactosaminide alpha-2,6-sialyltransferase 3 | 1.64E-05 | 3.11 |
| TC0100017844.hg.1 | NID1 | nidogen 1 | 5.53E-04 | 3.11 |
| TC1300007752.hg.1 | CLDN10 | claudin 10 | 7.80E-05 | 3.10 |
| TC0200007458.hg.1 | EPAS1 | endothelial PAS domain protein 1 | 1.72E-04 | 3.09 |
| TC0200015266.hg.1 | TMEFF2 | transmembrane protein with EGF-like and two follistatin-like domains 2 | 1.59E-06 | 3.08 |
| TSUnmapped00000390.hg.1 | RPS6KA1 | ribosomal protein S6 kinase, 90kDa, polypeptide 1 | 8.41E-06 | 3.05 |
| TC0500013261.hg.1 | GALNT10 | polypeptide N-acetylgalactosaminyltransferase 10 | 2.71E-03 | 3.04 |
| TC0500013280.hg.1 | ZDHHC11B | zinc finger, DHHC-type containing 11B | 1.54E-04 | 3.02 |
| TC0400012821.hg.1 | C4orf22 | chromosome 4 open reading frame 22 | 2.67E-04 | 3.02 |
| TC0100018403.hg.1 | ERRFI1 | ERBB receptor feedback inhibitor 1 | 2.37E-04 | 3.01 |
| TC0100014191.hg.1 | RAB3B | RAB3B, member RAS oncogene family | 3.49E-04 | 3.01 |
| TC0100010720.hg.1 | BRINP2 | bone morphogenetic protein/retinoic acid inducible neural-specific 2 | 1.75E-05 | 3.00 |
| TC1600008607.hg.1 | PLCG2 | phospholipase C, gamma 2 (phosphatidylinositol-specific) | 5.11E-03 | 3.00 |
| TC0500013089.hg.1 | GFPT2 | glutamine-fructose-6-phosphate transaminase 2 | 9.61E-04 | 2.99 |
| TC0300012392.hg.1 | EFCAB12 | EF-hand calcium binding domain 12 | 1.03E-03 | 2.97 |
| TC1000008715.hg.1 | ELOVL3 | ELOVL fatty acid elongase 3 | 5.02E-05 | 2.96 |
| TC0100016121.hg.1 | KCNJ10 | potassium channel, inwardly rectifying subfamily J, member 10 | 1.82E-05 | 2.95 |
| TC0800010685.hg.1 | MYBL1 | v-myb avian myeloblastosis viral oncogene homolog-like 1 | 5.57E-05 | 2.95 |
| TC1700008262.hg.1 | CACNA1G | calcium channel, voltage-dependent, T type, alpha 1G subunit | 2.96E-06 | 2.94 |
| TC0100009101.hg.1 | ABCD3 | Transcript Identified by AceView, Entrez Gene ID(s) 5825 | 2.61E-04 | 2.94 |
| TC1300008824.hg.1 | SLC25A30 | solute carrier family 25, member 30 | 5.59E-04 | 2.92 |
| TC0300009855.hg.1 | IL1RAP | interleukin 1 receptor accessory protein | 7.02E-05 | 2.91 |
| TC0900011938.hg.1 | C9orf116 | chromosome 9 open reading frame 116 | 8.56E-05 | 2.91 |
| TC1200011993.hg.1 | SLC8B1 | solute carrier family 8 (sodium/lithium/calcium exchanger), member B1 | 4.44E-05 | 2.90 |
| TC0200011040.hg.1 | ITM2C | integral membrane protein 2C | 1.01E-04 | 2.90 |
| TC2200007952.hg.1 | CLTCL1 | clathrin, heavy chain-like 1 | 3.40E-06 | 2.89 |
| TC0100015707.hg.1 | HIST2H4A | histone cluster 2, H4a | 2.37E-06 | 2.89 |
| TC0700006795.hg.1 | AHR | aryl hydrocarbon receptor | 6.51E-03 | 2.88 |
| TC0100009870.hg.1 | HIST2H4B | histone cluster 2, H4b | 2.61E-06 | 2.86 |
| TC0X00009637.hg.1 | PIM2 | Pim-2 proto-oncogene, serine/threonine kinase | 6.69E-05 | 2.85 |
| TC1900010593.hg.1 | DPF1 | D4, zinc and double PHD fingers family 1 | 3.02E-05 | 2.85 |
| TC0400006685.hg.1 | WFS1 | Wolfram syndrome 1 (wolframin) | 3.02E-04 | 2.85 |
| TC0400012640.hg.1 | CCDC110 | coiled-coil domain containing 110 | 2.23E-04 | 2.85 |
| TC0500013282.hg.1 | ZDHHC11 | zinc finger, DHHC-type containing 11 | 5.11E-05 | 2.84 |
| TC0X00006715.hg.1 | SCML1 | sex comb on midleg-like 1 (Drosophila) | 4.32E-04 | 2.84 |
| TC1500007977.hg.1 | HYKK | hydroxylysine kinase | 9.66E-05 | 2.82 |
| TC2000009218.hg.1 | SDC4 | syndecan 4 | 4.76E-04 | 2.82 |
| TC0700013224.hg.1 | LMBR1 | Transcript Identified by AceView, Entrez Gene ID(s) 64327 | 6.14E-05 | 2.81 |
| TC1200010559.hg.1 | VDR | vitamin D (1,25- dihydroxyvitamin D3) receptor | 8.32E-04 | 2.81 |
| TC0800011597.hg.1 | EXT1 | Jeck2013 ALT_ACCEPTOR, ALT_DONOR, coding, INTERNAL, intronic best transcript NM_000127 | 1.40E-04 | 2.81 |
| TC0600009819.hg.1 | ULBP2 | UL16 binding protein 2 | 1.13E-04 | 2.80 |
| TC0100006787.hg.1 | PIK3CD | phosphatidylinositol-4,5-bisphosphate 3-kinase, catalytic subunit delta | 4.88E-05 | 2.80 |
| TC0200007048.hg.1 | MAPRE3 | microtubule-associated protein, RP/EB family, member 3 | 4.12E-06 | 2.80 |
| TC1900010782.hg.1 | ATP1A3 | ATPase, Na+/K+ transporting, alpha 3 polypeptide | 7.60E-06 | 2.78 |
| TC1000008891.hg.1 | DUSP5 | dual specificity phosphatase 5 | 2.51E-04 | 2.78 |
| TC1900007040.hg.1 | CNN1 | calponin 1, basic, smooth muscle | 1.51E-05 | 2.78 |
| TC0900007006.hg.1 | PRSS3 | protease, serine, 3 | 4.85E-03 | 2.77 |
| TC0100016973.hg.1 | CYB5R1 | cytochrome b5 reductase 1 | 4.83E-04 | 2.77 |
| TC1900009155.hg.1 | GAMT | guanidinoacetate N-methyltransferase | 2.63E-05 | 2.76 |
| TC0400010618.hg.1 | TEC | tec protein tyrosine kinase | 1.73E-05 | 2.76 |
| TC0600014151.hg.1 | SMIM8 | small integral membrane protein 8 | 5.52E-04 | 2.73 |

| | | | | |
|---|---|---|---|---|
| TC1900007426.hg.1 | NCAN | neurocan | 1.02E-03 | 2.73 |
| TC0700010899.hg.1 | POLR2J4 | polymerase (RNA) II (DNA directed) polypeptide J4, pseudogene | 2.57E-05 | 2.73 |
| TC0300013719.hg.1 | CEP19 | centrosomal protein 19kDa | 1.88E-04 | 2.73 |
| 23076588 | NA | NA | 4.21E-04 | 2.71 |
| TC0800009212.hg.1 | MAPK15 | mitogen-activated protein kinase 15 | 4.21E-04 | 2.71 |
| TC0100013076.hg.1 | PADI2 | peptidyl arginine deiminase, type II | 4.04E-05 | 2.70 |
| TC0200009662.hg.1 | TNFAIP6 | tumor necrosis factor, alpha-induced protein 6 | 9.48E-03 | 2.70 |
| TC0600007262.hg.1 | HIST1H3A | histone cluster 1, H3a | 9.23E-04 | 2.69 |
| TSUnmapped00000572.hg.1 | RPS6KA1 | ribosomal protein S6 kinase, 90kDa, polypeptide 1 | 6.93E-06 | 2.69 |
| TC0200016571.hg.1 | OSBPL6 | oxysterol binding protein-like 6 | 4.83E-05 | 2.68 |
| TC1300006919.hg.1 | FREM2 | FRAS1 related extracellular matrix protein 2 | 4.95E-06 | 2.68 |
| TC1600009246.hg.1 | ROGDI | rogdi homolog | 5.23E-06 | 2.67 |
| TC0100011378.hg.1 | RASSF5 | Ras association (RalGDS/AF-6) domain family member 5 | 1.76E-04 | 2.67 |
| TC0200013602.hg.1 | TSGA10 | testis specific 10 | 1.16E-04 | 2.67 |
| TC0700008261.hg.1 | RUNDC3B | RUN domain containing 3B | 3.09E-05 | 2.67 |
| TC1200012111.hg.1 | TAOK3 | TAO kinase 3 | 1.05E-03 | 2.67 |
| TC1500010941.hg.1 | LYSMD4 | LysM, putative peptidoglycan-binding, domain containing 4 | 1.57E-03 | 2.67 |
| TC1700011436.hg.1 | ERN1 | endoplasmic reticulum to nucleus signaling 1 | 7.18E-04 | 2.66 |
| TC0200009829.hg.1 | GCA | grancalcin, EF-hand calcium binding protein | 1.08E-05 | 2.65 |
| TC0X00006723.hg.1 | CDKL5 | cyclin-dependent kinase-like 5 | 2.16E-03 | 2.65 |
| TC1200010612.hg.1 | RHEBL1 | Ras homolog enriched in brain like 1 | 6.48E-04 | 2.65 |
| TC1000007176.hg.1 | MAP3K8 | mitogen-activated protein kinase kinase kinase 8 | 8.54E-04 | 2.65 |
| TC1500008120.hg.1 | SH3GL3 | SH3-domain GRB2-like 3 | 1.39E-04 | 2.64 |
| TC0300007973.hg.1 | CADM2 | cell adhesion molecule 2 | 1.71E-04 | 2.64 |
| TC0600014362.hg.1 | TFB1M | transcription factor B1, mitochondrial | 3.11E-03 | 2.64 |
| TC1100011282.hg.1 | CD248 | CD248 molecule, endosialin | 6.78E-05 | 2.64 |
| TC0400008725.hg.1 | PCDH10 | protocadherin 10 | 4.84E-03 | 2.64 |
| TC0900011370.hg.1 | BRINP1 | bone morphogenetic protein/retinoic acid inducible neural-specific 1 | 4.60E-04 | 2.63 |
| TC1900007688.hg.1 | CCNE1 | cyclin E1 | 1.86E-05 | 2.63 |
| TC0100015102.hg.1 | COL11A1 | collagen, type XI, alpha 1 | 2.97E-04 | 2.63 |
| TC0200013298.hg.1 | ST3GAL5 | ST3 beta-galactoside alpha-2,3-sialyltransferase 5 | 1.88E-04 | 2.62 |
| TC1100008541.hg.1 | AAMDC | adipogenesis associated, Mth938 domain containing | 5.06E-05 | 2.62 |
| TC0100018418.hg.1 | IFFO2 | intermediate filament family orphan 2 | 3.40E-05 | 2.61 |
| TC0600011133.hg.1 | HIST1H2BE | Memczak2013 ANTISENSE, CDS, coding, upstream_start, UTR3, UTR5 best transcript NM_003523 | 3.55E-03 | 2.60 |
| TC0600007610.hg.1 | MSH5 | mutS homolog 5 | 1.94E-03 | 2.60 |
| TC2200007318.hg.1 | H1F0 | H1 histone family, member 0 | 7.34E-05 | 2.59 |
| TC1100007394.hg.1 | CD82 | CD82 molecule | 3.89E-04 | 2.59 |
| TC1200007359.hg.1 | CNTN1 | contactin 1 | 7.69E-04 | 2.59 |
| TC1600006525.hg.1 | CACNA1H | calcium channel, voltage-dependent, T type, alpha 1H subunit | 3.24E-04 | 2.59 |
| TC0100015803.hg.1 | TDRKH | tudor and KH domain containing | 5.20E-04 | 2.58 |
| TC0100014127.hg.1 | SPATA6 | spermatogenesis associated 6 | 1.96E-04 | 2.58 |
| TC1200007809.hg.1 | GDF11 | growth differentiation factor 11 | 2.14E-05 | 2.58 |
| TC1500010744.hg.1 | GCOM1 | GRINL1A complex locus 1 | 1.14E-05 | 2.57 |
| TC0300008853.hg.1 | TMEM108 | transmembrane protein 108 | 2.42E-05 | 2.57 |
| TC1200010413.hg.1 | ABCD2 | ATP binding cassette subfamily D member 2 | 3.38E-04 | 2.57 |
| TC0200011837.hg.1 | FAM49A | family with sequence similarity 49, member A | 3.33E-04 | 2.57 |
| TC1900007127.hg.1 | IER2 | immediate early response 2 | 2.80E-06 | 2.57 |
| TSUnmapped00000109.hg.1 | ATG16L1 | autophagy related 16-like 1 | 2.53E-04 | 2.57 |
| TC2000009023.hg.1 | SAMHD1 | SAM domain and HD domain 1 | 4.64E-05 | 2.56 |
| TC0100018466.hg.1 | DENND2C | DENN/MADD domain containing 2C | 7.74E-03 | 2.56 |
| TC1400009184.hg.1 | TXNDC16 | thioredoxin domain containing 16 | 7.92E-06 | 2.56 |
| TC1600008737.hg.1 | FLJ30679 | uncharacterized protein FLJ30679 | 5.84E-03 | 2.56 |
| TC0400008106.hg.1 | HERC5 | HECT and RLD domain containing E3 ubiquitin protein ligase 5 | 2.41E-03 | 2.56 |
| TC1900009670.hg.1 | DOCK6 | dedicator of cytokinesis 6 | 7.54E-05 | 2.55 |
| TC1700011060.hg.1 | FAM117A | family with sequence similarity 117, member A | 4.15E-05 | 2.54 |
| TC2100006585.hg.1 | RBM11 | RNA binding motif protein 11 | 2.16E-05 | 2.54 |
| TC2200007987.hg.1 | CLDN5 | claudin 5 | 1.51E-04 | 2.54 |
| TC0600012815.hg.1 | PPIL6 | peptidylprolyl isomerase (cyclophilin)-like 6 | 1.20E-04 | 2.53 |
| TC1900006507.hg.1 | CNN2 | calponin 2 | 9.01E-05 | 2.53 |
| TC0600014371.hg.1 | RNASET2 | ribonuclease T2 | 3.07E-04 | 2.53 |
| 23075523 | NA | NA | 1.93E-03 | 2.53 |
| 23075806 | NA | NA | 1.93E-03 | 2.53 |
| 23076508 | NA | NA | 1.93E-03 | 2.53 |
| TC0100016649.hg.1 | NMNAT2 | nicotinamide nucleotide adenylyltransferase 2 | 2.45E-03 | 2.53 |
| TC2000008242.hg.1 | RNF24 | ring finger protein 24 | 1.39E-04 | 2.53 |
| TC0500007549.hg.1 | ZSWIM6 | zinc finger, SWIM-type containing 6 | 4.30E-04 | 2.52 |
| TC1900009369.hg.1 | PLIN3 | perilipin 3 | 4.72E-04 | 2.52 |
| TC0200008870.hg.1 | BCL2L11 | BCL2-like 11 (apoptosis facilitator) | 4.96E-04 | 2.51 |
| TC1100006899.hg.1 | ARNTL | aryl hydrocarbon receptor nuclear translocator-like | 1.12E-03 | 2.50 |
| TC0800007311.hg.1 | ADGRA2 | adhesion G protein-coupled receptor A2 | 7.02E-06 | 2.49 |
| TSUnmapped00000435.hg.1 | RPS6KA1 | ribosomal protein S6 kinase, 90kDa, polypeptide 1 | 7.48E-05 | 2.49 |
| HTA2-neg-47419722_st | NA | NA | 4.96E-04 | 2.49 |
| TC1100011192.hg.1 | ATG2A | autophagy related 2A | 6.83E-04 | 2.49 |
| 23075724 | NA | NA | 6.16E-03 | 2.48 |
| TC0100015925.hg.1 | KCNN3 | potassium channel, calcium activated intermediate/small conductance subfamily N alpha, member 3 | 1.24E-03 | 2.48 |
| TC1600010603.hg.1 | RRAD | Ras-related associated with diabetes | 4.08E-03 | 2.48 |
| TC0700009680.hg.1 | TMEM176A | transmembrane protein 176A | 2.27E-04 | 2.48 |
| TC0200007954.hg.1 | PCYOX1 | prenylcysteine oxidase 1 | 5.04E-05 | 2.48 |
| TC1900008328.hg.1 | PPM1N | protein phosphatase, Mg2+/Mn2+ dependent, 1N (putative) | 1.89E-03 | 2.48 |
| TC0900008197.hg.1 | GALNT12 | polypeptide N-acetylgalactosaminyltransferase 12 | 1.04E-05 | 2.47 |
| TC1100009242.hg.1 | ABCG4 | ATP binding cassette subfamily G member 4 | 8.66E-05 | 2.47 |
| TC0600011943.hg.1 | ENPP5 | ectonucleotide pyrophosphatase/phosphodiesterase 5 (putative) | 5.24E-04 | 2.47 |
| TC1900011892.hg.1 | RTBDN | retbindin | 2.70E-04 | 2.47 |
| TC1100012019.hg.1 | SESN3 | sestrin 3 | 4.19E-03 | 2.47 |
| TC0100013205.hg.1 | ECE1 | endothelin converting enzyme 1 | 5.39E-05 | 2.47 |
| TC1500008029.hg.1 | FAH | fumarylacetoacetate hydrolase (fumarylacetoacetase) | 5.26E-04 | 2.46 |
| TC2000008268.hg.1 | SLC23A2 | solute carrier family 23 (ascorbic acid transporter), member 2 | 2.45E-04 | 2.46 |
| TC2100006659.hg.1 | CXADR | coxsackie virus and adenovirus receptor | 9.67E-05 | 2.46 |
| 23075965 | NA | NA | 1.64E-03 | 2.46 |
| TC0600007012.hg.1 | CD83 | CD83 molecule | 5.16E-04 | 2.46 |
| TC1500010909.hg.1 | STARD5 | StAR-related lipid transfer domain containing 5 | 1.23E-05 | 2.46 |
| TC0200007053.hg.1 | KHK | ketohexokinase | 1.41E-05 | 2.45 |
| TC1900008826.hg.1 | CACNG7 | calcium channel, voltage-dependent, gamma subunit 7 | 1.56E-04 | 2.45 |
| TC1900010360.hg.1 | ANKRD27 | ankyrin repeat domain 27 (VPS9 domain) | 8.41E-04 | 2.45 |
| TC1000012536.hg.1 | MPP7 | membrane protein, palmitoylated 7 | 9.84E-05 | 2.45 |
| TC0300009916.hg.1 | HES1 | hes family bHLH transcription factor 1 | 3.34E-04 | 2.44 |
| TC0800009194.hg.1 | RHPN1 | rhophilin, Rho GTPase binding protein 1 | 3.02E-04 | 2.43 |
| TC0600013709.hg.1 | SERAC1 | serine active site containing 1 | 6.74E-04 | 2.43 |
| TC0200010745.hg.1 | IGFBP2 | insulin like growth factor binding protein 2 | 5.48E-04 | 2.43 |
| TC1100013054.hg.1 | RAD9A | RAD9 checkpoint clamp component A | 2.15E-04 | 2.43 |
| TC0200013595.hg.1 | MGAT4A | mannosyl (alpha-1,3-)-glycoprotein beta-1,4-N-acetylglucosaminyltransferase, isozyme A | 4.58E-04 | 2.43 |
| TC2200007150.hg.1 | TIMP3 | TIMP metallopeptidase inhibitor 3 | 7.98E-04 | 2.43 |
| TC2000010026.hg.1 | TMEM189 | transmembrane protein 189 | 6.34E-04 | 2.43 |
| TC0800010506.hg.1 | PLAG1 | pleiomorphic adenoma gene 1 | 3.57E-03 | 2.42 |
| TC0600008166.hg.1 | SLC25A27 | solute carrier family 25, member 27 | 3.78E-04 | 2.42 |
| TC1800007066.hg.1 | MAPRE2 | microtubule-associated protein, RP/EB family, member 2 | 9.14E-04 | 2.42 |
| TC0100009215.hg.1 | CDC14A | cell division cycle 14A | 1.75E-05 | 2.42 |
| 23076189 | NA | NA | 8.05E-03 | 2.41 |
| TC0100009959.hg.1 | TUFT1 | tuftelin 1 | 1.93E-03 | 2.41 |
| TC1900007103.hg.1 | GCDH | glutaryl-CoA dehydrogenase | 1.76E-04 | 2.41 |
| TC1300007707.hg.1 | GPC5 | glypican 5 | 1.70E-04 | 2.41 |
| TC0X00010643.hg.1 | 44080 | septin 6 | 1.16E-04 | 2.40 |
| TC0200012412.hg.1 | C1GALT1C1L | C1GALT1-specific chaperone 1 like | 2.68E-05 | 2.40 |
| TC0900008243.hg.1 | PLPPR1 | phospholipid phosphatase related 1 | 2.05E-03 | 2.39 |
| TC0600013193.hg.1 | STX7 | syntaxin 7 | 2.40E-05 | 2.39 |
| TC0700010504.hg.1 | OSBPL3 | oxysterol binding protein-like 3 | 1.02E-04 | 2.39 |
| TC0300014085.hg.1 | TBCCD1 | TBCC domain containing 1 | 2.27E-05 | 2.38 |
| TC0100006571.hg.1 | SKI | Memczak2013 ALT_ACCEPTOR, ALT_DONOR, coding, INTERNAL, intronic best transcript NM_003036 | 5.47E-05 | 2.38 |
| TC0300006961.hg.1 | CMTM8 | CKLF-like MARVEL transmembrane domain containing 8 | 5.30E-05 | 2.38 |

| | | | | |
|---|---|---|---|---|
| TC1700010358.hg.1 | MYO1D | myosin ID | 1.40E-04 | 2.38 |
| TC1600010452.hg.1 | DOK4 | docking protein 4 | 7.19E-05 | 2.38 |
| TC0200015815.hg.1 | EPHA4 | EPH receptor A4 | 3.21E-03 | 2.38 |
| TC1900011844.hg.1 | MFSD12 | major facilitator superfamily domain containing 12 | 2.66E-03 | 2.37 |
| TC0700008252.hg.1 | CROT | carnitine O-octanoyltransferase | 3.50E-03 | 2.37 |
| TC0200011972.hg.1 | C2orf44 | chromosome 2 open reading frame 44 | 1.57E-03 | 2.37 |
| TC1600008661.hg.1 | CRISPLD2 | cysteine-rich secretory protein LCCL domain containing 2 | 1.58E-03 | 2.37 |
| TC0600011234.hg.1 | HIST1H4L | histone cluster 1, H4l | 1.94E-03 | 2.37 |
| TC1300009026.hg.1 | PCDH8 | protocadherin 8 | 3.03E-04 | 2.37 |
| TC1100009374.hg.1 | VWA5A | von Willebrand factor A domain containing 5A | 9.83E-03 | 2.36 |
| TC2000007177.hg.1 | RALY | Jeck2013 ALT_DONOR, coding, INTERNAL, intronic best transcript NM_016732 | 2.39E-04 | 2.36 |
| TC0700013047.hg.1 | TMEM176B | transmembrane protein 176B | 6.07E-05 | 2.36 |
| TC0500012023.hg.1 | KIF3A | kinesin family member 3A | 1.16E-04 | 2.36 |
| TC0900010933.hg.1 | CTSV | cathepsin V | 2.70E-03 | 2.36 |
| TC1900006605.hg.1 | ZNF57 | zinc finger protein 57 | 1.18E-04 | 2.36 |
| TC0600012379.hg.1 | ELOVL4 | ELOVL fatty acid elongase 4 | 3.75E-03 | 2.36 |
| TC0500012842.hg.1 | DUSP1 | dual specificity phosphatase 1 | 3.52E-04 | 2.35 |
| TC0200012140.hg.1 | ALK | anaplastic lymphoma receptor tyrosine kinase | 7.32E-05 | 2.35 |
| TC0300011525.hg.1 | RYBP | RING1 and YY1 binding protein | 2.94E-04 | 2.35 |
| TC0900010969.hg.1 | CORO2A | coronin, actin binding protein, 2A | 4.32E-05 | 2.35 |
| TC1500010776.hg.1 | IDH3A | isocitrate dehydrogenase 3 (NAD+) alpha | 3.43E-04 | 2.35 |
| TC0200015242.hg.1 | STAT1 | signal transducer and activator of transcription 1 | 1.39E-05 | 2.34 |
| TC1000007061.hg.1 | GPR158 | G protein-coupled receptor 158 | 2.51E-03 | 2.34 |
| TC0100008052.hg.1 | CFAP57 | cilia and flagella associated protein 57 | 2.31E-03 | 2.34 |
| TC1700008661.hg.1 | PRKCA | protein kinase C, alpha | 8.04E-05 | 2.34 |
| TC1400010708.hg.1 | TTC5 | tetratricopeptide repeat domain 5 | 1.31E-03 | 2.33 |
| TC1400008116.hg.1 | GSKIP | GSK3B interacting protein | 6.76E-04 | 2.33 |
| TC1300009004.hg.1 | ATP7B | ATPase, Cu++ transporting, beta polypeptide | 1.30E-04 | 2.33 |
| TC1500010841.hg.1 | CHRFAM7A | CHRNA7 (cholinergic receptor, nicotinic, alpha 7, exons 5-10) and FAM7A (family with sequence similarity 7A, exons A-E) fusion | 1.64E-05 | 2.33 |
| TC1700009954.hg.1 | TOM1L2 | target of myb1 like 2 membrane trafficking protein | 1.28E-03 | 2.33 |
| TC1900006588.hg.1 | GADD45B | growth arrest and DNA-damage-inducible, beta | 3.41E-03 | 2.33 |
| TC0800011595.hg.1 | EXT1 | Jeck2013 ALT_ACCEPTOR, ALT_DONOR, coding, INTERNAL, intronic best transcript NM_000127 | 7.56E-03 | 2.33 |
| TC0600009144.hg.1 | KIAA1919 | KIAA1919 | 3.00E-04 | 2.33 |
| TC1300009712.hg.1 | EFNB2 | ephrin-B2 | 4.40E-03 | 2.32 |
| TC0100015155.hg.1 | VAV3 | vav 3 guanine nucleotide exchange factor | 1.76E-04 | 2.32 |
| TC0100017310.hg.1 | ESRRG | estrogen-related receptor gamma | 1.76E-04 | 2.31 |
| TC1100011190.hg.1 | EHD1 | EH domain containing 1 | 8.27E-04 | 2.31 |
| TC0100010874.hg.1 | RGL1 | ral guanine nucleotide dissociation stimulator-like 1 | 7.61E-03 | 2.31 |
| TC2200009257.hg.1 | TCN2 | transcobalamin II | 5.18E-03 | 2.31 |
| TC1400008452.hg.1 | ADSSL1 | adenylosuccinate synthase like 1 | 8.14E-03 | 2.31 |
| TC1200009300.hg.1 | BRI3BP | BRI3 binding protein | 9.52E-04 | 2.31 |
| TC1100011652.hg.1 | GDPD5 | glycerophosphodiester phosphodiesterase domain containing 5 | 1.98E-05 | 2.30 |
| TC1700009402.hg.1 | SMYD4 | SET and MYND domain containing 4 | 1.59E-04 | 2.30 |
| TC0X00008910.hg.1 | ASMTL | acetylserotonin O-methyltransferase-like | 2.40E-05 | 2.30 |
| TC0Y00006884.hg.1 | ASMTL | acetylserotonin O-methyltransferase-like | 2.40E-05 | 2.30 |
| 23067838 | NA | NA | 4.96E-03 | 2.30 |
| TC1400008058.hg.1 | PPP4R4 | protein phosphatase 4, regulatory subunit 4 | 4.96E-03 | 2.30 |
| TC0X00008468.hg.1 | CCDC160 | coiled-coil domain containing 160 | 4.84E-04 | 2.30 |
| TC0100013028.hg.1 | EPHA2 | EPH receptor A2 | 3.58E-04 | 2.29 |
| TC2200007068.hg.1 | SLC35E4 | solute carrier family 35, member E4 | 3.37E-04 | 2.29 |
| TC1900011150.hg.1 | NOSIP | nitric oxide synthase interacting protein | 4.20E-04 | 2.29 |
| TC0200012009.hg.1 | DTNB | dystrobrevin beta | 8.50E-05 | 2.29 |
| TC0100011325.hg.1 | TMCC2 | transmembrane and coiled-coil domain family 2 | 1.60E-05 | 2.29 |
| TC0700012163.hg.1 | EFCAB10 | EF-hand calcium binding domain 10 | 1.23E-04 | 2.29 |
| TC0400007280.hg.1 | WDR19 | WD repeat domain 19 | 9.27E-05 | 2.28 |
| TC0800009094.hg.1 | DENND3 | DENN/MADD domain containing 3 | 1.10E-04 | 2.28 |
| TC0100017241.hg.1 | TMEM206 | transmembrane protein 206 | 1.16E-04 | 2.28 |
| TC2100007999.hg.1 | DNAJC28 | DnaJ (Hsp40) homolog, subfamily C, member 28 | 4.91E-04 | 2.27 |
| TC1100008627.hg.1 | ANKRD42 | ankyrin repeat domain 42 | 4.34E-03 | 2.27 |
| TC1100006820.hg.1 | SWAP70 | SWAP switching B-cell complex 70kDa subunit | 4.81E-05 | 2.27 |
| TC0600013147.hg.1 | ARHGAP18 | Rho GTPase activating protein 18 | 6.74E-05 | 2.27 |
| TC0400008424.hg.1 | C4orf32 | chromosome 4 open reading frame 32 | 7.86E-04 | 2.27 |
| TC0300007596.hg.1 | FLNB | filamin B, beta | 7.87E-05 | 2.27 |
| TC0600009622.hg.1 | CCDC28A | coiled-coil domain containing 28A | 8.94E-04 | 2.27 |
| TC1900009603.hg.1 | OLFM2 | olfactomedin 2 | 1.43E-04 | 2.26 |
| TC0600007274.hg.1 | HIST1H2BD | histone cluster 1, H2bd | 6.37E-05 | 2.26 |
| 23075524 | NA | NA | 1.44E-05 | 2.26 |
| TC0200007535.hg.1 | PPP1R21 | protein phosphatase 1, regulatory subunit 21 | 1.24E-04 | 2.26 |
| TC1100009521.hg.1 | APLP2 | amyloid beta (A4) precursor-like protein 2 | 5.01E-04 | 2.26 |
| 23076191 | NA | NA | 7.84E-03 | 2.25 |
| TC0200016582.hg.1 | NABP1 | nucleic acid binding protein 1 | 4.60E-05 | 2.25 |
| TC0X00009387.hg.1 | RPGR | retinitis pigmentosa GTPase regulator | 3.15E-03 | 2.25 |
| 23076268 | NA | NA | 9.50E-06 | 2.25 |
| TC0200010615.hg.1 | CCNYL1 | cyclin Y like 1 | 5.34E-04 | 2.25 |
| TC0300011540.hg.1 | SHQ1 | SHQ1, H/ACA ribonucleoprotein assembly factor | 3.16E-03 | 2.25 |
| TC1200007653.hg.1 | NR4A1 | nuclear receptor subfamily 4, group A, member 1 | 2.04E-03 | 2.25 |
| TC1000011567.hg.1 | CRTAC1 | cartilage acidic protein 1 | 4.13E-04 | 2.25 |
| TC0600010797.hg.1 | MAK | male germ cell-associated kinase | 5.30E-05 | 2.25 |
| TC1400010012.hg.1 | LGMN | legumain | 4.93E-04 | 2.24 |
| TC1100011367.hg.1 | UNC93B1 | unc-93 homolog B1 (C. elegans) | 5.23E-03 | 2.24 |
| TC1700011439.hg.1 | TEX2 | testis expressed 2 | 1.43E-04 | 2.24 |
| TC1000008056.hg.1 | VCL | vinculin | 4.90E-03 | 2.24 |
| TC0500009824.hg.1 | ZDHHC11 | Transcript Identified by AceView, Entrez Gene ID(s) 79844 | 5.13E-03 | 2.24 |
| TC0500008356.hg.1 | KCNN2 | potassium channel, calcium activated intermediate/small conductance subfamily N alpha, member 2 | 9.29E-04 | 2.24 |
| TC1900009806.hg.1 | PRKACA | protein kinase, cAMP-dependent, catalytic, alpha | 2.58E-04 | 2.24 |
| TC0200013046.hg.1 | EXOC6B | exocyst complex component 6B | 1.15E-05 | 2.23 |
| TC0800011312.hg.1 | RRM2B | ribonucleotide reductase M2 B (TP53 inducible) | 6.59E-04 | 2.23 |
| TC1300009583.hg.1 | DOCK9 | dedicator of cytokinesis 9 | 6.70E-03 | 2.23 |
| TC1700010198.hg.1 | UNC119 | unc-119 lipid binding chaperone | 1.74E-05 | 2.23 |
| 23074629 | NA | NA | 3.83E-04 | 2.23 |
| TC0400009376.hg.1 | WDR17 | WD repeat domain 17 | 3.82E-04 | 2.23 |
| TC0400010558.hg.1 | ATP8A1 | ATPase, aminophospholipid transporter (APLT), class I, type 8A, member 1 | 8.83E-03 | 2.23 |
| TC0200010907.hg.1 | ACSL3 | acyl-CoA synthetase long-chain family member 3 | 1.07E-04 | 2.23 |
| TC0100018210.hg.1 | SH3D21 | SH3 domain containing 21 | 3.56E-04 | 2.23 |
| TC0500013200.hg.1 | RFESD | Rieske (Fe-S) domain containing | 3.74E-03 | 2.23 |
| TC0X00006593.hg.1 | CLCN4 | chloride channel, voltage-sensitive 4 | 5.72E-05 | 2.22 |
| TC1200008574.hg.1 | APAF1 | apoptotic peptidase activating factor 1 | 1.29E-04 | 2.22 |
| TC1900006819.hg.1 | TRIP10 | thyroid hormone receptor interactor 10 | 2.06E-04 | 2.22 |
| TC0500012282.hg.1 | HDAC3 | histone deacetylase 3 | 1.25E-04 | 2.22 |
| TC0700009079.hg.1 | TSPAN33 | tetraspanin 33 | 4.83E-05 | 2.21 |
| TC1000009092.hg.1 | INPP5F | inositol polyphosphate-5-phosphatase F | 1.53E-03 | 2.21 |
| TC1900011139.hg.1 | SLC17A7 | solute carrier family 17 (vesicular glutamate transporter), member 7 | 2.02E-03 | 2.21 |
| HTA2-pos-2985890_st | NA | NA | 3.54E-04 | 2.21 |
| 23076478 | NA | NA | 4.81E-03 | 2.21 |
| TC0800007416.hg.1 | GINS4 | GINS complex subunit 4 (Sld5 homolog) | 5.24E-05 | 2.21 |
| TC1100007943.hg.1 | PLCB3 | phospholipase C, beta 3 (phosphatidylinositol-specific) | 4.25E-05 | 2.21 |
| 23075555 | NA | NA | 4.86E-03 | 2.20 |
| 23075693 | NA | NA | 4.86E-03 | 2.20 |
| 23075866 | NA | NA | 4.86E-03 | 2.20 |
| 23076012 | NA | NA | 4.86E-03 | 2.20 |
| 23076157 | NA | NA | 4.86E-03 | 2.20 |
| 23076325 | NA | NA | 4.86E-03 | 2.20 |
| TC1900011866.hg.1 | RAB3D | RAB3D, member RAS oncogene family | 2.39E-04 | 2.20 |
| TC0600014152.hg.1 | LINC01590 | long intergenic non-protein coding RNA 1590 | 5.92E-04 | 2.20 |
| TC0300008375.hg.1 | ZDHHC23 | zinc finger, DHHC-type containing 23 | 2.50E-04 | 2.20 |
| TC0300013763.hg.1 | RUBCN | RUN domain and cysteine-rich domain containing, Beclin 1-interacting protein | 1.64E-04 | 2.20 |
| TC1100007026.hg.1 | ZDHHC13 | zinc finger, DHHC-type containing 13 | 2.45E-03 | 2.20 |
| TC1600008712.hg.1 | IRF8 | interferon regulatory factor 8 | 4.52E-03 | 2.19 |
| TC0800007269.hg.1 | UNC5D | unc-5 netrin receptor D | 9.18E-04 | 2.19 |

| | | | | |
|---|---|---|---|---|
| TC0300014082.hg.1 | ETV5 | ets variant 5 | 6.18E-04 | 2.19 |
| HTA2-pos-2985912_st | NA | NA | 5.33E-03 | 2.18 |
| TC0400008686.hg.1 | C4orf33 | chromosome 4 open reading frame 33 | 1.68E-03 | 2.18 |
| TC0600013537.hg.1 | RAET1G | retinoic acid early transcript 1G | 3.95E-05 | 2.18 |
| 23066337 | NA | NA | 6.39E-04 | 2.18 |
| TC0100017500.hg.1 | TMEM63A | transmembrane protein 63A | 2.44E-03 | 2.18 |
| TC0100007295.hg.1 | EPHB2 | EPH receptor B2 | 2.24E-03 | 2.17 |
| TC0500011758.hg.1 | CDO1 | cysteine dioxygenase type 1 | 2.51E-04 | 2.17 |
| TC0200016530.hg.1 | PLEKHB2 | pleckstrin homology domain containing, family B (evectins) member 2 | 4.90E-04 | 2.17 |
| TC1400010629.hg.1 | FLVCR2 | feline leukemia virus subgroup C cellular receptor family, member 2 | 3.89E-03 | 2.17 |
| TC0700008552.hg.1 | AGFG2 | ArfGAP with FG repeats 2 | 4.12E-04 | 2.17 |
| TC0500010731.hg.1 | ARL15 | ADP-ribosylation factor like GTPase 15 | 1.86E-04 | 2.16 |
| TC0700013223.hg.1 | LMBR1 | Transcript Identified by AceView, Entrez Gene ID(s) 64327 | 8.54E-03 | 2.16 |
| TC1100009707.hg.1 | POLR2L | polymerase (RNA) II (DNA directed) polypeptide L, 7.6kDa | 9.40E-05 | 2.16 |
| 23075726 | NA | NA | 5.60E-03 | 2.16 |
| TC0900007047.hg.1 | DNAI1 | dynein, axonemal, intermediate chain 1 | 1.92E-04 | 2.16 |
| TC0300010821.hg.1 | ULK4 | unc-51 like kinase 4 | 3.07E-04 | 2.16 |
| TC0700011784.hg.1 | FAM133B | Transcript Identified by AceView, Entrez Gene ID(s) 257415 | 8.17E-04 | 2.16 |
| TC1300009674.hg.1 | KDELC1 | KDEL (Lys-Asp-Glu-Leu) containing 1 | 3.48E-04 | 2.15 |
| TC0500006816.hg.1 | FAM105A | family with sequence similarity 105, member A | 7.76E-05 | 2.15 |
| TC0100017849.hg.1 | ERO1B | endoplasmic reticulum oxidoreductase beta | 1.37E-04 | 2.15 |
| TC0X00007251.hg.1 | CCNB3 | cyclin B3 | 3.41E-03 | 2.15 |
| TC0100014009.hg.1 | TESK2 | testis-specific kinase 2 | 2.22E-04 | 2.15 |
| TSUnmapped00000262.hg.1 | MLXIP | MLX interacting protein | 2.80E-04 | 2.14 |
| TC0600007060.hg.1 | MYLIP | myosin regulatory light chain interacting protein | 5.21E-03 | 2.14 |
| TC0900007604.hg.1 | OSTF1 | osteoclast stimulating factor 1 | 1.21E-04 | 2.14 |
| TC0100007678.hg.1 | HDAC1 | histone deacetylase 1 | 4.12E-05 | 2.14 |
| TC0600011463.hg.1 | NEU1 | sialidase 1 (lysosomal sialidase) | 4.11E-03 | 2.14 |
| TC0700013394.hg.1 | CCDC146 | coiled-coil domain containing 146 | 6.09E-03 | 2.14 |
| TC0X00008836.hg.1 | PLXNA3 | plexin A3 | 8.25E-05 | 2.14 |
| TC0200016494.hg.1 | CNNM4 | cyclin and CBS domain divalent metal cation transport mediator 4 | 6.99E-03 | 2.14 |
| 23075635 | NA | NA | 5.16E-05 | 2.14 |
| TC0700006781.hg.1 | BZW2 | basic leucine zipper and W2 domains 2 | 2.92E-03 | 2.14 |
| TC2200009352.hg.1 | LOC400927 | TPTE and PTEN homologous inositol lipid phosphatase pseudogene | 3.17E-03 | 2.13 |
| TC0700011463.hg.1 | TYW1B | tRNA-yW synthesizing protein 1 homolog B (S. cerevisiae) | 4.08E-04 | 2.13 |
| TC1700012025.hg.1 | CEP131 | centrosomal protein 131kDa | 3.09E-04 | 2.13 |
| TC1100008085.hg.1 | PELI3 | pellino E3 ubiquitin protein ligase family member 3 | 5.43E-05 | 2.13 |
| TC0700011556.hg.1 | HIP1 | huntingtin interacting protein 1 | 3.45E-04 | 2.13 |
| TC0200012936.hg.1 | FBXO48 | F-box protein 48 | 5.96E-03 | 2.13 |
| TC1700009042.hg.1 | AFMID | arylformamidase | 1.01E-03 | 2.13 |
| TC0100012089.hg.1 | GPR137B | G protein-coupled receptor 137B | 1.02E-03 | 2.13 |
| TC0800010379.hg.1 | EFCAB1 | EF-hand calcium binding domain 1 | 1.10E-03 | 2.13 |
| TC1200010415.hg.1 | SLC2A13 | solute carrier family 2 (facilitated glucose transporter), member 13 | 1.15E-04 | 2.13 |
| HTA2-pos-47421984_st | NA | NA | 1.98E-03 | 2.12 |
| 23076565 | NA | NA | 3.30E-04 | 2.12 |
| 23076624 | NA | NA | 3.36E-03 | 2.12 |
| TC1600011574.hg.1 | GINS2 | GINS complex subunit 2 (Psf2 homolog) | 1.61E-03 | 2.12 |
| TC2000008885.hg.1 | SNTA1 | syntrophin, alpha 1 | 3.58E-05 | 2.12 |
| TC0600007613.hg.1 | HSPA1A | heat shock 70kDa protein 1A | 3.20E-05 | 2.12 |
| TC1000009169.hg.1 | ACADSB | acyl-CoA dehydrogenase, short/branched chain | 2.49E-04 | 2.12 |
| TC1100006492.hg.1 | PNPLA2 | patatin-like phospholipase domain containing 2 | 5.64E-04 | 2.12 |
| TC0900010849.hg.1 | BARX1 | BARX homeobox 1 | 2.87E-04 | 2.11 |
| TC1300007491.hg.1 | KLF5 | Kruppel-like factor 5 (intestinal) | 2.35E-04 | 2.11 |
| TC2000007704.hg.1 | PARD6B | par-6 family cell polarity regulator beta | 1.06E-04 | 2.11 |
| TC0100015771.hg.1 | SEMA6C | sema domain, transmembrane domain (TM), and cytoplasmic domain, (semaphorin) 6C | 4.02E-05 | 2.11 |
| TC1600008675.hg.1 | KIAA0513 | KIAA0513 | 1.46E-03 | 2.11 |
| TC1500007980.hg.1 | CHRNA5 | cholinergic receptor, nicotinic alpha 5 | 3.30E-04 | 2.11 |
| TC0200009626.hg.1 | LYPD6 | LY6/PLAUR domain containing 6 | 1.48E-04 | 2.10 |
| TC1600007811.hg.1 | PAPD5 | PAP associated domain containing 5 | 8.05E-04 | 2.10 |
| TC0700013603.hg.1 | RASA4 | RAS p21 protein activator 4 | 1.51E-03 | 2.10 |
| TC2000009748.hg.1 | CABLES2 | Cdk5 and Abl enzyme substrate 2 | 1.92E-05 | 2.09 |
| TC1100006495.hg.1 | TSPAN4 | tetraspanin 4 | 1.44E-04 | 2.09 |
| TC1200009122.hg.1 | SIRT4 | sirtuin 4 | 2.46E-04 | 2.09 |
| TC1900007363.hg.1 | MAST3 | microtubule associated serine/threonine kinase 3 | 4.60E-04 | 2.09 |
| TC0300009724.hg.1 | VPS8 | vacuolar protein sorting 8 homolog (S. cerevisiae) | 9.03E-05 | 2.09 |
| TC1700009150.hg.1 | BAIAP2 | BAI1-associated protein 2 | 3.59E-04 | 2.09 |
| TC0600014154.hg.1 | SLC35A1 | solute carrier family 35 (CMP-sialic acid transporter), member A1 | 3.78E-04 | 2.09 |
| TC1200009768.hg.1 | LPCAT3 | lysophosphatidylcholine acyltransferase 3 | 1.29E-03 | 2.09 |
| TC0300007466.hg.1 | ALAS1 | 5-aminolevulinate synthase 1 | 1.57E-03 | 2.09 |
| TC0200011419.hg.1 | ATG4B | autophagy related 4B, cysteine peptidase | 1.28E-04 | 2.09 |
| TC1900010998.hg.1 | PRKD2 | protein kinase D2 | 4.77E-04 | 2.09 |
| TC0200015958.hg.1 | DNER | delta/notch like EGF repeat containing | 6.61E-04 | 2.08 |
| TC0900011501.hg.1 | NR6A1 | nuclear receptor subfamily 6, group A, member 1 | 5.36E-03 | 2.08 |
| TC0100010775.hg.1 | SOAT1 | sterol O-acyltransferase 1 | 2.28E-04 | 2.08 |
| TC1200006946.hg.1 | H2AFJ | H2A histone family, member J | 1.19E-03 | 2.08 |
| TC0400012882.hg.1 | ZNF732 | zinc finger protein 732 | 4.69E-04 | 2.08 |
| TC0700011519.hg.1 | STAG3L2 | stromal antigen 3-like 2 (pseudogene) | 3.00E-05 | 2.08 |
| TC1900009824.hg.1 | DNAJB1 | DnaJ (Hsp40) homolog, subfamily B, member 1 | 1.51E-03 | 2.08 |
| TC0100018285.hg.1 | NBPF19 | neuroblastoma breakpoint family, member 19 | 1.50E-04 | 2.08 |
| TC0700013509.hg.1 | DAGLB | diacylglycerol lipase, beta | 1.83E-03 | 2.08 |
| TC0400006540.hg.1 | FGFR3 | fibroblast growth factor receptor 3 | 1.73E-04 | 2.07 |
| TC0600007400.hg.1 | ZSCAN16 | zinc finger and SCAN domain containing 16 | 3.14E-05 | 2.07 |
| TC0300007720.hg.1 | KBTBD8 | kelch repeat and BTB (POZ) domain containing 8 | 1.05E-03 | 2.07 |
| TC1200006520.hg.1 | TSPAN9 | tetraspanin 9 | 4.70E-03 | 2.07 |
| TC1900011653.hg.1 | MCOLN1 | mucolipin 1 | 2.63E-04 | 2.07 |
| TC2000008910.hg.1 | EIF2S2 | Zhang2013 ALT_ACCEPTOR, ALT_DONOR, coding, INTERNAL, intronic best transcript NM_003908 | 1.79E-03 | 2.07 |
| TC0200016624.hg.1 | KIDINS220 | kinase D-interacting substrate 220kDa | 3.84E-04 | 2.07 |
| TC1700008668.hg.1 | PRKCA | Jeck2013 ALT_ACCEPTOR, ALT_DONOR, coding, INTERNAL, intronic best transcript NM_002737 | 2.29E-03 | 2.07 |
| TC1100009445.hg.1 | FAM118B | family with sequence similarity 118, member B | 2.36E-03 | 2.07 |
| TC0700011924.hg.1 | TMEM130 | transmembrane protein 130 | 4.78E-03 | 2.07 |
| TC0100009442.hg.1 | WNT2B | wingless-type MMTV integration site family, member 2B | 4.71E-04 | 2.07 |
| TC0700013323.hg.1 | SUN1 | Sad1 and UNC84 domain containing 1 | 3.33E-04 | 2.07 |
| HTA2-pos-47421925_st | NA | NA | 3.86E-03 | 2.07 |
| TC1100012020.hg.1 | SESN3 | Transcript Identified by AceView, Entrez Gene ID(s) 143686 | 8.09E-04 | 2.06 |
| TC0700010189.hg.1 | CYTH3 | cytohesin 3 | 2.12E-04 | 2.06 |
| TC0200015226.hg.1 | HIBCH | Transcript Identified by AceView, Entrez Gene ID(s) 26275 | 1.68E-03 | 2.06 |
| 23075672 | NA | NA | 6.85E-04 | 2.06 |
| 23075844 | NA | NA | 6.85E-04 | 2.06 |
| 23075993 | NA | NA | 6.85E-04 | 2.06 |
| 23076303 | NA | NA | 6.85E-04 | 2.06 |
| 23076462 | NA | NA | 6.85E-04 | 2.06 |
| TC0900008851.hg.1 | DNM1 | dynamin 1 | 5.19E-04 | 2.06 |
| TC1600009620.hg.1 | SMG1 | SMG1 phosphatidylinositol 3-kinase-related kinase | 7.01E-04 | 2.05 |
| TC1200011991.hg.1 | IQCD | IQ motif containing D | 4.18E-05 | 2.05 |
| TC2000008367.hg.1 | PAK7 | p21 protein (Cdc42/Rac)-activated kinase 7 | 1.17E-04 | 2.05 |
| TC1900011861.hg.1 | S1PR2 | sphingosine-1-phosphate receptor 2 | 1.25E-03 | 2.05 |
| TC0700008072.hg.1 | RHBDD2 | rhomboid domain containing 2 | 1.10E-03 | 2.05 |
| HTA2-pos-47421981_st | NA | NA | 8.35E-04 | 2.05 |
| TC0100017445.hg.1 | TLR5 | toll-like receptor 5 | 7.04E-05 | 2.05 |
| TC1900007025.hg.1 | PLPPR2 | phospholipid phosphatase related 2 | 1.28E-03 | 2.04 |
| TC1200010341.hg.1 | PKP2 | plakophilin 2 | 2.13E-03 | 2.04 |
| TC0200014991.hg.1 | GPR155 | G protein-coupled receptor 155 | 3.47E-04 | 2.04 |
| TC1100010397.hg.1 | CCDC34 | coiled-coil domain containing 34 | 4.75E-04 | 2.04 |
| TC1900011891.hg.1 | HOOK2 | hook microtubule-tethering protein 2 | 6.04E-05 | 2.04 |
| TC0300009459.hg.1 | GPR160 | G protein-coupled receptor 160 | 2.41E-04 | 2.04 |
| TC1900006985.hg.1 | PDE4A | phosphodiesterase 4A, cAMP-specific | 2.68E-05 | 2.04 |
| TC1800006891.hg.1 | C18orf8 | chromosome 18 open reading frame 8 | 1.22E-04 | 2.04 |
| TC1700007262.hg.1 | MAP2K3 | mitogen-activated protein kinase kinase 3 | 9.91E-03 | 2.04 |
| 23075807 | NA | NA | 5.82E-05 | 2.04 |

| | | | | |
|---|---|---|---|---|
| TC1200008686.hg.1 | CHST11 | Transcript Identified by AceView, Entrez Gene ID(s) 50515 | 7.98E-03 | 2.04 |
| TSUnmapped00000495.hg.1 | ZDHHC3 | zinc finger, DHHC-type containing 3 | 9.34E-04 | 2.04 |
| TC1600008011.hg.1 | KATNB1 | katanin p80 (WD repeat containing) subunit B 1 | 3.93E-05 | 2.04 |
| TC1200011470.hg.1 | DUSP6 | dual specificity phosphatase 6 | 6.62E-05 | 2.04 |
| HTA2-pos-2985916_st | NA | NA | 9.47E-03 | 2.03 |
| TC1700010252.hg.1 | CORO6 | coronin 6 | 6.83E-03 | 2.03 |
| TC1100013053.hg.1 | RAD9A | RAD9 checkpoint clamp component A | 2.94E-04 | 2.03 |
| TC0900008148.hg.1 | TDRD7 | tudor domain containing 7 | 2.26E-03 | 2.03 |
| TC0500007552.hg.1 | LOC100421561 | family with sequence similarity 133, member A pseudogene | 1.41E-03 | 2.03 |
| TC0600014148.hg.1 | CYB5R4 | cytochrome b5 reductase 4 | 1.20E-03 | 2.03 |
| TC1300008253.hg.1 | CRYL1 | crystallin lambda 1 | 1.63E-03 | 2.03 |
| TC0100007909.hg.1 | BMP8A | bone morphogenetic protein 8a | 8.93E-05 | 2.03 |
| TC1100007063.hg.1 | ANO5 | anoctamin 5 | 7.39E-03 | 2.02 |
| TC0300011264.hg.1 | IL17RD | interleukin 17 receptor D | 4.87E-03 | 2.02 |
| TC0700013526.hg.1 | FAM126A | family with sequence similarity 126, member A | 1.76E-04 | 2.02 |
| TC1900011845.hg.1 | MFSD12 | major facilitator superfamily domain containing 12 | 1.61E-03 | 2.02 |
| TC0600012428.hg.1 | UBE3D | ubiquitin protein ligase E3D | 8.14E-05 | 2.02 |
| TC0900008425.hg.1 | ZNF483 | zinc finger protein 483 | 6.43E-03 | 2.02 |
| TC0700007352.hg.1 | BLVRA | biliverdin reductase A | 2.40E-03 | 2.02 |
| TC1900009823.hg.1 | GIPC1 | GIPC PDZ domain containing family, member 1 | 3.23E-03 | 2.02 |
| TC1100012528.hg.1 | THY1 | Thy-1 cell surface antigen | 2.09E-04 | 2.02 |
| TC0700006783.hg.1 | TSPAN13 | tetraspanin 13 | 8.01E-03 | 2.02 |
| TC0900011163.hg.1 | PTPN3 | protein tyrosine phosphatase, non-receptor type 3 | 1.24E-04 | 2.02 |
| TC1100011372.hg.1 | CHKA | choline kinase alpha | 3.06E-03 | 2.02 |
| 23066331 | NA | NA | 9.70E-03 | 2.02 |
| TC1300009165.hg.1 | PCDH9 | protocadherin 9 | 3.18E-04 | 2.02 |
| TC1500007640.hg.1 | IQCH | IQ motif containing H | 2.11E-03 | 2.02 |
| TC0100013578.hg.1 | ADGRB2 | adhesion G protein-coupled receptor B2 | 1.12E-03 | 2.02 |
| TC0300013636.hg.1 | ACAP2 | Transcript Identified by AceView, Entrez Gene ID(s) 23527 | 3.89E-03 | 2.01 |
| TC0100010543.hg.1 | ATP1B1 | ATPase, Na+/K+ transporting, beta 1 polypeptide | 1.72E-03 | 2.01 |
| TC1100012389.hg.1 | CADM1 | cell adhesion molecule 1 | 2.07E-04 | 2.01 |
| TC1200008373.hg.1 | TMTC3 | transmembrane and tetratricopeptide repeat containing 3 | 6.27E-03 | 2.01 |
| TC1900008113.hg.1 | LTBP4 | latent transforming growth factor beta binding protein 4 | 1.36E-03 | 2.01 |
| TC0800011579.hg.1 | EXT1 | exostosin glycosyltransferase 1 | 4.28E-04 | 2.01 |
| TC0700008745.hg.1 | PRKAR2B | protein kinase, cAMP-dependent, regulatory, type II, beta | 9.09E-03 | 2.01 |
| TC0100010806.hg.1 | XPR1 | xenotropic and polytropic retrovirus receptor 1 | 5.34E-03 | 2.01 |
| TC1500009597.hg.1 | MYO1E | myosin IE | 4.68E-04 | 2.01 |
| TC0500008121.hg.1 | ARSK | arylsulfatase family, member K | 6.20E-04 | 2.01 |
| TC0300013547.hg.1 | FGF12 | fibroblast growth factor 12 | 1.25E-03 | 2.01 |
| TC0400007128.hg.1 | TBC1D19 | TBC1 domain family, member 19 | 6.07E-05 | 2.00 |
| TC0500011596.hg.1 | SLCO4C1 | solute carrier organic anion transporter family, member 4C1 | 3.73E-03 | 2.00 |
| TC1600006610.hg.1 | CCNF | cyclin F | 2.05E-03 | 2.00 |
| TC0100016120.hg.1 | PIGM | phosphatidylinositol glycan anchor biosynthesis class M | 9.69E-05 | 2.00 |
| TC1600009074.hg.1 | MSRB1 | methionine sulfoxide reductase B1 | 2.72E-03 | 2.00 |
| TC0100007954.hg.1 | SMAP2 | small ArfGAP2 | 6.18E-03 | 2.00 |
| TC0800008916.hg.1 | EFR3A | EFR3 homolog A | 7.97E-04 | 2.00 |
| TC0200007746.hg.1 | B3GNT2 | UDP-GlcNAc:betaGal beta-1,3-N-acetylglucosaminyltransferase 2 | 7.22E-05 | 2.00 |
| TC0500013425.hg.1 | C5orf45 | chromosome 5 open reading frame 45 | 1.01E-04 | 2.00 |
| TC1700012315.hg.1 | SLC25A10 | solute carrier family 25 (mitochondrial carrier; dicarboxylate transporter), member 10 | 2.46E-03 | 1.99 |
| TC0100018208.hg.1 | ZBTB8B | zinc finger and BTB domain containing 8B | 7.89E-04 | 1.99 |
| TC0100009373.hg.1 | SLC6A17 | solute carrier family 6 (neutral amino acid transporter), member 17 | 9.29E-03 | 1.99 |
| HTA2-neg-47421950_st | NA | NA | 7.14E-03 | 1.99 |
| TC1000007703.hg.1 | BICC1 | BicC family RNA binding protein 1 | 7.78E-04 | 1.99 |
| TC0X00007116.hg.1 | RP2 | retinitis pigmentosa 2 (X-linked recessive) | 8.30E-05 | 1.99 |
| TC2100008459.hg.1 | FTCD | formimidoyltransferase cyclodeaminase | 7.92E-03 | 1.99 |
| TC1100007220.hg.1 | DEPDC7 | DEP domain containing 7 | 7.68E-04 | 1.99 |
| TC1700008440.hg.1 | PPM1E | protein phosphatase, Mg2+/Mn2+ dependent, 1E | 1.74E-04 | 1.99 |
| TC0800008663.hg.1 | MAL2 | mal, T-cell differentiation protein 2 (gene/pseudogene) | 1.20E-04 | 1.98 |
| TC0200016637.hg.1 | SMC6 | structural maintenance of chromosomes 6 | 7.13E-03 | 1.98 |
| TC0100017059.hg.1 | RBBP5 | retinoblastoma binding protein 5 | 2.22E-03 | 1.98 |
| TC1700006911.hg.1 | ARHGAP44 | Rho GTPase activating protein 44 | 6.21E-03 | 1.98 |
| TC1200008795.hg.1 | ACACB | acetyl-CoA carboxylase beta | 3.98E-03 | 1.98 |
| TC0800008150.hg.1 | CPNE3 | copine III | 2.71E-05 | 1.98 |
| 23076506 | NA | NA | 6.38E-03 | 1.98 |
| TC1200012680.hg.1 | ACSS3 | acyl-CoA synthetase short-chain family member 3 | 9.18E-04 | 1.98 |
| TC0900009324.hg.1 | CACNA1B | calcium channel, voltage-dependent, N type, alpha 1B subunit | 5.56E-04 | 1.98 |
| TC1200011599.hg.1 | LTA4H | leukotriene A4 hydrolase | 3.85E-04 | 1.98 |
| TC0800007439.hg.1 | POLB | polymerase (DNA directed), beta | 1.79E-04 | 1.98 |
| TC1400009175.hg.1 | NID2 | nidogen 2 (osteonidogen) | 2.49E-03 | 1.98 |
| TC1300007312.hg.1 | PCDH17 | protocadherin 17 | 5.12E-04 | 1.98 |
| TC1900009201.hg.1 | MKNK2 | MAP kinase interacting serine/threonine kinase 2 | 7.47E-04 | 1.98 |
| TC0200014790.hg.1 | GRB14 | growth factor receptor bound protein 14 | 5.48E-04 | 1.97 |
| TC0800012323.hg.1 | CA13 | carbonic anhydrase XIII | 8.93E-04 | 1.97 |
| TC0200010275.hg.1 | GLS | glutaminase | 1.88E-03 | 1.97 |
| TC0800012405.hg.1 | FUT10 | fucosyltransferase 10 (alpha (1,3) fucosyltransferase) | 1.93E-03 | 1.97 |
| TC0700011050.hg.1 | FIGNL1 | fidgetin-like 1 | 2.28E-04 | 1.97 |
| TC1900007382.hg.1 | PGPEP1 | pyroglutamyl-peptidase I | 2.97E-04 | 1.97 |
| TC0600014150.hg.1 | SMIM8 | small integral membrane protein 8 | 1.08E-03 | 1.97 |
| TC1900011981.hg.1 | ERCC2 | excision repair cross-complementation group 2 | 2.26E-04 | 1.97 |
| TC0200006983.hg.1 | EFR3B | EFR3 homolog B | 6.39E-03 | 1.96 |
| TC0500012160.hg.1 | NME5 | NME/NM23 family member 5 | 4.96E-04 | 1.96 |
| TC0400010733.hg.1 | KDR | kinase insert domain receptor | 5.51E-03 | 1.96 |
| TC0600007103.hg.1 | KDM1B | lysine (K)-specific demethylase 1B | 8.28E-05 | 1.96 |
| TC1500010755.hg.1 | ANKDD1A | ankyrin repeat and death domain containing 1A | 5.63E-05 | 1.96 |
| TC0500009521.hg.1 | CPEB4 | cytoplasmic polyadenylation element binding protein 4 | 6.45E-04 | 1.96 |
| TC1900011874.hg.1 | ZNF878 | zinc finger protein 878 | 3.69E-04 | 1.96 |
| TC1500008470.hg.1 | ARRDC4 | arrestin domain containing 4 | 7.79E-04 | 1.96 |
| TC1400009198.hg.1 | DDHD1 | DDHD domain containing 1 | 3.54E-03 | 1.96 |
| TC0200009071.hg.1 | CFAP221 | cilia and flagella associated protein 221 | 3.56E-04 | 1.96 |
| HTA2-pos-47421982_st | NA | NA | 2.10E-03 | 1.96 |
| TC1200006905.hg.1 | FAM234B | family with sequence similarity 234, member B | 5.14E-04 | 1.96 |
| TC0900011340.hg.1 | ASTN2 | astrotactin 2 | 6.21E-03 | 1.96 |
| TC0700013575.hg.1 | STAG3L3 | stromal antigen 3-like 3 (pseudogene) | 3.45E-05 | 1.96 |
| TC2000007695.hg.1 | PTPN1 | protein tyrosine phosphatase, non-receptor type 1 | 1.32E-03 | 1.96 |
| TC0100015701.hg.1 | HIST2H3A | histone cluster 2, H3a | 2.84E-03 | 1.96 |
| TC0300012765.hg.1 | HLTF | Transcript Identified by AceView, Entrez Gene ID(s) 6596 | 2.61E-03 | 1.96 |
| TC0100009181.hg.1 | PLPPR4 | phospholipid phosphatase related 4 | 8.78E-03 | 1.96 |
| TSUnmapped00000050.hg.1 | MLXIP | MLX interacting protein | 2.68E-03 | 1.95 |
| TC2000007670.hg.1 | SNAI1 | snail family zinc finger 1 | 1.88E-04 | 1.95 |
| TC2200008449.hg.1 | DUSP18 | dual specificity phosphatase 18 | 1.46E-03 | 1.95 |
| TC0200006715.hg.1 | PQLC3 | PQ loop repeat containing 3 | 2.71E-03 | 1.95 |
| TC1700011448.hg.1 | POLG2 | polymerase (DNA directed), gamma 2, accessory subunit | 2.32E-04 | 1.95 |
| TC0700013327.hg.1 | CHST12 | carbohydrate (chondroitin 4) sulfotransferase 12 | 8.89E-04 | 1.95 |
| TC2100007241.hg.1 | ABCG1 | ATP binding cassette subfamily G member 1 | 5.03E-04 | 1.95 |
| TC0300008715.hg.1 | EEFSEC | eukaryotic elongation factor, selenocysteine-tRNA-specific | 2.16E-04 | 1.95 |
| TC1100011052.hg.1 | FADS3 | fatty acid desaturase 3 | 7.12E-04 | 1.95 |
| TSUnmapped00000128.hg.1 | ZNF852 | zinc finger protein 852 | 2.75E-03 | 1.95 |
| TC0700013388.hg.1 | STAG3L1 | stromal antigen 3-like 1 (pseudogene) | 3.26E-05 | 1.95 |
| TC1100007812.hg.1 | LRRC10B | leucine rich repeat containing 10B | 1.03E-04 | 1.95 |
| TC0600009143.hg.1 | SLC16A10 | solute carrier family 16 (aromatic amino acid transporter), member 10 | 5.01E-04 | 1.94 |
| TC1500007615.hg.1 | MAP2K1 | Transcript Identified by AceView, Entrez Gene ID(s) 5604 | 2.56E-03 | 1.94 |
| TC1600010987.hg.1 | CMIP | Memczak2013 ANTISENSE, coding, INTERNAL, intronic best transcript NM_198390 | 5.28E-05 | 1.94 |
| TC0900011523.hg.1 | HSPA5 | heat shock 70kDa protein 5 (glucose-regulated protein, 78kDa) | 8.38E-04 | 1.94 |
| TC1700012212.hg.1 | CCDC144NL-AS1 | CCDC144NL antisense RNA 1 | 8.49E-03 | 1.94 |
| TC1100010418.hg.1 | KIF18A | kinesin family member 18A | 8.09E-03 | 1.94 |
| TC0500010780.hg.1 | SLC38A9 | Transcript Identified by AceView, Entrez Gene ID(s) 153129 | 4.57E-03 | 1.94 |
| TC1600007240.hg.1 | PRKCB | protein kinase C, beta | 4.72E-03 | 1.94 |
| TC0200007296.hg.1 | GALM | galactose mutarotase (aldose 1-epimerase) | 1.27E-04 | 1.94 |
| TC1200010156.hg.1 | CASC1 | cancer susceptibility candidate 1 | 1.97E-04 | 1.94 |

| | | | | |
|---|---|---|---|---|
| TC0800010163.hg.1 | FGFR1 | fibroblast growth factor receptor 1 | 4.10E-04 | 1.94 |
| TC0X00009117.hg.1 | FANCB | Fanconi anemia complementation group B | 1.52E-04 | 1.94 |
| TC0700011780.hg.1 | PEX1 | peroxisomal biogenesis factor 1 | 2.68E-04 | 1.94 |
| TC1400009787.hg.1 | SPTLC2 | serine palmitoyltransferase, long chain base subunit 2 | 4.15E-03 | 1.93 |
| TC0100013417.hg.1 | SLC9A1 | solute carrier family 9, subfamily A (NHE1, cation proton antiporter 1), member 1 | 3.24E-03 | 1.93 |
| TC1100006576.hg.1 | CD81 | CD81 molecule | 2.67E-04 | 1.93 |
| 23075476 | NA | NA | 8.27E-04 | 1.93 |
| TC1700008441.hg.1 | PPM1E | Transcript Identified by AceView, Entrez Gene ID(s) 22843 | 5.08E-03 | 1.93 |
| TC0600011127.hg.1 | HIST1H1C | histone cluster 1, H1c | 1.77E-03 | 1.93 |
| TC0900008933.hg.1 | TOR1B | torsin family 1, member B (torsin B) | 3.83E-04 | 1.93 |
| TC1100007390.hg.1 | EXT2 | exostosin glycosyltransferase 2 | 1.22E-03 | 1.93 |
| TC0800007328.hg.1 | BAG4 | BCL2-associated athanogene 4 | 5.77E-03 | 1.93 |
| TC0400011043.hg.1 | CDKL2 | cyclin-dependent kinase-like 2 (CDC2-related kinase) | 6.18E-03 | 1.93 |
| TC0200009402.hg.1 | MGAT5 | mannosyl (alpha-1,6-)-glycoprotein beta-1,6-N-acetyl-glucosaminyltransferase | 1.11E-03 | 1.93 |
| TC0100014365.hg.1 | CYP2J2 | cytochrome P450, family 2, subfamily J, polypeptide 2 | 8.91E-04 | 1.93 |
| TC0100018261.hg.1 | GSTM4 | glutathione S-transferase mu 4 | 4.10E-03 | 1.93 |
| TC0100011463.hg.1 | SYT14 | synaptotagmin XIV | 6.69E-03 | 1.93 |
| TC1900007356.hg.1 | KCNN1 | potassium channel, calcium activated intermediate/small conductance subfamily N alpha, member 1 | 4.77E-04 | 1.92 |
| TC1400007307.hg.1 | ARID4A | AT rich interactive domain 4A (RBP1-like) | 3.50E-04 | 1.92 |
| TC0700008617.hg.1 | SH2B2 | SH2B adaptor protein 2 | 4.50E-03 | 1.92 |
| TC0100007634.hg.1 | ZCCHC17 | zinc finger, CCHC domain containing 17 | 3.23E-03 | 1.92 |
| TC0100018479.hg.1 | NOTCH2NL | notch 2 N-terminal like | 2.60E-04 | 1.92 |
| TC1800006715.hg.1 | IMPA2 | inositol(myo)-1(or 4)-monophosphatase 2 | 2.15E-04 | 1.92 |
| TC1200007236.hg.1 | METTL20 | methyltransferase like 20 | 4.57E-03 | 1.92 |
| TC0100018437.hg.1 | MKNK1 | MAP kinase interacting serine/threonine kinase 1 | 1.26E-04 | 1.92 |
| TC1200012700.hg.1 | C12orf75 | chromosome 12 open reading frame 75 | 4.32E-03 | 1.92 |
| TC0800012351.hg.1 | WDYHV1 | WDYHV motif containing 1 | 1.91E-04 | 1.92 |
| TC0500010203.hg.1 | MYO10 | myosin X | 2.52E-04 | 1.92 |
| TC0X00010532.hg.1 | AMOT | angiomotin | 1.03E-03 | 1.92 |
| TC1400010759.hg.1 | ATP6V1D | ATPase, H+ transporting, lysosomal 34kDa, V1 subunit D | 6.91E-04 | 1.92 |
| TC0300006520.hg.1 | LMCD1 | LIM and cysteine-rich domains 1 | 6.19E-04 | 1.92 |
| TC0100018417.hg.1 | ALDH4A1 | aldehyde dehydrogenase 4 family, member A1 | 3.08E-04 | 1.92 |
| TC1100012535.hg.1 | PVRL1 | poliovirus receptor-related 1 (herpesvirus entry mediator C) | 1.60E-03 | 1.92 |
| TC1700008228.hg.1 | ITGA3 | integrin alpha 3 | 6.51E-03 | 1.92 |
| TC0100017716.hg.1 | FAM89A | family with sequence similarity 89, member A | 5.05E-05 | 1.92 |
| TC0900011286.hg.1 | DFNB31 | deafness, autosomal recessive 31 | 3.20E-04 | 1.92 |
| TC0200015776.hg.1 | RESP18 | regulated endocrine-specific protein 18 | 4.84E-03 | 1.92 |
| TC0600013727.hg.1 | EZR | ezrin | 4.74E-03 | 1.92 |
| TC1900010391.hg.1 | PEPD | peptidase D | 3.50E-04 | 1.92 |
| TC1600007957.hg.1 | MT2A | metallothionein 2A | 1.18E-03 | 1.92 |
| TC1900009433.hg.1 | TUBB4A | tubulin, beta 4A class IVa | 3.65E-04 | 1.91 |
| TC0500007136.hg.1 | SPEF2 | sperm flagellar 2 | 8.73E-04 | 1.91 |
| TC0100009656.hg.1 | FCGR1B | Fc fragment of IgG, high affinity Ib, receptor (CD64) | 4.89E-03 | 1.91 |
| TC0700006977.hg.1 | NFE2L3 | nuclear factor, erythroid 2-like 3 | 5.01E-03 | 1.91 |
| TC0900009307.hg.1 | ARRDC1 | arrestin domain containing 1 | 3.94E-04 | 1.91 |
| TC0X00007707.hg.1 | ATP7A | ATPase, Cu++ transporting, alpha polypeptide | 2.87E-03 | 1.91 |
| TC1400007201.hg.1 | CDKN3 | cyclin-dependent kinase inhibitor 3 | 1.46E-04 | 1.91 |
| TC1600008020.hg.1 | USB1 | U6 snRNA biogenesis 1 | 2.01E-04 | 1.91 |
| TC1100012052.hg.1 | CCDC82 | coiled-coil domain containing 82 | 3.91E-04 | 1.91 |
| TC1500010753.hg.1 | TLN2 | talin 2 | 5.67E-04 | 1.91 |
| TC0300007432.hg.1 | MAPKAPK3 | mitogen-activated protein kinase-activated protein kinase 3 | 1.14E-04 | 1.91 |
| TC0100015246.hg.1 | KCNA3 | potassium channel, voltage gated shaker related subfamily A, member 3 | 2.44E-03 | 1.91 |
| TC1700007949.hg.1 | ARL4D | ADP-ribosylation factor like GTPase 4D | 5.27E-03 | 1.91 |
| TC1600006893.hg.1 | CLEC16A | C-type lectin domain family 16, member A | 3.72E-04 | 1.91 |
| TC0900011260.hg.1 | ALAD | aminolevulinate dehydratase | 6.15E-05 | 1.91 |
| TC0100014218.hg.1 | SCP2 | Jeck2013 ANTISENSE, coding, INTERNAL, OVEXON, UTR3 best transcript NM_001007098 | 2.24E-04 | 1.90 |
| TC0800012306.hg.1 | MTFR1 | mitochondrial fission regulator 1 | 1.86E-03 | 1.90 |
| TC0600012452.hg.1 | CEP162 | centrosomal protein 162kDa | 9.77E-05 | 1.90 |
| TC2100008533.hg.1 | PCBP3 | poly(rC) binding protein 3 | 4.90E-04 | 1.90 |
| TC1200009452.hg.1 | MMP17 | matrix metallopeptidase 17 (membrane-inserted) | 9.46E-04 | 1.90 |
| TC0400010784.hg.1 | NOA1 | nitric oxide associated 1 | 4.97E-03 | 1.90 |
| TC0600013604.hg.1 | RGS17 | regulator of G-protein signaling 17 | 2.14E-04 | 1.90 |
| TC0100017058.hg.1 | TMEM81 | transmembrane protein 81 | 4.54E-03 | 1.90 |
| TC0100007365.hg.1 | NIPAL3 | NIPA-like domain containing 3 | 8.47E-04 | 1.90 |
| TC0300009353.hg.1 | PPM1L | protein phosphatase, Mg2+/Mn2+ dependent, 1L | 3.31E-03 | 1.90 |
| TC1600009944.hg.1 | KCTD13 | potassium channel tetramerization domain containing 13 | 3.20E-04 | 1.90 |
| TC1200008906.hg.1 | TRAFD1 | TRAF-type zinc finger domain containing 1 | 3.58E-03 | 1.90 |
| TC1000010499.hg.1 | ZFAND4 | zinc finger, AN1-type domain 4 | 4.17E-04 | 1.90 |
| TC0400008331.hg.1 | NPNT | nephronectin | 4.58E-03 | 1.90 |
| TC0300013025.hg.1 | BCHE | butyrylcholinesterase | 2.24E-03 | 1.90 |
| TC0100018302.hg.1 | EFNA4 | ephrin-A4 | 6.83E-04 | 1.89 |
| TC0100009877.hg.1 | HIST2H3A | histone cluster 2, H3a | 3.64E-03 | 1.89 |
| TC0200011013.hg.1 | FBXO36 | F-box protein 36 | 1.97E-03 | 1.89 |
| TC1600009351.hg.1 | GRIN2A | Transcript Identified by AceView, Entrez Gene ID(s) 2903 | 1.65E-04 | 1.89 |
| TSUnmapped00000488.hg.1 | ZDHHC3 | zinc finger, DHHC-type containing 3 | 1.31E-03 | 1.89 |
| TC0100015180.hg.1 | WDR47 | WD repeat domain 47 | 2.22E-03 | 1.89 |
| TC0100011269.hg.1 | LINC00260 | long intergenic non-protein coding RNA 260 | 5.62E-04 | 1.89 |
| TC0400011579.hg.1 | PLA2G12A | phospholipase A2, group XIIA | 8.41E-04 | 1.89 |
| TC1700011181.hg.1 | MMD | monocyte to macrophage differentiation-associated | 1.23E-03 | 1.89 |
| TC0100016659.hg.1 | APOBEC4 | apolipoprotein B mRNA editing enzyme, catalytic polypeptide-like 4 (putative) | 5.63E-04 | 1.89 |
| TC0600012040.hg.1 | TRAM2 | translocation associated membrane protein 2 | 7.31E-04 | 1.89 |
| TC0300012319.hg.1 | goyborbu | Transcript Identified by AceView | 1.30E-03 | 1.88 |
| TC1100012906.hg.1 | IGSF9B | immunoglobulin superfamily, member 9B | 1.44E-04 | 1.88 |
| TC0100009097.hg.1 | ABCD3 | ATP binding cassette subfamily D member 3 | 2.34E-03 | 1.88 |
| TC0500013186.hg.1 | PTCD2 | pentatricopeptide repeat domain 2 | 2.33E-03 | 1.88 |
| TC1100008091.hg.1 | CCS | copper chaperone for superoxide dismutase | 7.87E-03 | 1.88 |
| TC0X00011278.hg.1 | PRRG1 | proline rich Gla (G-carboxyglutamic acid) 1 | 1.13E-03 | 1.88 |
| TC1900012045.hg.1 | RDH13 | retinol dehydrogenase 13 (all-trans/9-cis) | 6.82E-04 | 1.88 |
| TC0X00007053.hg.1 | MAOA | monoamine oxidase A | 1.44E-04 | 1.88 |
| TC1800008862.hg.1 | PIGN | phosphatidylinositol glycan anchor biosynthesis class N | 2.53E-03 | 1.88 |
| TC0900006762.hg.1 | ACER2 | alkaline ceramidase 2 | 5.58E-04 | 1.88 |
| TC0300010038.hg.1 | SENP5 | Transcript Identified by AceView, Entrez Gene ID(s) 205564 | 4.99E-03 | 1.88 |
| TC1700011277.hg.1 | TRIM37 | Transcript Identified by AceView, Entrez Gene ID(s) 4591 | 8.95E-03 | 1.88 |
| TC0900010925.hg.1 | ZNF782 | zinc finger protein 782 | 8.02E-04 | 1.88 |
| TC0600014194.hg.1 | TIAM2 | T-cell lymphoma invasion and metastasis 2 | 5.12E-03 | 1.88 |
| TC1600009642.hg.1 | GDE1 | glycerophosphodiester phosphodiesterase 1 | 6.99E-03 | 1.87 |
| TC1900009297.hg.1 | TBXA2R | thromboxane A2 receptor | 6.77E-04 | 1.87 |
| TC0200015556.hg.1 | FZD5 | frizzled class receptor 5 | 3.03E-03 | 1.87 |
| TC1100011867.hg.1 | FZD4 | frizzled class receptor 4 | 4.90E-04 | 1.87 |
| TC0X00008235.hg.1 | LONRF3 | LON peptidase N-terminal domain and ring finger 3 | 6.60E-03 | 1.87 |
| TC0200011402.hg.1 | FARP2 | FERM, ARH/RhoGEF and pleckstrin domain protein 2 | 6.91E-04 | 1.87 |
| TC0300008979.hg.1 | CLSTN2 | calsyntenin 2 | 1.52E-03 | 1.87 |
| TC1500007695.hg.1 | PAQR5 | progestin and adipoQ receptor family member V | 1.29E-04 | 1.87 |
| TC0100006861.hg.1 | FBXO44 | F-box protein 44 | 8.33E-04 | 1.87 |
| TC0X00008394.hg.1 | BCORL1 | BCL6 corepressor-like 1 | 5.95E-05 | 1.87 |
| TC1100012526.hg.1 | USP2 | ubiquitin specific peptidase 2 | 1.28E-03 | 1.87 |
| HTA2-pos-47421983_st | NA | NA | 9.79E-04 | 1.87 |
| TC0400009001.hg.1 | FAM160A1 | family with sequence similarity 160, member A1 | 5.37E-03 | 1.87 |
| TC0700010352.hg.1 | ANKMY2 | ankyrin repeat and MYND domain containing 2 | 7.81E-03 | 1.86 |
| TC0700010582.hg.1 | HIBADH | Salzman2013 ALT_ACCEPTOR, ALT_DONOR, coding, INTERNAL, intronic best transcript NM_152740 | 6.59E-04 | 1.86 |
| TC2000009014.hg.1 | NDRG3 | NDRG family member 3 | 4.66E-04 | 1.86 |
| TC0X00007507.hg.1 | EFNB1 | ephrin-B1 | 3.45E-03 | 1.86 |
| TC0800009853.hg.1 | BIN3 | bridging integrator 3 | 2.83E-04 | 1.86 |
| TC0900008109.hg.1 | HABP4 | hyaluronan binding protein 4 | 1.41E-04 | 1.86 |
| TC0100013281.hg.1 | ASAP3 | ArfGAP with SH3 domain, ankyrin repeat and PH domain 3 | 1.44E-03 | 1.86 |
| TC1200010839.hg.1 | ITGA5 | integrin alpha 5 | 5.29E-04 | 1.86 |
| TC2000006688.hg.1 | SNAP25 | synaptosome associated protein 25kDa | 3.73E-03 | 1.86 |
| TC1700008372.hg.1 | SCPEP1 | serine carboxypeptidase 1 | 2.86E-03 | 1.86 |
| TC1500010846.hg.1 | GOLGA8K | golgin A8 family, member K | 3.46E-04 | 1.86 |

| | | | | |
|---|---|---|---|---|
| TC0100009418.hg.1 | RAP1A | RAP1A, member of RAS oncogene family | 7.42E-04 | 1.86 |
| TC1000008005.hg.1 | MCU | mitochondrial calcium uniporter | 1.85E-03 | 1.86 |
| TC1100010175.hg.1 | RRAS2 | related RAS viral (r-ras) oncogene homolog 2 | 7.78E-03 | 1.86 |
| TC0500009046.hg.1 | ADRB2 | adrenoceptor beta 2, surface | 2.53E-03 | 1.86 |
| TC0900008865.hg.1 | CERCAM | cerebral endothelial cell adhesion molecule | 4.84E-03 | 1.86 |
| TC2000010027.hg.1 | TMEM189-UBE2V1 | TMEM189-UBE2V1 readthrough | 2.70E-03 | 1.85 |
| TC0200009687.hg.1 | ARL6IP6 | ADP-ribosylation factor like GTPase 6 interacting protein 6 | 4.49E-04 | 1.85 |
| TC0200014851.hg.1 | LRP2 | LDL receptor related protein 2 | 5.25E-03 | 1.85 |
| 23071946 | NA | NA | 1.38E-04 | 1.85 |
| TC0400011749.hg.1 | BBS7 | Bardet-Biedl syndrome 7 | 1.67E-03 | 1.85 |
| TC0100018481.hg.1 | NOTCH2NL | notch 2 N-terminal like | 4.19E-04 | 1.85 |
| TC1600011292.hg.1 | FANCA | Fanconi anemia complementation group A | 7.03E-03 | 1.85 |
| TC0600008571.hg.1 | MYO6 | Transcript Identified by AceView, Entrez Gene ID(s) 4646 | 2.90E-03 | 1.84 |
| HTA2-pos-3513255_st | NA | NA | 8.30E-04 | 1.84 |
| TC0100015211.hg.1 | GSTM3 | glutathione S-transferase mu 3 (brain) | 2.78E-03 | 1.84 |
| TC1100011643.hg.1 | ARRB1 | arrestin, beta 1 | 2.53E-03 | 1.84 |
| TC0300012764.hg.1 | HLTF | helicase-like transcription factor | 9.11E-03 | 1.84 |
| TC1300008834.hg.1 | SIAH3 | siah E3 ubiquitin protein ligase family member 3 | 2.54E-04 | 1.84 |
| TC0600010960.hg.1 | TPMT | thiopurine S-methyltransferase | 7.89E-03 | 1.84 |
| HTA2-pos-PSR12013086.hg.1 | NA | NA | 7.93E-03 | 1.84 |
| TC1600008943.hg.1 | TMEM8A | transmembrane protein 8A | 2.65E-04 | 1.84 |
| TC0300009713.hg.1 | EPHB3 | EPH receptor B3 | 2.78E-03 | 1.84 |
| TC0500013088.hg.1 | MAPK9 | mitogen-activated protein kinase 9 | 1.08E-03 | 1.84 |
| TC0600011809.hg.1 | CCND3 | cyclin D3 | 1.12E-03 | 1.84 |
| TC2000008267.hg.1 | RASSF2 | Ras association (RalGDS/AF-6) domain family member 2 | 2.35E-04 | 1.84 |
| TC2200007909.hg.1 | MICAL3 | microtubule associated monooxygenase, calponin and LIM domain containing 3 | 1.91E-04 | 1.84 |
| 23071925 | NA | NA | 1.65E-04 | 1.84 |
| TC0200015314.hg.1 | STK17B | serine/threonine kinase 17b | 4.72E-03 | 1.84 |
| HTA2-pos-3371082_st | NA | NA | 7.01E-03 | 1.84 |
| TC0800010045.hg.1 | TEX15 | testis expressed 15 | 9.68E-03 | 1.84 |
| TC1300006643.hg.1 | ATP8A2 | ATPase, aminophospholipid transporter, class I, type 8A, member 2 | 2.58E-04 | 1.83 |
| TC0700012731.hg.1 | DGKI | diacylglycerol kinase, iota | 8.11E-04 | 1.83 |
| TC1900010025.hg.1 | CRLF1 | cytokine receptor-like factor 1 | 5.14E-04 | 1.83 |
| TC0400007016.hg.1 | SLIT2 | slit guidance ligand 2 | 3.28E-03 | 1.83 |
| TC0300011628.hg.1 | ROBO1 | roundabout guidance receptor 1 | 2.94E-04 | 1.83 |
| TC0700007137.hg.1 | BBS9 | Bardet-Biedl syndrome 9 | 8.93E-03 | 1.83 |
| TC1000009908.hg.1 | RSU1 | Ras suppressor protein 1 | 5.86E-04 | 1.83 |
| TC1900009302.hg.1 | PIP5K1C | phosphatidylinositol-4-phosphate 5-kinase, type I, gamma | 6.56E-04 | 1.83 |
| TC0900007113.hg.1 | RECK | reversion-inducing-cysteine-rich protein with kazal motifs | 2.59E-03 | 1.83 |
| TC1100011101.hg.1 | NXF1 | nuclear RNA export factor 1 | 2.11E-03 | 1.83 |
| HTA2-pos-PSR02023982.hg.1 | NA | NA | 8.77E-04 | 1.83 |
| TC1600011465.hg.1 | ERVK13-1 | endogenous retrovirus group K13, member 1 | 1.08E-03 | 1.83 |
| TC0100010159.hg.1 | SLC25A44 | solute carrier family 25, member 44 | 2.31E-03 | 1.83 |
| TC0100006675.hg.1 | KCNAB2 | potassium channel, voltage gated subfamily A regulatory beta subunit 2 | 2.45E-03 | 1.82 |
| TC2100006714.hg.1 | NCAM2 | neural cell adhesion molecule 2 | 3.03E-04 | 1.82 |
| TC2100007420.hg.1 | COL18A1 | collagen, type XVIII, alpha 1 | 6.89E-04 | 1.82 |
| TC1700006767.hg.1 | WRAP53 | WD repeat containing, antisense to TP53 | 1.14E-03 | 1.82 |
| TC1900010005.hg.1 | RAB3A | RAB3A, member RAS oncogene family | 1.72E-03 | 1.82 |
| TC0700008295.hg.1 | CFAP69 | cilia and flagella associated protein 69 | 8.32E-04 | 1.82 |
| HTA2-pos-2978678_st | NA | NA | 1.35E-03 | 1.82 |
| TC1100010105.hg.1 | GALNT18 | polypeptide N-acetylgalactosaminyltransferase 18 | 7.38E-04 | 1.82 |
| HTA2-pos-PSR02023992.hg.1 | NA | NA | 3.79E-03 | 1.82 |
| TC0200011054.hg.1 | ARMC9 | armadillo repeat containing 9 | 5.11E-03 | 1.82 |
| TC0100015483.hg.1 | NOTCH2 | notch 2 | 4.04E-04 | 1.82 |
| TC1800009026.hg.1 | FBXO15 | F-box protein 15 | 6.86E-03 | 1.82 |
| TC2200006907.hg.1 | SGSM1 | small G protein signaling modulator 1 | 3.55E-03 | 1.82 |
| TC0700011500.hg.1 | STX1A | syntaxin 1A (brain) | 8.03E-04 | 1.82 |
| TC0100013804.hg.1 | BMP8B | bone morphogenetic protein 8b | 2.50E-03 | 1.82 |
| TC1100009302.hg.1 | TECTA | tectorin alpha | 8.57E-04 | 1.82 |
| TC0500012285.hg.1 | FCHSD1 | FCH and double SH3 domains 1 | 3.66E-03 | 1.82 |
| TC0200007929.hg.1 | GMCL1 | germ cell-less, spermatogenesis associated 1 | 9.87E-03 | 1.81 |
| TC1200006570.hg.1 | KCNA1 | potassium channel, voltage gated shaker related subfamily A, member 1 | 6.96E-03 | 1.81 |
| HTA2-pos-2978682_st | NA | NA | 7.71E-04 | 1.81 |
| TC1700008254.hg.1 | ACSF2 | acyl-CoA synthetase family member 2 | 8.92E-03 | 1.81 |
| TC1400010261.hg.1 | MOK | MOK protein kinase | 9.63E-04 | 1.81 |
| TC0200007353.hg.1 | EML4 | echinoderm microtubule associated protein like 4 | 6.15E-04 | 1.81 |
| TC1500010157.hg.1 | ADAMTS7 | ADAM metallopeptidase with thrombospondin type 1 motif 7 | 9.56E-05 | 1.81 |
| TC2000007476.hg.1 | STK4 | serine/threonine kinase 4 | 4.58E-04 | 1.81 |
| HTA2-pos-2985914_st | NA | NA | 2.73E-03 | 1.81 |
| TC0100018303.hg.1 | EFNA3 | ephrin-A3 | 1.05E-04 | 1.81 |
| HTA2-neg-47421381_st | NA | NA | 7.62E-03 | 1.81 |
| TC1400010620.hg.1 | SNAPC1 | small nuclear RNA activating complex polypeptide 1 | 2.38E-03 | 1.81 |
| TC0X00010130.hg.1 | MAGT1 | magnesium transporter 1 | 1.72E-04 | 1.81 |
| TC1100010718.hg.1 | LRP4 | LDL receptor related protein 4 | 4.22E-04 | 1.81 |
| TC1100011609.hg.1 | P4HA3 | prolyl 4-hydroxylase, alpha polypeptide III | 7.97E-04 | 1.81 |
| TC2100008536.hg.1 | PCBP3 | poly(rC) binding protein 3 | 1.53E-03 | 1.81 |
| TC1500010908.hg.1 | STARD5 | StAR-related lipid transfer domain containing 5 | 4.86E-04 | 1.81 |
| TC0800008467.hg.1 | ATP6V1C1 | ATPase, H+ transporting, lysosomal 42kDa, V1 subunit C1 | 8.28E-03 | 1.81 |
| TC0200015434.hg.1 | ALS2 | ALS2, alsin Rho guanine nucleotide exchange factor | 6.35E-04 | 1.80 |
| TC0600008156.hg.1 | ENPP4 | ectonucleotide pyrophosphatase/phosphodiesterase 4 (putative) | 2.52E-03 | 1.80 |
| TC0400008329.hg.1 | GSTCD | glutathione S-transferase, C-terminal domain containing | 6.43E-04 | 1.80 |
| 23072018 | NA | NA | 2.68E-04 | 1.80 |
| 23072048 | NA | NA | 2.68E-04 | 1.80 |
| TC0100006805.hg.1 | RBP7 | retinol binding protein 7, cellular | 5.01E-03 | 1.80 |
| TC0600006870.hg.1 | SNRNP48 | small nuclear ribonucleoprotein, U11/U12 48KDa subunit | 1.11E-03 | 1.80 |
| TC0100018434.hg.1 | MYCBP | MYC binding protein | 1.33E-03 | 1.80 |
| TC0300007386.hg.1 | APEH | acylaminoacyl-peptide hydrolase | 7.39E-03 | 1.80 |
| TC1200010812.hg.1 | CALCOCO1 | calcium binding and coiled-coil domain 1 | 3.83E-03 | 1.80 |
| TC1200011099.hg.1 | GNS | glucosamine (N-acetyl)-6-sulfatase | 6.33E-04 | 1.80 |
| TC2100006976.hg.1 | ITSN1 | intersectin 1 | 9.49E-04 | 1.80 |
| TC1400010754.hg.1 | C14orf37 | chromosome 14 open reading frame 37 | 9.61E-05 | 1.80 |
| TC1600008169.hg.1 | PARD6A | par-6 family cell polarity regulator alpha | 3.52E-04 | 1.80 |
| TSUnmapped00000726.hg.1 | ZDHHC3 | zinc finger, DHHC-type containing 3 | 1.57E-03 | 1.80 |
| TC2100007964.hg.1 | SYNJ1 | synaptojanin 1 | 3.03E-03 | 1.79 |
| TC0900012276.hg.1 | SCAI | suppressor of cancer cell invasion | 4.69E-04 | 1.79 |
| TC0200015773.hg.1 | PTPRN | protein tyrosine phosphatase, receptor type, N | 3.12E-03 | 1.79 |
| TC0500008278.hg.1 | FER | fer (fps/fes related) tyrosine kinase | 5.24E-04 | 1.79 |
| TC1200006472.hg.1 | ADIPOR2 | adiponectin receptor 2 | 2.63E-03 | 1.79 |
| TC1200009031.hg.1 | RNFT2 | ring finger protein, transmembrane 2 | 5.99E-04 | 1.79 |
| TC0600007374.hg.1 | HIST1H2AI | histone cluster 1, H2ai | 9.98E-03 | 1.79 |
| TC1800008578.hg.1 | EPG5 | ectopic P-granules autophagy protein 5 homolog (C. elegans) | 7.16E-04 | 1.79 |
| TC0X00010877.hg.1 | FAM122B | family with sequence similarity 122B | 1.93E-03 | 1.79 |
| TC0400012781.hg.1 | STIM2 | stromal interaction molecule 2 | 2.19E-03 | 1.79 |
| TC0500007050.hg.1 | PDZD2 | PDZ domain containing 2 | 6.41E-04 | 1.79 |
| TC1600008498.hg.1 | VAT1L | vesicle amine transport 1-like | 7.09E-03 | 1.78 |
| TC2000008230.hg.1 | C20orf27 | chromosome 20 open reading frame 27 | 6.77E-03 | 1.78 |
| TC1600009199.hg.1 | SLX4 | SLX4 structure-specific endonuclease subunit | 1.55E-03 | 1.78 |
| TC1700010879.hg.1 | MAP3K14 | mitogen-activated protein kinase kinase kinase 14 | 7.01E-03 | 1.78 |
| TC0800007065.hg.1 | CDCA2 | cell division cycle associated 2 | 1.18E-03 | 1.78 |
| TC0100012675.hg.1 | ACOT7 | acyl-CoA thioesterase 7 | 6.26E-03 | 1.78 |
| TC0800010382.hg.1 | SNAI2 | snail family zinc finger 2 | 8.55E-04 | 1.78 |
| TC0300008046.hg.1 | ARL13B | ADP-ribosylation factor like GTPase 13B | 3.40E-04 | 1.78 |
| TC1400007950.hg.1 | CALM1 | calmodulin 1 (phosphorylase kinase, delta) | 2.66E-04 | 1.78 |
| TC0200009895.hg.1 | CERS6 | ceramide synthase 6 | 1.35E-03 | 1.78 |
| TC1500007067.hg.1 | CKMT1A | creatine kinase, mitochondrial 1A | 2.96E-04 | 1.78 |
| TC0900008795.hg.1 | RALGPS1 | Ral GEF with PH domain and SH3 binding motif 1 | 8.77E-04 | 1.78 |
| TC1500009238.hg.1 | FRMD5 | FERM domain containing 5 | 2.49E-03 | 1.78 |
| TC0100013072.hg.1 | ATP13A2 | ATPase type 13A2 | 2.86E-04 | 1.78 |
| TC0200015616.hg.1 | ERBB4 | erb-b2 receptor tyrosine kinase 4 | 8.36E-04 | 1.78 |

| | | | | |
|---|---|---|---|---|
| TC0300008369.hg.1 | ATP6V1A | ATPase, H+ transporting, lysosomal 70kDa, V1 subunit A | 1.26E-03 | 1.78 |
| TC1200007137.hg.1 | FGFR1OP2 | FGFR1 oncogene partner 2 | 5.55E-03 | 1.78 |
| TC0300013932.hg.1 | SRGAP3 | SLIT-ROBO Rho GTPase activating protein 3 | 1.12E-03 | 1.77 |
| TC1500009984.hg.1 | NPTN | neuroplastin | 4.70E-04 | 1.77 |
| TC1200009981.hg.1 | HEBP1 | heme binding protein 1 | 1.83E-03 | 1.77 |
| TC0100013634.hg.1 | TMEM54 | transmembrane protein 54 | 3.18E-03 | 1.77 |
| TC0100013763.hg.1 | POU3F1 | POU class 3 homeobox 1 | 3.26E-04 | 1.77 |
| TC1900009266.hg.1 | ZNF77 | zinc finger protein 77 | 5.83E-03 | 1.77 |
| TC1000006704.hg.1 | TAF3 | TATA box binding protein associated factor 3 | 1.52E-03 | 1.77 |
| TC1000007925.hg.1 | EIF4EBP2 | eukaryotic translation initiation factor 4E binding protein 2 | 8.27E-04 | 1.77 |
| TC1700009378.hg.1 | PITPNA | phosphatidylinositol transfer protein, alpha | 4.93E-04 | 1.77 |
| TC0900011259.hg.1 | HDHD3 | haloacid dehalogenase-like hydrolase domain containing 3 | 9.43E-03 | 1.77 |
| TC0300010908.hg.1 | ZDHHC3 | zinc finger, DHHC-type containing 3 | 2.62E-03 | 1.77 |
| TC1400007706.hg.1 | FOS | FBJ murine osteosarcoma viral oncogene homolog | 4.68E-03 | 1.77 |
| TC1800009203.hg.1 | PQLC1 | PQ loop repeat containing 1 | 3.06E-04 | 1.77 |
| TC0900009673.hg.1 | MLLT3 | myeloid/lymphoid or mixed-lineage leukemia; translocated to, 3 | 4.89E-03 | 1.77 |
| TC1100006906.hg.1 | FAR1 | fatty acyl CoA reductase 1 | 9.49E-04 | 1.77 |
| TC0600013010.hg.1 | MAN1A1 | mannosidase, alpha, class 1A, member 1 | 1.84E-03 | 1.77 |
| TC0800011679.hg.1 | ATAD2 | ATPase family, AAA domain containing 2 | 5.20E-03 | 1.77 |
| TC0600010040.hg.1 | FNDC1 | fibronectin type III domain containing 1 | 8.83E-04 | 1.77 |
| TC0600008127.hg.1 | SLC29A1 | solute carrier family 29 (equilibrative nucleoside transporter), member 1 | 3.01E-03 | 1.77 |
| TSUnmapped00000569.hg.1 | ZDHHC3 | zinc finger, DHHC-type containing 3 | 8.04E-03 | 1.77 |
| TC0100015194.hg.1 | SORT1 | sortilin 1 | 1.45E-03 | 1.77 |
| TC0X00008674.hg.1 | AFF2 | AF4/FMR2 family, member 2 | 7.15E-03 | 1.77 |
| TC0200014717.hg.1 | WDSUB1 | WD repeat, sterile alpha motif and U-box domain containing 1 | 4.23E-03 | 1.77 |
| TC0100013302.hg.1 | GALE | UDP-galactose-4-epimerase | 8.37E-03 | 1.77 |
| TC0600011138.hg.1 | HIST1H1D | histone cluster 1, H1d | 1.46E-03 | 1.77 |
| TSUnmapped00000178.hg.1 | SLC16A1 | solute carrier family 16 (monocarboxylate transporter), member 1 | 5.71E-03 | 1.77 |
| TC0600009343.hg.1 | NKAIN2 | Na+/K+ transporting ATPase interacting 2 | 2.78E-03 | 1.76 |
| TC0900010971.hg.1 | TBC1D2 | TBC1 domain family, member 2 | 3.58E-03 | 1.76 |
| TC1700012089.hg.1 | DCXR | dicarbonyl/L-xylulose reductase | 1.85E-03 | 1.76 |
| TC0X00007936.hg.1 | CENPI | centromere protein I | 2.28E-03 | 1.76 |
| TC0600007799.hg.1 | FANCE | Fanconi anemia complementation group E | 2.89E-03 | 1.76 |
| TC0100009408.hg.1 | PIFO | primary cilia formation | 1.60E-03 | 1.76 |
| TC1000009248.hg.1 | FANK1 | fibronectin type III and ankyrin repeat domains 1 | 8.61E-04 | 1.76 |
| TC1300009012.hg.1 | THSD1 | thrombospondin type 1 domain containing 1 | 2.25E-03 | 1.76 |
| TC1700007677.hg.1 | DUSP14 | dual specificity phosphatase 14 | 3.66E-03 | 1.76 |
| HTA2-neg-47420655_st | NA | NA | 1.47E-03 | 1.76 |
| TC0100016350.hg.1 | CCDC181 | coiled-coil domain containing 181 | 8.29E-03 | 1.76 |
| TC0600009697.hg.1 | PHACTR2 | Jeck2013 ALT_ACCEPTOR, ALT_DONOR, coding, INTERNAL, intronic best transcript NM_001100164 | 8.78E-03 | 1.76 |
| TC1900007063.hg.1 | ZNF788 | zinc finger family member 788 | 3.60E-03 | 1.76 |
| TC0300007610.hg.1 | PXK | PX domain containing serine/threonine kinase | 1.17E-04 | 1.76 |
| TC0500011260.hg.1 | SERINC5 | serine incorporator 5 | 1.96E-04 | 1.76 |
| TC0900008803.hg.1 | SLC2A8 | solute carrier family 2 (facilitated glucose transporter), member 8 | 8.36E-04 | 1.76 |
| TC0500008602.hg.1 | LYRM7 | LYR motif containing 7 | 6.13E-04 | 1.76 |
| TC0500011702.hg.1 | STARD4 | StAR-related lipid transfer domain containing 4 | 8.21E-04 | 1.76 |
| TC0200012489.hg.1 | PIGF | phosphatidylinositol glycan anchor biosynthesis class F | 2.40E-04 | 1.76 |
| 23071874 | NA | NA | 3.49E-04 | 1.75 |
| TC1200010968.hg.1 | DDIT3 | DNA-damage-inducible transcript 3 | 9.99E-04 | 1.75 |
| TC0800009854.hg.1 | BIN3-IT1 | BIN3 intronic transcript 1 | 3.74E-03 | 1.75 |
| TC2200006827.hg.1 | GNAZ | guanine nucleotide binding protein (G protein), alpha z polypeptide | 4.13E-03 | 1.75 |
| TSUnmapped00000068.hg.1 | ZNF780A | zinc finger protein 780A | 9.88E-03 | 1.75 |
| TC1400009214.hg.1 | BMP4 | bone morphogenetic protein 4 | 7.06E-04 | 1.75 |
| TC1200011908.hg.1 | GPN3 | GPN-loop GTPase 3 | 1.80E-03 | 1.75 |
| TC0600008632.hg.1 | BCKDHB | branched chain keto acid dehydrogenase E1, beta polypeptide | 6.59E-04 | 1.75 |
| TC0900011655.hg.1 | ZER1 | zyg-11 related, cell cycle regulator | 5.79E-03 | 1.75 |
| TC0100016567.hg.1 | TOR1AIP2 | torsin A interacting protein 2 | 1.46E-03 | 1.75 |
| TC1900006586.hg.1 | TMPRSS9 | transmembrane protease, serine 9 | 9.55E-03 | 1.75 |
| TC0900009776.hg.1 | MOB3B | MOB kinase activator 3B | 5.63E-03 | 1.75 |
| TC1400007500.hg.1 | ARG2 | arginase 2 | 3.42E-03 | 1.75 |
| TC1600010000.hg.1 | STX1B | syntaxin 1B | 3.89E-03 | 1.75 |
| TC0900010808.hg.1 | BICD2 | bicaudal D homolog 2 (Drosophila) | 2.33E-03 | 1.75 |
| TC0600009800.hg.1 | GINM1 | glycoprotein integral membrane 1 | 3.90E-04 | 1.75 |
| TC1200008275.hg.1 | SYT1 | synaptotagmin I | 2.46E-03 | 1.75 |
| TC0600009102.hg.1 | FIG4 | FIG4 phosphoinositide 5-phosphatase | 3.69E-04 | 1.75 |
| TC0700012099.hg.1 | RASA4 | Salzman2013 ALT_ACCEPTOR, ALT_DONOR, coding, INTERNAL, intronic best transcript NM_006989 | 3.28E-03 | 1.75 |
| TC0200014204.hg.1 | HS6ST1 | heparan sulfate 6-O-sulfotransferase 1 | 3.83E-04 | 1.75 |
| TC1700006702.hg.1 | TXNDC17 | thioredoxin domain containing 17 | 1.36E-03 | 1.75 |
| TC0800012447.hg.1 | AZIN1 | antizyme inhibitor 1 | 1.36E-03 | 1.74 |
| TC0X00011211.hg.1 | G6PD | glucose-6-phosphate dehydrogenase | 7.10E-03 | 1.74 |
| TC1500010845.hg.1 | OTUD7A | OTU deubiquitinase 7A | 1.42E-03 | 1.74 |
| TC2000006571.hg.1 | SMOX | spermine oxidase | 1.43E-03 | 1.74 |
| TC0500009161.hg.1 | SAP30L | SAP30-like | 5.92E-03 | 1.74 |
| TC0300009856.hg.1 | IL1RAP | interleukin 1 receptor accessory protein | 5.36E-03 | 1.74 |
| TC1700011208.hg.1 | TRIM25 | tripartite motif containing 25 | 1.11E-03 | 1.74 |
| TC1500010128.hg.1 | TBC1D2B | TBC1 domain family, member 2B | 4.03E-03 | 1.74 |
| HTA2-neg-47423726_st | NA | NA | 3.37E-03 | 1.74 |
| TC0700013525.hg.1 | FAM126A | family with sequence similarity 126, member A | 6.89E-03 | 1.74 |
| TSUnmapped00000085.hg.1 | CCDC84 | coiled-coil domain containing 84 | 3.27E-03 | 1.74 |
| TC1900008259.hg.1 | ZNF233 | zinc finger protein 233 | 1.29E-03 | 1.74 |
| TC0X00009057.hg.1 | MID1 | midline 1 | 4.24E-04 | 1.74 |
| TC1500010160.hg.1 | CTSH | cathepsin H | 2.66E-03 | 1.74 |
| 23071987 | NA | NA | 2.65E-04 | 1.74 |
| TC0900011379.hg.1 | MEGF9 | multiple EGF-like-domains 9 | 6.83E-04 | 1.74 |
| TC1200010011.hg.1 | HIST4H4 | histone cluster 4, H4 | 6.26E-03 | 1.74 |
| TC1400010331.hg.1 | XRCC3 | X-ray repair complementing defective repair in Chinese hamster cells 3 | 1.30E-03 | 1.74 |
| TC0800007351.hg.1 | TACC1 | transforming, acidic coiled-coil containing protein 1 | 8.48E-04 | 1.74 |
| 23071967 | NA | NA | 2.02E-04 | 1.74 |
| TC0900012277.hg.1 | AK1 | adenylate kinase 1 | 9.06E-04 | 1.74 |
| TC0700010581.hg.1 | HIBADH | 3-hydroxyisobutyrate dehydrogenase | 3.24E-03 | 1.74 |
| TC1200008814.hg.1 | ANKRD13A | ankyrin repeat domain 13A | 2.28E-04 | 1.74 |
| TC0100009241.hg.1 | S1PR1 | sphingosine-1-phosphate receptor 1 | 7.88E-04 | 1.74 |
| 23075636 | NA | NA | 3.05E-03 | 1.74 |
| 23076269 | NA | NA | 3.05E-03 | 1.74 |
| TC1900006848.hg.1 | CAMSAP3 | calmodulin regulated spectrin-associated protein family, member 3 | 5.66E-03 | 1.74 |
| TC2000009960.hg.1 | SRXN1 | sulfiredoxin 1 | 6.66E-03 | 1.73 |
| TC1600010593.hg.1 | CMTM4 | CKLF-like MARVEL transmembrane domain containing 4 | 4.89E-04 | 1.73 |
| TC1100010491.hg.1 | FBXO3 | F-box protein 3 | 4.43E-03 | 1.73 |
| TC1400010615.hg.1 | MNAT1 | MNAT CDK-activating kinase assembly factor 1 | 7.80E-04 | 1.73 |
| TC1400009705.hg.1 | PGF | placental growth factor | 4.77E-04 | 1.73 |
| TC1200007835.hg.1 | COQ10A | coenzyme Q10A | 1.20E-03 | 1.73 |
| TC0200011107.hg.1 | EFHD1 | EF-hand domain family member D1 | 3.68E-03 | 1.73 |
| HTA2-pos-PSR02023981.hg.1 | NA | NA | 7.91E-03 | 1.73 |
| TC1300008443.hg.1 | LNX2 | ligand of numb-protein X 2 | 5.08E-03 | 1.73 |
| TC0100009958.hg.1 | CGN | cingulin | 8.82E-03 | 1.73 |
| TC1900007366.hg.1 | MPV17L2 | MPV17 mitochondrial membrane protein-like 2 | 1.40E-03 | 1.73 |
| TC1000008743.hg.1 | SUFU | Transcript Identified by AceView, Entrez Gene ID(s) 51684 | 9.10E-04 | 1.73 |
| TC1000011660.hg.1 | MGEA5 | Salzman2013 ALT_ACCEPTOR, ALT_DONOR, coding, INTERNAL, intronic best transcript NM_012215 | 8.00E-03 | 1.73 |
| 23075808 | NA | NA | 2.01E-03 | 1.73 |
| TC1400010716.hg.1 | HOMEZ | homeobox and leucine zipper encoding | 1.64E-03 | 1.73 |
| TC2200006678.hg.1 | SNAP29 | synaptosome associated protein 29kDa | 2.91E-03 | 1.72 |
| TC0700008495.hg.1 | BUD31 | Transcript Identified by AceView, Entrez Gene ID(s) 8896 | 3.94E-03 | 1.72 |
| TC0700008292.hg.1 | STEAP1 | six transmembrane epithelial antigen of the prostate 1 | 9.99E-03 | 1.72 |
| TC1200007594.hg.1 | ASIC1 | acid sensing ion channel 1 | 3.16E-03 | 1.72 |
| HTA2-pos-3529035_st | NA | NA | 4.43E-03 | 1.72 |
| HTA2-pos-PSR19012462.hg.1 | NA | NA | 2.66E-03 | 1.72 |
| TC0100018480.hg.1 | NBPF14 | neuroblastoma breakpoint family, member 14 | 2.06E-04 | 1.72 |
| TC0200013765.hg.1 | UXS1 | UDP-glucuronate decarboxylase 1 | 9.23E-04 | 1.72 |
| TC1700012307.hg.1 | TSEN54 | TSEN54 tRNA splicing endonuclease subunit | 3.02E-04 | 1.72 |

| | | | | |
|---|---|---|---|---|
| TC0300011550.hg.1 | PDZRN3 | PDZ domain containing ring finger 3 | 4.64E-03 | 1.72 |
| TC0500012294.hg.1 | PCDH1 | protocadherin 1 | 8.99E-04 | 1.72 |
| TC1700011910.hg.1 | CYTH1 | cytohesin 1 | 1.35E-03 | 1.72 |
| TC1900006602.hg.1 | ZNF555 | zinc finger protein 555 | 9.45E-04 | 1.72 |
| 23071900 | NA | NA | 3.56E-04 | 1.72 |
| TC0200011720.hg.1 | ODC1 | ornithine decarboxylase 1 | 4.88E-03 | 1.72 |
| TC1600011315.hg.1 | FAM234A | family with sequence similarity 234, member A | 8.91E-04 | 1.72 |
| TC0700006512.hg.1 | INTS1 | Memczak2013 ANTISENSE, CDS, coding, INTERNAL best transcript NM_001080453 | 2.77E-03 | 1.72 |
| TC1700007189.hg.1 | ALDH3A2 | aldehyde dehydrogenase 3 family, member A2 | 2.17E-03 | 1.72 |
| TC0100016122.hg.1 | IGSF8 | immunoglobulin superfamily, member 8 | 1.22E-03 | 1.72 |
| TC0X00006816.hg.1 | PDK3 | pyruvate dehydrogenase kinase, isozyme 3 | 1.66E-03 | 1.72 |
| TC1500008983.hg.1 | SLC12A6 | solute carrier family 12 (potassium/chloride transporter), member 6 | 7.36E-03 | 1.72 |
| TC1600009989.hg.1 | CCDC189 | coiled-coil domain containing 189 | 3.15E-03 | 1.72 |
| TC1100008010.hg.1 | CDC42EP2 | CDC42 effector protein (Rho GTPase binding) 2 | 9.86E-04 | 1.72 |
| TC1900009584.hg.1 | ZNF560 | zinc finger protein 560 | 6.08E-03 | 1.72 |
| TC1000010982.hg.1 | ASCC1 | Transcript Identified by AceView, Entrez Gene ID(s) 51008 | 3.45E-03 | 1.72 |
| TC1600010685.hg.1 | SMPD3 | sphingomyelin phosphodiesterase 3, neutral membrane (neutral sphingomyelinase II) | 4.72E-03 | 1.72 |
| TC1200011573.hg.1 | NR2C1 | nuclear receptor subfamily 2, group C, member 1 | 8.76E-03 | 1.72 |
| TC1000007132.hg.1 | BAMBI | BMP and activin membrane-bound inhibitor | 2.02E-03 | 1.72 |
| TC0100006954.hg.1 | KAZN | kazrin, periplakin interacting protein | 9.43E-03 | 1.71 |
| 23075955 | NA | NA | 3.86E-03 | 1.71 |
| TC1400009829.hg.1 | CEP128 | centrosomal protein 128kDa | 7.84E-03 | 1.71 |
| TC0300013853.hg.1 | ST3GAL6 | ST3 beta-galactoside alpha-2,3-sialyltransferase 6 | 3.59E-03 | 1.71 |
| TC0200006738.hg.1 | LPIN1 | Transcript Identified by AceView, Entrez Gene ID(s) 23175 | 5.26E-03 | 1.71 |
| TC0300009131.hg.1 | HPS3 | Hermansky-Pudlak syndrome 3 | 2.02E-03 | 1.71 |
| TC1900011867.hg.1 | TMEM205 | transmembrane protein 205 | 7.15E-04 | 1.71 |
| TC0900009489.hg.1 | GLDC | glycine dehydrogenase (decarboxylating) | 5.19E-03 | 1.71 |
| TC0200007419.hg.1 | CAMKMT | calmodulin-lysine N-methyltransferase | 3.81E-03 | 1.71 |
| TC1100008899.hg.1 | CEP126 | centrosomal protein 126kDa | 3.03E-03 | 1.71 |
| TC0600008255.hg.1 | EFHC1 | EF-hand domain (C-terminal) containing 1 | 2.73E-03 | 1.71 |
| TC1900010067.hg.1 | PBX4 | pre-B-cell leukemia homeobox 4 | 5.71E-04 | 1.71 |
| TC0700009020.hg.1 | SND1 | staphylococcal nuclease and tudor domain containing 1 | 7.46E-04 | 1.71 |
| TC2200007109.hg.1 | DEPDC5 | DEP domain containing 5 | 4.24E-04 | 1.71 |
| TC0800012176.hg.1 | PLEC | plectin | 2.76E-03 | 1.71 |
| TC0500012826.hg.1 | SH3PXD2B | SH3 and PX domains 2B | 1.84E-03 | 1.71 |
| TC0500008054.hg.1 | POLR3G | polymerase (RNA) III (DNA directed) polypeptide G (32kD) | 7.86E-03 | 1.71 |
| TC1700009417.hg.1 | SMG6 | SMG6 nonsense mediated mRNA decay factor | 1.29E-03 | 1.71 |
| TC1700008769.hg.1 | KCNJ2 | potassium channel, inwardly rectifying subfamily J, member 2 | 1.34E-03 | 1.71 |
| TC0700007398.hg.1 | ZMIZ2 | zinc finger, MIZ-type containing 2 | 1.18E-03 | 1.71 |
| TC1100012966.hg.1 | MICAL2 | microtubule associated monooxygenase, calponin and LIM domain containing 2 | 1.35E-03 | 1.71 |
| TC1600006760.hg.1 | UBN1 | ubinuclein 1 | 5.22E-03 | 1.71 |
| TC0100018233.hg.1 | FPGT | fucose-1-phosphate guanylyltransferase | 3.01E-03 | 1.70 |
| TC0500011408.hg.1 | TMEM161B | transmembrane protein 161B | 1.34E-03 | 1.70 |
| TC1500007034.hg.1 | SNAP23 | synaptosome associated protein 23kDa | 5.23E-03 | 1.70 |
| TC0100009324.hg.1 | FAM102B | family with sequence similarity 102, member B | 9.34E-03 | 1.70 |
| TC1400010630.hg.1 | TTLL5 | tubulin tyrosine ligase-like family member 5 | 5.04E-03 | 1.70 |
| TC1700009333.hg.1 | NXN | nucleoredoxin | 1.05E-03 | 1.70 |
| TC0100017617.hg.1 | HIST3H2A | histone cluster 3, H2a | 7.49E-04 | 1.70 |
| HTA2-neg-47419773_st | NA | NA | 2.37E-03 | 1.70 |
| TSUnmapped00000732.hg.1 | SLC37A4 | solute carrier family 37 (glucose-6-phosphate transporter), member 4 [Source:HGNC Symbol;Acc:HGNC:4061] | 8.24E-03 | 1.70 |
| TC0200015876.hg.1 | SERPINE2 | serpin peptidase inhibitor, clade E (nexin, plasminogen activator inhibitor type 1), member 2 | 4.49E-04 | 1.70 |
| HTA2-neg-47420711_st | NA | NA | 9.17E-03 | 1.70 |
| TC0400012381.hg.1 | NEK1 | NIMA-related kinase 1 | 5.81E-03 | 1.70 |
| TC0100013837.hg.1 | RIMS3 | regulating synaptic membrane exocytosis 3 | 9.68E-04 | 1.70 |
| TC0200010624.hg.1 | PIKFYVE | phosphoinositide kinase, FYVE finger containing | 4.39E-03 | 1.70 |
| TC0100009969.hg.1 | RIIAD1 | regulatory subunit of type II PKA R-subunit (RIIa) domain containing 1 | 1.45E-03 | 1.70 |
| TC0700011066.hg.1 | COBL | cordon-bleu WH2 repeat protein | 4.42E-04 | 1.70 |
| TC0100010798.hg.1 | QSOX1 | quiescin Q6 sulfhydryl oxidase 1 | 1.70E-03 | 1.70 |
| TC1200006670.hg.1 | CLSTN3 | calsyntenin 3 | 2.48E-03 | 1.70 |
| TC1700011815.hg.1 | MXRA7 | matrix-remodelling associated 7 | 9.72E-03 | 1.70 |
| TC0900008887.hg.1 | TBC1D13 | TBC1 domain family, member 13 | 2.09E-03 | 1.70 |
| TC0900006937.hg.1 | ACO1 | aconitase 1, soluble | 6.80E-04 | 1.70 |
| TC1900006859.hg.1 | EVI5L | ecotropic viral integration site 5-like | 2.77E-04 | 1.69 |
| TC0200016587.hg.1 | PLCL1 | phospholipase C-like 1 | 3.19E-03 | 1.69 |
| TC0700008367.hg.1 | PEG10 | paternally expressed 10 | 8.46E-03 | 1.69 |
| TC0300008948.hg.1 | ESYT3 | extended synaptotagmin-like protein 3 | 4.56E-04 | 1.69 |
| TC0900011461.hg.1 | STRBP | spermatid perinuclear RNA binding protein | 1.06E-03 | 1.69 |
| TC1900011172.hg.1 | VRK3 | vaccinia related kinase 3 | 8.07E-04 | 1.69 |
| TC0400009856.hg.1 | LYAR | Ly1 antibody reactive | 5.00E-03 | 1.69 |
| TC1500010886.hg.1 | CALML4 | calmodulin-like 4 | 1.79E-03 | 1.69 |
| TC0900008684.hg.1 | GPR21 | G protein-coupled receptor 21 | 8.89E-04 | 1.69 |
| TC1200012622.hg.1 | SLC48A1 | solute carrier family 48 (heme transporter), member 1 | 2.67E-04 | 1.69 |
| TSUnmapped00000530.hg.1 | ATG16L1 | autophagy related 16-like 1 | 6.13E-03 | 1.69 |
| TC2200009324.hg.1 | DRICH1 | aspartate-rich 1 | 2.40E-03 | 1.69 |
| TC1700009274.hg.1 | FN3KRP | fructosamine 3 kinase related protein | 3.26E-03 | 1.69 |
| TC1200011968.hg.1 | HECTD4 | HECT domain containing E3 ubiquitin protein ligase 4 | 4.21E-03 | 1.69 |
| TC0400009132.hg.1 | ETFDH | electron-transferring-flavoprotein dehydrogenase | 6.71E-04 | 1.68 |
| TC0200007028.hg.1 | GAREM2 | GRB2 associated regulator of MAPK1 2 | 2.30E-03 | 1.68 |
| TC2000008130.hg.1 | TBC1D20 | TBC1 domain family, member 20 | 5.97E-03 | 1.68 |
| TC2100006980.hg.1 | ITSN1 | Transcript Identified by AceView, Entrez Gene ID(s) 6453 | 1.36E-03 | 1.68 |
| TC0800008197.hg.1 | OTUD6B | OTU domain containing 6B | 1.46E-03 | 1.68 |
| TC0100011245.hg.1 | ADORA1 | adenosine A1 receptor | 4.27E-04 | 1.68 |
| TC0300011939.hg.1 | IFT57 | intraflagellar transport 57 | 3.35E-03 | 1.68 |
| TC1100009245.hg.1 | CBL | Cbl proto-oncogene, E3 ubiquitin protein ligase | 3.52E-03 | 1.68 |
| TC2000009780.hg.1 | TCFL5 | transcription factor-like 5 (basic helix-loop-helix) | 5.09E-03 | 1.68 |
| TC0600007616.hg.1 | HSPA1B | heat shock 70kDa protein 1B | 5.60E-04 | 1.68 |
| TC1100010661.hg.1 | CHST1 | carbohydrate (keratan sulfate Gal-6) sulfotransferase 1 | 1.38E-03 | 1.68 |
| TC0100011552.hg.1 | VASH2 | vasohibin 2 | 2.21E-03 | 1.68 |
| TC0600008670.hg.1 | PRSS35 | protease, serine 35 | 2.23E-04 | 1.68 |
| TC0600008569.hg.1 | MYO6 | myosin VI | 6.41E-03 | 1.68 |
| TC0100012846.hg.1 | DFFA | DNA fragmentation factor, 45kDa, alpha polypeptide | 1.50E-03 | 1.68 |
| TC1200010865.hg.1 | ITGA7 | integrin alpha 7 | 6.24E-03 | 1.68 |
| TC0400011518.hg.1 | INTS12 | integrator complex subunit 12 | 2.46E-03 | 1.68 |
| TC0600013228.hg.1 | SLC2A12 | solute carrier family 2 (facilitated glucose transporter), member 12 | 6.97E-03 | 1.68 |
| TC0100016631.hg.1 | RGS16 | regulator of G-protein signaling 16 | 9.97E-04 | 1.68 |
| TC0200016746.hg.1 | TTC21B | tetratricopeptide repeat domain 21B | 4.60E-03 | 1.68 |
| TC0300014050.hg.1 | NPHP3 | nephronophthisis 3 (adolescent) | 7.19E-03 | 1.68 |
| TC2000007132.hg.1 | DNMT3B | DNA (cytosine-5-)-methyltransferase 3 beta | 4.19E-03 | 1.68 |
| TC0200009091.hg.1 | RALB | v-ral simian leukemia viral oncogene homolog B | 6.37E-03 | 1.68 |
| TC0100009333.hg.1 | GPSM2 | G-protein signaling modulator 2 | 4.08E-03 | 1.68 |
| TC1000010993.hg.1 | MICU1 | mitochondrial calcium uptake 1 | 1.59E-03 | 1.68 |
| TC0100007240.hg.1 | NBPF3 | neuroblastoma breakpoint family, member 3 | 2.05E-03 | 1.68 |
| TC0500009580.hg.1 | SIMC1 | SUMO-interacting motifs containing 1 | 2.09E-03 | 1.67 |
| TC1900007447.hg.1 | ZNF101 | zinc finger protein 101 | 6.00E-03 | 1.67 |
| TC0200010511.hg.1 | NBEAL1 | neurobeachin like 1 | 2.69E-03 | 1.67 |
| TC1600011409.hg.1 | PDP2 | pyruvate dehyrogenase phosphatase catalytic subunit 2 | 5.98E-04 | 1.67 |
| TC1700011956.hg.1 | TBC1D16 | TBC1 domain family, member 16 | 2.73E-04 | 1.67 |
| TC0400011203.hg.1 | HPSE | heparanase | 7.91E-04 | 1.67 |
| TC0300010223.hg.1 | PRRT3 | proline-rich transmembrane protein 3 | 3.54E-04 | 1.67 |
| TC0300007445.hg.1 | GRM2 | glutamate receptor, metabotropic 2 | 1.22E-03 | 1.67 |
| TC1700008014.hg.1 | ADAM11 | ADAM metallopeptidase domain 11 | 1.30E-03 | 1.67 |
| TC1100010410.hg.1 | BDNF | brain-derived neurotrophic factor | 5.88E-03 | 1.67 |
| TC0X00009215.hg.1 | EIF1AX | eukaryotic translation initiation factor 1A, X-linked | 6.46E-03 | 1.67 |
| TC1500008772.hg.1 | GABRB3 | gamma-aminobutyric acid (GABA) A receptor, beta 3 | 6.86E-03 | 1.67 |
| TC0900011013.hg.1 | ERP44 | endoplasmic reticulum protein 44 | 4.17E-03 | 1.67 |
| TC1400006521.hg.1 | PNP | purine nucleoside phosphorylase | 7.39E-04 | 1.67 |
| TC1500010915.hg.1 | GOLGA6L5P | golgin A6 family-like 5, pseudogene | 6.02E-03 | 1.67 |
| TC0900010580.hg.1 | GKAP1 | G kinase anchoring protein 1 | 2.59E-03 | 1.67 |
| TC1100012811.hg.1 | NFRKB | nuclear factor related to kappaB binding protein | 5.44E-04 | 1.67 |

| | | | | |
|---|---|---|---|---|
| TC0200012809.hg.1 | PELI1 | pellino E3 ubiquitin protein ligase 1 | 2.81E-03 | 1.67 |
| TC0100010735.hg.1 | RASAL2 | RAS protein activator like 2 | 1.13E-03 | 1.67 |
| TC2000009890.hg.1 | ANKEF1 | ankyrin repeat and EF-hand domain containing 1 | 6.25E-03 | 1.67 |
| TC1700012184.hg.1 | CHRNB1 | cholinergic receptor, nicotinic beta 1 | 7.66E-03 | 1.67 |
| 23074909 | NA | NA | 5.03E-03 | 1.67 |
| TC2000008023.hg.1 | SLCO4A1 | solute carrier organic anion transporter family, member 4A1 | 4.87E-03 | 1.67 |
| TC0100012002.hg.1 | KIAA1804 | mixed lineage kinase 4 | 9.75E-04 | 1.67 |
| TC1500007004.hg.1 | RTF1 | RTF1 homolog, Paf1/RNA polymerase II complex component | 6.93E-03 | 1.67 |
| TC1700009256.hg.1 | NARF | nuclear prelamin A recognition factor | 1.46E-03 | 1.67 |
| TC0100014457.hg.1 | JAK1 | Janus kinase 1 | 9.52E-04 | 1.67 |
| TC0600007375.hg.1 | HIST1H3H | histone cluster 1, H3h | 9.84E-03 | 1.67 |
| TC0300011183.hg.1 | NEK4 | NIMA-related kinase 4 | 6.57E-04 | 1.67 |
| TC1700008539.hg.1 | MRC2 | mannose receptor, C type 2 | 7.04E-04 | 1.67 |
| HTA2-pos-PSR03025032.hg.1 | NA | NA | 3.42E-03 | 1.67 |
| TC1600011168.hg.1 | ZCCHC14 | zinc finger, CCHC domain containing 14 | 2.81E-03 | 1.66 |
| TC0900011120.hg.1 | KLF4 | Kruppel-like factor 4 (gut) | 1.20E-03 | 1.66 |
| TC0100018478.hg.1 | NBPF10 | neuroblastoma breakpoint family, member 10 | 7.75E-04 | 1.66 |
| TC1600006941.hg.1 | SNX29 | sorting nexin 29 | 2.58E-03 | 1.66 |
| TC0100007691.hg.1 | KIAA1522 | KIAA1522 | 5.39E-04 | 1.66 |
| TC1500008521.hg.1 | MEF2A | myocyte enhancer factor 2A | 6.81E-04 | 1.66 |
| TC0X00010351.hg.1 | TAF7L | TATA box binding protein associated factor 7 like | 5.21E-03 | 1.66 |
| TC2200007910.hg.1 | MICAL3 | Transcript Identified by AceView, Entrez Gene ID(s) 57553 | 1.76E-03 | 1.66 |
| TC0800007738.hg.1 | SDCBP | syndecan binding protein | 1.03E-03 | 1.66 |
| TC1200008726.hg.1 | TCP11L2 | t-complex 11, testis-specific-like 2 | 1.50E-03 | 1.66 |
| TC1100009618.hg.1 | GLB1L2 | galactosidase beta 1 like 2 | 6.26E-03 | 1.66 |
| TC0300010557.hg.1 | NEK10 | NIMA-related kinase 10 | 9.57E-03 | 1.66 |
| TC0600010066.hg.1 | IGF2R | insulin-like growth factor 2 receptor | 7.12E-04 | 1.66 |
| TC0600011635.hg.1 | FKBP5 | FK506 binding protein 5 | 1.00E-02 | 1.66 |
| TC1600008189.hg.1 | PLA2G15 | phospholipase A2, group XV | 4.27E-03 | 1.66 |
| TC0900008463.hg.1 | HSDL2 | hydroxysteroid dehydrogenase like 2 | 3.49E-03 | 1.66 |
| TC1300006774.hg.1 | MEDAG | mesenteric estrogen-dependent adipogenesis | 9.53E-04 | 1.66 |
| TC0300006618.hg.1 | SYN2 | synapsin II | 2.27E-03 | 1.66 |
| TC0700013604.hg.1 | UPK3BL | uroplakin 3B-like | 8.38E-04 | 1.66 |
| TC0200008223.hg.1 | DNAH6 | dynein, axonemal, heavy chain 6 | 6.51E-03 | 1.65 |
| TC0300007406.hg.1 | SEMA3F | sema domain, immunoglobulin domain (Ig), short basic domain, secreted, (semaphorin) 3F | 3.63E-03 | 1.65 |
| TC1900009808.hg.1 | ADGRL1 | adhesion G protein-coupled receptor L1 | 2.83E-03 | 1.65 |
| TC1500010712.hg.1 | GOLGA8N | golgin A8 family, member N | 3.21E-03 | 1.65 |
| TC1600011047.hg.1 | HSDL1 | hydroxysteroid dehydrogenase like 1 | 1.14E-03 | 1.65 |
| TC1200011846.hg.1 | CORO1C | coronin, actin binding protein, 1C | 6.26E-03 | 1.65 |
| TC0100017073.hg.1 | SLC45A3 | solute carrier family 45, member 3 | 8.67E-04 | 1.65 |
| HTA2-neg-47422238_st | NA | NA | 7.05E-03 | 1.65 |
| TC1600010864.hg.1 | WDR59 | WD repeat domain 59 | 1.66E-03 | 1.65 |
| TC0100018367.hg.1 | CHRM3 | cholinergic receptor, muscarinic 3 | 7.33E-03 | 1.65 |
| TC1500007851.hg.1 | ARID3B | AT rich interactive domain 3B (BRIGHT-like) | 2.75E-03 | 1.65 |
| TC1900007328.hg.1 | SLC27A1 | solute carrier family 27 (fatty acid transporter), member 1 | 3.01E-03 | 1.65 |
| TC1200011574.hg.1 | FGD6 | FYVE, RhoGEF and PH domain containing 6 | 2.53E-03 | 1.65 |
| TC0200014728.hg.1 | LY75-CD302 | LY75-CD302 readthrough | 3.69E-03 | 1.65 |
| TC0100011219.hg.1 | PPP1R12B | protein phosphatase 1, regulatory subunit 12B | 5.16E-04 | 1.65 |
| TC0100009048.hg.1 | RPL5 | Memczak2013 ALT_ACCEPTOR, ALT_DONOR, coding, INTERNAL, intronic best transcript NM_000969 | 4.11E-03 | 1.65 |
| TC1200011321.hg.1 | OSBPL8 | oxysterol binding protein-like 8 | 8.33E-04 | 1.65 |
| TC1500006643.hg.1 | GABRA5 | gamma-aminobutyric acid (GABA) A receptor, alpha 5 | 6.32E-03 | 1.65 |
| 23071983 | NA | NA | 4.10E-03 | 1.64 |
| TC1500010236.hg.1 | GOLGA6L10 | golgin A6 family-like 10 | 2.94E-03 | 1.64 |
| TC0100013000.hg.1 | CASP9 | caspase 9 | 6.82E-03 | 1.64 |
| TC0500009268.hg.1 | TTC1 | tetratricopeptide repeat domain 1 | 5.83E-03 | 1.64 |
| TC1600007955.hg.1 | MT3 | metallothionein 3 | 4.21E-03 | 1.64 |
| HTA2-pos-PSR02023963.hg.1 | NA | NA | 9.49E-04 | 1.64 |
| TC0600006659.hg.1 | BPHL | biphenyl hydrolase-like (serine hydrolase) | 1.98E-03 | 1.64 |
| TC0300006994.hg.1 | FBXL2 | F-box and leucine-rich repeat protein 2 | 3.32E-03 | 1.64 |
| TC2100006951.hg.1 | OLIG1 | oligodendrocyte transcription factor 1 | 9.05E-04 | 1.64 |
| TC0800009921.hg.1 | KCTD9 | potassium channel tetramerization domain containing 9 | 1.05E-03 | 1.64 |
| TC1700008452.hg.1 | GDPD1 | glycerophosphodiester phosphodiesterase domain containing 1 | 2.04E-03 | 1.64 |
| TC0600012536.hg.1 | SRSF12 | serine/arginine-rich splicing factor 12 | 8.31E-03 | 1.64 |
| TC1700008033.hg.1 | HEXIM1 | hexamethylene bis-acetamide inducible 1 | 2.61E-03 | 1.64 |
| TC0500006479.hg.1 | NKD2 | naked cuticle homolog 2 (Drosophila) | 3.43E-03 | 1.64 |
| TC2200006630.hg.1 | TANGO2 | transport and golgi organization 2 homolog | 4.92E-04 | 1.64 |
| TC1400008462.hg.1 | C14orf79 | chromosome 14 open reading frame 79 | 8.11E-03 | 1.64 |
| TC1700008467.hg.1 | VMP1 | vacuole membrane protein 1 | 8.69E-04 | 1.63 |
| TC1600007317.hg.1 | KIAA0556 | KIAA0556 | 9.19E-04 | 1.63 |
| TC0100016917.hg.1 | CSRP1 | cysteine and glycine-rich protein 1 | 5.96E-03 | 1.63 |
| TC0500009609.hg.1 | FGFR4 | fibroblast growth factor receptor 4 | 1.31E-03 | 1.63 |
| TC0X00010670.hg.1 | TMEM255A | transmembrane protein 255A | 8.51E-03 | 1.63 |
| TC1700008455.hg.1 | YPEL2 | yippee like 2 | 8.42E-04 | 1.63 |
| TC0700013602.hg.1 | POLR2J3 | polymerase (RNA) II (DNA directed) polypeptide J3 | 7.20E-04 | 1.63 |
| TC1700006638.hg.1 | ARRB2 | arrestin, beta 2 | 9.51E-03 | 1.63 |
| TC0500011150.hg.1 | GFM2 | G elongation factor, mitochondrial 2 | 7.22E-03 | 1.63 |
| TC1400009697.hg.1 | NPC2 | Niemann-Pick disease, type C2 | 6.12E-03 | 1.63 |
| TC2000006464.hg.1 | FAM110A | family with sequence similarity 110, member A | 6.09E-04 | 1.63 |
| TC1500010723.hg.1 | CHAC1 | ChaC glutathione-specific gamma-glutamylcyclotransferase 1 | 7.58E-03 | 1.63 |
| TC0100012490.hg.1 | C1orf233 | chromosome 1 open reading frame 233 | 5.95E-04 | 1.63 |
| TC0200007590.hg.1 | CHAC2 | ChaC, cation transport regulator homolog 2 (E. coli) | 2.78E-03 | 1.63 |
| TC0900012179.hg.1 | SPTAN1 | spectrin, alpha, non-erythrocytic 1 | 1.76E-03 | 1.63 |
| TC1900007100.hg.1 | MAST1 | microtubule associated serine/threonine kinase 1 | 9.42E-03 | 1.63 |
| TC0300010744.hg.1 | ACAA1 | acetyl-CoA acyltransferase 1 | 6.68E-03 | 1.63 |
| TC1000009880.hg.1 | NMT2 | N-myristoyltransferase 2 | 3.33E-04 | 1.63 |
| TC0200010837.hg.1 | ANKZF1 | ankyrin repeat and zinc finger domain containing 1 | 9.71E-03 | 1.63 |
| TC1700011919.hg.1 | CEP295NL | CEP295 N-terminal like | 5.41E-03 | 1.63 |
| HTA2-neg-47424241_st | NA | NA | 7.96E-03 | 1.63 |
| TC0400011180.hg.1 | SCD5 | stearoyl-CoA desaturase 5 | 2.65E-03 | 1.63 |
| TC0500007792.hg.1 | HEXB | hexosaminidase B (beta polypeptide) | 4.11E-03 | 1.63 |
| TC1000009894.hg.1 | FAM188A | family with sequence similarity 188, member A | 5.43E-03 | 1.63 |
| TC1900011724.hg.1 | ZNF540 | zinc finger protein 540 | 2.10E-03 | 1.63 |
| TC1900007049.hg.1 | ZNF440 | zinc finger protein 440 | 5.44E-03 | 1.63 |
| TC0900007547.hg.1 | C9orf85 | chromosome 9 open reading frame 85 | 8.91E-03 | 1.62 |
| TC0100012889.hg.1 | MAD2L2 | MAD2 mitotic arrest deficient-like 2 (yeast) | 3.33E-03 | 1.62 |
| TC0400008041.hg.1 | PTPN13 | protein tyrosine phosphatase, non-receptor type 13 (APO-1/CD95 (Fas)-associated phosphatase) | 7.63E-03 | 1.62 |
| TC0800006447.hg.1 | FBXO25 | F-box protein 25 | 1.25E-03 | 1.62 |
| TC1600010778.hg.1 | HYDIN | HYDIN, axonemal central pair apparatus protein | 5.80E-04 | 1.62 |
| TC0800009726.hg.1 | MTMR7 | myotubularin related protein 7 | 7.41E-04 | 1.62 |
| TC2000007435.hg.1 | IFT52 | intraflagellar transport 52 | 1.04E-03 | 1.62 |
| TC0100008897.hg.1 | WDR63 | WD repeat domain 63 | 3.78E-03 | 1.62 |
| TC0500008342.hg.1 | DCP2 | decapping mRNA 2 | 7.88E-04 | 1.62 |
| TC0700013219.hg.1 | LMBR1 | limb development membrane protein 1 | 1.20E-03 | 1.62 |
| TC1800007385.hg.1 | WDR7 | WD repeat domain 7 | 7.90E-04 | 1.62 |
| TC0700013337.hg.1 | UMAD1 | UBAP1-MVB12-associated (UMA) domain containing 1 | 4.62E-03 | 1.62 |
| TC0900011613.hg.1 | FAM102A | family with sequence similarity 102, member A | 8.47E-03 | 1.62 |
| TC1200012859.hg.1 | RHOF | ras homolog family member F (in filopodia) | 4.18E-03 | 1.62 |
| TC1100012269.hg.1 | ARHGAP20 | Rho GTPase activating protein 20 | 3.53E-03 | 1.62 |
| TC1000010699.hg.1 | IPMK | inositol polyphosphate multikinase | 4.48E-03 | 1.62 |
| TC1600011494.hg.1 | ARL6IP1 | ADP-ribosylation factor like GTPase 6 interacting protein 1 | 1.49E-03 | 1.62 |
| TC2000008530.hg.1 | DZANK1 | double zinc ribbon and ankyrin repeat domains 1 | 8.79E-04 | 1.62 |
| TC1600010633.hg.1 | ATP6V0D1 | ATPase, H+ transporting, lysosomal 38kDa, V0 subunit d1 | 2.30E-03 | 1.62 |
| TC1500006645.hg.1 | GABRG3 | gamma-aminobutyric acid (GABA) A receptor, gamma 3 | 3.50E-03 | 1.62 |
| TC0100017079.hg.1 | SLC41A1 | solute carrier family 41 (magnesium transporter), member 1 | 7.48E-03 | 1.62 |
| 23071942 | NA | NA | 5.21E-03 | 1.62 |
| TC1800006905.hg.1 | CABYR | calcium binding tyrosine-(Y)-phosphorylation regulated | 3.95E-03 | 1.62 |
| TC0900008482.hg.1 | SLC31A2 | solute carrier family 31 (copper transporter), member 2 | 2.32E-03 | 1.62 |
| TC1200007510.hg.1 | CCDC184 | coiled-coil domain containing 184 | 6.10E-04 | 1.62 |
| TC1100012509.hg.1 | SLC37A4 | solute carrier family 37 (glucose-6-phosphate transporter), member 4 | 5.34E-03 | 1.61 |

| | | | | |
|---|---|---|---|---|
| TC0900012294.hg.1 | TPRN | taperin | 3.54E-03 | 1.61 |
| TC1300006503.hg.1 | IFT88 | intraflagellar transport 88 | 2.46E-03 | 1.61 |
| TC2000009447.hg.1 | DPM1 | dolichyl-phosphate mannosyltransferase polypeptide 1, catalytic subunit | 6.33E-04 | 1.61 |
| TC0900011404.hg.1 | STOM | stomatin | 7.25E-04 | 1.61 |
| TC0500012948.hg.1 | SNCB | synuclein beta | 1.07E-03 | 1.61 |
| TC1900011766.hg.1 | MARK4 | MAP/microtubule affinity-regulating kinase 4 | 1.04E-03 | 1.61 |
| TC0100006650.hg.1 | AJAP1 | adherens junctions associated protein 1 | 9.10E-03 | 1.61 |
| TC1000011487.hg.1 | SORBS1 | sorbin and SH3 domain containing 1 | 3.43E-03 | 1.61 |
| TC1100009068.hg.1 | NCAM1 | neural cell adhesion molecule 1 | 1.03E-03 | 1.61 |
| TC1700010684.hg.1 | HAP1 | huntingtin-associated protein 1 | 6.54E-03 | 1.61 |
| TC1500008511.hg.1 | LRRC28 | leucine rich repeat containing 28 | 4.56E-03 | 1.61 |
| TC2100008560.hg.1 | DONSON | downstream neighbor of SON | 7.50E-04 | 1.61 |
| TC1500007659.hg.1 | PIAS1 | protein inhibitor of activated STAT 1 | 1.75E-03 | 1.61 |
| TC0700013429.hg.1 | PILRA | paired immunoglobin-like type 2 receptor alpha | 3.20E-03 | 1.61 |
| TC2100008534.hg.1 | PCBP3 | poly(rC) binding protein 3 | 2.77E-03 | 1.61 |
| TC0100011316.hg.1 | NFASC | neurofascin | 6.58E-04 | 1.61 |
| TC1100013065.hg.1 | KRTAP5-9 | keratin associated protein 5-9 | 5.53E-03 | 1.61 |
| TC0100016113.hg.1 | IGSF9 | immunoglobulin superfamily, member 9 | 3.25E-03 | 1.61 |
| TC0700010162.hg.1 | FBXL18 | F-box and leucine-rich repeat protein 18 | 5.49E-04 | 1.61 |
| TC0100013049.hg.1 | NBPF1 | neuroblastoma breakpoint family, member 1 | 1.67E-03 | 1.61 |
| TC0300013855.hg.1 | NFKBIZ | nuclear factor of kappa light polypeptide gene enhancer in B-cells inhibitor, zeta | 8.32E-03 | 1.60 |
| TC2200006552.hg.1 | TMEM191B | transmembrane protein 191B | 9.81E-03 | 1.60 |
| TC1900007342.hg.1 | FCHO1 | FCH domain only 1 | 2.52E-03 | 1.60 |
| TC0100006857.hg.1 | PTCHD2 | patched domain containing 2 | 8.02E-03 | 1.60 |
| TC1400010171.hg.1 | CCDC85C | coiled-coil domain containing 85C | 1.17E-03 | 1.60 |
| TC0900010390.hg.1 | ABHD17B | abhydrolase domain containing 17B | 8.45E-03 | 1.60 |
| TC0700012296.hg.1 | C7orf60 | chromosome 7 open reading frame 60 | 3.47E-03 | 1.60 |
| TC1500007005.hg.1 | ITPKA | inositol-trisphosphate 3-kinase A | 7.25E-04 | 1.60 |
| TC1500009951.hg.1 | GRAMD2 | GRAM domain containing 2 | 2.45E-03 | 1.60 |
| TC0200007032.hg.1 | HADHB | hydroxyacyl-CoA dehydrogenase/3-ketoacyl-CoA thiolase/enoyl-CoA hydratase (trifunctional protein), beta subunit | 1.07E-03 | 1.60 |
| TC0400008834.hg.1 | SCOC | short coiled-coil protein | 6.52E-03 | 1.60 |
| TC0X00008795.hg.1 | ABCD1 | ATP binding cassette subfamily D member 1 | 2.61E-03 | 1.60 |
| TC0100013889.hg.1 | ZMYND12 | zinc finger, MYND-type containing 12 | 9.57E-03 | 1.60 |
| TC0800007119.hg.1 | ESCO2 | establishment of sister chromatid cohesion N-acetyltransferase 2 | 2.73E-03 | 1.60 |
| TC0200016166.hg.1 | IQCA1 | IQ motif containing with AAA domain 1 | 5.32E-03 | 1.60 |
| HTA2-neg-47420640_st | NA | NA | 3.51E-03 | 1.60 |
| TC1900007087.hg.1 | WDR83 | WD repeat domain 83 | 1.03E-03 | 1.60 |
| TC1700012482.hg.1 | SIRT7 | sirtuin 7 | 4.05E-03 | 1.60 |
| HTA2-neg-47419576_st | NA | NA | 5.10E-03 | 1.60 |
| TC1600008454.hg.1 | GABARAPL2 | GABA(A) receptor-associated protein like 2 | 5.16E-04 | 1.59 |
| TC0100013995.hg.1 | EIF2B3 | eukaryotic translation initiation factor 2B, subunit 3 gamma, 58kDa | 8.51E-03 | 1.59 |
| TC1900006994.hg.1 | DNM2 | dynamin 2 | 1.65E-03 | 1.59 |
| TC1100008082.hg.1 | NPAS4 | neuronal PAS domain protein 4 | 4.51E-03 | 1.59 |
| TC1900007419.hg.1 | SLC25A42 | solute carrier family 25, member 42 | 9.59E-03 | 1.59 |
| TC2000007572.hg.1 | NCOA3 | nuclear receptor coactivator 3 | 7.86E-03 | 1.59 |
| HTA2-neg-47422675_st | NA | NA | 8.52E-03 | 1.59 |
| TC1200007864.hg.1 | NXPH4 | neurexophilin 4 | 9.20E-03 | 1.59 |
| TC0100018286.hg.1 | NBPF19 | neuroblastoma breakpoint family, member 19 | 8.50E-04 | 1.59 |
| TC1500006994.hg.1 | CHP1 | calcineurin-like EF-hand protein 1 | 5.81E-03 | 1.59 |
| TC0600008663.hg.1 | DOPEY1 | dopey family member 1 | 3.90E-03 | 1.59 |
| TC1500009781.hg.1 | UBAP1L | ubiquitin associated protein 1 like | 3.35E-03 | 1.59 |
| TC1000006796.hg.1 | SEC61A2 | Sec61 translocon alpha 2 subunit | 1.93E-03 | 1.59 |
| TC1900008416.hg.1 | INAFM1 | InaF-motif containing 1 | 4.89E-03 | 1.59 |
| TC0400010626.hg.1 | FRYL | FRY like transcription coactivator | 4.55E-03 | 1.59 |
| TC1800008952.hg.1 | DSEL | dermatan sulfate epimerase-like | 5.40E-03 | 1.59 |
| TC0200008534.hg.1 | ANKRD36 | ankyrin repeat domain 36 | 3.05E-03 | 1.59 |
| TC1900010701.hg.1 | C19orf47 | chromosome 19 open reading frame 47 | 5.99E-03 | 1.59 |
| TC0700012086.hg.1 | POLR2J | polymerase (RNA) II (DNA directed) polypeptide J, 13.3kDa | 7.39E-04 | 1.59 |
| HTA2-pos-PSR02018287.hg.1 | NA | NA | 2.53E-03 | 1.59 |
| TC1000007698.hg.1 | CISD1 | CDGSH iron sulfur domain 1 | 6.84E-03 | 1.59 |
| TC0100014175.hg.1 | EPS15 | epidermal growth factor receptor pathway substrate 15 | 3.40E-03 | 1.59 |
| TC1100009241.hg.1 | HINFP | histone H4 transcription factor | 3.62E-03 | 1.59 |
| TC1100013061.hg.1 | NADSYN1 | NAD synthetase 1 | 5.39E-03 | 1.59 |
| TC0900008793.hg.1 | ZBTB34 | zinc finger and BTB domain containing 34 | 1.50E-03 | 1.59 |
| TC0300010889.hg.1 | ZNF445 | zinc finger protein 445 | 8.74E-03 | 1.59 |
| TC1700006735.hg.1 | SLC2A4 | solute carrier family 2 (facilitated glucose transporter), member 4 | 3.06E-03 | 1.59 |
| TC0200016626.hg.1 | MBOAT2 | membrane bound O-acyltransferase domain containing 2 | 2.10E-03 | 1.58 |
| TC1900011029.hg.1 | MEIS3 | Meis homeobox 3 | 3.37E-03 | 1.58 |
| TC1500007546.hg.1 | SNX22 | sorting nexin 22 | 9.23E-04 | 1.58 |
| TC0800008032.hg.1 | ZC2HC1A | zinc finger, C2HC-type containing 1A | 5.56E-03 | 1.58 |
| TC1100009858.hg.1 | CHRNA10 | cholinergic receptor, nicotinic alpha 10 | 2.88E-03 | 1.58 |
| 23075426 | NA | NA | 6.35E-03 | 1.58 |
| TC1800006650.hg.1 | RAB31 | RAB31, member RAS oncogene family | 4.46E-03 | 1.58 |
| TC0100018224.hg.1 | FAM183A | family with sequence similarity 183, member A | 9.35E-03 | 1.58 |
| TC0100017693.hg.1 | PGBD5 | piggyBac transposable element derived 5 | 9.35E-04 | 1.58 |
| TC2200008831.hg.1 | PMM1 | phosphomannomutase 1 | 4.65E-03 | 1.58 |
| TC1200012707.hg.1 | ALDH2 | aldehyde dehydrogenase 2 family (mitochondrial) | 2.35E-03 | 1.58 |
| TC0200016039.hg.1 | PDE6D | phosphodiesterase 6D, cGMP-specific, rod, delta | 2.84E-03 | 1.58 |
| TC1600010500.hg.1 | GOT2 | glutamic-oxaloacetic transaminase 2, mitochondrial | 8.41E-03 | 1.58 |
| TC0100007963.hg.1 | EXO5 | exonuclease 5 | 5.14E-03 | 1.58 |
| TC0900011705.hg.1 | ASB6 | ankyrin repeat and SOCS box containing 6 | 2.60E-03 | 1.58 |
| TSUnmapped00000027.hg.1 | MLXIP | MLX interacting protein | 6.93E-03 | 1.58 |
| TC0X00007310.hg.1 | TSPYL2 | TSPY-like 2 | 1.26E-03 | 1.58 |
| TC1000012050.hg.1 | FGFR2 | fibroblast growth factor receptor 2 | 2.99E-03 | 1.58 |
| TC1000007564.hg.1 | MAPK8 | mitogen-activated protein kinase 8 | 1.27E-03 | 1.57 |
| TC1900011806.hg.1 | ZNF582-AS1 | ZNF582 antisense RNA 1 (head to head) | 2.65E-03 | 1.57 |
| TC1500008514.hg.1 | LRRC28 | Transcript Identified by AceView, Entrez Gene ID(s) 123355 | 2.91E-03 | 1.57 |
| TC0300011815.hg.1 | DCBLD2 | discoidin, CUB and LCCL domain containing 2 | 1.54E-03 | 1.57 |
| TSUnmapped00000166.hg.1 | LRP6 | LDL receptor related protein 6 | 3.85E-03 | 1.57 |
| TC1000012549.hg.1 | PARG | poly (ADP-ribose) glycohydrolase | 8.61E-03 | 1.57 |
| TC0200008569.hg.1 | INPP4A | inositol polyphosphate-4-phosphatase type I A | 1.70E-03 | 1.57 |
| TC2000009886.hg.1 | PANK2 | pantothenate kinase 2 | 1.16E-03 | 1.57 |
| TC1500010319.hg.1 | KLHL25 | kelch-like family member 25 | 3.54E-03 | 1.57 |
| TC0600014101.hg.1 | MICA | MHC class I polypeptide-related sequence A | 9.39E-03 | 1.57 |
| TC1400007566.hg.1 | SMOC1 | SPARC related modular calcium binding 1 | 8.41E-04 | 1.57 |
| TC1000009873.hg.1 | DCLRE1C | DNA cross-link repair 1C | 2.74E-03 | 1.57 |
| TC0600010331.hg.1 | FAM120B | family with sequence similarity 120B | 5.39E-03 | 1.57 |
| TC0500013336.hg.1 | SSBP2 | single-stranded DNA binding protein 2 | 3.29E-03 | 1.57 |
| TC1000012577.hg.1 | LIPA | lipase A, lysosomal acid, cholesterol esterase | 2.35E-03 | 1.57 |
| HTA2-pos-PSR05025584.hg.1 | NA | NA | 5.09E-03 | 1.57 |
| 23071963 | NA | NA | 6.13E-03 | 1.57 |
| TC1500010796.hg.1 | SYNM | synemin, intermediate filament protein | 9.47E-03 | 1.57 |
| TC1900010050.hg.1 | SUGP1 | SURP and G-patch domain containing 1 | 6.20E-03 | 1.57 |
| TC0800009619.hg.1 | CTSB | cathepsin B | 1.76E-03 | 1.57 |
| TC0200015402.hg.1 | FAM126B | family with sequence similarity 126, member B | 5.61E-03 | 1.57 |
| TC0600008957.hg.1 | BVES-AS1 | BVES antisense RNA 1 | 1.67E-03 | 1.57 |
| TC0200012252.hg.1 | STRN | striatin, calmodulin binding protein | 3.55E-03 | 1.57 |
| TC1900009304.hg.1 | PIP5K1C | Zhang2013 ALT_ACCEPTOR, ALT_DONOR, coding, INTERNAL, intronic best transcript NM_012398 | 2.38E-03 | 1.57 |
| TC0200014192.hg.1 | SAP130 | Sin3A associated protein 130kDa | 2.32E-03 | 1.57 |
| TC0X00009256.hg.1 | KLHL15 | kelch-like family member 15 | 1.89E-03 | 1.57 |
| TC0800007870.hg.1 | CSPP1 | centrosome and spindle pole associated protein 1 | 1.19E-03 | 1.57 |
| TC0500012153.hg.1 | FAM13B | family with sequence similarity 13, member B | 9.65E-03 | 1.57 |
| TC0100011698.hg.1 | DISP1 | dispatched homolog 1 (Drosophila) | 2.37E-03 | 1.57 |
| TC1100013040.hg.1 | TM7SF2 | transmembrane 7 superfamily member 2 | 1.43E-03 | 1.56 |
| TC0100014627.hg.1 | ST6GALNAC3 | Memczak2013 ANTISENSE, coding, INTERNAL, intronic best transcript NM_152996 | 1.83E-03 | 1.56 |
| TC0700013598.hg.1 | LRCH4 | leucine-rich repeats and calponin homology (CH) domain containing 4 | 1.21E-03 | 1.56 |
| TC0100006865.hg.1 | AGTRAP | angiotensin II receptor-associated protein | 3.72E-03 | 1.56 |
| TC0700008450.hg.1 | LMTK2 | lemur tyrosine kinase 2 | 4.39E-03 | 1.56 |
| 23071896 | NA | NA | 6.95E-03 | 1.56 |

| | | | | |
|---|---|---|---|---|
| TC0200011125.hg.1 | ATG16L1 | autophagy related 16-like 1 | 3.53E-03 | 1.56 |
| TC1700010565.hg.1 | FBXL20 | F-box and leucine-rich repeat protein 20 | 1.29E-03 | 1.56 |
| TC0600009980.hg.1 | ZDHHC14 | zinc finger, DHHC-type containing 14 | 5.54E-03 | 1.56 |
| TC2000007666.hg.1 | RNF114 | ring finger protein 114 | 3.65E-03 | 1.56 |
| TC1400008415.hg.1 | ZFYVE21 | zinc finger, FYVE domain containing 21 | 1.69E-03 | 1.56 |
| TC0400012992.hg.1 | MFAP3L | microfibrillar associated protein 3 like | 8.84E-03 | 1.56 |
| TC2200008907.hg.1 | SCUBE1 | signal peptide, CUB domain, EGF-like 1 | 7.52E-03 | 1.56 |
| TC0100018249.hg.1 | AMY2B | amylase, alpha 2B (pancreatic) | 7.38E-03 | 1.56 |
| TC0100018242.hg.1 | HS2ST1 | heparan sulfate 2-O-sulfotransferase 1 | 4.42E-03 | 1.56 |
| TC1700008984.hg.1 | SEC14L1 | SEC14-like lipid binding 1 | 1.17E-03 | 1.56 |
| TC1600009958.hg.1 | NPIPB4 | nuclear pore complex interacting protein family, member B4 | 2.57E-03 | 1.56 |
| TC0100012465.hg.1 | CPSF3L | cleavage and polyadenylation specific factor 3-like | 2.67E-03 | 1.56 |
| TC1100010274.hg.1 | SPTY2D1 | SPT2 chromatin protein domain containing 1 | 3.61E-03 | 1.56 |
| TC0300009468.hg.1 | PRKCI | protein kinase C, iota | 1.34E-03 | 1.55 |
| 23075525 | NA | NA | 2.94E-03 | 1.55 |
| TC1800007518.hg.1 | ZCCHC2 | zinc finger, CCHC domain containing 2 | 7.52E-03 | 1.55 |
| TC0X00007012.hg.1 | ATP6AP2 | ATPase, H+ transporting, lysosomal accessory protein 2 | 9.88E-03 | 1.55 |
| TC0100014528.hg.1 | DIRAS3 | DIRAS family, GTP-binding RAS-like 3 | 7.72E-03 | 1.55 |
| TC1700012183.hg.1 | FGF11 | fibroblast growth factor 11 | 8.03E-03 | 1.55 |
| TC1500009200.hg.1 | TTBK2 | tau tubulin kinase 2 | 8.21E-03 | 1.55 |
| TC1000009152.hg.1 | HTRA1 | HtrA serine peptidase 1 | 6.76E-03 | 1.55 |
| TC0300008904.hg.1 | PPP2R3A | protein phosphatase 2, regulatory subunit B, alpha | 3.65E-03 | 1.55 |
| TC1000011948.hg.1 | PDZD8 | PDZ domain containing 8 | 3.87E-03 | 1.55 |
| HTA2-neg-47422968_st | NA | NA | 6.81E-03 | 1.55 |
| TC0X00007190.hg.1 | PORCN | porcupine homolog (Drosophila) | 2.68E-03 | 1.55 |
| TC2000010008.hg.1 | NFS1 | NFS1 cysteine desulfurase | 3.21E-03 | 1.55 |
| HTA2-pos-PSR02018288.hg.1 | NA | NA | 2.01E-03 | 1.55 |
| TC1100013206.hg.1 | STARD10 | StAR-related lipid transfer domain containing 10 | 5.15E-03 | 1.55 |
| HTA2-pos-PSR02018286.hg.1 | NA | NA | 3.50E-03 | 1.55 |
| TC1200006604.hg.1 | CD9 | CD9 molecule | 8.02E-03 | 1.55 |
| TC1600011445.hg.1 | SPG7 | spastic paraplegia 7 (pure and complicated autosomal recessive) | 5.36E-03 | 1.55 |
| TC1200008147.hg.1 | CCT2 | chaperonin containing TCP1, subunit 2 (beta) | 2.53E-03 | 1.55 |
| TC0300009282.hg.1 | TIPARP | TCDD-inducible poly(ADP-ribose) polymerase | 3.81E-03 | 1.55 |
| TC1200010252.hg.1 | TMTC1 | transmembrane and tetratricopeptide repeat containing 1 | 2.18E-03 | 1.55 |
| TC0800006866.hg.1 | VPS37A | vacuolar protein sorting 37 homolog A (S. cerevisiae) | 5.53E-03 | 1.55 |
| TC1200007567.hg.1 | KCNH3 | potassium channel, voltage gated eag related subfamily H, member 3 | 3.81E-03 | 1.55 |
| TC0800011881.hg.1 | NDRG1 | N-myc downstream regulated 1 | 2.49E-03 | 1.55 |
| TC1700008846.hg.1 | KIF19 | kinesin family member 19 | 7.11E-03 | 1.55 |
| TC0X00008521.hg.1 | SLC9A6 | solute carrier family 9, subfamily A (NHE6, cation proton antiporter 6), member 6 | 5.07E-03 | 1.54 |
| TC0300013877.hg.1 | EPHB1 | EPH receptor B1 | 2.58E-03 | 1.54 |
| TC1600009533.hg.1 | MYH11 | myosin, heavy chain 11, smooth muscle | 4.56E-03 | 1.54 |
| TC1500010687.hg.1 | OR4M2 | olfactory receptor, family 4, subfamily M, member 2 | 5.81E-03 | 1.54 |
| TC1500006726.hg.1 | ULK4P1 | ULK4 pseudogene 1 | 2.22E-03 | 1.54 |
| TC1000006639.hg.1 | FBXO18 | F-box protein, helicase, 18 | 2.60E-03 | 1.54 |
| TC1200012256.hg.1 | ABCB9 | ATP binding cassette subfamily B member 9 | 2.49E-03 | 1.54 |
| TC1100007030.hg.1 | NAV2 | neuron navigator 2 | 3.81E-03 | 1.54 |
| TC2100007039.hg.1 | DOPEY2 | dopey family member 2 | 8.93E-03 | 1.54 |
| TC1500008309.hg.1 | ZNF774 | zinc finger protein 774 | 2.31E-03 | 1.54 |
| TC1100011629.hg.1 | XRRA1 | X-ray radiation resistance associated 1 | 9.65E-03 | 1.54 |
| 23074628 | NA | NA | 3.39E-03 | 1.54 |
| TC1000012162.hg.1 | UROS | uroporphyrinogen III synthase | 7.55E-03 | 1.54 |
| TC0900008156.hg.1 | NCBP1 | nuclear cap binding protein subunit 1 | 9.88E-04 | 1.54 |
| TC1100010077.hg.1 | SBF2 | SET binding factor 2 | 2.02E-03 | 1.54 |
| TC2200008856.hg.1 | NAGA | N-acetylgalactosaminidase, alpha- | 7.83E-03 | 1.54 |
| TC0900010607.hg.1 | AGTPBP1 | ATP/GTP binding protein 1 | 4.99E-03 | 1.54 |
| TC1300007861.hg.1 | PCCA | propionyl-CoA carboxylase alpha subunit | 2.25E-03 | 1.54 |
| 23075826 | NA | NA | 1.86E-03 | 1.54 |
| TC0200015082.hg.1 | FKBP7 | FK506 binding protein 7 | 9.95E-03 | 1.54 |
| TC1200012663.hg.1 | RAB3IP | RAB3A interacting protein | 2.38E-03 | 1.54 |
| TC0100009646.hg.1 | NBPF26 | neuroblastoma breakpoint family, member 26 | 1.79E-03 | 1.54 |
| TC1700010828.hg.1 | FAM171A2 | family with sequence similarity 171, member A2 | 7.79E-03 | 1.54 |
| TC1600011346.hg.1 | SEC14L5 | SEC14-like lipid binding 5 | 1.76E-03 | 1.54 |
| TC0700013502.hg.1 | SNX8 | sorting nexin 8 | 2.53E-03 | 1.54 |
| TC1900011007.hg.1 | AP2S1 | adaptor-related protein complex 2 sigma 1 subunit | 4.90E-03 | 1.54 |
| 23076566 | NA | NA | 3.24E-03 | 1.54 |
| TC0300009004.hg.1 | RASA2 | RAS p21 protein activator 2 | 2.83E-03 | 1.54 |
| TC1900007988.hg.1 | SIPA1L3 | signal-induced proliferation-associated 1 like 3 | 3.36E-03 | 1.54 |
| TC0200012068.hg.1 | SLC30A3 | solute carrier family 30 (zinc transporter), member 3 | 3.68E-03 | 1.53 |
| TC0X00006697.hg.1 | SYAP1 | synapse associated protein 1 | 2.53E-03 | 1.53 |
| TC2100007996.hg.1 | TMEM50B | transmembrane protein 50B | 3.61E-03 | 1.53 |
| TC0900009076.hg.1 | STKLD1 | serine/threonine kinase-like domain containing 1 | 4.34E-03 | 1.53 |
| TC0400012857.hg.1 | RAPGEF2 | Rap guanine nucleotide exchange factor 2 | 1.61E-03 | 1.53 |
| TC0100006983.hg.1 | CTRC | chymotrypsin C (caldecrin) | 5.93E-03 | 1.53 |
| TC0500012523.hg.1 | ATOX1 | antioxidant 1 copper chaperone | 1.82E-03 | 1.53 |
| TC1500010906.hg.1 | RP11-351M8.2 | --- | 4.34E-03 | 1.53 |
| TC0400010242.hg.1 | PPARGC1A | peroxisome proliferator-activated receptor gamma, coactivator 1 alpha | 3.67E-03 | 1.53 |
| TC0700012584.hg.1 | UBE2H | ubiquitin conjugating enzyme E2H | 7.36E-03 | 1.53 |
| TC1100007865.hg.1 | METTL12 | methyltransferase like 12 | 6.29E-03 | 1.53 |
| TC2000009604.hg.1 | PMEPA1 | prostate transmembrane protein, androgen induced 1 | 6.67E-03 | 1.53 |
| TC1700008331.hg.1 | HLF | hepatic leukemia factor | 4.80E-03 | 1.53 |
| TC0700009560.hg.1 | CNTNAP2 | contactin associated protein-like 2 | 1.33E-03 | 1.53 |
| TC1600007210.hg.1 | HS3ST2 | heparan sulfate (glucosamine) 3-O-sulfotransferase 2 | 9.64E-03 | 1.53 |
| TC1300007774.hg.1 | MBNL2 | muscleblind-like splicing regulator 2 | 8.89E-03 | 1.53 |
| TC1400010169.hg.1 | SETD3 | SET domain containing 3 | 6.56E-03 | 1.53 |
| TC1600011399.hg.1 | MT1X | metallothionein 1X | 3.79E-03 | 1.53 |
| TC1700010248.hg.1 | GIT1 | G protein-coupled receptor kinase interacting ArfGAP 1 | 2.54E-03 | 1.53 |
| TC1900010997.hg.1 | DACT3 | dishevelled-binding antagonist of beta-catenin 3 | 3.95E-03 | 1.53 |
| TC0300009701.hg.1 | PSMD2 | proteasome 26S subunit, non-ATPase 2 | 1.25E-03 | 1.53 |
| TC0500013350.hg.1 | RAPGEF6 | Rap guanine nucleotide exchange factor 6 | 2.29E-03 | 1.53 |
| TC1700010811.hg.1 | HDAC5 | histone deacetylase 5 | 4.84E-03 | 1.53 |
| TC0400012792.hg.1 | NIPAL1 | NIPA-like domain containing 1 | 7.09E-03 | 1.53 |
| TC1000007007.hg.1 | ARMC3 | armadillo repeat containing 3 | 2.70E-03 | 1.53 |
| TC1000011556.hg.1 | AVPI1 | arginine vasopressin-induced 1 | 5.50E-03 | 1.53 |
| TC1100007906.hg.1 | RTN3 | reticulon 3 | 1.22E-03 | 1.52 |
| TC0800007431.hg.1 | AP3M2 | adaptor-related protein complex 3, mu 2 subunit | 2.87E-03 | 1.52 |
| TC1700007787.hg.1 | WIPF2 | WAS/WASL interacting protein family, member 2 | 2.23E-03 | 1.52 |
| TC0600010853.hg.1 | GFOD1 | glucose-fructose oxidoreductase domain containing 1 | 1.72E-03 | 1.52 |
| TC1300007158.hg.1 | MLNR | motilin receptor | 3.48E-03 | 1.52 |
| TC1900008245.hg.1 | ZNF221 | zinc finger protein 221 | 7.05E-03 | 1.52 |
| TC0300010739.hg.1 | PLCD1 | phospholipase C, delta 1 | 5.95E-03 | 1.52 |
| TC0500007868.hg.1 | SCAMP1 | secretory carrier membrane protein 1 | 5.19E-03 | 1.52 |
| TC0200011130.hg.1 | DGKD | diacylglycerol kinase, delta 130kDa | 1.29E-03 | 1.52 |
| 23075655 | NA | NA | 4.15E-03 | 1.52 |
| TC1100008502.hg.1 | LOC100506127 | putative uncharacterized protein FLJ37770-like | 4.42E-03 | 1.52 |
| TC1200008924.hg.1 | DTX1 | deltex 1, E3 ubiquitin ligase | 5.17E-03 | 1.52 |
| TC0100018235.hg.1 | FPGT-TNNI3K | FPGT-TNNI3K readthrough | 3.27E-03 | 1.52 |
| TC0500009061.hg.1 | PCYOX1L | prenylcysteine oxidase 1 like | 5.20E-03 | 1.52 |
| HTA2-neg-47419616_st | NA | NA | 7.38E-03 | 1.52 |
| TC1900011690.hg.1 | ZNF493 | zinc finger protein 493 | 8.34E-03 | 1.52 |
| TC1000009865.hg.1 | FAM107B | Memczak2013 ALT_ACCEPTOR, ALT_DONOR, coding, INTERNAL, intronic best transcript NM_031453 | 1.76E-03 | 1.52 |
| TC1900009628.hg.1 | CDC37 | cell division cycle 37 | 6.04E-03 | 1.52 |
| TC0600009683.hg.1 | AIG1 | androgen-induced 1 | 9.90E-03 | 1.52 |
| TC1200009959.hg.1 | LRP6 | LDL receptor related protein 6 | 6.14E-03 | 1.52 |
| TC1600008869.hg.1 | ZNF778 | zinc finger protein 778 | 9.42E-03 | 1.51 |
| TC0200008263.hg.1 | USP39 | ubiquitin specific peptidase 39 | 3.14E-03 | 1.51 |
| TC0200013863.hg.1 | NPHP1 | nephronophthisis 1 (juvenile) | 7.14E-03 | 1.51 |
| TC1300008338.hg.1 | MIPEP | mitochondrial intermediate peptidase | 9.55E-03 | 1.51 |
| TC1000012117.hg.1 | CHST15 | carbohydrate (N-acetylgalactosamine 4-sulfate 6-O) sulfotransferase 15 | 7.87E-03 | 1.51 |
| TC0500013320.hg.1 | GTF2H2 | general transcription factor IIH subunit 2 | 9.78E-03 | 1.51 |

| | | | | |
|---|---|---|---|---|
| TC0X00006590.hg.1 | WWC3 | WWC family member 3 | 6.69E-03 | 1.51 |
| TC1500009753.hg.1 | OAZ2 | ornithine decarboxylase antizyme 2 | 3.76E-03 | 1.51 |
| TC0200016606.hg.1 | SCLY | selenocysteine lyase | 3.86E-03 | 1.51 |
| TC0700012427.hg.1 | FAM3C | family with sequence similarity 3, member C | 1.70E-03 | 1.51 |
| TC2200007278.hg.1 | KCTD17 | potassium channel tetramerization domain containing 17 | 7.94E-03 | 1.51 |
| TC1600006619.hg.1 | PDPK1 | 3-phosphoinositide dependent protein kinase 1 | 9.36E-03 | 1.51 |
| TC0800009237.hg.1 | GPAA1 | glycosylphosphatidylinositol anchor attachment 1 | 6.70E-03 | 1.51 |
| TC0600011725.hg.1 | GLO1 | glyoxalase I | 2.49E-03 | 1.51 |
| TC0100007493.hg.1 | TRNP1 | TMF1-regulated nuclear protein 1 | 1.64E-03 | 1.51 |
| TC0200016562.hg.1 | PHOSPHO2 | phosphatase, orphan 2 | 3.28E-03 | 1.51 |
| TC0100010267.hg.1 | DUSP23 | dual specificity phosphatase 23 | 1.40E-03 | 1.51 |
| HTA2-pos-3374492_st | NA | NA | 7.37E-03 | 1.51 |
| TC1200010292.hg.1 | AMN1 | antagonist of mitotic exit network 1 homolog | 8.15E-03 | 1.51 |
| TSUnmapped00000399.hg.1 | ZNF852 | zinc finger protein 852 | 1.87E-03 | 1.51 |
| TC1900011194.hg.1 | KCNC3 | potassium channel, voltage gated Shaw related subfamily C, member 3 | 7.42E-03 | 1.51 |
| TC0400007938.hg.1 | BMP2K | BMP2 inducible kinase | 1.59E-03 | 1.51 |
| TC1700008844.hg.1 | TTYH2 | tweety family member 2 | 6.76E-03 | 1.51 |
| TC0300008242.hg.1 | ALCAM | activated leukocyte cell adhesion molecule | 7.60E-03 | 1.51 |
| TC1900008555.hg.1 | AP2A1 | adaptor-related protein complex 2, alpha 1 subunit | 7.95E-03 | 1.51 |
| TC0600007038.hg.1 | JARID2 | jumonji, AT rich interactive domain 2 | 1.90E-03 | 1.51 |
| TC1400007172.hg.1 | STYX | serine/threonine/tyrosine interacting protein | 4.41E-03 | 1.50 |
| HTA2-pos-PSR05025585.hg.1 | NA | NA | 7.29E-03 | 1.50 |
| TC0300013635.hg.1 | ACAP2 | ArfGAP with coiled-coil, ankyrin repeat and PH domains 2 | 3.22E-03 | 1.50 |
| TC0500009622.hg.1 | GRK6 | G protein-coupled receptor kinase 6 | 6.70E-03 | 1.50 |
| TC0X00011413.hg.1 | L1CAM | L1 cell adhesion molecule | 2.23E-03 | 1.50 |
| TC0800008371.hg.1 | SPAG1 | sperm associated antigen 1 | 5.38E-03 | 1.50 |
| TC1100007876.hg.1 | TMEM179B | transmembrane protein 179B | 4.04E-03 | 1.50 |
| TC1200007686.hg.1 | KRT18 | keratin 18, type I | 1.30E-03 | 1.50 |
| TC0600013531.hg.1 | NUP43 | nucleoporin 43kDa | 2.42E-03 | 1.50 |
| TC1700012308.hg.1 | LLGL2 | lethal giant larvae homolog 2 (Drosophila) | 9.14E-03 | 1.50 |
| TC0700010887.hg.1 | COA1 | cytochrome c oxidase assembly factor 1 homolog | 9.58E-03 | 1.50 |
